# Peopling of Tibet Plateau and multiple waves of admixture of Tibetans inferred from both modern and ancient genome-wide data

**DOI:** 10.1101/2020.07.03.185884

**Authors:** Mengge Wang, Xing Zou, Hui-Yuan Ye, Zheng Wang, Yan Liu, Jing Liu, Fei Wang, Hongbin Yao, Pengyu Chen, Ruiyang Tao, Shouyu Wang, Lan-Hai Wei, Renkuan Tang, Chuan-Chao Wang, Guanglin He

## Abstract

Archeologically attested human occupation on the Tibet Plateau (TP) can be traced back to 160 thousand years ago (kya, Xiahe) via archaic people and 30~40 kya via anatomically modern human in Nwya Devu. However, the past human movements and peopling of the TP keep in its infancy in the modern/ancient DNA studies. Here, we performed the first modern/ancient genomic meta-analysis among 3,017 Paleolithic to present-day eastern Eurasian genomes (2,444 modern individuals from 183 populations (including 98 Ü-Tsang/Ando/Kham Tibetans) and 573 ancients (including 161 Chinese ancients first meta-analyzed here)). Closer genetic connection between ancient-modern highland Tibetans and lowland island/coastal Neolithic northern East Asians was identified, reflecting the main ancestry of high-altitude Tibeto-Burman speakers originated from the ancestors of Houli/Yangshao/Longshan ancients in the middle and lower Yellow River basin, consistent with the common North-China origin of Sino-Tibetan language and dispersal pattern of millet farmers. Although the shared common northern East Asian lineage between Tibetans and lowland East Asians, we still identified genetic differentiation between Highlanders and lowland northern East Asians, the former harboring more deeply diverged Hoabinhian/Onge ancestry and the latter possessing more modern Neolithic southern East Asian and Siberian ancestry, which suggested the co-existence of Paleolithic and Neolithic ancestries in modern and Neolithic East Asian Highlanders. Tibetans from Ü-Tsang/Ando/Kham Tibetan regions showed strong population stratifications consistent with their cultural backgrounds and geographic terrains (showed as barriers for human movements): stronger Chokhopani affinity in Ü-Tsang Tibetans, more western Eurasian ancestry in Ando and greater Neolithic southern East Asian ancestry in Kham Tibetan. Modern combined ancient genomes documented multiple waves of human migrations in TP past: the first layer of local Hunter-Gatherer mixed with Qijia Farmer arose the Chokhopani-associated Proto-Tibetan-Burman, admixture with the additional genetic materials from the western Eurasian steppe, Yellow River and Yangtze River respectively gave rise to modern Ando, Ü-Tsang and Kham Tibetans.

## Introduction

The Tibet Plateau (TP), widely known as the Third Pole of the world, forms the high-altitude core of Asia with an average elevation of more than 4,000 meters above sea level (masl) and represents one of the most demanding environments for human settlement due to perennial low temperatures, extreme aridity, and severe hypoxia. However, archeological and genetic studies have indicated that archaic hominins occupied the TP had well adapted to the high-altitude hypoxic environment long before the arrival of modern *Homo sapiens* and present-day Tibetan Highlanders have adapted uniquely to extreme high-altitude conditions since the initial colonization of the TP(Qi, et al. 2013; Jeong, et al. 2016; Gnecchi-Ruscone, et al. 2018; Chen, Welker, et al. 2019). Besides, recent linguistic evidence suggested that Tibeto-Burman populations diverged from Han Chinese with an average coalescence age of approximately 5.9 thousand years ago (kya). At present, over seven million indigenous Tibetans (2016 census) have settled in the Plateau and are successfully adapted to the high-altitude hypoxic environment. Genomic evidence supported that multiple variants may jointly deliver the fitness of the modern Tibetans on the TP, and Denisovan introgression into modern Tibetans and surrounding populations including positively selected haplotypes of *HIF-1α prolyl hydroxylase1 (EGLN1)* and *Endothelial PAS domain protein 1 (EPAS1)* is significantly associated with the high-altitude adaptation to hypoxia(Simonson, et al. 2010; Xu, et al. 2011; Xiang, et al. 2013; Huerta-Sanchez, et al. 2014; Lu, et al. 2016; Gnecchi-Ruscone, et al. 2018; Chen, Welker, et al. 2019; Deng, et al. 2019). Compared to other parts of East Asia(Reich 2018; Ning, et al. 2019; Jeong, et al. 2020; Ning, et al. 2020; Wang, Yeh, et al. 2020; Yang, et al. 2020), the greatest problem facing researchers is the lack of excavated archaeological sites on the TP, which means that certain types of critical data, such as zooarchaeological and archaeobotanical data for reconstructing the subsistence strategy, ancient DNA (aDNA) for dissecting the genomic correlation between ancient individuals and modern Tibetan-like Highlanders, are in short supply.

To date, whence and how the early human colonizers conquered the TP and who modern Tibetans descended from are two key questions that remain to be solved, however, archaeological, paleoanthropological and genetic researches on the peopling of the TP and demographic history of Tibetan Highlanders are still in a developmental stage. As revealed by archaeological evidence, handprints and footprints of *Homo sapiens* found in the southern TP (Quesang site) at 4,200 masl suggested that the TP retains traces of an intermittent human presence from at least 20 kya(Zhang and Li 2002), but some scholars supporting at the early Holocene(Meyer, et al. 2017). The Nwya Devu site, located nearly 4,600 masl in central Tibet, could be dated to at least 30 kya, which deepens considerably the history of the peopling of the TP and the antiquity of human high-altitude adaptations(Zhang, et al. 2018). The palaeoproteomic analysis of a Xiahe Denisovan mandible indicated that the prehistoric colonization of archaic hominins on the TP could be traced back to the Middle Pleistocene epoch (around 160 kya)(Chen, Welker, et al. 2019). Additionally, genomic evidence strongly suggested that modern humans did exist on the TP before the Last Glacial Maximum (LGM), and the existence of genetic relics of the Upper Paleolithic inhabitants in modern Tibetans indicated some genetic continuity between the initial Paleolithic settlers and modern Tibetan Highlanders(Zhao, et al. 2009; Qin, et al. 2010; Qi, et al. 2013; Li, et al. 2015; Lu, et al. 2016). The archaeogenetic investigation of prehistoric Himalayan populations provided supporting evidence for the high-elevation East Asian origin of the first inhabitants of the Himalayas, indirectly indicating the pre-Neolithic human activities on the TP(Jeong, et al. 2016).

In contrast to the Late Pleistocene hunter-gatherer colonization, the timing and dynamics of the permanent human occupation of the TP have provoked much debate(Aldenderfer 2011; Qi, et al. 2013; Chen, et al. 2015; d’Alpoim Guedes 2015; Lu 2016; Rhode 2016; Hu, et al. 2019; Li, Tian, et al. 2019a; Ren, et al. 2020). Archaeological and genomic findings revealed that the permanent settlement of the TP was a relatively recent occurrence that coincided with the establishment of farming and pastoralism on the Plateau(Aldenderfer 2011; Qi, et al. 2013; Chen, et al. 2015; Lu 2016; Li, Tian, et al. 2019a). Chen et al. reported archaeobotanical and zooarchaeological data from 53 archaeological sites in the northeastern TP (NETP) and illustrated that the novel agropastoral subsistence strategy facilitated year-round living on the TP after 3.6 kya(Chen, et al. 2015). The first comprehensive and in-depth genomic investigation of the Tibet sheep revealed a stepwise pattern of permanent human occupation on the TP through the Tang-Bo Ancient Road (~3,100 years ago, from northern China to the NETP; and ~1,300 years ago, from the NETP to southwestern areas of the TP)(Hu, et al. 2019). However, it remains unknown who brought the cold-tolerant barley agriculture and livestock to the TP, and how indigenous foragers interacted with incoming farmers remains unclear. The archaeological observations demonstrated that incoming farmer groups did not replace the local foragers, and the two populations co-existed for extended periods(Gao, et al. 2020; Ren, et al. 2020). The mitochondrial evidence and radiocarbon dates of the cereal remains revealed that millet farmers adopted and brought barley agriculture to the TP around 3.6–3.3 kya, and contemporary Tibetans could trace their ancestry back to the Neolithic millet farmers(Li, Tian, et al. 2019a). Xu et al. conducted a series of typical population genomic studies focused on the demographic history of modern Tibetans and other high-altitude adaptive Highlanders and concluded that Tibetans arose from a mixture of multiple ancestral gene pools and Paleolithic and Neolithic ancestries co-existed in the Tibetan gene pool(Lu, et al. 2016). Moreover, the studies of genetic variations of modern Tibetan groups have also been performed based on forensically available markers(He, Wang, Su, et al. 2018; He, Wang, Zou, et al. 2018; Wang, He, et al. 2018; Zou, et al. 2018; Li, Ye, et al. 2019; Wang, Du, et al. 2020; Wang, Wang, et al. 2020; Zou, et al. 2020), however, the low-resolution of these markers hinders the comprehensive understanding of prehistoric human activities on the TP and impedes the dissection of ancestral components of Tibetans. Collectively, previous studies pave the way towards a better understanding of Middle Pleistocene arrival, Paleolithic colonization, and Neolithic permanent settlement on the TP. However, most of the previously archeological investigations have focused primarily on NETP (< 4000 masl), and the lack of discussion of ancient samples from the TP and the uncomprehensive analysis of ancient individuals from East Asia hindered our ability to connect geographically and temporally dispersed ancient East Asians and modern Tibetans. Thus, we comprehensively meta-analyzed the genetic variations of modern and ancient Highlanders from TP and surrounding lowland eastern Eurasians and explored the phylogenetic relationship between East Highlander and reference worldwide populations. By analyzing genome-wide data of Neolithic to Historic individuals from East Asia and modern Tibetans, we shed light on the genetic transition, turnover or continuity, ancestral composition and demographic history of Tibetan Highlanders.

## Results

### Genome-wide data of both modern and ancient Tibetans showed their closed genetic affinity with northern East Asians

To explore the genomic history of modern Tibetans and elucidate the peopling of Qinghai-Tibet Plateau, we used the genome-wide data from 98 modern Tibetans (**Figure 1A**) collecting from eleven geographically different regions and cultural backgrounds from Tibet Autonomous Region (five), Qinghai (two), Gansu (one), Sichuan (two) and Yunnan (One), which were used to reconstruct the genetic background of modern East Asians in our recent ancient genome research of the deep genomic history of East Asia(Wang, Yeh, et al. 2020). Besides, we merged our data with other modern and ancient East Asians(Patterson, et al. 2012; Lipson, Cheronet, et al. 2018; Jeong, et al. 2019; Liu, et al. 2020), in which modern samples from Altai-speaking (also referred as Mongolic, Tungusic, and Turkic language families by other scholars), Sino-Tibetan (Sinitic and Tibeto-Burman), Hmong-Mien, Austronesian, Austroasiatic, and Tai-Kadai language families, and ancient populations included eight individuals from Nepal(Jeong, et al. 2016) (Chokhopani, Samdzong, and Mebrak cultures), eighty-four samples from Yellow River(Ning, et al. 2020; Wang, Yeh, et al. 2020; Yang, et al. 2020), Amur River and West Liao River in the coastal and inland northern East Asia (including Houli, Yangshao, Longshan, Qijia, Hongshan, Yumin, and other cultures), fifty-eight individuals(Ning, et al. 2020; Wang, Yeh, et al. 2020; Yang, et al. 2020) belonged to Tanshishan and other cultures in the coastal southeast China, islands of Taiwan strait and Taiwan. We also included the Neolithic to Bronze Age or Iron Age populations from Southwest Asia(Lipson, Cheronet, et al. 2018; McColl, et al. 2018) and Siberia(Allentoft, et al. 2015; Mathieson, et al. 2015; Damgaard, et al. 2018; de Barros Damgaard, et al. 2018; Sikora, et al. 2019) in some of the following comprehensive population genetic analyses. All Tibetans and Neolithic to historic East Asians were grouped in the East Asian genetic cline along the second component in the Eurasian PCA (Data not shown here). Focused on the genetic variations of East Asian, we constructed East Asian PCA based on genetic variations from 106 modern populations from East Asia and the island and mainland Southeast Asia (**Figure 1B**). We found that modern East Asians grouped into four genetic clines or clusters (Mongolic/Tungusic genetic clines consisting of populations from northeast Asia; South China/Southeast Asian genetic cluster comprising of Austronesian, Austroasiatic, Tai-Kadai, and Hmong-Mien speakers; Sinitic-related north-to-south genetic cline and Tibeto-Burman cluster), which were consistent with the linguistic divisions and geographical regions. Tibetan populations were grouped together and showed a relatively close relationship with some of Mongolic and Tungusic speakers in North China and northern Han Chinese and other lowland Tibeto-Burman speakers. Focused on the population substructures of Tibetan populations, we further observed three different sub-clusters which also were consistent with the geographical position of sampling places. Here, we referred to as to high-altitude adaptive Tibet Tibetan or Ü-Tsang Tibetan cluster (Lhasa, Nagqu, Shannan and Shigatse), Gan-Qing or Ando Tibetan genetic cluster in northeastern TP (Xunhua, Gangcha, and Gannan) and lowland southeast genetic cluster or Kham Tibetans (Chamdo, Xinlong, Yajiang, and Yunnan).

**Figure 1.**
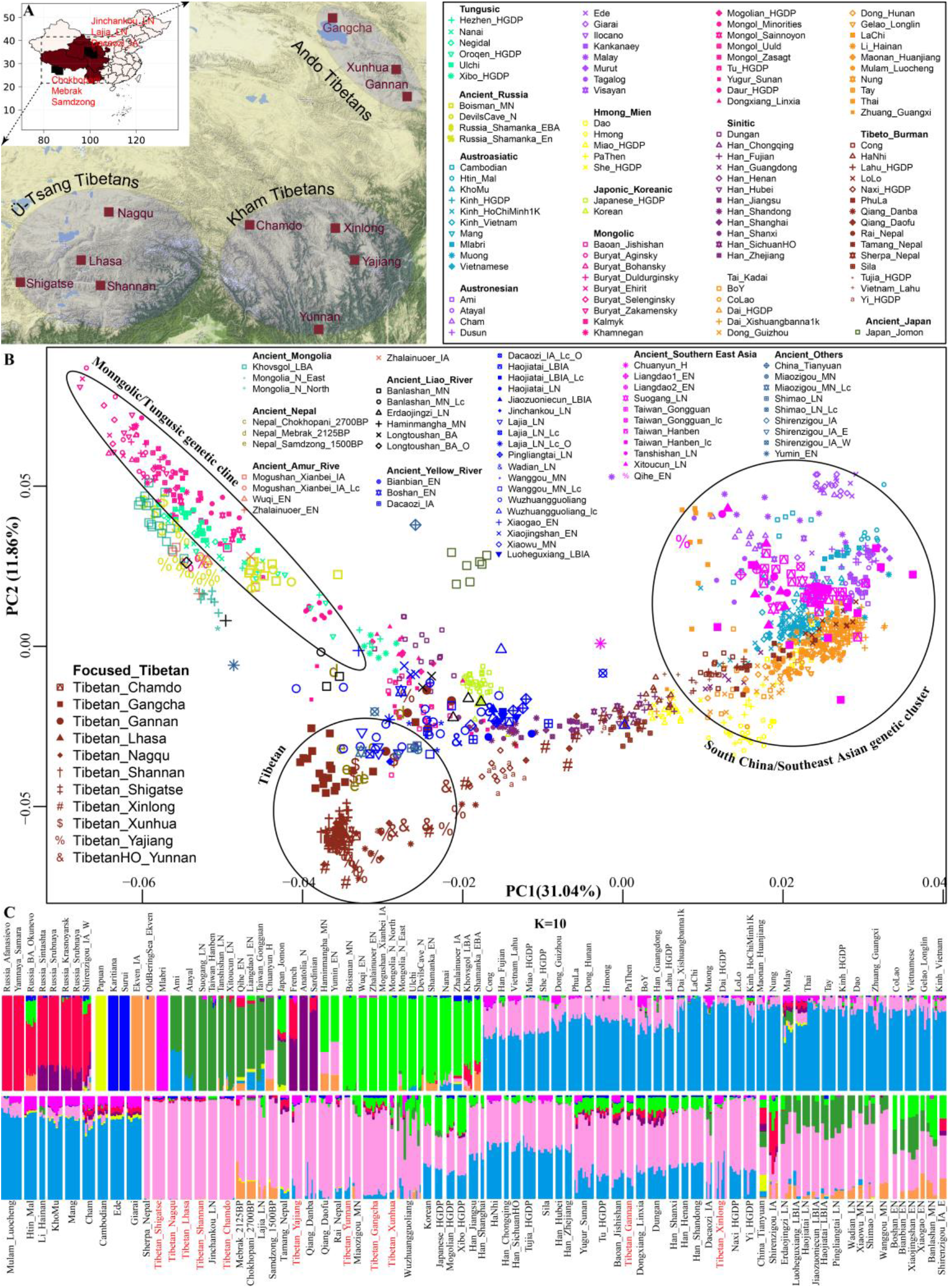
Geographical position of Tibetans and genetic patterns of East Asians. (**A**) Sampling place of eleven modern Tibetans mainly discussed in the present study from Tibet Tibetan Autonomous Region, Qinghai, Gansu, Sichuan and Yunnan provinces. (**B**). Principal component analysis (PCA) showed the genetic similarities and differences between modern and ancient East Asians from geographically/linguistically/ culturally different populations. Spatial-temporally diverse ancient populations were projected onto the two-dimensional genetic background of modern East Asian. (**C**). Admixture ancestry estimation based on model-based ADMIXTURE. Here, the optimal predefined ten ancestral populations were used.

We subsequently explored the patterns of genomic affinity between ancient populations and modern East Asians and projected all included ancient individuals (243 eastern Eurasian ancients) onto the genetic background of the aforementioned patterns of modern population genetic relationships. It should be pointed that this was the first comprehensive meta-analysis of modern and ancient genomes of East Asians. Here, we found four ancient population genetic clusters, Neolithic to historic southern East Asians (including Hanben and Gongguan from Taiwan, and mainland late Neolithic Tanshishan and Xitoucun people) grouped together and clustered with modern Tai-Kadai, Austronesian, Austroasiatic speakers. Neolithic to Bronze Age/Iron Age northern East Asians (both inland and coastal Houli, Yangshao, Longshan and Qijia people) grouped together and projected close to the juncture position of three main East Asian genetic lines and the northmost end of Han Chinese genetic cline, and we observed a close genetic relationship between early Neolithic Houli individuals from Shandong province associated the main subsistence strategy of the Hunter-Gathering and the Yangshao and Longshan middle and late Neolithic farmers in the geographically close Henan province, which indicated the genetic continuity of Neolithic transition from foragers to millet farmers in the early Neolithic northern China. We also identified the subtle genetic differences of these Neolithic to Iron Age individuals from northern China. These Shandong Houli individuals were localized closely with modern Mongolic-speaking Baoan, Tu, Yugur and Dongxiang, while the early Neolithic Xiaogao individuals were posited close with modern Tungusic-speaking Hezhen and Xibo. All Shandong Neolithic ancient populations were localized distant from the modern Shandong Han Chinese and shifted to modern northern Chinese minorities, which indicated modern northern Han received additional gene flow from southern East Asian ancestral lineage or ancient Houli individuals harbored more Siberian-associated ancestral lineage. Henan late Neolithic Longshan individuals (Pingliangtai, Haojiatai, and Wadian) and Bronze/Iron Age individuals (Haojiatai, Jiaozuoniecun, and Luoheguxiang) were grouped and shifted to Han Chinese genetic cline and partially overlapped with Han Chinese from Shanxi and Shandong provinces. This observed genetic similarities from late Neolithic to present-day northern East Asians from Central Plain (Henan, Shanxi, and Shandong provinces) indicated the genetic stability in the core region of Chinese civilization since the late Neolithic period. Middle Neolithic Henan Yangshao individuals (Xiaowu and Wanggou) grouped with some of Wuzhuangguoliang individuals and shifted to more northern modern minorities, and more inland middle and late Neolithic northern East Asians from Shaanxi Shimao, Inner Mongolia Miaozigou and upper Yellow River (Lajia and Jinchankou) clustered together and shifted to modern Tibetans and ancient Nepal high-altitude adaptive ancestral lineage, which was partially overlapped with modern geographically close Tibetans (Gangcha Tibetan and Xunha Tibetan from Qinghai and Gannan Tibetan from Gansu) and also showed the close genetic affinity with Nepal ancients (Mebrak, Samdzong, and Chokhopani).

For ancient populations from West Liao River, three different patterns of genetic affinity can be identified in the projected results: northern cluster (Haminmangha_MN and Longtoushan_BA_O) showed a genetic affinity with Shamanka and Mongolia Neolithic people, middle Hongshan cluster localized between Mongolia minorities and modern Gangcha Tibetans, and southern cluster (Upper Xiajiadian Longtoushan_BA and Erdaojingzi_LN), which was suggested both northern Mongolia Plateau Neolithic ancients associated with steppe pastoralists and Yellow River millet farmers have participated the formation of late Neolithic and subsequent populations in the West Liao River basin. These population movements, interaction and admixture recently have been fully elucidated via Ning et al(Ning, et al. 2020). Here, we observed the late Neolithic populations in the southern cluster was localized between Coastal early Neolithic northern East Asians and inland Neolithic Yangshao and Longshan individuals, which indicated millet farmers from middle and lower Yellow Rivers (Henan and Shandong) have played an important role in the formation of Hongshan people or their descendants via both inland and coastal northward human population migrations. For ancient populations from Mongolia Plateau, Russia Far East, Baikal-Region, and Amur River basin, all included forty-six individuals (Neolithic to Bronze Age Shamanka, Mongolia, DevilsCave, Bosman and others) were clustered closed to modern Tungusic language speakers (Nanai and Ulchi) and some Mongolic speakers. Jomon individuals were grouped together and localized far away from modern Japanese populations in the intermediate position between Russia coastal Neolithic people and modern Taiwan Hanben and coastal Neolithic southern East Asians.

Patterns of genetic relationship revealed from the top two components (extracting 43% variation: PC1: 31.04% and PC2: 11.86%) revealed a genomic affinity between modern Tibetans, ancient Nepal populations, and modern/ancient East Asians and Siberians. To further explore the genetic structure and corresponding population relationships, we estimated the ancestry composition and cluster patterns according to the model-based maximizing likelihood clustering algorithm (**Figure 1C** and **Figure S1**). We observed two northern and two southern East Asian dominant ancestries. Coastal northern East Asian ancestry (light green ancestry) maximized in Neolithic Siberians (Boisman_MN, Wuqi_EN, Zhalainuoer_EN, Mongolia_N_North, Mongolia_N_East, DevilsCave_N, and Shamanka_EN) and modern Tungusic speakers (Ulchi and Nanai). Light green ancestry also existed in the Bronze Age to present-day populations from northeast China and Russia, as well as Coastal early Neolithic northern East Asian from Shandong province with a high proportion. The other northern ancestry enriched in modern highland Tibetans and Qijia culture-related Lajia and Jinchankou late Neolithic populations, which also maximized in Nepal Bronze Age to historic individuals and ancient northern East Asians, as well as the lowland modern Sino-Tibetan speakers, inland Hmong-Mien and Tai-Kadai language speakers. We called this Tibetan-associated ancestry as inland northern East Asians, which was the direct indicator of close genetic affinity between Tibetan and modern and ancient northern East Asians. Deep green ancestry enriched in the Coastal early Neolithic southern East Asians, Iron Age Hanben and modern Austronesian Ami and Atayal, here, we referred it as to coastal southern East Asian ancestry. Blue ancestry maximized in LaChi as the counterpart of the coastal ancestry widely distributed in Hmong-Mien and Tai-Kadai-speaking populations, this blue inland southern East Asian ancestry existed in lowland Tibetans with a relatively high proportion, including Yunnan Tibetan, Yajiang and Xinlong Tibetans and Gannan Tibetan. Besides, we found Tibetans collecting from the northeast TP harbored more coastal northern East Asian ancestry. Some Austroasiatic-associated ancestry, maximized in Mlabri, and Steppe pastoralist-like red ancestry, enriched in Bronze Age Afanasievo and Yamnaya, were also identified in Sichuan and Yunnan Kham Tibetans, and Qinghai and Gansu Ando Tibetans respectively. Ancient Nepal populations had common ancestry associated with Iron Age Ekven people from northeast Siberia.

### Population differentiation between highland and lowland East Asians and substructure among Tibetans

To further explore the genetic differentiation between eleven modern Tibetans and modern or ancient reference populations, we first calculated the pairwise Fst genetic distance among 82 modern populations (**Table S1**, modern dataset) and 32 modern and ancient populations (**Table S2**, ancient dataset). We found a strong genetic affinity among geographically close populations. As shown in **Figures S2~3**, the high-altitude Ü-Tsang Tibetans from the south (Shigatse and Shannan), central (Lhasa), north or northeast (Nagqu and Chamdo) of Tibet Autonomous Region had the smallest Fst genetic distance with their geographical neighbors, followed by lowland Ando Tibetans from the northeast TP (Qinghai and Gansu) and the Kham Tibetan from the southeast region of TP (Sichuan and Yunnan) and other Tibeto-Burman speaking populations (Daofu Qiang, Tu and Yi), these observed patterns of genetic relationship was significantly different from the lowland populations, such as Hmong-Mien-speaking She sharing most ancestry with lowland East Asians (**Figure S3**). For Ando Tibetans from Qinghai and Gansu provinces, Gangcha Tibetan harbored a close genetic affinity with northern or northeastern Tibet Tibetans with the smallest Fst genetic distances (Chamdo and Nagqu), followed by Qiang and Tu or other geographically close Tibetans (**Figure S4**). Different patterns were observed in Gangcha Tibetan and Xunhua Tibetan, which showed the closest relationship with each other, and then followed by Tu and Yugur. We also found a relatively small genetic distance between Tibetans (Gannan and Xunhua) and Turkic-speaking Kazakh population, which showed a western Eurasian affinity of Tibetan from the northeast region of TP relative to the Tibetans from the central region of TP. **Figure S5** presented the patterns of genetic differentiation between lowland Kham Tibetans and their reference populations, and we found that Yajiang and Xinlong Tibetans from Sichuan harbored the close genetic affinity with the Tibeto-Burman speakers (Tibetan, Qiang, Yugur, and Tu) from Sichuan and Ganqing Region. Yunnan Tibetan had the smallest genetic distance with Gangcha and Chamdo Tibetans, followed by Qiang, Yi, and Tu. Among Tibetans and Neolithic to Iron Age East Asians (**Figure S6**), the top genetically closest populations harboring the smallest Fst values with each modern Tibetans is proved by the other modern Tibetans, and we also found Hanben population from Taiwan showed the closest relationship with modern Tibetans relative to other ancient East Asians.

Phylogenetic relationships were further estimated under the genetic variations of Eurasian modern populations and eastern Eurasian ancients via the TreeMix-based analyses. As shown in **Figure 2A**, a phylogenetic tree with no migration events showed that modern populations from similar language families tended to cluster into one group. Altai-speaking (Turkic and Mongolic) populations were clustered with Uralic speakers. Southern Austronesian East Asian first clustered with Tai-Kadai speakers and then clustered with Hmong-Mien and Austroasiatic speakers. Tibetans clustered first with each other, especially for high-altitude adaptive Ü-Tsang Tibetans and then clustered with the lowland East Asians, this observed geographical isolation showed the genetic differentiation between modern highland Tibetans and lowland East Asians could be identified although shared the common originated lineage. We further analyzed the population splits and gene flow events between modern Tibetan and 26 ancients from eastern Eurasia (except for Anatolia_N from Near East) with three predefined admixture events. Modern Tibetans except for Gannan and Xinlong Tibetans were first clustered with highland Nepal ancients and then clustered with lowland Neolithic northern East Asians and Neolithic to Bronze Age southern Siberians, which also showed a genetic division between highland modern Tibetans and lowland ancient northern East Asians (**Figure 2B**). The cluster patterns also showed a distant relationship between northern and southern East Asians, as well as highland modern and ancient Tibetans and lowland southern East Asians, which further provided evidence for some special connections or closer genetic relationships between Tibetans and northern East Asians.

**Figure 2.**
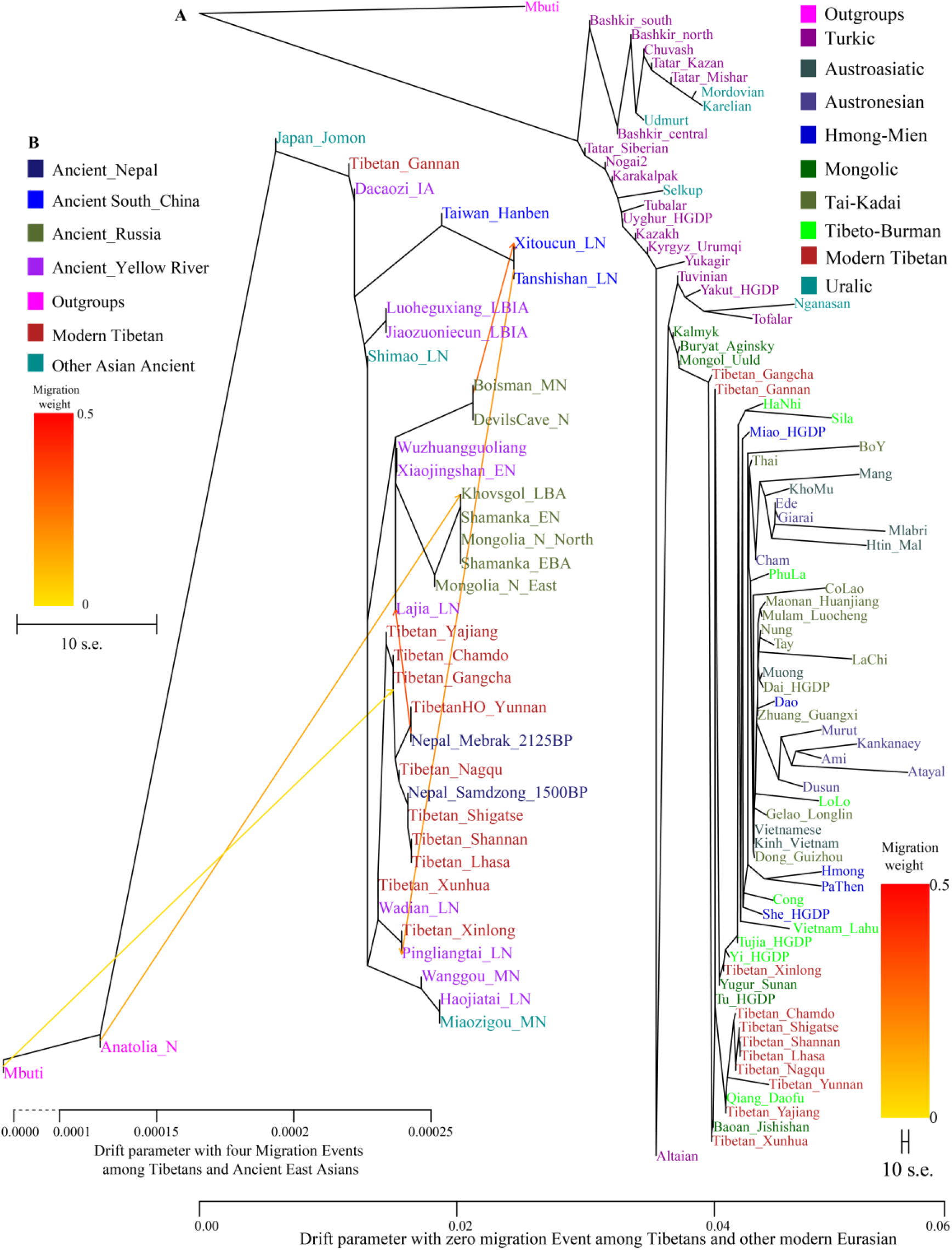
Maximum likelihood phylogeny reconstruction based on the genetic variation from both modern Tibetan and Eurasian modern reference populations. (A), modern Tibetan and Neolithic to historic East Asian (B). Mbuti was used as the root. Focused on the phylogenetic relationship among all modern populations, we used the patterns of genetic relationship with zero migration events. And evaluating the evolutionary history among modern Tibetan and ancient Chinese, we included three migration events. To better present our result, the drift branch length of Mlabri was shortened as the third of the truth drift branch length due to strong genetic drift occurred in Mlabri.

Genetic affinity was further evaluated via the outgroup-*f_3_*-statistics of the form *f_3_(modern Tibetans, Eurasian modern/ancient populations, Mbuti)*. Among 184 modern populations (**Table S3**), the top sharing for each Tibet Tibetans is provided by another geographically close Tibetans. Shannan Tibetan shared most alleles with Lhasa/Shigatse/Nagqu Tibetans, and the similar patterns of population affinity were identified in the other southern Shigatse Tibetan and central Lhasa Tibetan. However, Nagqu Tibetan shared most alleles with the northeastern Chamdo Kham Tibetan (followed by Tibetan-Burman-speaking Qiang from Sichuan province and other Tibetans or Sherpa), these patterns were consistent with the population features in Chamdo Tibetan. Following by genomic affinity within Tibetans, we also found that these five Tibet Tibetans shared the strongest genetic affinity with the lowland Han Chinese, which is consistent with the common origin from the middle and lower Yellow River basin of Sino-Tibetan language speakers. For lowland Kham Tibetans in Sichuan and Yunnan provinces, Xinlong Tibetan shared the most genetic drift with Sinitic-speaking populations (Han from Shanghai, Chongqing, Hubei, Jiangsu, and others) and other lowlands Tibeto-Burman-speaking Qiang and Tujia. Being different from Xinlong Tibetan, geographically close Yajiang and Yunnan Tibetans shared the most drifts with Qiang and geographically close Tibetans (Chamdo and Xinlong), followed by Sinitic speakers and other Tibetans. These lowland Han or southern East Asian affinity suggested that lowland Kham Tibetans from southwestern China harbored shifts in ancestry with southern East Asians in prehistoric and historic times via population migration and admixture. Ando Tibetan of Gangcha Tibetan not only showed the genetic affinity with Sinitic and Tibeto-Burman speakers but also showed the signals of Turkic-speaking population affinity. Allele sharing results from Gannan and Xunhua Tibetans showed the Han Chinese groups shared the most ancestry components with them.

Levels of allele sharing between modern Tibetans and 106 Paleolithic to historic Eurasian ancient populations (including 33 populations from Russia, 41 from China, 29 from Mongolia, three from Nepal) inferred from *outgroup-f_3_-statistics* showed that modern Tibetans had the clear connections with ancient Neolithic to Iron Age northern East Asians, which was consistent the patterns observed in the PCA, Fst, ADMIXTURE and modern population-based affinity estimations (**Table S3**). Middle-altitude Chamdo Tibetan shared the most genetic drift with Neolithic Wuzhuangguoliang, upper Yellow River late Neolithic farmers (Jinchankou and Lajia, which are the typically represented source populations for Qijia culture), followed by Iron Age Dacaozi people, Shimao people from Shaanxi, middle Neolithic Banlashan associated with Hongshan culture in North China and other northern East Asians from lower and middle Yellow River basin. Neolithic people from Russia and Mongolia and Bronze Age to historic Nepal ancients showed a relatively distant genetic relationship with modern Chamdo Tibetan (**Figure S7**). Different from the patterns of Chamdo Tibetan, southern and central Ü-Tsang Tibetans showed the increased ancestry associated with Nepal ancient people, and northern Nagqu Tibetan showed the intermediate trend of population affinity with 2700-year-old Chokhopani. As showed in **Figures S8~9**, lowland Tibetans from southwestern China and northeastern China showed a similar population affinity to northern East Asian ancients. The genomic affinity between modern Tibetan and 43 East Asians was visualized in **Figure 3**. Tibetans except for Xinlong Tibetan shared the most genetic drifts with each other and clustered together and then grouped with three Nepal ancients and formed the Highlander cluster, early Neolithic to Iron Age northern East Asian clustered first and then grouped with Highlander cluster. Amur and West Liao River ancient cluster also showed a closer relationship with highland Tibetans, and Xinjiang Shirenzigou people and southern East Asians kept a relatively distinct relationship with northern lowland East Asians and highland Tibetan clusters.

**Figure 3.**
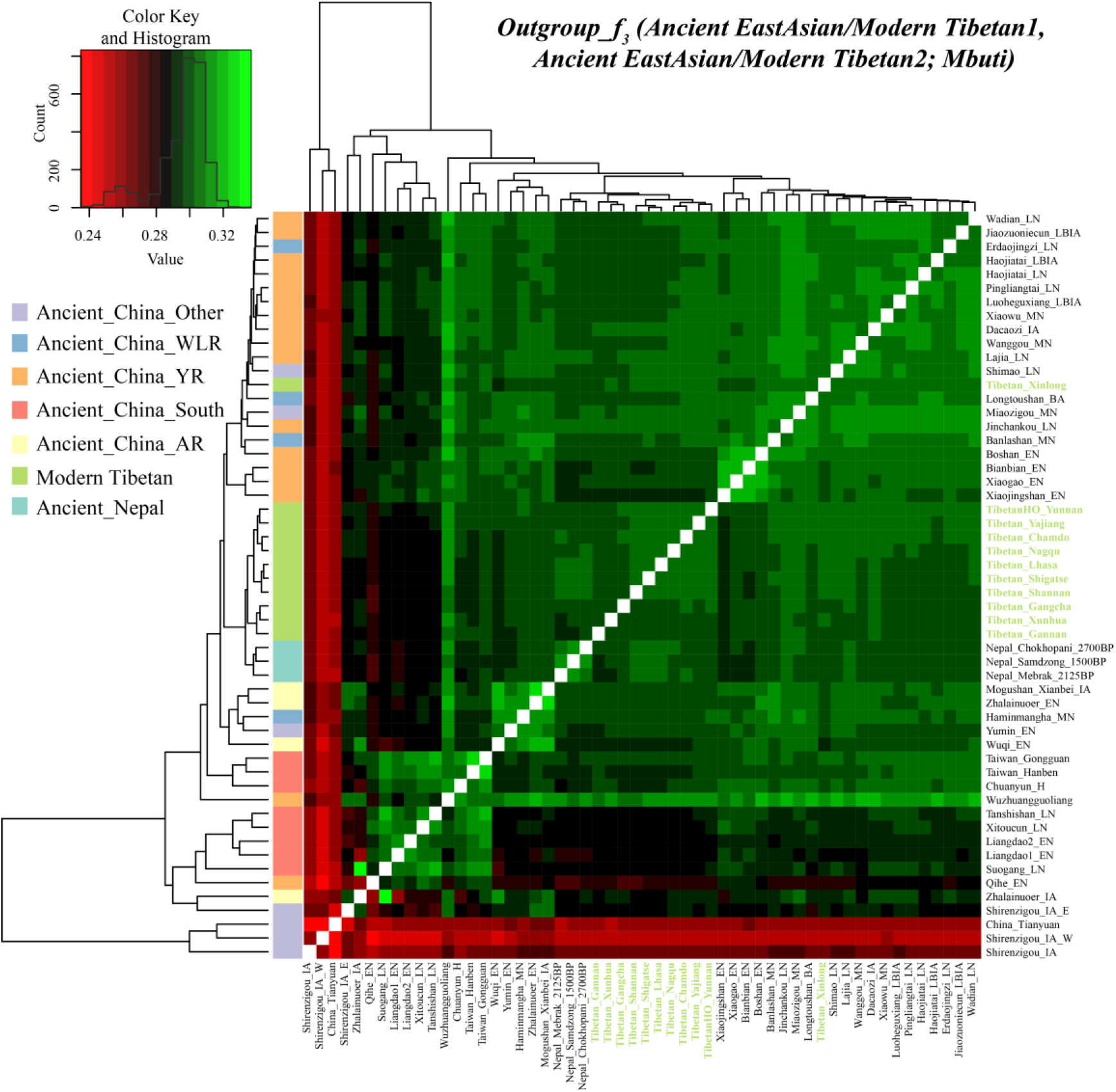
Genomic affinity between our eleven targeted Tibetan populations and other 43 spatial-temporally different East Asian populations. The color of the bar code in the left cluster showed the geographical origin of ancient samples.

### Admixture signatures of modern Tibetans and ancient populations from Qinghai-Tibet Plateau

To further explore whether any populations harbored the obvious evidence or signals for recent genetic admixture and determine their corresponding ancestral source populations, we carried out admixture-*f_3_-statistics* of the form *f_3_(Modern/ancient source population 1 Modern/ancient source population2; Targeted Tibetan populations)* to evaluated the extent to which Tibetans from different geographical divisions possess the shared derived alleles from their modern ancestral proximity or ancient source populations. We also re-evaluated the admixture signatures of eight individuals from Nepal (Chokhopani, Mebrak, and Samdzong with geographical affinity with Ü-Tsang Tibetan) and eleven individuals from Qinghai province (Late Neolithic Lajia people and Jinchankou people, and Iron Age Dacaozi population with geographical affinity with Ando Tibetans) using this three-population comparison method and our comprehensive modern and ancient reference source database. We found different patterns of admixture signals and source populations in highland and lowland modern and ancient Tibetans (**Tables S4~18**), besides, we could also identify small but significant differences among Tibetans from one geographical region or similar culture backgrounds (Ü-Tsang Tibetans from Tibet, Ando Tibetan from Gansu and Qinghai, Kham Tibetans from Sichuan and Yunnan). With the statistic significant level of Z-scores with three, no admixture signals were observed in southern Tibetans (Shannan and Shigatse) over forty thousand tested pairs, only four in central Lhasa Tibetan (one source from 1500-year-old Samdzong and other from Kham Tibetan/Qiang, or the combination of southern Tibet Tibetan with Neolithic northern East Asians or Baikal lake ancients, **Tables S4~6**). It is interesting to found that 188 tested population pairs showed statistically significant *f_3_*-statistic values with one source from Tibeto-Burman speakers and the other from Western Eurasians (Alan, Andronovo, Sintashta, Poltavka, Yamnaya and modern people) in *f_3_(Source1, Source2; Nagqu Tibetan)*. Tibetans from southern and central Tibet combined with lowland modern East Asians, not ancient northern and southern East Asians, could also produce significant admixture signals for Nagqu Tibetan (**Table S7**). Chamdo Tibetan localized the junction regions between Ü-Tsang Tibetan and Kham Tibetan possessed potential active cultural and human population movements and admixtures, but only one admixture signals observed here, *f_3_(Lhasa Tibetan, Yajiang Tibetan; Chamdo Tibetan) = −3.49*SE* (**Table S8**). Three Tibetans from the Ganqing region possessed admixture signatures from over several thousand population pairs with one from modern or ancient East Asians and the other from Western Eurasians (**Tables S9~11**). Results from *f_3_(Yumin_EN, Austronesian/Tai-Kadai; Ganqing Tibetans)* showed the inland Neolithic northern East Asian Yumin_EN as northern ancestral source combined with Austronesian/Tai-Kadai speakers as the southern ancestral source could produce significant*f_3_* values, these admixture signals could be also identified in *f_3_ (Neolithic northern East Asians, Neolithic Russian/modern Turkic/Mongolic/Indo-European speakers; Ganqing Tibetans)*. Tibetans from Sichuan only showed significant signals as the result of the admixture between northern and southern East Asians or the highland Tibeto-Burman speakers and lowland East Asians, i.e. *f_3_ (highland Tibeto-Burman speakers, lowland Tibeto-Burman speakers; Sichuan Tibetan)<-3*SE* (**Tables S12~13**). Similar to the results for the southern Tibet Tibetan, no obvious admixture signals were observed in Yunnan Tibetan, which may be caused by the genetic isolation or obvious genetic drift that occurred recently (**Table S14**). Tests focused on the ancient populations from TP showed seven admixture signals from Qinghai Iron Age Dacaozi people (**Table S15~18**), which are the admixture results of ancient northern East Asians and modern southern East Asians, or Chamdo Tibetan-related source and Taiwan Iron Age Hanben-like population.

### Intra population differentiation amongst high-altitude and low-altitude residing Tibetans inferred from *f_4_*-statistics

To gain insights into the population substructures among modern Tibetans, we first conducted symmetry-*f_4_* statistics of the form *f_4_(modern Tibetan1, modern Tibetan2; modern Tibetan3, Mbuti)*, in which we expected to observe the no significant positive or negative *f_4_* values if no significant differences existed between modern Tibetan1 and modern Tibetan2 relative to reference Tibetan3. As shown in **Table S19 and Figure S10**, in the tests of symmetry-*f_4_* statistics of the form *f_4_(Tibetan1, Chamdo Tibetan; Tibetan2, Mbuti)*, we observed that Chamdo Tibetan formed a clade with Tibetans from Nagqu and Yunnan compared with other Tibetan reference populations. All included Tibetans shared more alleles with Chamdo Tibetan compared with Gangcha Tibetan, Gannan Tibetan, and Xunhua Tibetans from Ganqing Ando Tibetan Region with significant negative *f_4_* values. Compared to related low-altitude Tibetans (Yajiang and Xinlong), Chamdo Tibetan had more high-altitude Tibetan-related ancestry (Lhasa, Nagqu, Shigatse, and Shannan), while Gannan Tibetan shared more alleles with Xinlong Tibetan compared with Chamdo Tibetan. Compared with high-altitude Tibetan, Chamdo Tibetan shared more alleles with other Tibetans from relatively low-altitude sampling places. Results from the symmetry-*f_4_* statistics of the form *f_4_(Shigatse/Shannan/Lhasa Tibetans, Shigatse/Shannan/Lhasa Tibetans; Tibetan2, Mbuti)* with non-significant Z-scores showed clear genetic homogeneity among Tibet central and southern-Ü-Tsang Tibetans (**Figures S11~12**). Negative values in *f_4_(Ganqing Ando Tibetans, Shigatse/Shannan/Lhasa Tibetan; Tibetans, Mbuti)* showed all included Tibetans shared more alleles with southern Tibet Tibetans relative to Ganqing Ando Tibetans. However, northern Tibet Tibetan formed a clade with Chamdo and Yunnan Tibetans and received more high-altitude Tibetan-related derived alleles compared with Ganqing and Sichuan Tibetans. For lowland Tibetans, northwestern Chinese Gangcha and Xunhua Tibetans formed one clade, i.e., all absolute Z-scores of *f_4_(Gangcha, Xunhua Tibetan; Tibetan2, Mbuti)* are less than three (**Figure S13**). Compared with Gannan Tibetans, Qinghai Tibetans harbored more ancestry shared with Tibet Tibetan. We found no Tibetan populations shared more alleles with Gannan Tibetans relative to other Tibetans, as all *f_4_(Tibetan1, Gannan Tibetan; Tibetan2, Mbuti)* values were larger than zero. Southwestern Chinese Yunnan Tibetan formed one clade with Chamdo/Xinlong and Yajiang Tibetan, all of them belonged to Kham Tibetan (**Figures S14~15**). Lowland Sichuan and Yunnan Tibetans harbored increased Tibetan-related derived alleles compared with Ganqing Tibetans and more ancestry related to highland Tibetans compared with other highland Tibet Tibetans.

We additionally explored the observed genetic affinity and population substructure among highland and lowland Tibetans using ancient Eurasian populations (mainly collected from China, Mongolia, and eastern Siberia and some steppe pastoralists from western Eurasia) via *f_4_(Modern Tibetan1, Modern Tibetan2; Ancient Eurasians, Mbuti)*. Results with no significant positive or negative Z-scores in *f_4_(Ü-Tsang Tibetans1, Ü-Tsang Tibetans2; Ancient Eurasians, Mbuti)* confirmed the patterns of genomic affinity within high-altitude adaptive Ü-Tsang Tibetans (Shannan, Shigatse, Lhasa, and Nagqu), we could also identify obvious more affinity with Nepal ancients in Ü-Tsang Tibetans relative to Ando and Kham Tibetans (**Figures S16~19**). Compared with Shannan Tibetan, Nagqu Tibetan harbored increased ancestry associated with lowland ancient populations of Neolithic/Historic southern East Asians in the southeastern coastal region of South China (Tanshishan_LN and Chuanyun_H) to as north far as Baikal Region (Russia_UstIda_LN). Compared to Tibetans from Qinghai Ando Tibetans, Nagqu Tibetan owned both increased Nepal ancient-related ancestry and increased Late Neolithic Lajia related ancestry relative to Xunhua Tibetan, and it also harbored additionally increased ancestry related to coastal Late Neolithic southern East Asians of Tanshishan_LN, middle Yellow River Middle Neolithic to Iron Age ancient populations (Wanggou_MN, Haojiatai_LBIA, and Jiaozuoniecun_LBIA), Upper Xiajiadian culture of Longtoushan_BA, inland Neolithic northern East Asians (Yumin_EN and Shimao_LN) and other upper Yellow River Late Neolithic Jinchankou and Iron Age Dacaozi. Here, we found a closer affinity between upper Yellow River ancient populations with Nagqu Tibetan, not with geographically close Gangcha Tibetan, which suggested that ancient populations from Lajia, Jinchankou and Dacaozi may be the direct ancestors of modern Nagqu Tibetan. Significant negative *f_4_* values were observed in Chinese Ando Tibetans via *f_4_(modern Tibetan1, Ganqing Ando Tibetans; Bronze Age stepped pastoralists, Mbuti)*, which suggested that Ando Tibetan harbored increased ancestry with Sintashta-like, Yamnaya-related and other ancestry related to middle and late Bronze Age Steppe pastoralists (Afanasievo, Srubnaya, Andronovo and Xinjiang Iron Age Shirenzigou populations). Although strong genetic affinity among Ando Tibetans was confirmed with the similar patterns of *f_4_*-based sharing alleles and non-significant statistical results in symmetry-*f_4_* statistics. Negative *f_4_* values with Z-scores larger than three in *f_4_(Gangcha Tibetan, Gannan Tibetan; Ami/Atayal/Hanben/ Gongguan/Tanshishan_LN/Qihe_EN, Mbuti)* showed that Gannan Tibetan harbored increased southern East Asian ancestry represented by modern Austronesian or Proto-Austronesian-related early Neolithic to present-day southeastern coastal/island populations (**Figures S20~22**). The same southern East Asian affinity of Gannan Tibetan was also identified compared with Tibet Ü-Tsang Tibetans.

Results of the four-population comparison analysis focused on Kham Tibetans were presented in **Figure 4** and **Figures S23~25**. The results of *f_4_(Nagqu Tibetan, Chamdo Tibetan; eastern Eurasian ancients, Mbuti)* suggested that Nagqu Tibetan formed one clade with Chamdo Tibetan except for the reference population of Banlashan-MN with significant negative *f_4_* value, which meant that Chamdo Tibetan harbored increased middle Neolithic northern East Asian Banlashan-related ancestry, and Banlashan people was evidenced to be associated with archeologically attested Hongshan culture. Compared with Lhasa Tibetan, *f_4_(Lhasa Tibetan, Chamdo Tibetan; eastern Eurasian ancients of DevilsCave_N/Mongolia_N_East/Banlashan_MN/Wanggou_MN/Xiaowu_MN/Pingliangtai_LN/Lajia_ LN, Mbuti)* showed significant negative statistical results, which suggested increased Russia- or Mongolia-related Neolithic ancestries (DevilsCave_N/Mongolia_N_East), middle Neolithic Hongshan culture-related ancestry (Banlashan_MN), middle Yellow River middle to late Neolithic Yangshao or Longshan farmer-related ancestry (Wanggou_MN/Xiaowu_MN/Pingliangtai_LN) and upper Yellow River late Neolithic Qijia culture-related ancestry (Lajia_LN) in Chamdo Tibetan relative to Lhasa Tibetan. The genetic affinity showed a link between ancient populations from the TP and northern East Asia during the early Neolithic period. Compared with southern Ü-Tsang (Shannan) Tibetans, Chamdo Tibetan harbored increased ancestry related to different ancestral populations from the lowland East Asian, as *f_4_(Shannan Tibetan, Chamdo Tibetan; both southern and northern ancient East Asians and ancient Siberians, Mbuti)* showed significant negative *f_4_*-statistic values. First, coastal late Neolithic southern East Asian Tanshishan, Iron Age Taiwan Hanben islander, and historic Chuanyun people shared more drift with Chamdo Tibetan than with Shannan Tibetan. Second, Coastal early Neolithic northern East Asians from Shandong province (Xiaojingshan_EN, Xiaogao_EN, Bianbian_EN, and Boshan_EN) shared more genetic drift with Chamdo Tibetan. Third, middle Neolithic to late Bronze Age and Iron Age ancient populations from Henan province in the middle and lower Yellow River shared more derived alleles with Chamdo Tibetan, such as middle Neolithic people of Wanggou and Xiaowu, late Neolithic people of Wadian, Pingliangtai and Haojiatai, and Late Bronze Age and Iron Age Jiaozuoniecun and Luoheguxiang. Fourth, Neolithic populations from the middle Yellow River basin (Wuzhuangguoliang and Shimao_LN) shared more alleles with Chamdo Tibetan. Fifth, three populations from late Neolithic (Lajia_LN and Jinchankou_LN) to Iron Age (Dacaozi_IA) in the upper Yellow River shared more alleles with Chamdo Tibetan than with Shannan Tibetan. Sixth, three Neolithic populations from West Liao River, including Haminmangha_MN, Banlashan_MN, and Erdaojingzi_LN, shared more alleles with Chamdo Tibetans. Seventh, ancestral populations related to the Neolithic to present-day people from Mongolia and Russia, including Mongolia_N_East, Russia_UstIda_LN, Mongolia_N_North, Mongolia_Medieval, DevilsCave_N, Boisman_MN, Russia_Shamanka_EBA, Shamanka_Eneolithic and modern Ulchi, shared more alleles with Chamdo Tibetans. Compared with Shigatse Tibetan, similar patterns of sharing derived alleles were observed. Compared with Ando Tibetans, Chamdo Tibetan shared an increased ancestry component associated with both highland and lowland ancient populations. And compared with Tibetans from Sichuan province (Xinlong Yajiang Tibetans), Chamdo Tibetan shared more alleles with 2125-year-old Mebrak and 1500-year-old Samdzong. Similar to the patterns of genetic affinity observed in Chamdo Tibetan, the other three Kham Tibetan also had increased both northern and southern East Asian ancestry.

**Figure 4.**
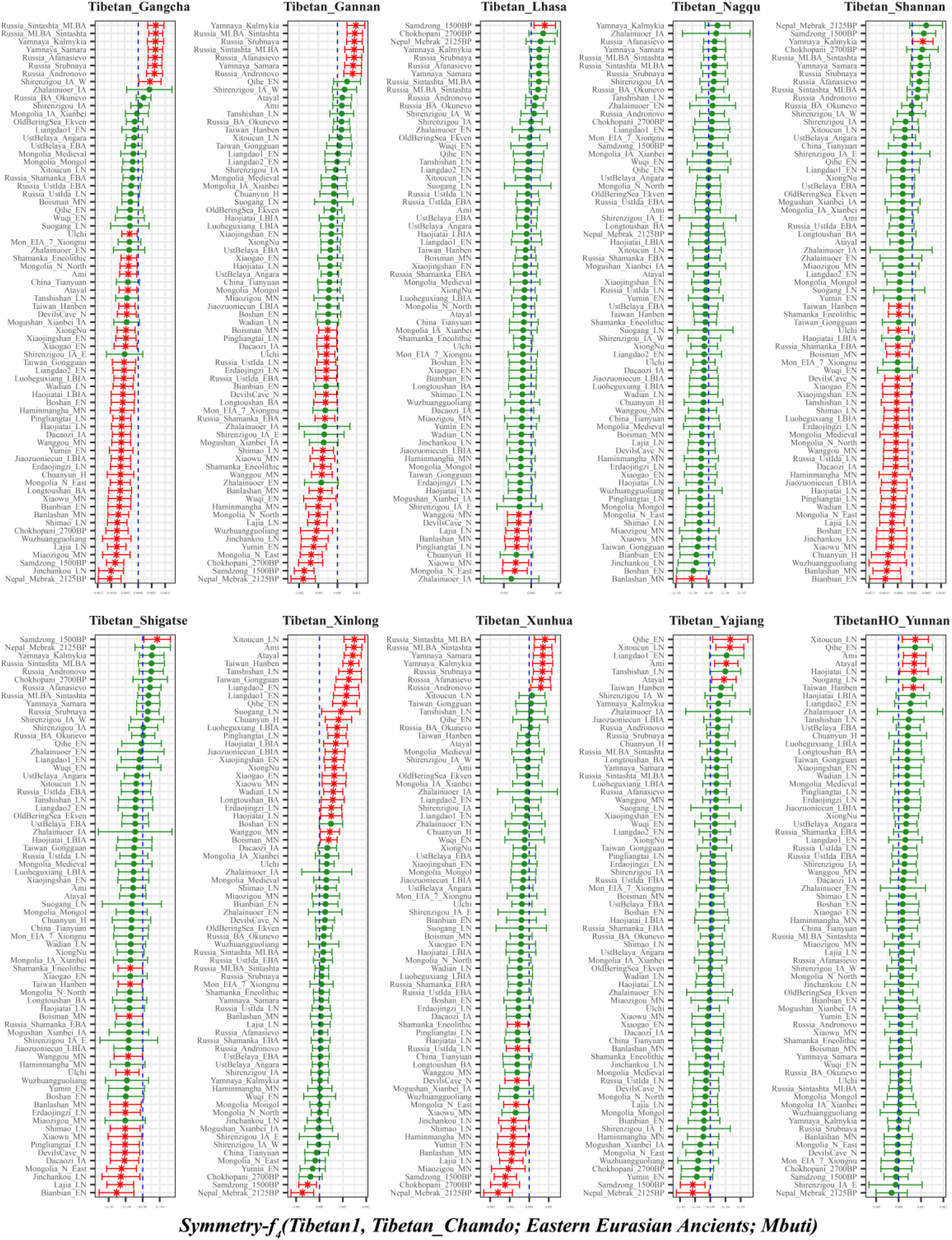
Genomic affinity between modern Tibetans and eastern Eurasian ancient populations inferred from four population symmetry-f_4_ statistics of the form f_4_(Tibetan1, Chamdo Tibetan; eastern Eurasian ancients, Mbuti). Here, overlapping SNP loci included in the Affymetrix Human Origins platform among four analyzed populations were used. We used the genetic variation of Mbuti as the outgroup. Red asterisk point meant the significant value (Absolute value of Z-scores larger than three or equal to three) observed in the symmetry-f_4_ statistics and green circle point denoted the non-significant f_4_-statistic values (Absolute value of Z-scores less than three). All Tibetan2 were listed along the Y-axis and f_4_ values were labeled along the X-axis. All tested population pairs were faceted or grouped via the Tibetan1. Significant negative f_4_ values indicated that the included eastern Eurasian ancient population shared more alleles with Chamdo Tibetan compared with Tibetan1 or Chamdo Tibetan harbored increased eastern Eurasian ancient population-related ancestry compared with Tibetan1, and significant positive f4 value indicated that the included eastern Eurasian ancient population shared more derived alleles with Tibetan1 compared with Chamdo Tibetan or elucidated as Chamdo Tibetan had increased eastern Eurasian ancient population-related ancestry relative to Tibetan1. The value off_4_-statistics equal to zero was marked as the blue dash line. The bar indicated three standard errors.

### Spatiotemporal comparison analysis among modern Tibetan and all Paleolithic to Historic East Asians and genetic admixture and continuity of modern Tibetans

To clearly elucidate the patterns of genetic structures and population dynamics of overall East Asians and provide new insights into the origin of culturally/geographically diverse Tibetans, we carried out both spatial and temporal difference explorations via *f_4_*-statistics. Focused on four early coastal Neolithic northern East Asians from Shandong province, *f_4_(Coastal Neolithic northern East Asian1, Coastal Neolithic northern East Asian2; Modern Tibetans/Ancient East Asians, Mbuti)* revealed the similar genetic relationship between modern Tibetans and these Neolithic northern East Asians (**Figure S26**). Increased coastal Neolithic southern East Asian related ancestry could be identified in Xiaojingshan_EN people, which showed a close connection with coastal populations in East Asia and was wonderfully illustrated in a recent ancient study by Fu et al(Yang, et al. 2020). Results from *f_4_(Bronze/Iron Age Henan populations, Neolithic to Iron Age Henan populations; Eastern Modern Tibetan/Ancient East Asians, Mbuti)* only revealed Luoheguxiang people had increased ancestry associated with modern Austronesian-speaking Ami (**Figures S27~29**) relative to Wanggou_MN. Late Neolithic Haojiatai population had more southern East Asian ancestry related to Xitoucun_LN and Hanben people compared with Wanggou_MN (**Figure S30**), similar southern coastal population (Ami, Atayal, and Hanben-related sources) affinity was observed in Pingliangtai_LN, but not in Wadian_LN and middle Neolithic Wanggou_MN and Xiaowu_EN (**Figures S31~34**). Focused on ancients from Shaanxi and Inner Mongolia, we found modern Tibetans and northern and southern East Asians from the Yellow River and South China shared more alleles with late Neolithic Shimao populations (**Figure S35**). Temporal analysis among upper Yellow River ancients showed all modern Tibetans showed the similar relationship with them, although Iron Age Dacaozi people harbored more southern East Asian ancestry, as revealed by significant positive *f_4_* values in *f_4_(Dacaozi_IA, Lajia_LN; Neolithic Qihe and Xitoucun/Iron Age Gongguan and Hanben/ modern Ami and Atayal, Mbuti)*. These results suggested that population movements from South China have a significant influence on the gene pool of northeastern populations in the TP at least from Iron Age (**Figure S36**). Symmetrical relationships among East Asians with temporally different Nepal ancient populations was evidenced in **Figure S37**.

Next, we also explored the similarities and differences of the shared genetic profiles related to northern Neolithic East Asians via the spatial comparison analysis in modern Tibetans and all available ancient East Asians. We sought up a series of symmetry *f_4_*-statistics, where we compared all eleven modern Tibetans and other ancient East Asians against the geographically different ancient northern East Asians and ancient Tibetans. **Figures S38~41** showed the shared alleles between targeted populations against the lowland early Neolithic northern East Asians and others. Symmetry *f_4_ (Northern East Asians, Chokhopani; Modern Tibetan/Neolithic to Historic East Asians, Mbuti)* was used to determine the lowland and highland East Asian affinity. Compared with four coastal Neolithic Shandong populations, we found that Ü-Tsang Tibetans had the strong highland East Asian affinity. Besides, comparing against coastal and inland ancients revealed that modern Tibetans had a strong inland northern East Asian affinity, especially with late Neolithic Lajia people from the upper Yellow River. This Lajia affinity or inland northern East Asian affinity persisted when we substituted inland Yumin_MN with the coastal Neolithic northern East Asians (**Figure S42**) but disappeared when we substituted the latter Neolithic with the early Neolithic northern East Asians (**Figures S43~48**). We summarized the overall highland and lowland East Asian affinities of Tibetans in **Figure S49**, which showed the Ando and Kham Tibetans had lowland northern East Asian affinity, Ü-Tsang Tibetans with Nepal ancient affinity.

Aforementioned population genomic studies have identified population substructures among modern Tibetans (Ü-Tsang, Ando and Kham Tibetans) and their closest relationship with northern East Asians and affinity to southern East Asians and Siberians, which was confirmed with the negative *f_4_(Reference populations, modern Tibetans; northern/southern East Asians and Siberians, Mbuti)* values. Here, consistent with three subgroups respectively showing the affinity to northern and southern East Asians and Siberians, we further assumed that they descended directly from one of these source populations without additional genetic admixture, and thus we would expect non-significant *f_4_* values in *(Source population, modern Tibetans; Reference populations, Mbuti)*, as negative values denoted additional gene flow into modern Tibetans. We first assumed modern Tibetans were the direct descendants of southern East Asians which is associated with the Yangtze Rice farmer ancestry. As shown in **Figures S50~58**, we observed significant negative *f_4_* values when we used northern East Asians or Siberians as the reference populations, which indicated obvious gene flow events from these reference populations and close genetic relationship. To dissect the additional gene flow events when we assumed that Tibetans’ direct ancestor is coastal Neolithic northern East Asian related ancestral populations, we conducted *f_4_(Shandong ancients, Modern Tibetans; Neolithic to Historic East Asians, Mbuti)* and found only Nepal ancients showed a negative *f_4_* values, which is consistent with the common origin from middle and lower Yellow River basin of Sino-Tibetan speakers (**Figures S59~62**). The patterns were confirmed when we assumed Yangshao and Longshan farmers or their related populations (**Figure S63~71**), Shaanxi ancients (**Figure S72~74**) and other ancient northern East Asians and southern Siberians (**Figures S75~88**) as the direct ancestor of modern Tibetans. As shown in **Figures S75 and S88**, assuming Yumin or Ulchi as their direct ancestor, additional ancestral gene flows from the southern East Asians (Hanben and Tanshishan et.al.) and Yellow River farmers were identified. Assuming the Nepal ancients as direct ancestors, obvious additional gene flow events that occurred from the lowland ancient East Asians were detected in Kham Tibetans (**Figures S89~91**). Additional predefined ancestral populations from Russia and Chinese Xinjiang further confirmed the strong East Asian affinity (**Figures S92~104**).

### Ancestry compositions of modern and ancient Tibetans via *qpWave/qpAdm* and *qpGraph*

Considering the close connections of modern Tibetan and Neolithic northern East Asians and paternity affinity between Tibetan, Onge, and Jomon(Shi, et al. 2008) (the latter two have been evidenced as the early Asian lineage with the close relationship with 7700-year-old Hoabinhian from southeast Asia(McColl, et al. 2018)), we further explored the number of ancestral populations of modern Tibetans, Nepal ancients and Jomon Hunter-gatherer from the Japanese archipelago using the *qpWave* and estimated their corresponding ancestry proportion under one-way, two-day and three-way admixture models via *qpAdm*. The *qpWave* results (p_rank<0.05) showed that there were needed at least two ancestral populations to explain the observed genetic variations in targeted populations. We first employed the two-way model of Onge and six inland/costal early Neolithic northern East Asians and found inland Yumin failed to fit our targeted populations’ genetic variations (all p values <0.05). Xiaogao_EN-Onge two-way model could be well fitted all modern Tibetans except for Gannan Tibetan with the Xiaogao-related ancestry proportion ranging from 0.846 in Shannan Tibetan to 0.906 in Xinlong Tibetan. 2700-year-old Chokhopani, similar to geographically close Shigatse Ü-Tsang Tibetan, could be fitted as the result of admixture of 0.861 northern East Asian Xiaogao-related ancestry and 0.139 Onge related ancestry (**Table S20 and Figure 5**). Younger Nepal ancient could be modeled as higher ancestry from Onge-related ancestry and lower ancestry associated with northern East Asian lineage. Jomon could be modeled as deriving 0.484 of its ancestry from populations related to Xiaogao_EN and 0.516 from groups related to Onge with marginal statistical significance. We substituted Boshan_EN and Bianbian_EN with Xiaogao_EN, we could obtain similar results, however, when we substituted Xiaojingshan_EN with Boshan_EN, 1500-year-old Samdzong failed to fit our two-way model (p_rank1= 0.00007). Zhalainuoer_EN-Onge could be successfully fitted highland Tibet Tibetan and Yunnan Tibetan with higher Onge-related ancestry but failed to the other Ando and Kham Tibetans. Using middle Neolithic East Asian as the source, the Xiaowu_MN-Onge model failed to all targets, and the DevilsCave_N-Onge model could be only fitted the Sichuan Tibetans, Jomon, and Chokhopani with a higher proportion of Onge-related ancestry. Except for populations with a western Eurasian affinity (Ando Tibetans and Samdzong), all remaining modern and ancient populations could be fitted as the admixture between Onge and middle Neolithic Wanggou_MN (Tibetan: ranging from 0.898 in Shannan Tibetan to 0.96 in Xinlong Tibetan; 0.518 for Jomon; 0.889 for Mebrak and 0.914 for Chokhopani), Banlashan_MN (0.795 in Shannan to 0.847 in Xinlong for Tibetan, 0.458 for Jomon and 0.8 for Chokhopani) and Miaozigou_MN (0.906 to 0.952 for Tibetans, 0.615 for Jomon, 0.906 for Mebrak and 0.933 for Chokhopani). We further used the late Neolithic northern East Asian as the source, Gangcha, and Gannan Tibetans and Samdzong failed in all models except for Gangcha Tibetan in Wadian_LN-Onge model (0.932: 0.068) and Haojiatai_LN-Onge (0.973: 0.027), Samdzong in Haojiatai_LN-Onge (0.908: 0.092). All the remaining populations could be fitted as the admixture of higher late Neolithic East Asians related ancestry and smaller Onge related ancestry. We additionally substituted Hoabinhian with Onge as the southern source representative for deep lineage and used aforementioned early Neolithic to late Neolithic northern East Asians as the other source to perform the two-way admixture model for estimating ancestry proportion of modern Tibetan without Gangcha and Gannan Tibetans and Nepal ancients except for Samdzong and Jomon. As shown in **Figure 5**, the good fit could be acquired with slightly variable ancestry composition compared with Onge-based two-way models. We finally employed the Afanasievo (significant negative *f_3_* value in admixture-*f_3_*-statistics) as the western Eurasian source in a three-way admixture model to fit the genetic variations in Ando Gangcha and Gannan Tibetans and Samdzong. All three populations could be successfully fitted when we introduced the Bronze Age steppe pastoralists related ancestry.

**Figure 5.**
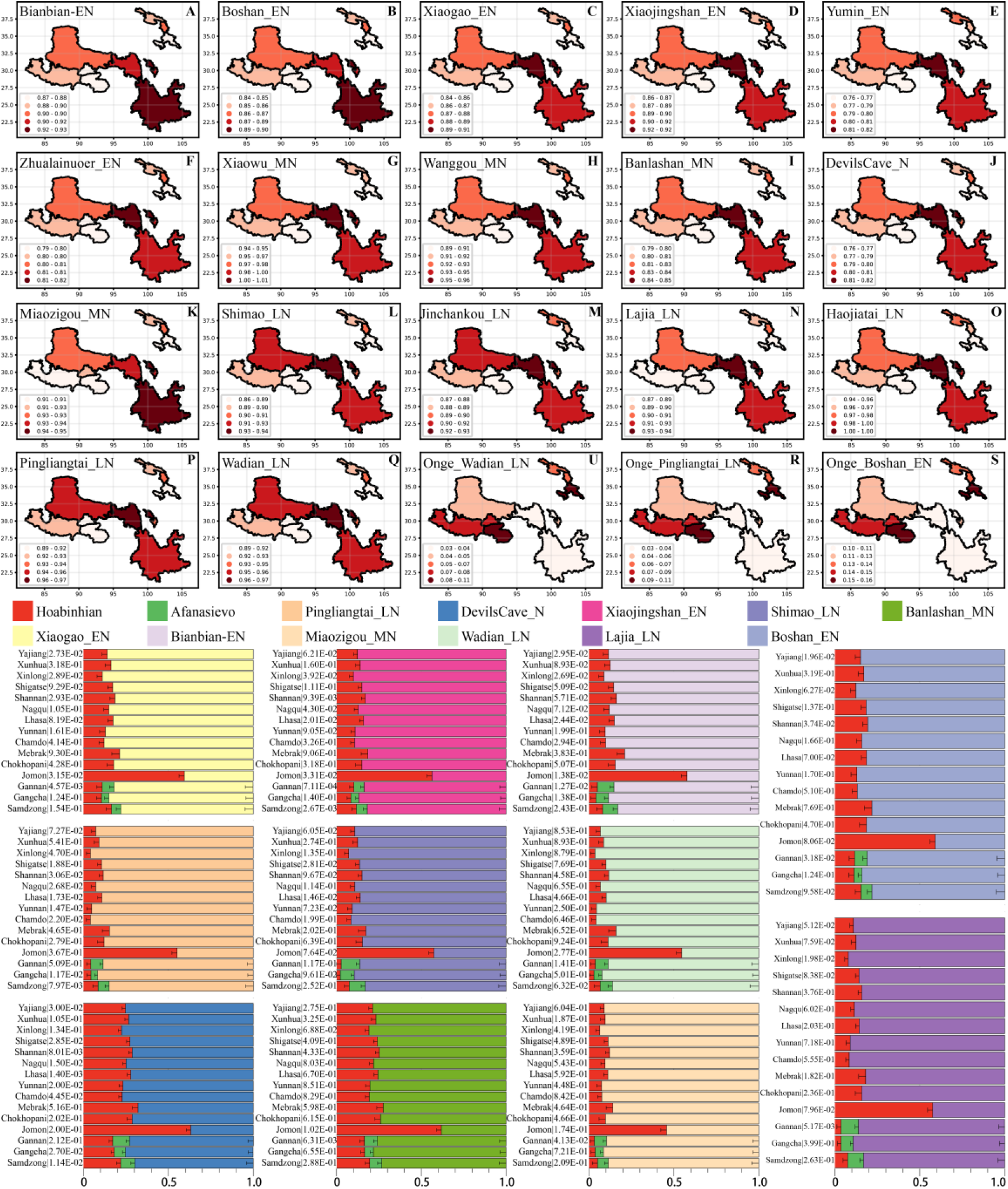
Results of qpAdm showed the main ancestry composition of modern and ancient Tibetans and Jomon Hunter-Gatherer were the results of the mixing of ancient northern East Asian and one deep lineage associated with South Asian Hunter-Gatherer Onge or Southeast Hunter-Gatherer Hoabinhian (the early Asian). Heatmap showed the Northern East Asian related ancestry in the two-way admixture model of Onge and the early Neolithic East Asian (**A~F**), middle Neolithic northern East Asian (**J~K**) and late Neolithic northern East Asian (**L~Q**). Onge-related ancestry was presented with three cases (**U~S**). Bar plots showed the ancestry composition of two-way model of Hoabinhian and East Asian for modern Tibetan, Jomon and Ancient Nepal Mebrak and Samdzong people, and three-way model for Qinghai and Gansu Tibetans.

Finally, to comprehensively summarize the phylogenetic relationships and reconstruct the population history between Neolithic East Asians and modern Tibetans in one phylogenetic framework, we built a series of admixture graph models via *qpGraph*. The core model of our admixture graph included archaic Denisovan and central African Mbuti as the roots, Loschbour as the representative of western Eurasian, modern Onge Hunter-Gatherer from Andaman island and 40,000-year-old Tianyuan (3% ancestry from Denisovan) as representatives of deep lineages of southern East Eurasian and northern East Eurasian. As shown in **Figure 6A**, East Asians diverged into northern lineage (represented by East Mongolia Neolithic population with 1% gene flow from western Eurasian) and southern lineage (represented by Liangdao2_EN with 35% ancestry deriving from lineages close to Onge). Here, late Neolithic Qijia-related Lajia people could be fitted as the admixture of 84% lineage related to northern East Asians and 16% lineage associated with south Asian Onge, and 2750-year-old Chokhopani could be modeled as driving 86% of its ancestry from Lajia_LN and 14% from Onge side. Our model provided ancient genomic evidence of the co-existence of both Paleolithic Hunter-Gather ancestry associated with the indigenous TP people and Neolithic northern East Asian ancestry in Chokhopani culture-related ancient Tibetan and late Neolithic Lajia people. We subsequently added all eleven modern Tibetans to the aforementioned scaffold model and found all Ü-Tsang Tibetans and Kham Tibetans except for Xinlong Tibetan could be fitted as direct descendants from 2,700-year-old Chokhopani with additional gene flow from one northern East Asian ancestry populations, which also contributed additional 33% ancestry to Iron Age Hanben people. This gene flow could be regarded as the epitome of the second wave of Neolithic expansion into TP. Thus, results from **Figure 6** suggested that seven Tibetans could be fitted well with three sources of ancestry: Onge-related, Lajia_LN-related and second wave of northern East Asian lineage-related, in respective proportion of 0.1235, 0.8265, and 0.05 (Shannan); 0.144, 0.816, and 0.04 (Shigatse); 0.1344, 0.8256, and 0.04 (Lhasa), 0.1176, 0.7224, and 0.16 (Nagqu); 0.1001, 0.6699, and 0.23 (Chamdo); 0.1106, 0.6794, and 0.21 (Yunnan); 0.1232, 0.7568, and 0.12 (Yajiang). We could obtain a good fit when considering one gene flow event for Ganqing Ando Tibetans with the Loschbour-related ancestry proportion varying from 2% to 3% (**Figure 7**). To further explore the best ancestral source proximity of the second migration wave, extended admixture graphs introducing inland/coastal northern and southern East Asian Neolithic populations were reconstrued. As shown in **Figure 8**, the second wave into lowland Kham Tibetans with Neolithic southern East Asian affinity could be well fitted as directly driving from Hanben-related ancestral population with the proportion ranging from 5% to 11%. We then added northern coastal early Neolithic Houli Boshan People, middle Neolithic Xiaowu Yangshao people, late Neolithic Wadian people and Bronze/Iron Age Haojiatai Shangzhou people to our core model in **Figure 6** and then fitted all Tibetans on it. We found that Yunnan Kham Tibetan harbored 33% additional ancestry associated with Longshan people, Sichuan Yajiang Kham Tibetan with 26% additional Longshan-related ancestry population (**Figure 9**). It is interesting to found that the gene pool of the Lhasa Ü-Tsang Tibetan was also influenced by the second population migration associated with Longshan people. This second gene flow event persisted when we substituted Longshan People as other Neolithic or Bronze/Iron Age populations with the acceptable ancestry proportions (**Figures S105~107**), these phenomena may be caused by genetic stability of the main ancestry in Central Plain (Henan and Shandong provinces).

**Figure 6.**
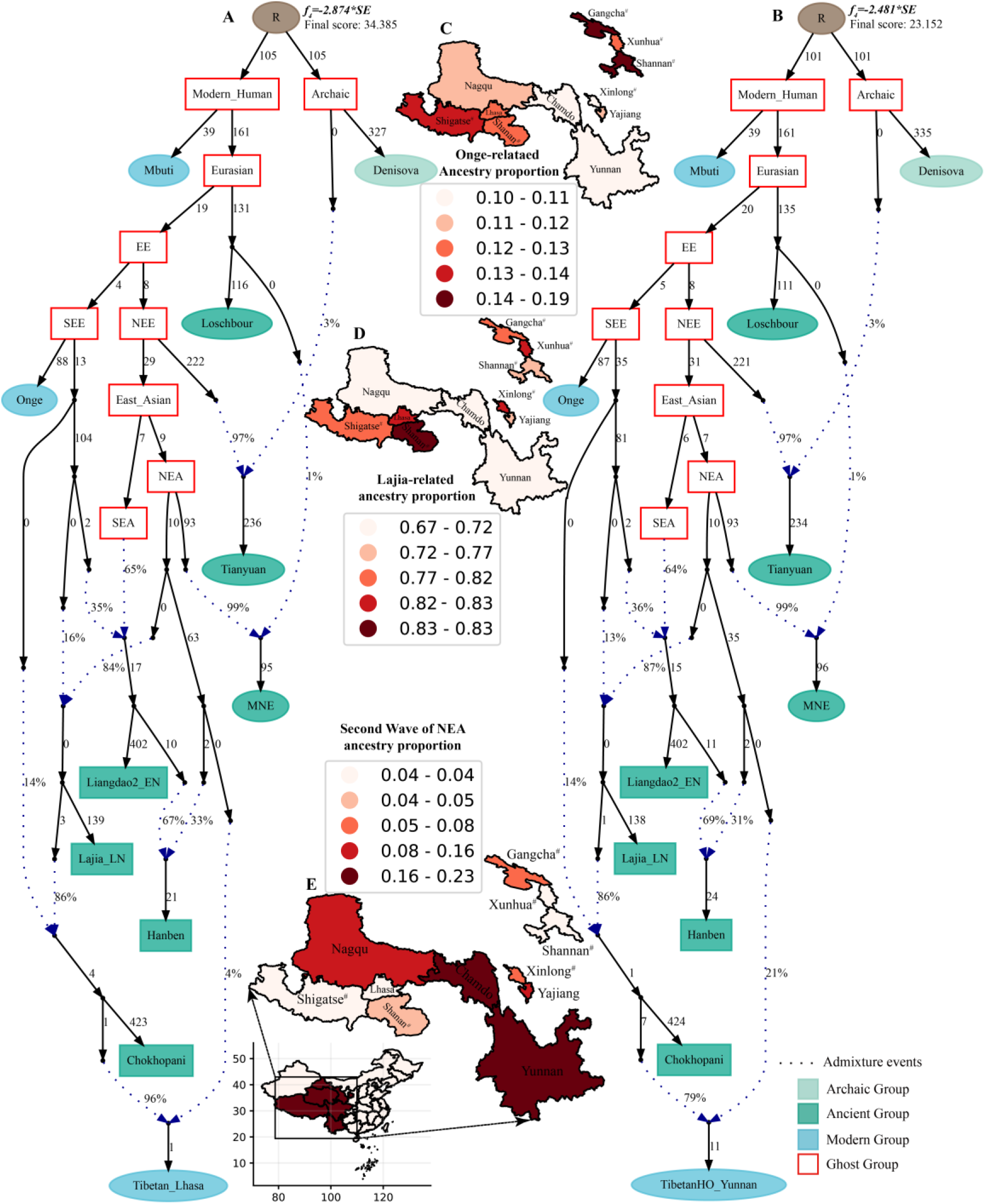
Admixture graph model of East Asians and modern Tibetans based on the Human Origin dataset. Admixture history of highland Tibetan from Lhasa (**A**) and lowland Tibetan from Yunnan (**B**). Heatmap showed the ancestry composition of modern Tibetans from three source populations: deep hunter-gatherer One-related ancestry (**C**), the first batch of Neolithic farmer associated ancestry (**D**) and the second batch of Neolithic farmer related ancestry (**E**). Denisovan and Central African of Mbuti were used as the Archaic and modern roots respectively. Western Eurasian was represented by Loschbour. Deep southern Eurasian (SEE) and northern Eurasian (NEE) were represented by South Asian Hunter-Gatherer of Onge and 40,000-year-old Tianyuan people. East Asian was subsequently diverged as northern East Asian (NEA) and southern East Asian (SEA). All f_4_-statistics of included populations are predicted to within 3 standard errors of their observed values. Branch lengths are given in units of 1000 times the f_2_ drift distance (rounded to the nearest integer). Pound signs denoted the modern populations added to the basic model of A or B with larger Z-scores or Zero internal branch length. Blue dotted lines denoted admixture events with admixture proportions as shown.

**Figure 7.**
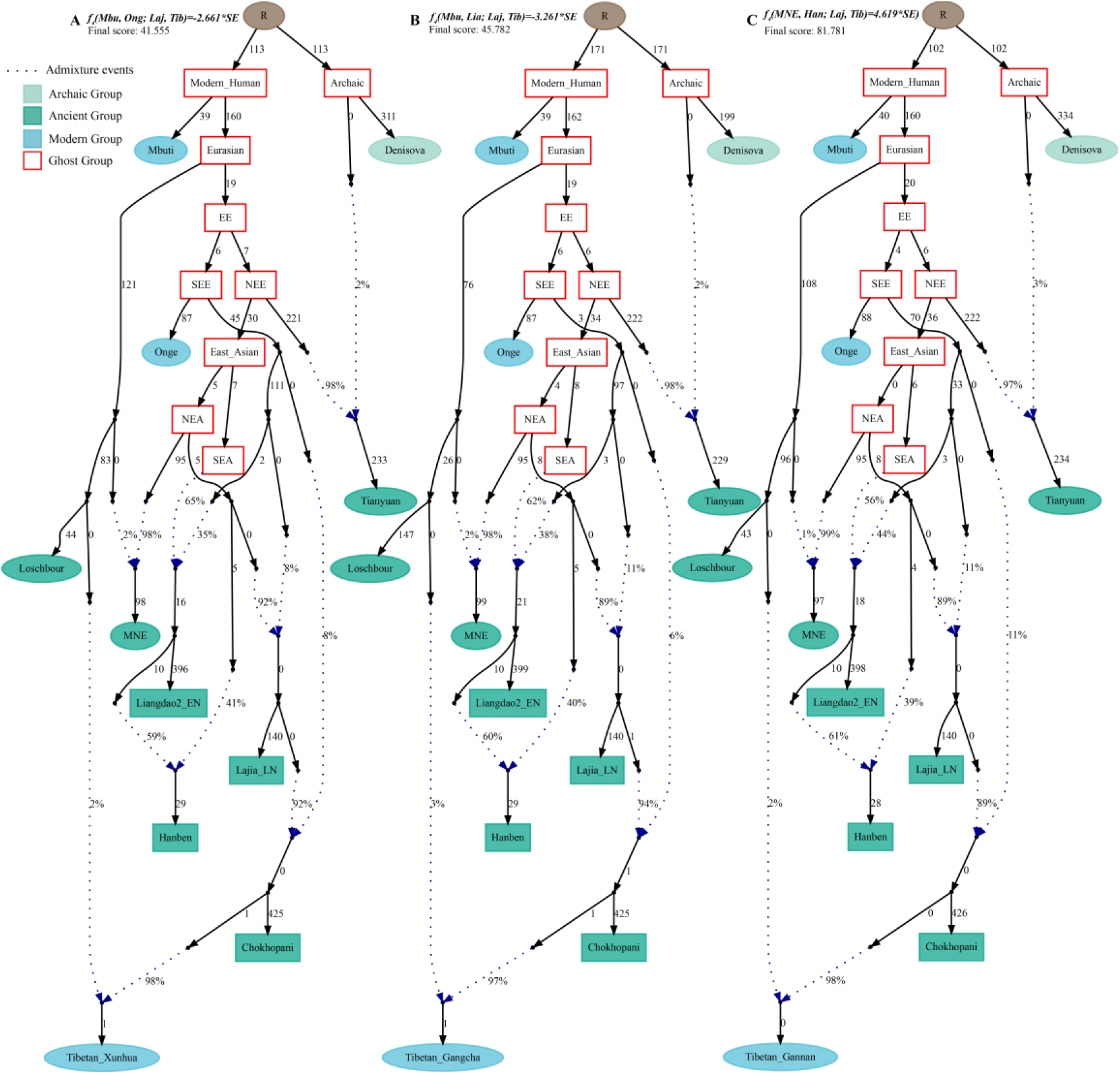
Admixture graph model of East Asians and modern Tibetans from the northeast Tibet Plateau based on the Human Origin dataset. Admixture history of Tibetan from Xunhua (**A**), Tibetan from Gangcha (**B**) and Tibetan from Gannan (**C**).

**Figure 8.**
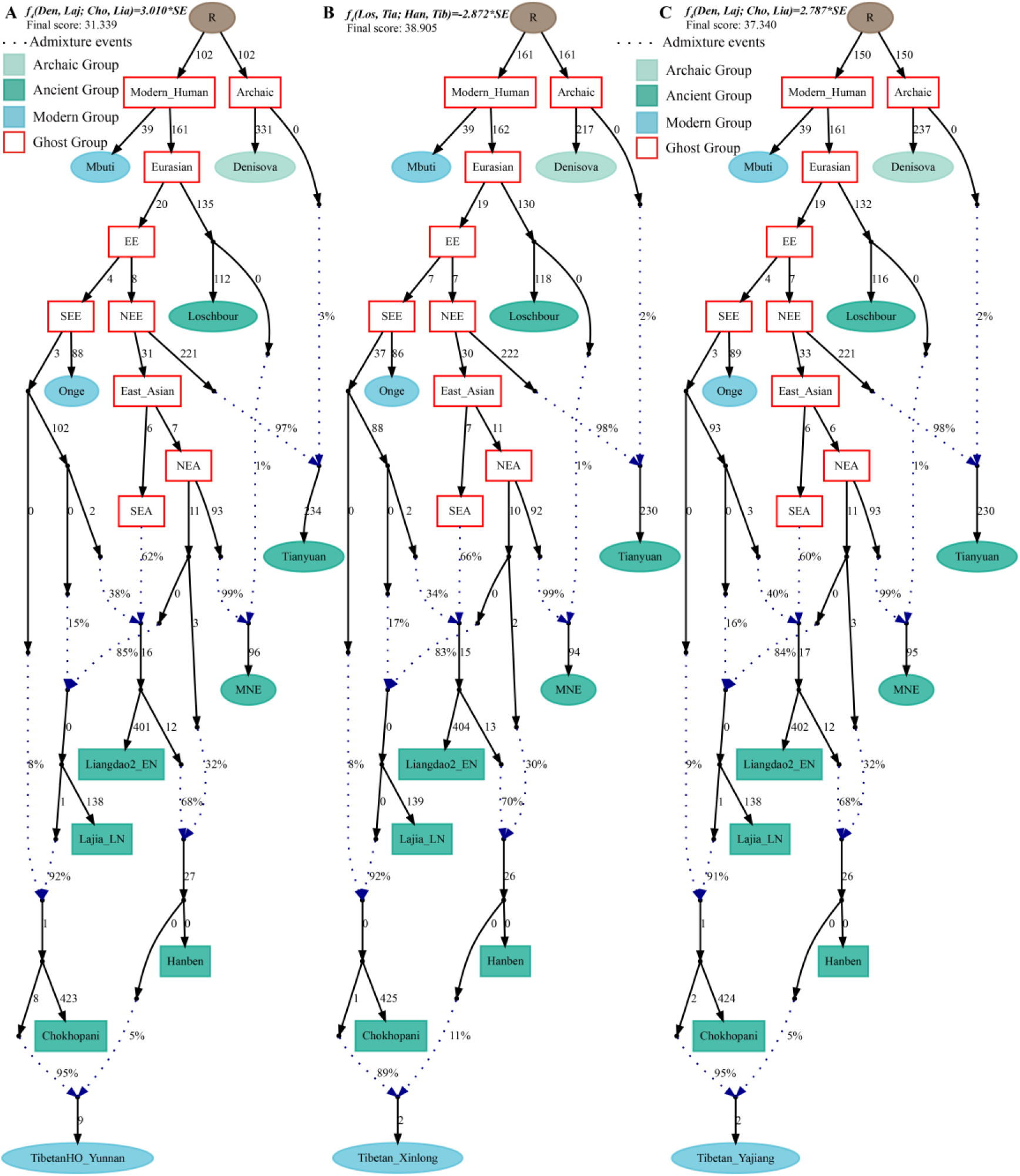
Admixture graph model of East Asians and modern lowland Tibetans based on the Human Origin dataset. Admixture history of lowland Tibetan from Yunnan (**A**), Tibetan from Xinlong (**B**) and Tibetan from Yajiang (**C**).

**Figure 9.**
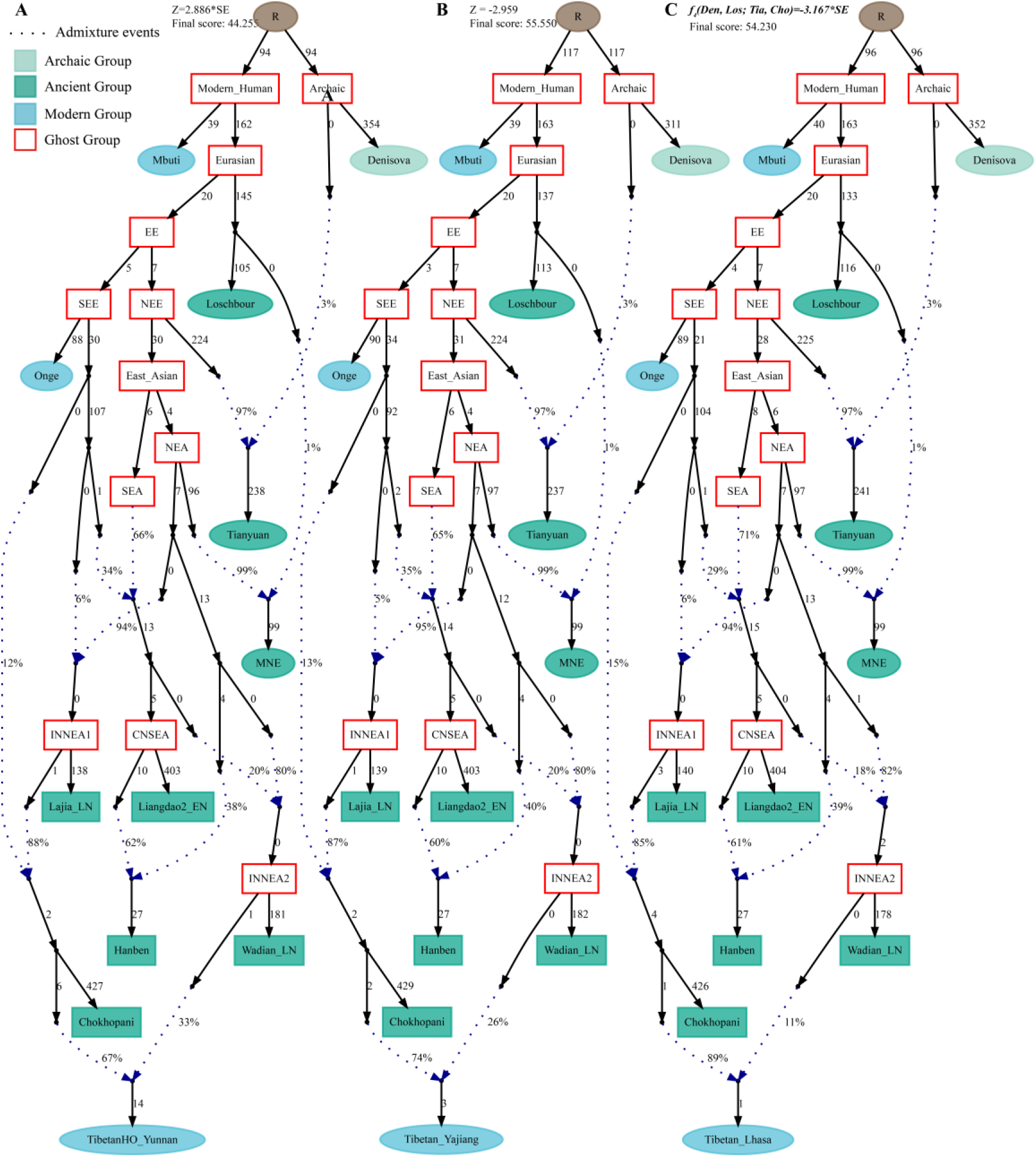
Admixture graph model of modern highland and lowland Tibetans based on the Human Origin dataset using late Neolithic Wadian people as the source of the second migration into Tibet Plateau. Admixture history of lowland Tibetan from Yunnan (**A**), Tibetan from Yajiang (**B**) and highland Tibetan from Lhasa (**C**).

## Discussion

Prehistoric human activities and the origin of the high-altitude adaptive modern Tibetans are the research topic in a variety of disciplines, mainly including genetics, archaeology, anthropology, history, and literature. Recent genome-wide sequencing and Paleo-genomic researches have been revolutionizing the knowledge of peopling of Europe(Narasimhan, et al. 2019), Central/South Asia(Narasimhan, et al. 2019), America(Nakatsuka, et al. 2020), Africa(Skoglund, et al. 2017) and Oceania(Lipson, Skoglund, et al. 2018). More and more ancient DNA studies from the surrounding of East Asia have been conducted and reported the population dynamics in Southeast Asia(Lipson, Cheronet, et al. 2018; McColl, et al. 2018) and South Siberia or Eurasia’s Eastern Steppe(Lazaridis, et al. 2014; Raghavan, et al. 2014; Mathieson, et al. 2015; Damgaard, et al. 2018; Sikora, et al. 2019), but lack in China. Fortunately, six ancient DNA studies from China(Yang, et al. 2017; Ning, et al. 2019; Ning, et al. 2020; Wang, Yeh, et al. 2020; Yang, et al. 2020) have been recently published to elucidate the prehistory of East Asian independently with 161 Paleolithic to historic (ranging from 40,000 ybp to 300 ybp). Yang et al. sequenced 40,000-year-old Tianyuan people from Beijing and found the early Asian population structures existed before the divergence between East Asian and Native American and the peopling of America by anatomically modern human populations(Yang, et al. 2017). Yang and Fu et al. recently conducted another ancient DNA work focused on 24 ancient genomes from Neolithic northern East Asia (eight samples), Neolithic southern East Asia (fifteen samples) and one historic Chuanyun people and found the north-south genetic differentiation among East Asian persisted since the early Neolithic period due to the observed significant genetic differences between Neolithic Shandong people and Fujian people based on multiple statistical methods(Yang, et al. 2020). Besides, they also identified southward migration from Shandong Houli populations and northward migration from Fujian Tanshishan populations, as well as a Neolithic coastal connection from southeastern Vietnam to Russia Far East, and a Pro-Austronesian connection between southern East Asians and southeast Pacific Vanuatu islanders. Besides, Ning et al. reported the population history of North China with fifteen ancient genomes from the Yellow River, West Liao River and Amur River, and discovered the subsistence strategy changes were associated with the population movements and admixtures(Ning, et al. 2020). Ning and Wang et.al. also reported ten Iron Age Shirenzigou people and found the Yamnaya-related steppe pastoralists mediated the population communications between East Asian and western Eurasian and dispersed Indo-European language into the Northwest China(Wang, Yeh, et al. 2020). Although these progresses have been achieved, the population history, genetic relationship and genetic differentiation between highland and lowland modern and ancient East Asians still kept in its infancy and remained to be clarified. Thus, we collected nineteen TP-related Neolithic to historic ancients, seventy-eight modern Tibetans from Ü-Tsang, Ando and Kham Tibetan regions, as well as all available eastern Eurasian ancients with different prehistoric human cultural backgrounds (including Tanshishan culture: Tanshishan and Xitoucun; Houli culture: Xiaogao, Xiaojingshan, Bianbian, and Boshan; Yangshao culture: Wanggou and Xiaowu; Longshan culture: Pingliangtai, Haojiatai, and Wadian; Hongshan culture: Banlashan) as well as modern Eurasians from Indo-European, Altai, Uralic, Sino-Tibetan, Austronesian, Austroasiatic, Hmong-Mien and Tai-Kadai language families and conducted one comprehensive Paleolithic to present-day ancient and modern genomic meta-analysis to provide new insights into the peopling of TP and clarify the relationship between high-altitude adaptive modern and ancients and lowland modern and ancient East Asians.

Modern and ancient genomes from TP showed a clear connection with northern modern Han Chinese and Neolithic northern East Asians (especially with coastal Houli people from Shandong, inland Yangshao and Longshan people from Henan and Qijia people from Ganqing region), which suggested the northern China-origin of modern Tibeto-Burman speaking populations. Three hypotheses have been proposed to elucidate the origin of the Sino-Tibetan language family based on the linguistic diversity and others(Zhang, et al. 2019), including northern China-origin associated with Yangshao/Majiayao hypothesis, southwestern Sichuan-origin hypothesis, and northeastern India-origin hypothesis. Based on the farming/language dispersal hypothesis, and the similarities of material culture assemblage among TP, East Asia, South/Central Asia, and Siberia, the origin of modern and ancient Tibetans is still confused and unclear(Jeong, et al. 2016). Shared ancestry revealed by our PCA, pairwise Fst genetic distance and outgroup-*f_3_* values, ADMIXTURE, and *f_4_*-statistics among modern and ancient highlanders and northern East Asian lowlanders showed their close relationship among them, which is consistent with genetic similarities revealed by forensic low-density genetic markers and uniparental haplotype/haplogroup data(Zou, et al. 2018; Chen, Wu, et al. 2019; He, et al. 2019). Direct evidence supported and confirmed this proposed common origin of Sino-Tibetan (northern China-origin associated with Yangshao/Majiayao hypothesis) was provided by the phylogenetic relationship reconstruction. Both TreeMix and qpGraph based on phylogenetic results supported the main ancestry in modern Tibetans, ancient TP people (Nepal and Qijia ancients) were derived from the common northern East Asian lineage related to East Mongolia Neolithic people and Yangshao/Longshan/Houli people from Central Plain. Thus, our results in this meta-genomic analysis supported the main lineage of TP people was originated from the lower and middle Yellow River with the Neolithic expansion of millet farmer. Our Neolithic to present-day autosomal genome-based findings confirmed the origin, diversification, and expansion of the modern Sino-Tibetan population revealed by mitochondrial and Y-chromosome variations(Wang, Lu, et al. 2018; Li, Tian, et al. 2019b).

Although strong evidence for the common origin of Sino-Tibetan speakers was provided, we still identified the differences in their ancestry composition. Compared with the Highlanders in TP, lowland late Neolithic to present-day harbored more ancestry related to Neolithic southern East Asians and Siberians. Iron Age Dacaozi people from Ganqing Region also showed a closer genetic affinity with southern people from Tanshishan culture, which showed the genetic trace of the northward dispersal of rice farmers. Compared with lowland Yangshao/Longshan or coastal Houli populations, the highland populations harbored a certain (8%~14%) proportion of Paleolithic Hunter-Gatherer ancestry related to Onge or Hoabinhian populations. Thus, our meta-analysis provided new evidence for the co-existence of both Paleolithic ancestries and Neolithic ancestries in the gene pool of East Asian Highlanders and a paleolithic colonization and Neolithic expansion of TP people, which was previously clarified via modern whole-sequence and mitochondrial and Y-chromosomal data(Qi, et al. 2013; Wang, Lu, et al. 2018; Li, Tian, et al. 2019b). Additionally, we also found obvious population substructures among modern Tibetans: Ü-Tsang Tibetans in Tibet core region haring predominant original Paleolithic and Neolithic ancestries, Ando Tibetans from Ganqing region in northwest China owing 2~3% western Eurasian admixture ancestry via qpGraph-based model and Kham Tibetans from Sichuan and Yunnan provinces possessing stronger southern Neolithic East Asian affinity. Thus, population substructures observed in modern Tibetan were consistent with the geographic and cultural divisions, which suggested that the complex cultural backgrounds and terrain to some extent served as the barriers for population movement and admixture. The second wave of population movements and admixtures well fitted in our qpGraph-based phylogeny revealed the gene flow from southern Iron Age East Asians to Kham Tibetans, from Neolithic northern East Asians to Kham and Ü-Tsang Tibetans, from western Eurasians to Ando Tibetans, which demonstrated multiple waves from Siberia, northern and southern East Asia have shaped the gene pool of East Asian highlander of Tibetan.

## Conclusion

Our comprehensive Neolithic to present-day genomic meta-analysis focused on eastern Eurasian (especially for China) was performed to clarify the relationships between Highlanders from TP and lowland East Asians and explore the peopling of TP for the first time. Results from our genetic survey showed a strong genetic affinity between ancient-modern Tibetans and Neolithic to present-day northern East Asian, which suggested that the main lineage of Tibeto-Burman speakers originated from Yangshao/Longshan people in the middle and lower Yellow River basin in North China with the common ancestor of Han Chinese and the dispersal of millet farmers and Sino-Tibetan languages. Although the shared ancestry persisted among ancient Tibetans and lowland Yangshao/Longshan/Houli people, we also found genetic differentiation among them: Highland Tibetans harboring deeply diverged eastern Eurasian Onge-related Hunter-Gatherer ancestry and lowland Neolithic to presented-day northern East Asians possessing more ancestry from Neolithic southern East Asian and Siberian ancestries, suggesting co-existence of Paleolithic and Neolithic ancestries in modern and ancient Tibetans and the population history of paleolithic colonization and Neolithic expansion. Besides, consistent with the geographic/linguistic divisions, three population substructures were identified in modern Tibetans: higher Onge/Hoabinhian ancestry in Ü-Tsang Tibetans, more western Eurasian-related ancestry in Ando Tibetans, and greater Neolithic southern East Asian ancestry in Kham Tibetan. Summarily, modern East Asian Highlanders derived ancestry from at least five ancient populations: Hoabinhian as the oldest layer; additional genetic materials from two Neolithic expansions (inland and coastal) from Northern East Asians, one Neolithic southern East Asian northwestward expansion and one western Eurasian eastward expansion.

## Methods and Materials

### Public dataset available

We collected 2,444 individuals from 183 geographically/culturally different populations(Patterson, et al. 2012; Lipson, Cheronet, et al. 2018; Jeong, et al. 2019; Liu, et al. 2020) belonging to fifteen language families or groups: Sinitic, Tai-Kadai, Tibeto-Burman, Tungusic, Turkic, Uralic, Austroasiatic, Austronesian, Caucasian, Chukotko-Kamchatkan, Eskimo-Aleut, Hmong-Mien, Indo-European, Japonic, Koreanic and Mongolic. 383 modern East Asians genotyped via Affymetrix Human Origins array (including 98 high-altitude adaptive Tibetans) by our group also used here(Wang, Yeh, et al. 2020). Besides, Paleolithic to historic ancient genomes from East Eurasian (Russia, China, Mongolia, Nepal and countries from Southeast Asia) were collected from recent ancient DNA studies or Reich Lab. A total of 161 Paleolithic to historic East Asians and eight Nepal ancients were collected and first comprehensively meta-analyzed and discussed in the present study(Jeong, et al. 2016; Yang, et al. 2017; Ning, et al. 2020; Wang, Yeh, et al. 2020; Yang, et al. 2020).

### Principal component analysis

We performed principal component analysis (PCA) with the *smartpca* program of the EIGENSOFT package(Patterson, et al. 2006), using default parameters and the lsqproject: YES and numoutlieriter: 0 options to project ancient samples onto the first two components.

### ADMIXTURE analysis

We carried out model-based clustering analysis using the ADMIXTURE (v. 1.3.0)(Alexander, et al. 2009) after pruning for linkage disequilibrium in PLINK v.1.9(Chang, et al. 2015) with parameters --indep-pairwise 200 25 0.4. We ran ADMIXTURE with the 10-fold cross-validation (--cv = 10), predefining the number of ancestral populations between K = 2 and K = 20 in 100 bootstraps with different random seeds.

### F-statistics

We computed *f_3_- and f_4_-statistics* using the *qp3Pop* and *qpDstat* programs as implemented in the ADMIXTOOLS(Reich, et al. 2009; Patterson, et al. 2012) with default parameters. We assessed standard errors using the weighted block jackknife approach.

### Admixture graph modeling

We ran TreeMix v.1.13(Pickrell and Pritchard 2012) with migration events ranging from 0 to 8 to construct the topology with the maximum likelihood. Based on the results of the *f-statistics*, we carried out graph-based admixture modeling using the *qpGraph* program as implemented in the ADMIXTOOLS package using Mbuti as an outgroup(Fu, et al. 2015).

### Streams of ancestry and inference of mixture proportions

We used the *qpWave* and *qpAdm* as implemented in the ADMIXTOOLS(Haak, et al. 2015) to estimate mixture proportions with respect to a basic set of outgroup populations: Mbuti, Ust_Ishim, Russia_Kostenki14, Papuan, Australian, Mixe, MA1 and Mongolia_N_East

## Acknowledgements

This study was supported by the National Natural Science Foundation of China (31801040 and 31760309), Nanqiang Outstanding Young Talents Program of Xiamen University (X2123302), Fundamental Research Funds for the Central Universities (ZK1144), and Lanzhou Talent Innovation and Entrepreneurship Project (2018-RC-113).

## Legends of Tables

Table S1. Pairwise Fst genetic distance between eleven modern Tibetans and other 71 worldwide reference populations.

Table S2. Pairwise Fst genetic distance between eleven modern Tibetans, Yoruba and other 20 East Asian Neolithic to Bronze/Iron Age reference populations.

Table S3. The results of outgroup-f3(Modern Tibetans/other 278 Eurasian modern and Ancient population, Modern Tibetans/other 278 Eurasian modern and Ancient population; Mbuti)

Table S4. Admxiture-f3-statistics of the form f3(Source population1, Source population2; Shigatse Tibetan) showed the admixture signals for high-altitude adaptive Shigatse Tibetan

Table S5. Admxiture-f3-statistics of the form f3(Source population1, Source population2; Shannan Tibetan) showed the admixture signals for high-altitude adaptive Shannan Tibetan

Table S6. Admxiture-f3-statistics of the form f3(Source population1, Source population2; Lhasa Tibetan) showed the admixture signals for high-altitude adaptive Lhasa Tibetan

Table S7. Admxiture-f3-statistics of the form f3(Source population1, Source population2; Nagqu Tibetan) showed the admixture signals for high-altitude adaptive Nagqu Tibetan

Table S8. Admxiture-f3-statistics of the form f3(Source population1, Source population2; Chamdo Tibetan) showed the admixture signals for high-altitude adaptive Chamdo Tibetan

Table S9. Admxiture-f3-statistics of the form f3(Source population1, Source population2; Gannan Tibetan) showed the admixture signals for lowland Gannan Tibetan from Gansu province

Table S10. Admxiture-f3-statistics of the form f3(Source population1, Source population2; Gangcha Tibetan) showed the admixture signals for lowland Gangcha Tibetan from Qinghai province

Table S11. Admxiture-f3-statistics of the form f3(Source population1, Source population2; Xunhua Tibetan) showed the admixture signals for lowland Xunhua Tibetan from Qinghai province

Table S12. Admxiture-f3-statistics of the form f3(Source population1, Source population2; Xinlong Tibetan) showed the admixture signals for lowland Xinlong Tibetan from Sichuan province

Table S13. Admxiture-f3-statistics of the form f3(Source population1, Source population2; Yajiang Tibetan) showed the admixture signals for lowland Yajiang Tibetan from Sichuan province

Table S14. Admxiture-f3-statistics of the form f3(Source population1, Source population2; Yunnan Tibetan) showed the admixture signals for lowland Yunnan Tibetan

Table S15. Admxiture-f3-statistics of the form f3(Source population1, Source population2; Nepal_Samdzong_1500BP) showed the admixture signals for high-altitude adaptive1500-year-old Samdzong population from Nepal

Table S16. Admxiture-f3-statistics of the form f3(Source population1, Source population2; Nepal_Mebrak_2125BP) showed the admixture signals for high-altitude adaptive 2125-year-old Mebrak population from Nepal

Table S17. Admxiture-f3-statistics of the form f3(Source population1, Source population2; Lajia_LN) showed the admixture signals for late Neolithic Lajia population from the upper Yellow River Basin

Table S18. Admxiture-f3-statistics of the form f3(Source population1, Source population2; Dacaozi_IA) showed the admixture signals for Iron Age Dacaozi population from the upper Yellow River Basin

Table S19. Results of four population texts focused on the genetic differentiation within Tibetans

Table S20. Ancestry composition of two-way admixture Model

