## Supplementary Figures S1-49 for "Peopling of Tibet Plateau and multiple waves of admixture of Tibetans inferred from both modern and ancient genome-wide data": Supplementary Figures S1-49.pdf

**Supplementary Figures S1-S49**

**Peopling of Tibet Plateau and multiple waves of admixture of Tibetans inferred from both modern and ancient genome-wide data**

Mengge Wang[1,*], Xing Zou[1,*], Hui-Yuan Ye[2,*], Zheng Wang[1], Yan Liu[3], Jing Liu[1], Fei Wang[1], Hongbin Yao[4], Pengyu Chen[5], Ruiyang Tao[1], Shouyu Wang[1], Lan-Hai Wei[6], Renkuan Tang[7,#], Chuan-Chao Wang[6,#] , Guanglin He[1,6,#]

[1]Institute of Forensic Medicine, West China School of Basic Science and Forensic Medicine, Sichuan University, Chengdu, China
[2]School of Humanities, Nanyang Technological University, Nanyang, 639798, Singapore
[3]College of Basic Medicine, Chuanbei Medical University
[4]Belt and Road Research Center for Forensic Molecular Anthropology, Key Laboratory of Evidence Science of Gansu Province, Gansu University of Political Science and Law, Lanzhou 730070, China
[5]Center of Forensic Expertise, Affiliated hospital of Zunyi Medical University, Zunyi, Guizhou, China
[6]Department of Anthropology and Ethnology, Institute of Anthropology, National Institute for Data Science in Health and Medicine, and School of Life Sciences, Xiamen University, Xiamen, China
[7]Department of Forensic Medicine, College of Basic Medicine, Chongqing Medical University, Chongqing, China

*These authors contributed equally to this work and should be considered co-first authors.

#Corresponding author
Renkuan Tang
Department of Forensic Medicine, College of Basic Medicine, Chongqing Medical University, Chongqing, China

Chuan-Chao Wang
Affiliation: Department of Anthropology and Ethnology, Institute of Anthropology, National Institute for Data Science in Health and Medicine, Xiamen University,
Xiamen, China.

Guanglin He
Affiliation: Department of Anthropology and Ethnology, Institute of Anthropology, National Institute for Data Science in Health and Medicine, Xiamen University,
Xiamen, China.


**Contents of Supplementary Figures S1-S49**

**Figure S1**. ADMIXTURE results of all included populations. All ancestry composition was the mean proportion of one population. 1

**Figure S2**. The pairwise genetic distances between three highland Tibetans from Tibet Tibetan Autonomous Region (from left to right: Lhasa Tibetan, Shannan Tibetan and Shigatse Tibetan) and other 79 modern reference populations. 2

**Figure S3**. The pairwise genetic distances between two highland Tibetans from Tibet Tibetan Autonomous Region and one lowland She (from left to right: She from HGDP, Chamdo Tibetan and Nagqu Tibetan) and other 79 modern reference populations. 3

**Figure S4**. The pairwise genetic distances between Tibetans from Qinghai and Gansu in the northeast Tibet Plateau (from left to right: Qinghai Gangcha Tibetan, Gansu Gannan Tibetan and Qinghai Xunhua Tibetan) and other 79 modern reference populations. 4

**Figure S5**. The pairwise genetic distances between Tibetans from Sichuan and Yunnan in the lowland region (from left to right: Sichuan Xinlong Tibetan, Sichuan Yajing Tibetan and Yunnan Tibetan) and other 79 modern reference populations. 5

**Figure S6**. The pairwise genetic distances between our studied eleven Tibetans and other 20 East Asian reference populations. 6

**Figure S7**. The shared genetic drift between our studied Tibet Tibetans and other 43 spatial-temporally different East Asian populations. 7

**Figure S8**. The shared genetic drift between our studied Gansu/Qinghai Tibetans and other 43 spatial-temporally different East Asian populations. 8

**Figure S9**. The shared genetic drift between our studied Sichuan/Yunnan Tibetans and other 43 spatial-temporally different East Asian populations. 9

Intra population differentiation among high-altitude residing and low-altitude residing Tibetans. 10

**Figure S10**. Genomic similarities and differences among both high-altitude residing Tibetan and low-altitude residing Tibetans inferred from four population symmetry-$f_4$ statistics of the form *$f_4$(Tibetan1, Chamdo Tibetan; Tibetan2, Mbuti)*. 10

**Figure S11**. Genomic similarities and differences among both high-altitude residing Tibetan and low-altitude residing Tibetans inferred from four population symmetry-$f_4$ statistics of the form *$f_4$(Tibetan1, Shigatse/Shannan Tibetan; Tibetan2, Mbuti)*. 11

**Figure S12**. Genomic similarities and differences among both high-altitude residing Tibetan and low-altitude residing Tibetans inferred from four population symmetry-$f_4$ statistics of the form *$f_4$(Tibetan1, Lhasa/Nagqu Tibetan; Tibetan2, Mbuti)*. 12

**Figure S13**. Genomic similarities and differences among both high-altitude residing Tibetan and low-altitude residing Tibetans inferred from four population symmetry-$f_4$ statistics of the form *$f_4$(Tibetan1, Gangcha/Xunhua Tibetan; Tibetan2, Mbuti)*. 14

**Figure S14**. Genomic similarities and differences among both high-altitude residing Tibetan and low-altitude residing Tibetans inferred from four population symmetry-$f_4$ statistics of the form *$f_4$(Tibetan1, Gannan/Yunnan Tibetan; Tibetan2, Mbuti)*. 15

**Figure S15**. Genomic similarities and differences among both high-altitude residing Tibetan and low-altitude residing Tibetans inferred from four population symmetry-$f_4$ statistics of the form *$f_4$(Tibetan1, Yajiang/Xinlong Tibetan; Tibetan2, Mbuti)*. 17

**Figure S16**. Genomic affinity between modern Tibetans and eastern Eurasian ancient populations inferred from four population symmetry-$f_4$ statistics of the form *$f_4$(Tibetan1, Shigatse Tibetan;*

*eastern Eurasian ancients, Mbuti)*. 18

**Figure S17**. Genomic affinity between modern Tibetans and eastern Eurasian ancient populations inferred from four population symmetry-$f_4$ statistics of the form $f_4$*(Tibetan1, Shannan Tibetan; eastern Eurasian ancients, Mbuti)*. 20

**Figure S18**. Genomic affinity between modern Tibetans and eastern Eurasian ancient populations inferred from four population symmetry-$f_4$ statistics of the form $f_4$*(Tibetan1, Lhasa Tibetan; eastern Eurasian ancients, Mbuti)*. 22

**Figure S19**. Genomic affinity between modern Tibetans and eastern Eurasian ancient populations inferred from four population symmetry-$f_4$ statistics of the form $f_4$*(Tibetan1, Nagqu Tibetan; eastern Eurasian ancients, Mbuti)*. 24

**Figure S20**. Genomic affinity between modern Tibetans and eastern Eurasian ancient populations inferred from four population symmetry-$f_4$ statistics of the form $f_4$*(Tibetan1, Gangcha Tibetan; eastern Eurasian ancients, Mbuti)*. 26

**Figure S21**. Genomic affinity between modern Tibetans and eastern Eurasian ancient populations inferred from four population symmetry-$f_4$ statistics of the form $f_4$*(Tibetan1, Xunhua Tibetan; eastern Eurasian ancients, Mbuti)*. 28

**Figure S22**. Genomic affinity between modern Tibetans and eastern Eurasian ancient populations inferred from four population symmetry-$f_4$ statistics of the form $f_4$*(Tibetan1, Gannan Tibetan; eastern Eurasian ancients, Mbuti)*. 29

**Figure S23**. Genomic affinity between modern Tibetans and eastern Eurasian ancient populations inferred from four population symmetry-$f_4$ statistics of the form $f_4$*(Tibetan1, Xinlong Tibetan; eastern Eurasian ancients, Mbuti)*. 31

**Figure S24**. Genomic affinity between modern Tibetans and eastern Eurasian ancient populations inferred from four population symmetry-$f_4$ statistics of the form $f_4$*(Tibetan1, Yajiang Tibetan; eastern Eurasian ancients, Mbuti)*. 32

**Figure S25**. Genomic affinity between modern Tibetans and eastern Eurasian ancient populations inferred from four population symmetry-$f_4$ statistics of the form $f_4$*(Tibetan1, Yunnan Tibetan; eastern Eurasian ancients, Mbuti)*. 34

**Figure S26**. Shared ancestry associated with coastal Neolithic northern East Asian in modern Tibetans and ancient East Asians compared with geographically different coastal Neolithic northern East Asian inferred from four population symmetry-$f_4$-statistics of the form $f_4$*(Coastal Neolithic northern East Asian1, Coastal Neolithic northern East Asian2; Eastern Modern Tibetan/Ancient East Asians, Mbuti)* 36

**Figure S27**. Shared ancestry associated with inland Neolithic to Iron Age northern East Asian from Henan province in modern Tibetans and ancient East Asians compared with geographically different ancient Henan populations inferred from four population symmetry-$f_4$-statistics of the form $f_4$*(Haojiatai_LBIA, Neolithic to Iron Age Henan populations; Eastern Modern Tibetan/Ancient East Asians, Mbuti)* 37

**Figure S28**. Shared ancestry associated with inland Neolithic to Iron Age northern East Asian from Henan province in modern Tibetans and ancient East Asians compared with geographically different ancient Henan populations inferred from four population symmetry-$f_4$-statistics of the form $f_4$*(Jiaozuoniecun_LBIA, Neolithic to Iron Age Henan populations; Eastern Modern Tibetan/Ancient East Asians, Mbuti)* 39

**Figure S29**. Shared ancestry associated with inland Neolithic to Iron Age northern East Asian from

**Figure S48**. Spatial comparison analysis showed the shared ancestry related to late northern Neolithic East Asians in modern Tibetans and all available ancient East Asians inferred from four population symmetry-$f_4$-statistics of the form *$f_4$(Shimao_LN, Ancient Northern East Asians; Modern*

*Tibetan/Neolithic to Historic East Asians, Mbuti)*. ........ 62
**Figure S49**. Spatial comparison analysis showed the shared ancestry related to the Ancient Nepal population in modern Tibetans and all available ancient East Asians inferred from four population symmetry-$f_4$-statistics of the form $f_4$*(Chokhopani, Ancient Northern East Asians; Modern Tibetan/Neolithic to Historic East Asians, Mbuti)*. ........ 64

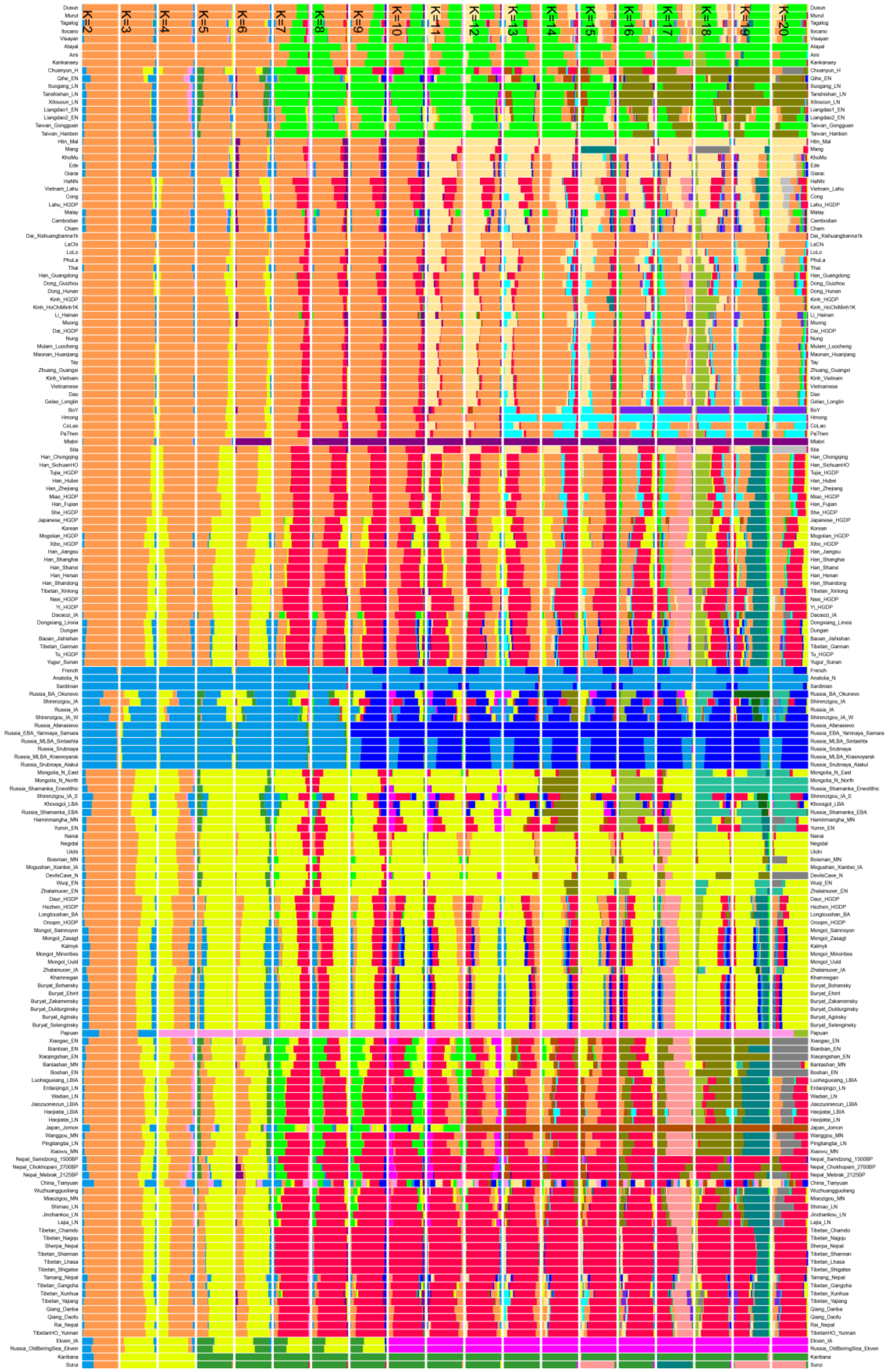

Figure S1. ADMIXTURE results of all included populations. All ancestry composition was the mean proportion of one population.

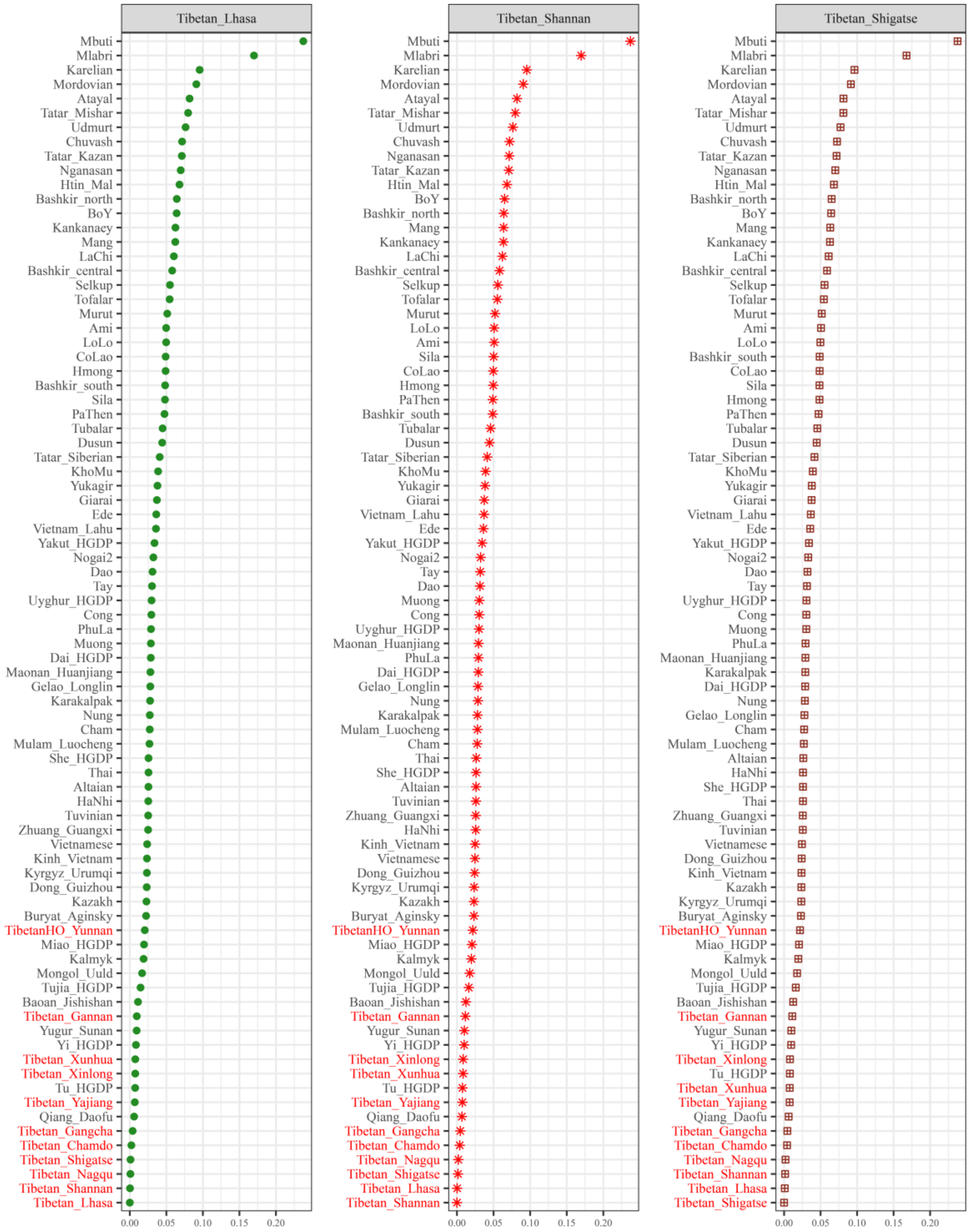

**Figure S2. The pairwise genetic distances between three highland Tibetans from Tibet Tibetan Autonomous Region (from left to right: Lhasa Tibetan, Shannan Tibetan and Shigatse Tibetan) and other 79 modern reference populations.**

All populations listed in the Y-axis were sorted according to the pairwise Fst genetic distances and the Fst values were marked in the X-axis. The small distance between our targeted Tibetan and reference populations showed the closer genetic relationship and stronger genetic affinity among them, otherwise, the larger Fst value denotes distant genetic relationships among them.

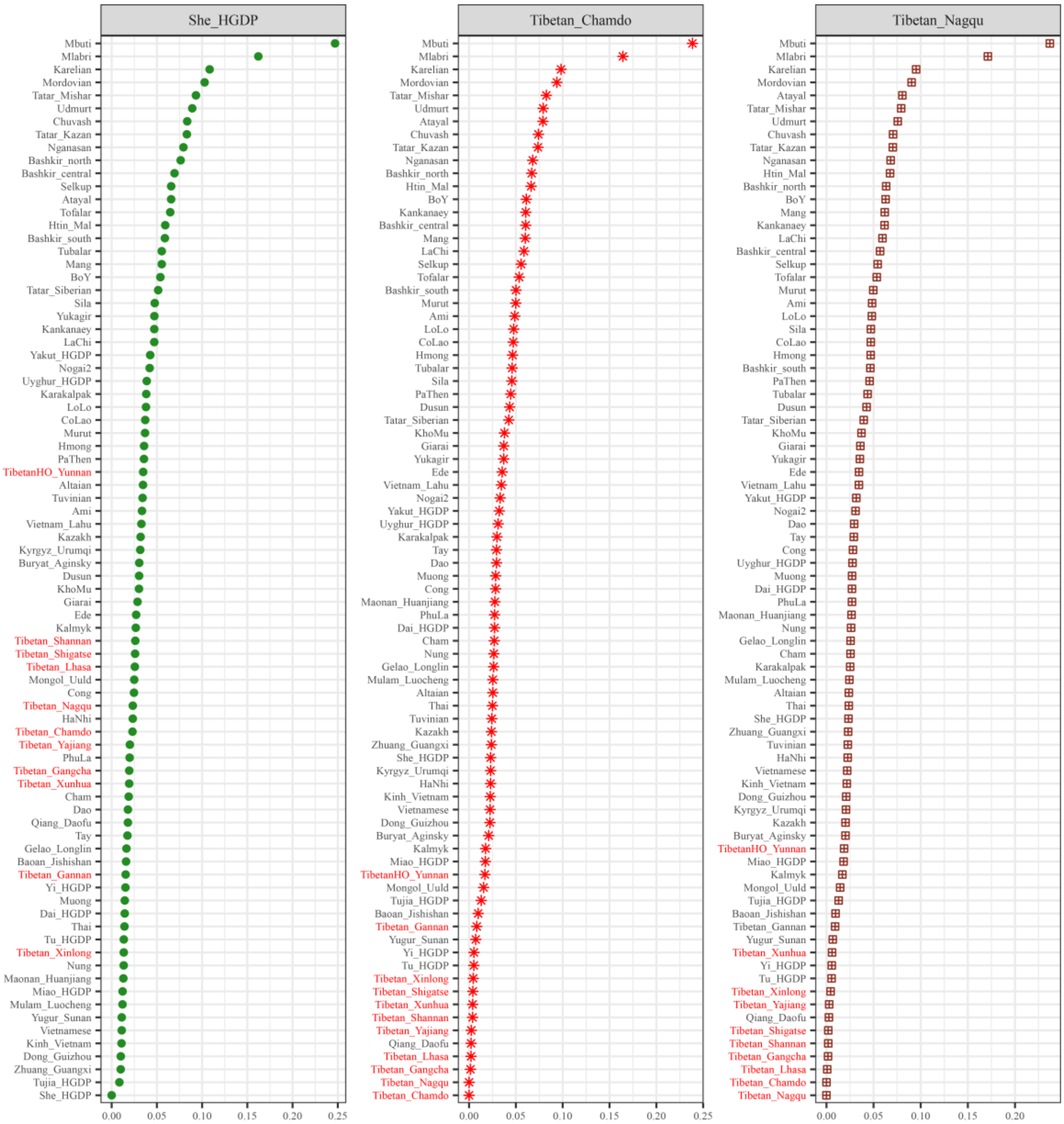

**Figure S3. The pairwise genetic distances between two highland Tibetans from Tibet Tibetan Autonomous Region and one lowland She (from left to right: She from HGDP, Chamdo Tibetan and Nagqu Tibetan) and other 79 modern reference populations.**

All populations listed in the Y-axis were sorted according to the pairwise Fst genetic distances and the Fst values were marked in the X-axis. The small distance between our targeted Tibetan and reference populations showed the closer genetic relationship and stronger genetic affinity among them, otherwise, the larger Fst value denotes distant genetic relationships among them.

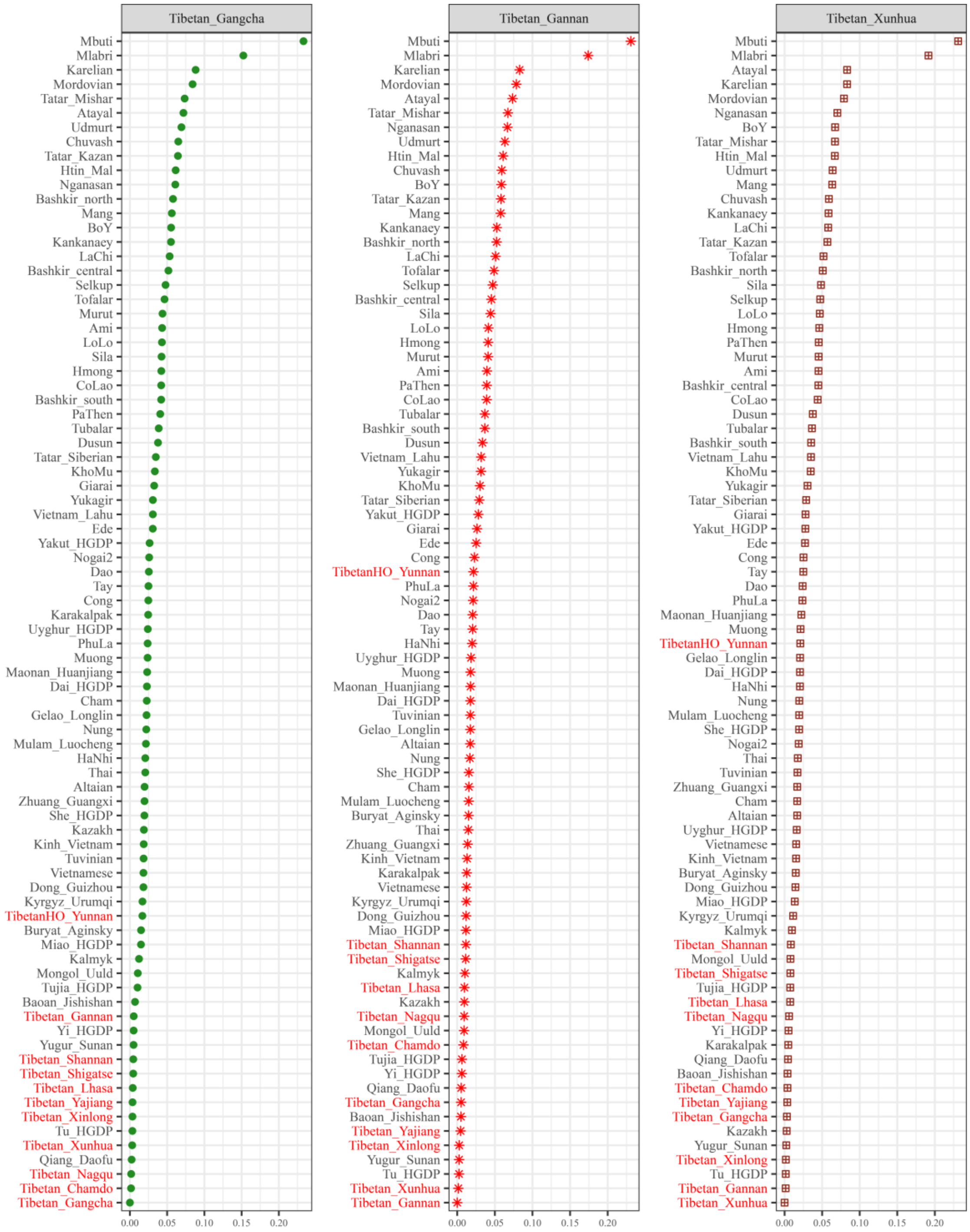

**Figure S4. The pairwise genetic distances between Tibetans from Qinghai and Gansu in the northeast Tibet Plateau (from left to right: Qinghai Gangcha Tibetan, Gansu Gannan Tibetan and Qinghai Xunhua Tibetan) and other 79 modern reference populations.**

All populations listed in the Y-axis were sorted according to the pairwise Fst genetic distances and the Fst values were marked in the X-axis. The small distance between our targeted Tibetan and reference populations showed the closer genetic relationship and stronger genetic affinity among them, otherwise, the larger Fst value denotes distant genetic relationships among them.

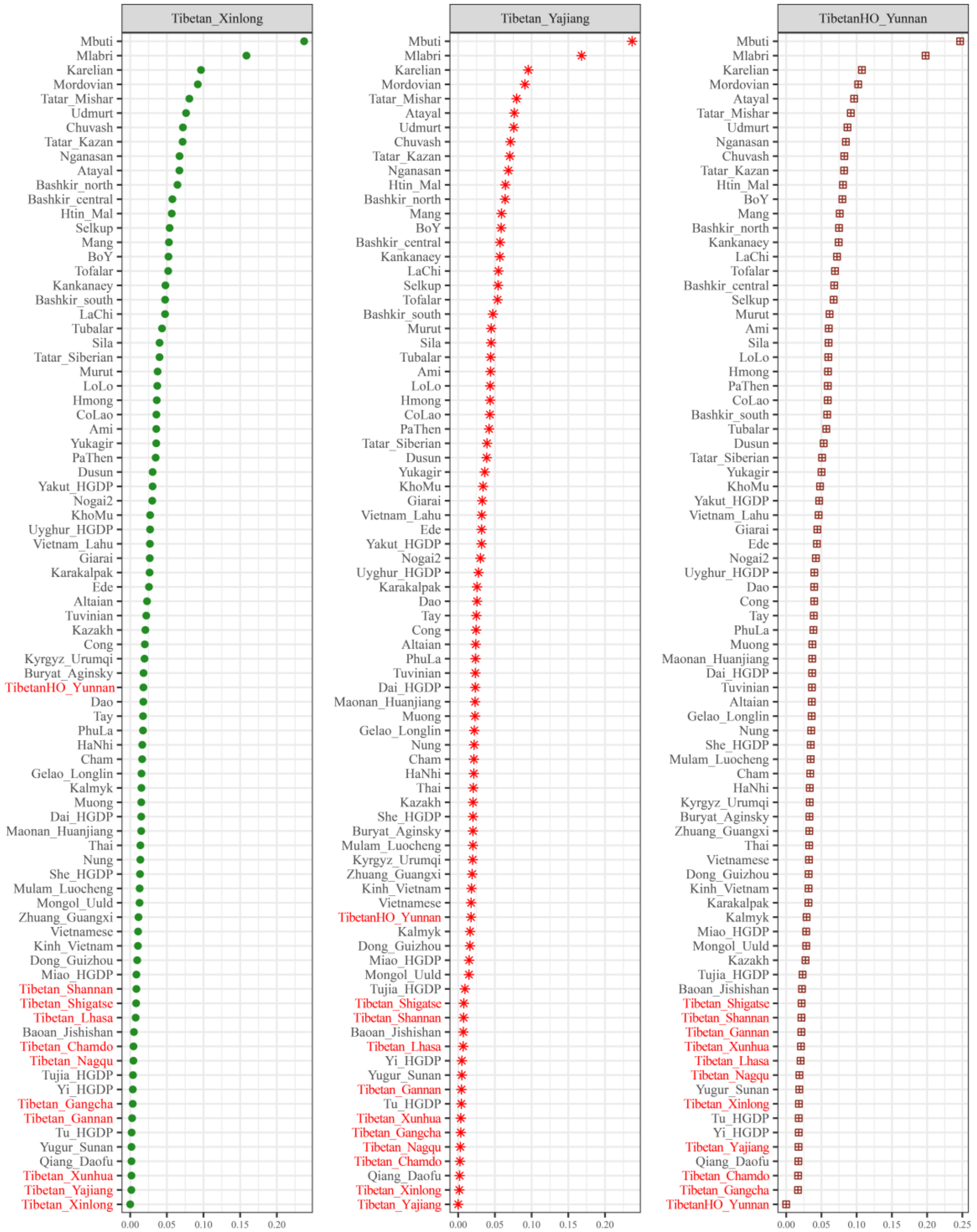

**Figure S5. The pairwise genetic distances between Tibetans from Sichuan and Yunnan in the lowland region (from left to right: Sichuan Xinlong Tibetan, Sichuan Yajing Tibetan and Yunnan Tibetan) and other 79 modern reference populations.**

All populations listed in the Y-axis were sorted according to the pairwise Fst genetic distances and the Fst values were marked in the X-axis. The small distance between our targeted Tibetan and reference populations showed the closer genetic relationship and stronger genetic affinity among them, otherwise, the larger Fst value denotes distant genetic relationships among them.

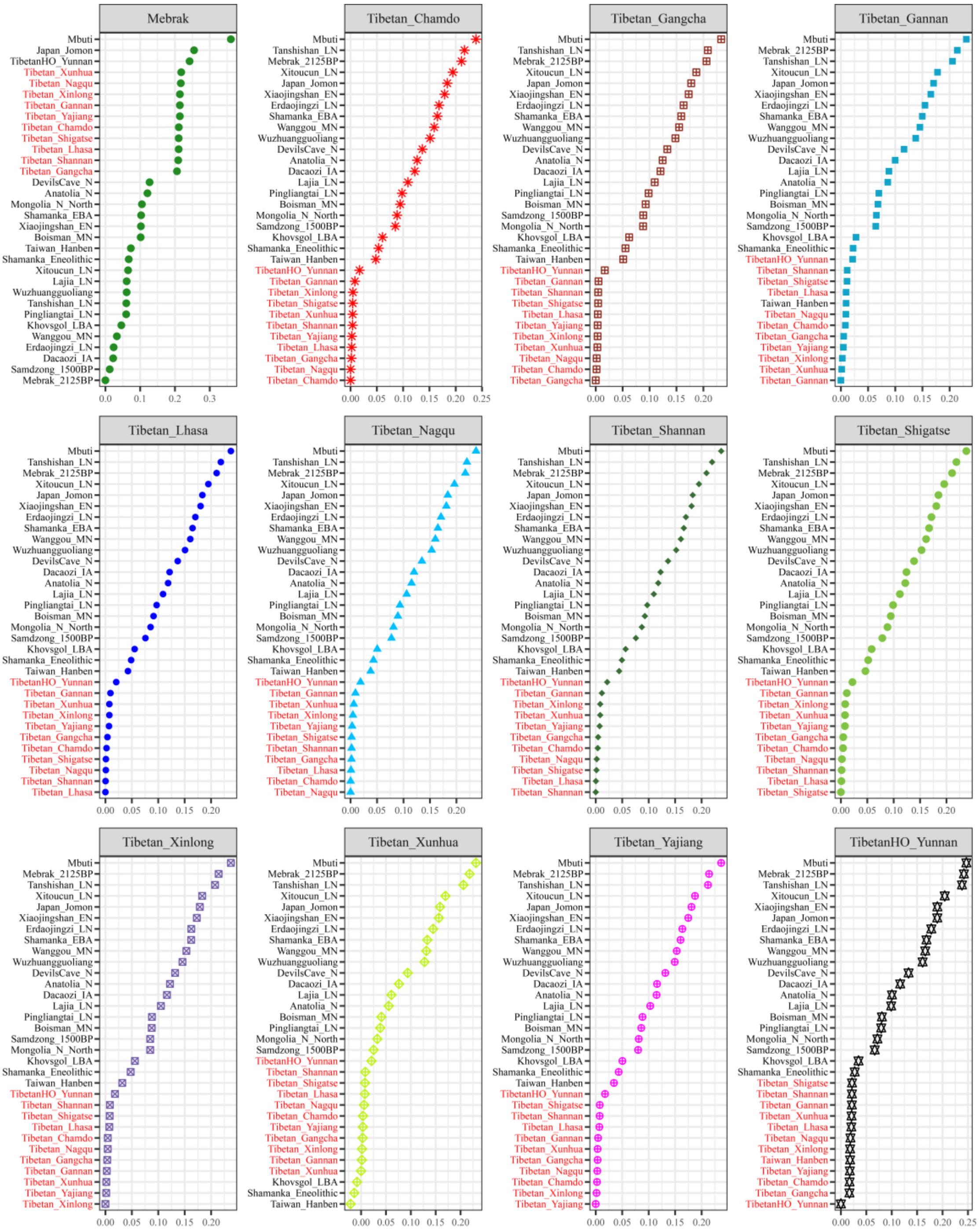

**Figure S6. The pairwise genetic distances between our studied eleven Tibetans and other 20 East Asian reference populations.**

All populations listed in the Y-axis were sorted according to the pairwise Fst genetic distances and the Fst values were marked in the X-axis. The small distance between our targeted Tibetan and reference populations showed the closer genetic relationship and stronger genetic affinity among them, otherwise, the larger Fst value denotes distant genetic relationships among them.

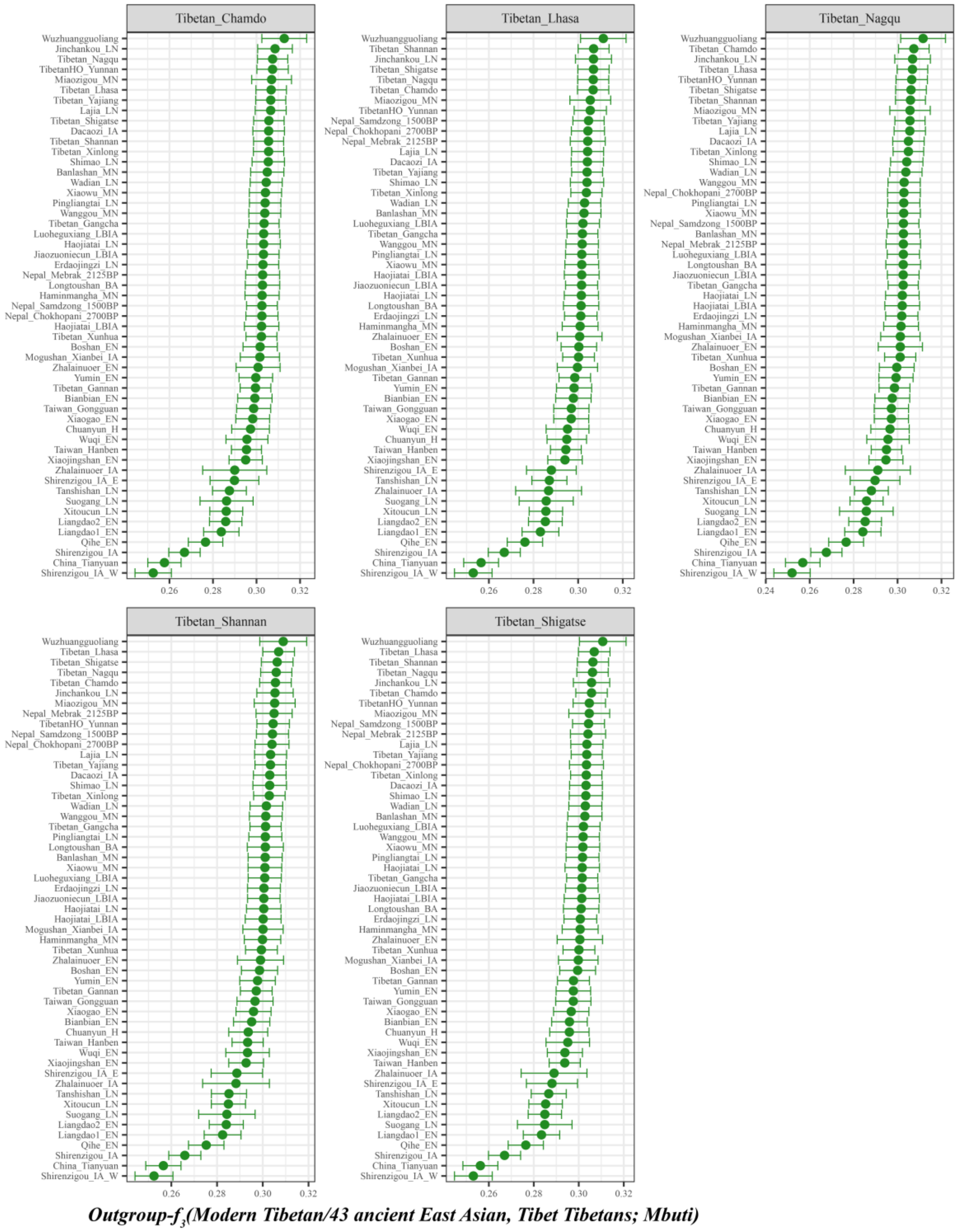

**Figure S7. The shared genetic drift between our studied Tibet Tibetans and other 43 spatial-temporally different East Asian populations.**

All populations listed in the Y-axis were sorted according to the outgroup-$f_3$ values and the outgroup-$f_3$ values were marked in the X-axis. The larger outgroup-$f_3$ value between our targeted Tibetan and reference populations showed the closer genetic relationship and stronger genetic affinity among them, otherwise, the smaller outgroup-$f_3$ value denotes distant genetic relationships among them.

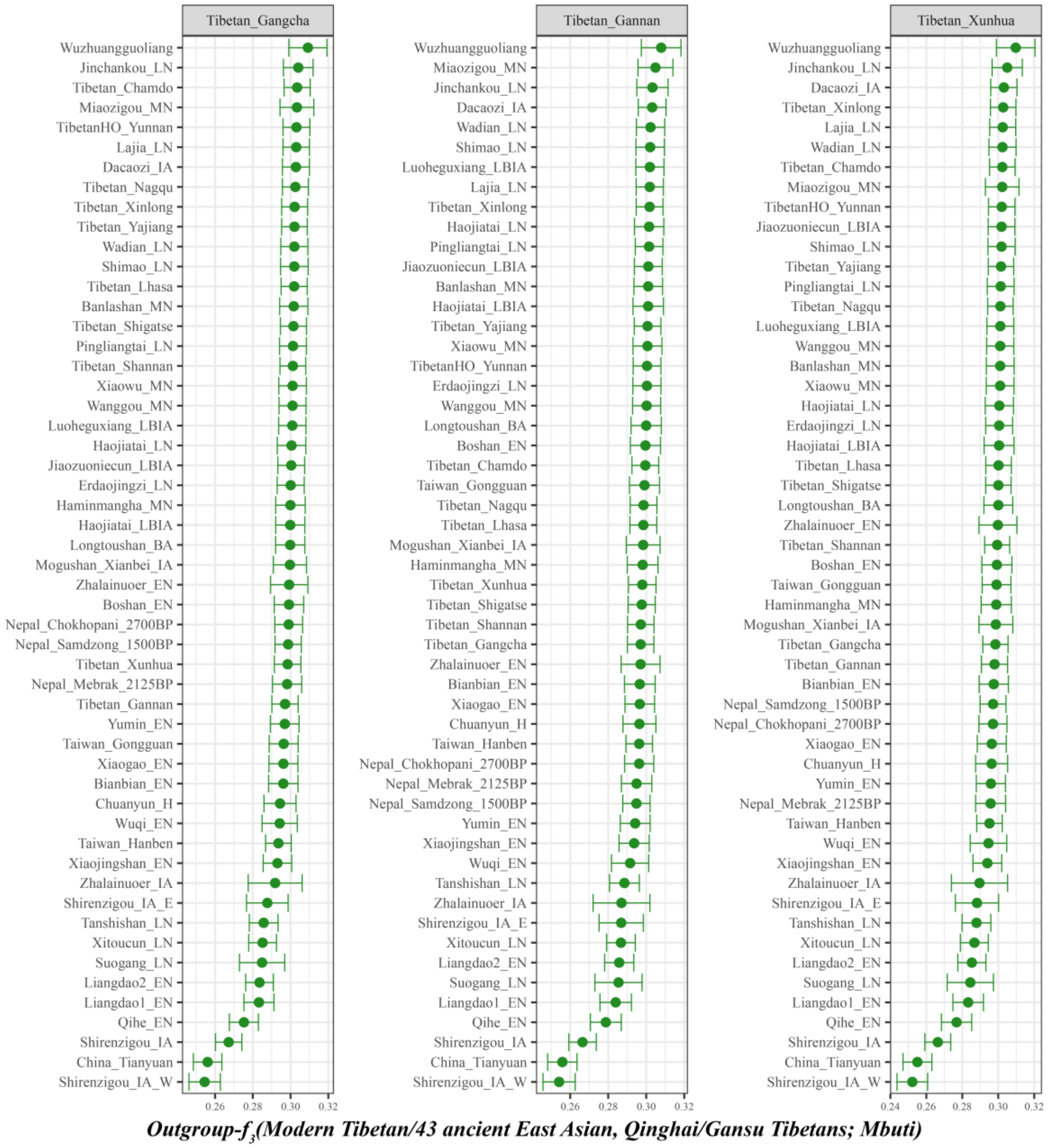

**Figure S8. The shared genetic drift between our studied Gansu/Qinghai Tibetans and other 43 spatial-temporally different East Asian populations.**

All populations listed in the Y-axis were sorted according to the outgroup-$f_3$ values and the outgroup-$f_3$ values were marked in the X-axis. The larger outgroup-$f_3$ value between our targeted Tibetan and reference populations showed the closer genetic relationship and stronger genetic affinity among them, otherwise, the smaller outgroup-$f_3$ value denotes distant genetic relationships among them.

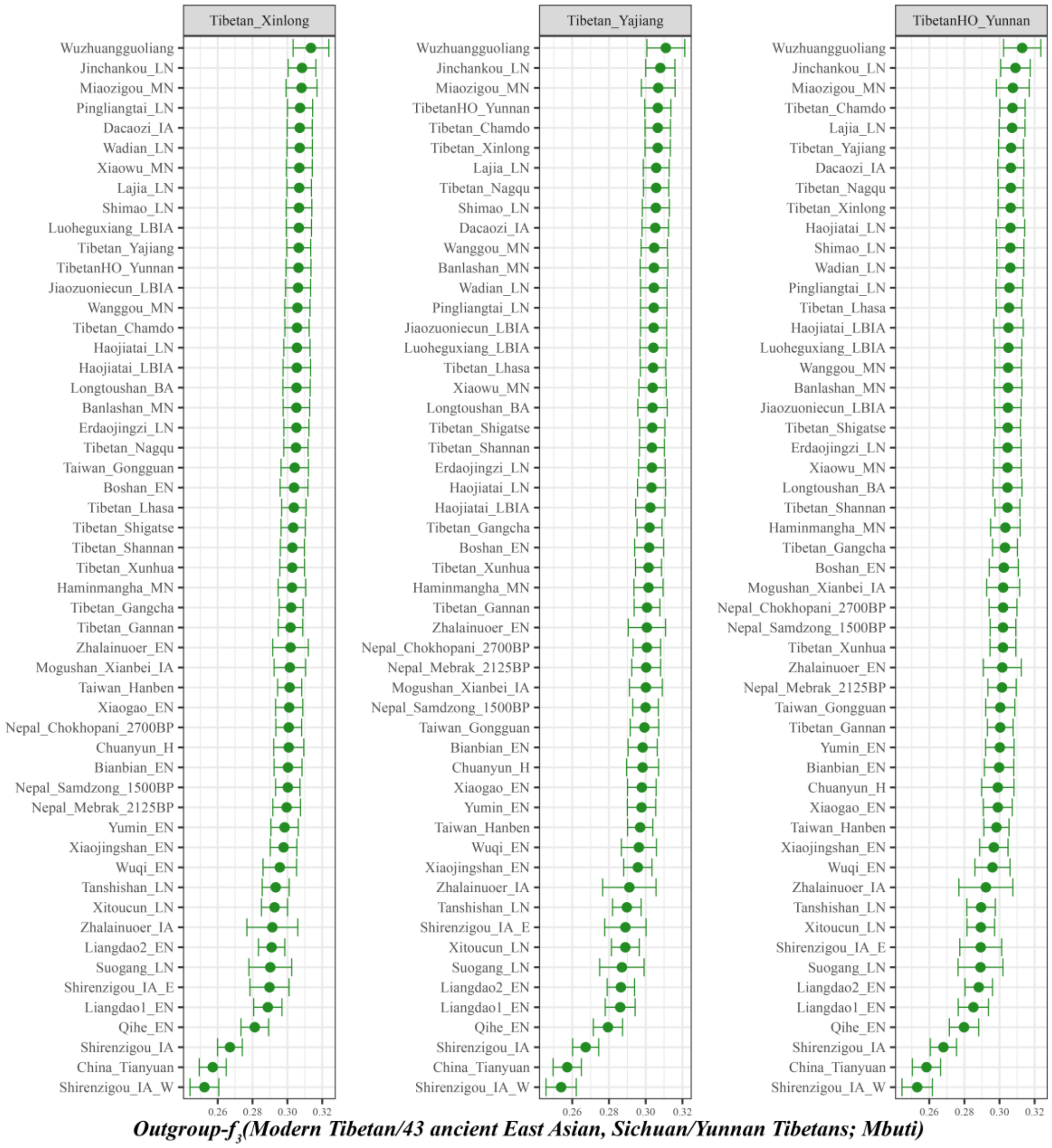

**Figure S9. The shared genetic drift between our studied Sichuan/Yunnan Tibetans and other 43 spatial-temporally different East Asian populations.**

All populations listed in the Y-axis were sorted according to the outgroup-$f_3$ values and the outgroup-$f_3$ values were marked in the X-axis. The larger outgroup-$f_3$ value between our targeted Tibetan and reference populations showed the closer genetic relationship and stronger genetic affinity among them, otherwise, the smaller outgroup-$f_3$ value denotes distant genetic relationships among them.

### Intra population differentiation among high-altitude residing and low-altitude residing Tibetans

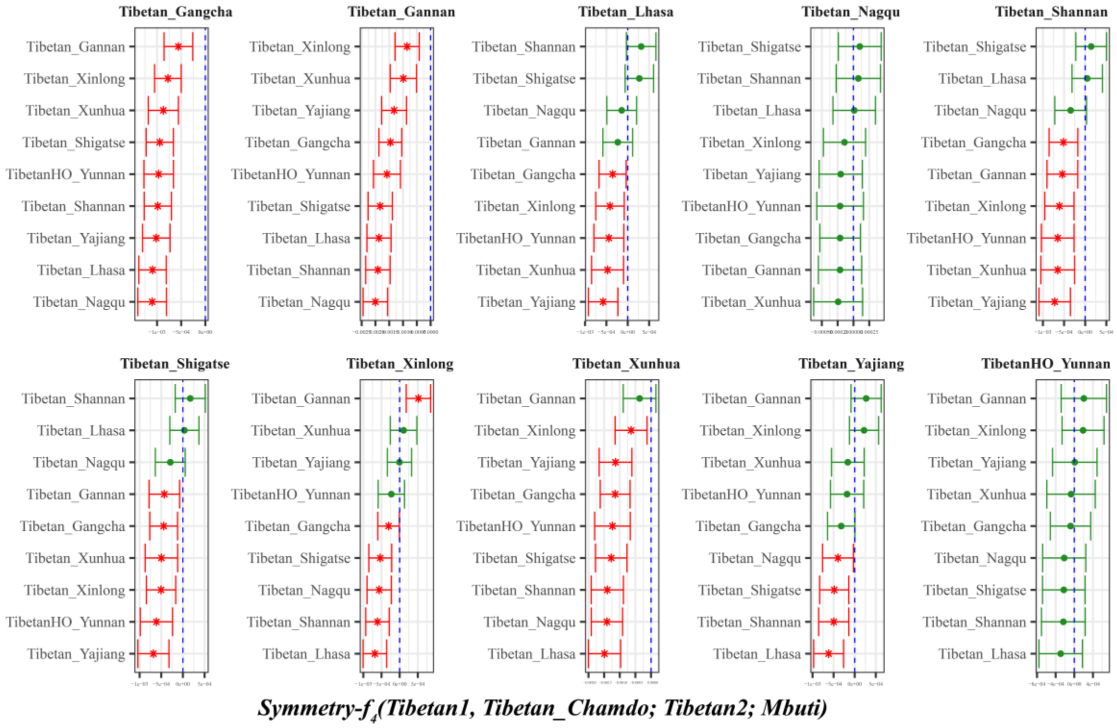

**Figure S10. Genomic similarities and differences among both high-altitude residing Tibetan and low-altitude residing Tibetans inferred from four population symmetry-$f_4$ statistics of the form *$f_4$(Tibetan1, Chamdo Tibetan; Tibetan2, Mbuti)*.**

Here, overlapping SNP loci included in the Affymetrix Human Origins platform among four analyzed populations were used. We used the genetic variation of Mbuti as the outgroup. Red asterisk point meant the significant value (Absolute value of Z-scores larger than three or equal to three) observed in the symmetry-$f_4$ statistics and green circle point denoted the non-significant $f_4$-statistic values (Absolute value of Z-scores less than three). All Tibetan2 were listed along the Y-axis and $f_4$ values were labeled along the X-axis. All tested population pairs were faceted or grouped via the Tibetan1. Significant negative $f_4$ values indicated that Tibetan2 shared more alleles with Chamdo Tibetan compared with Tibetan1 or compared with Tibetan1 Chamdo Tibetan harbored increased Tibetan2-related ancestry, and significant positive $f_4$ value indicated that Tibetan2 shared more derived alleles with Tibetan1 compared with Chamdo Tibetan, or elucidated as Chamdo Tibetan had increased Tibetan2-related ancestry relative to Tibetan1. The value of $f_4$-statistics-statistics equal to zero was marked as the blue dash line. The bar indicated three standard errors.

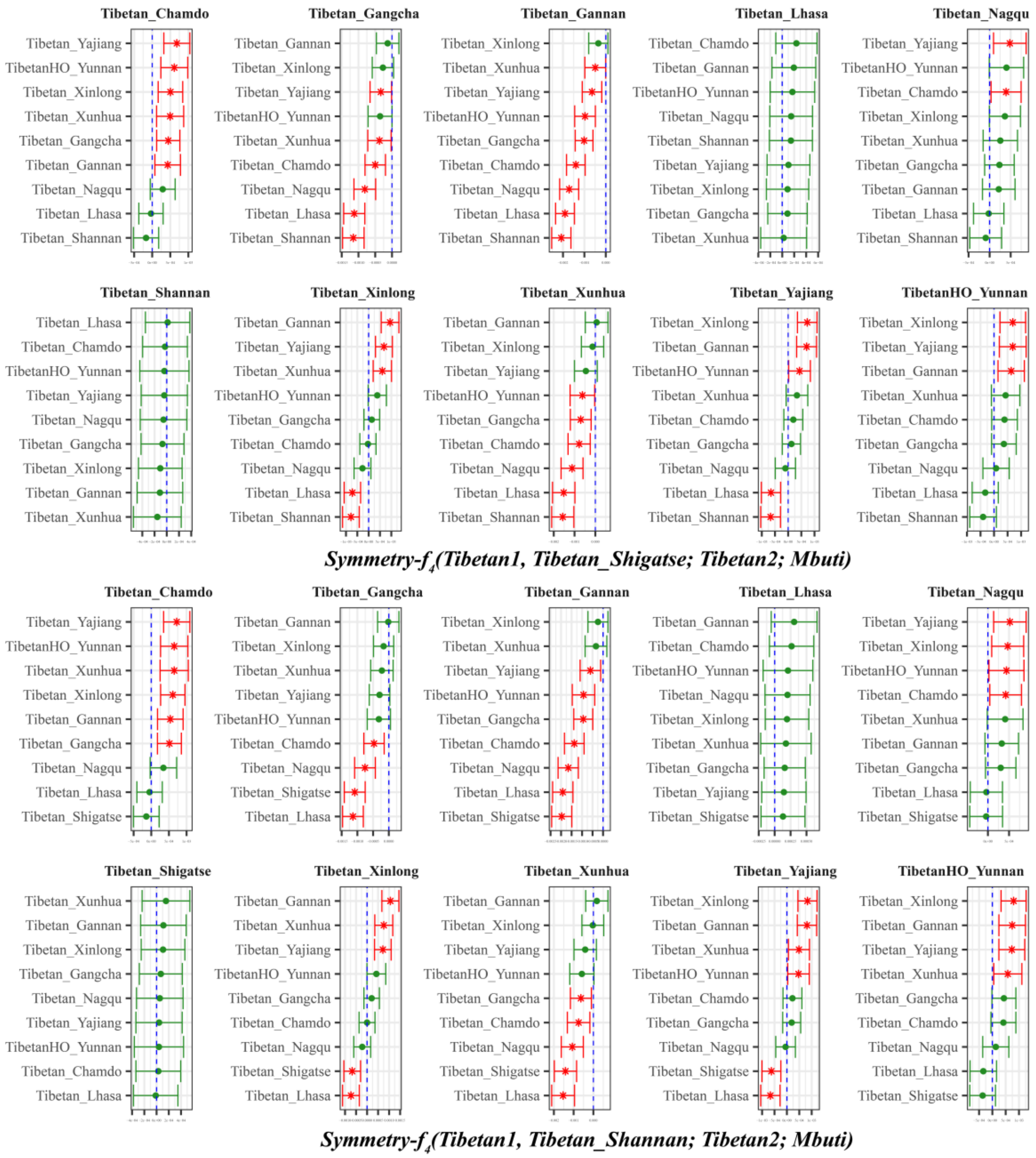

**Figure S11. Genomic similarities and differences among both high-altitude residing Tibetan and low-altitude residing Tibetans inferred from four population symmetry-$f_4$ statistics of the form *$f_4$(Tibetan1, Shigatse/Shannan Tibetan; Tibetan2, Mbuti)*.**

Here, overlapping SNP loci included in the Affymetrix Human Origins platform among four analyzed populations were used. We used the genetic variation of Mbuti as the outgroup. Red asterisk point meant the significant value (Absolute value of Z-scores larger than three or equal to three) observed in the symmetry-$f_4$ statistics and green circle point denoted the non-significant $f_4$-statistic values (Absolute value of Z-scores less than three). All Tibetan2 were listed along the Y-axis and $f_4$ values were labeled along the X-axis. All tested population pairs were faceted or grouped via the Tibetan1. Significant negative $f_4$ values indicated that Tibetan2 shared more alleles with Shigatse/Shannan Tibetan compared with Tibetan1 or Shigatse/Shannan Tibetan harbored increased Tibetan2-related ancestry compared with Tibetan1, and significant positive $f_4$ value indicated that Tibetan2 shared more derived alleles with Tibetan1 compared with Shigatse/Shannan Tibetan, or elucidated as Shigatse/Shannan Tibetan had increased Tibetan2-related ancestry relative to Tibetan1. The value of $f_4$ -statistics equal to zero was marked as the blue dash line. The bar indicated three standard errors.

We found that Shigatse, Shannan and Lhasa Tibetans form a clade compared with other reference Tibetan populations, due to no significant $f_4$-values were observed. Only negative or significant negative $f_4$ values were observed in Tibetan1 as Gangcha Tibetan, Xunhua Tibetan and Gannan Tibetan, which indicated that all included Tibetan populations shared more alleles with Shigatse Tibetan relative to them. Most of the values were positive or significantly positive when Chamdo Tibetan and Nagqu Tibetan used as Tibetan1, which indicated all other Tibetans shared more alleles with them compared with Shigatse Tibetans. When compared with low-altitude Tibetan from Sichuan and Yunnan Province, an asymmetrical distribution of values of $f_4$-statistics observed indicated that Shigatse Tibetan had more Lhasa/Shannan Tibetan-like ancestry component.

Among *$f_4$(Tibetan1, Shannan Tibetan; Tibetan2, Mbuti)*, Shigatse, Lhasa and Shannan Tibetans formed one clade. All Tibetans shared more alleles with Shannan Tibetans relative to the Gangcha, Gannan, Xunhua Tibetans, but all shared fewer alleles with Chamdo and Nagqu Tibetans. When we used Xinlong, Yajiang and Yunnan Tibetan as the reference populations, high-altitude Tibetan shared more alleles with Shannan Tibetans, but low altitude-resided Tibetan shared more alleles with low and mediate altitude-resided Tibetans.

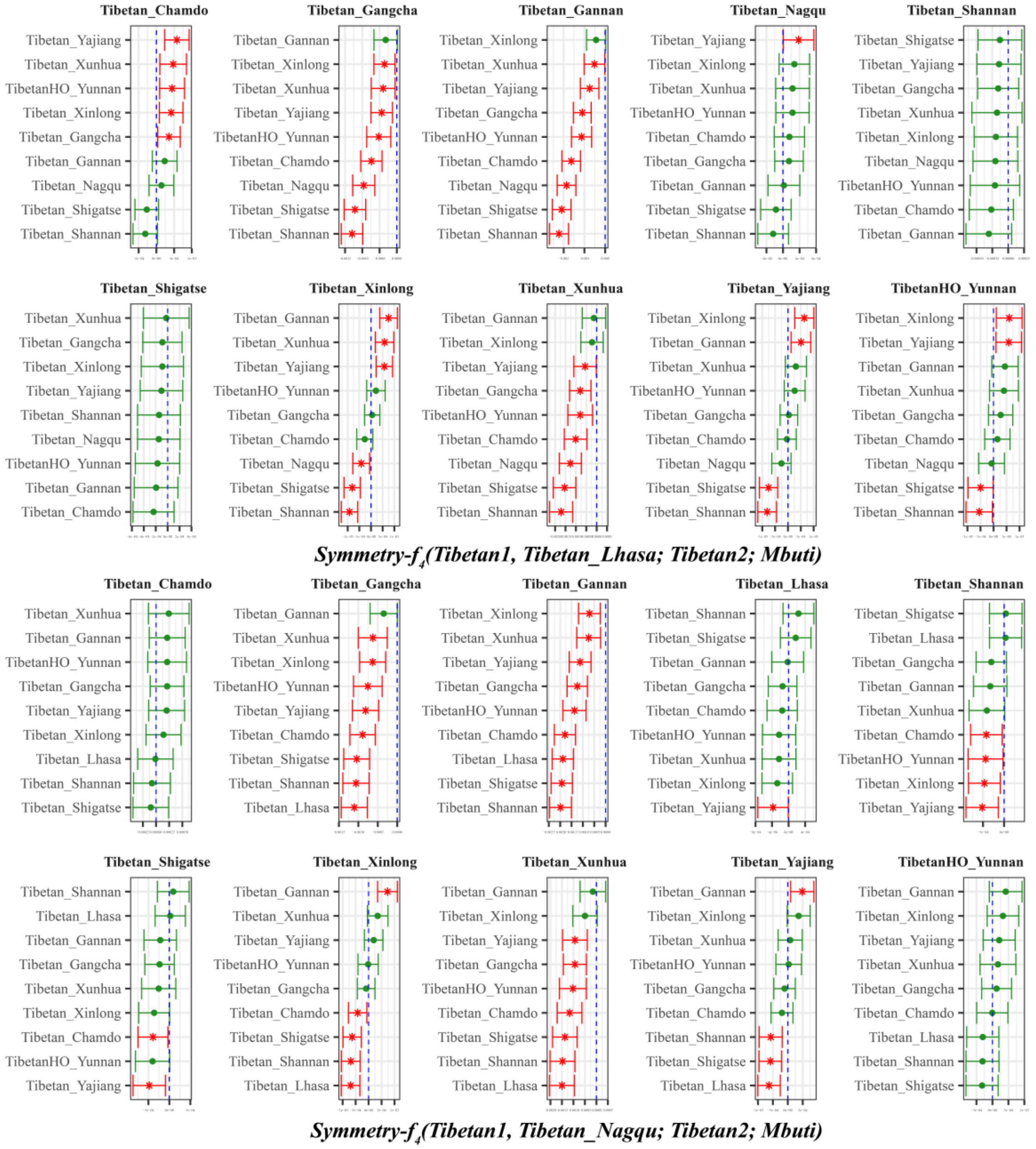

**Figure S12. Genomic similarities and differences among both high-altitude residing Tibetan and low-altitude residing Tibetans inferred from four population symmetry-$f_4$ statistics of the form *$f_4$(Tibetan1, Lhasa/Nagqu Tibetan; Tibetan2, Mbuti)*.**

Here, overlapping SNP loci included in the Affymetrix Human Origins platform among four analyzed populations were used. We used the genetic variation of Mbuti as the outgroup. Red asterisk point meant the significant value (Absolute value of Z-scores larger than three or equal to three) observed in the symmetry-$f_4$ statistics and green circle point denoted the non-significant $f_4$-statistic values (Absolute

value of Z-scores less than three). All Tibetan2 were listed along the Y-axis and $f_4$ values were labeled along the X-axis. All tested population pairs were faceted or grouped via the Tibetan1. Significant negative $f_4$ values indicated that Tibetan2 shared more alleles with Lhasa/Nagqu Tibetan compared with Tibetan1 or Lhasa/Nagqu Tibetan harbored increased Tibetan2-related ancestry compared with Tibetan1, and significant positive $f_4$ value indicated that Tibetan2 shared more derived alleles with Tibetan1 compared with Lhasa/Nagqu Tibetan, or elucidated as Lhasa/Nagqu Tibetan had increased Tibetan2-related ancestry relative to Tibetan1. The value of $f_4$-statistics equal to zero was marked as the blue dash line. The bar indicated three standard errors.

Lhasa, Shannan and Shigatse Tibetans formed one clade due to non-significant values observed. Compared with Gangcha, Gannan and Xunhua Tibetans, all negative or significant negative values suggested that all included Tibetans shared more alleles with Lhasa Tibetan. Non-consistent distributions of $f_4$-statistic values when we used Chamdo, Nagqu, Xinlong, Yajiang and Yunnan Tibetans as the reference populations of Tibetan1. Nagqu Tibetan and Lhasa Tibetan formed one clade except for Yajiang Tibetan which had more alleles with Nagqu Tibetan. Compared with Lhasa Tibetan, Chamdo Tibetan, Xinlong Tibetan, Yajiang and Yunnan Tibetan shared more genetic affinity with low-altitude Tibetan residents, Shigatse and Shannan Tibetans shared more alleles with Lhasa Tibetan.

Focused on Nagqu Tibetan from north of Tibet Province, we found that Chamdo Tibetan, Lhasa Tibetan (except for Yajiang (significantly negative)), Yunnan and Nagqu Tibetan formed one clade compared with other included reference Tibetans (no significantly negative or positive values). All included Tibetans shared more alleles with Nagqu Tibetans relative to Gangcha, Gannan, Shannan, Shigatse and Xunhua Tibetans due to negative or significant negative values were observed. Gannan Tibetan shared more alleles with Yajiang Tibetan and Xinlong Tibetans relative to Nagqu Tibetan.

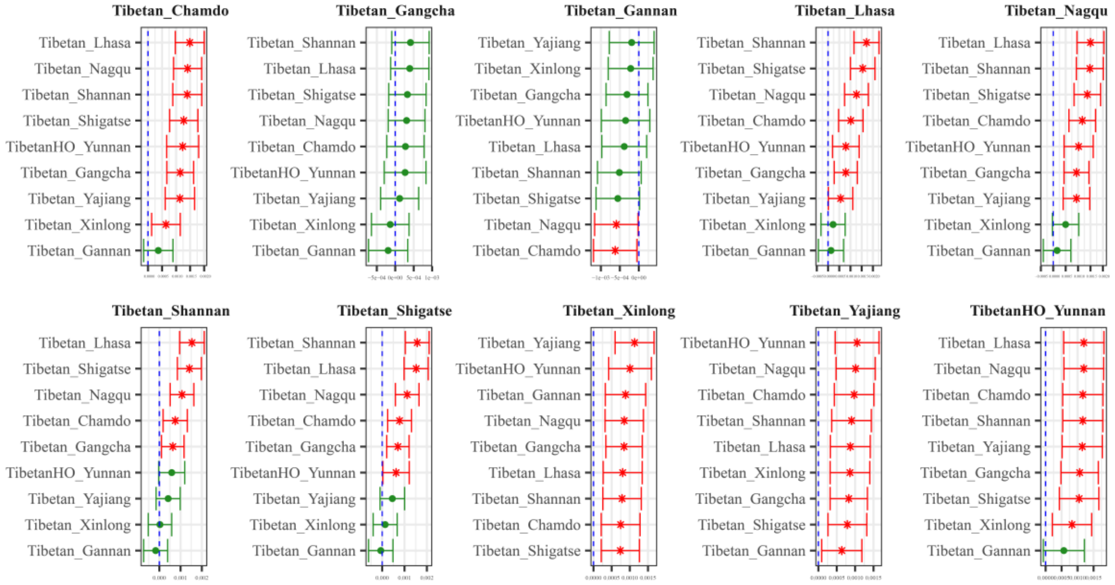

***Symmetry-$f_4$(Tibetan1, Tibetan_Xunhua; Tibetan2; Mbuti)***

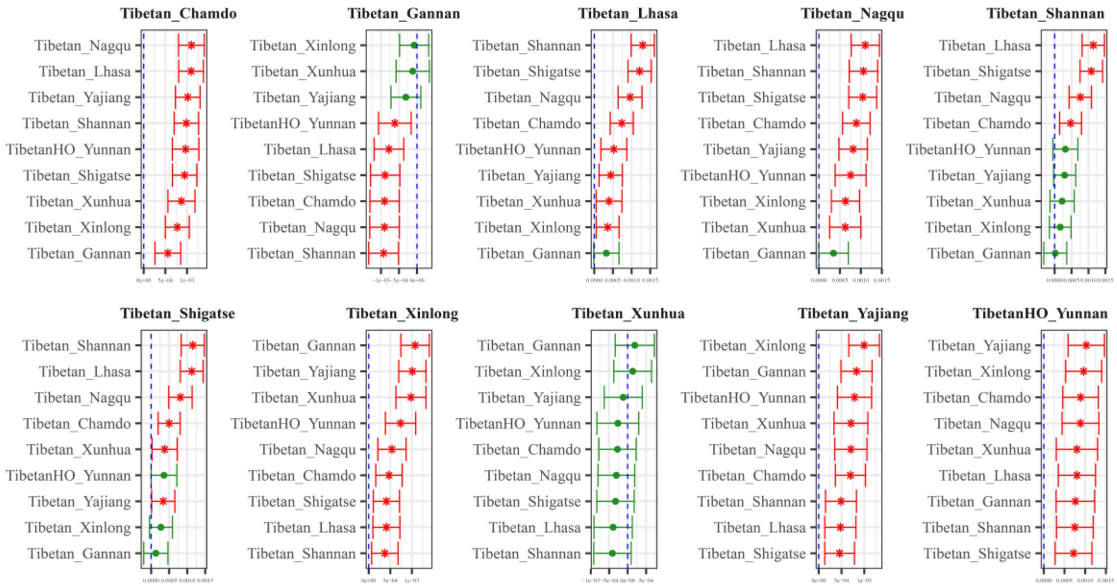

***Symmetry-$f_4$(Tibetan1, Tibetan_Gangcha; Tibetan2; Mbuti)***

**Figure S13. Genomic similarities and differences among both high-altitude residing Tibetan and low-altitude residing Tibetans inferred from four population symmetry-$f_4$ statistics of the form *$f_4$(Tibetan1, Gangcha/Xunhua Tibetan; Tibetan2, Mbuti)*.**

Here, overlapping SNP loci included in the Affymetrix Human Origins platform among four analyzed populations were used. We used the genetic variation of Mbuti as the outgroup. Red asterisk point meant the significant value (Absolute value of Z-scores larger than three or equal to three) observed in the symmetry-$f_4$ statistics and green circle point denoted the non-significant $f_4$-statistic values (Absolute value of Z-scores less than three). All Tibetan2 were listed along the Y-axis and $f_4$ values were labeled along the X-axis. All tested population pairs were faceted or grouped via the Tibetan1. Significant negative $f_4$ values indicated that Tibetan2 shared more alleles with Gangcha/Xunhua Tibetan compared with Tibetan1 or Gangcha/Xunhua Tibetan harbored increased Tibetan2-related ancestry compared with Tibetan1, and significant positive $f_4$ value indicated that Tibetan2 shared more derived alleles with Tibetan1 compared with Gangcha/Xunhua Tibetan, or elucidated as Gangcha/Xunhua Tibetan had increased Tibetan2-related ancestry relative to Tibetan1. The value of $f_4$-statistics equal to zero was marked as the blue dash line. The bar indicated three standard errors.
Gangcha Tibetan formed one clade with Xunhua Tibetan, as non-significant positive or negative $f_4$ values were observed. Significant positive values observed when Chamdo, Lhasa, Nagqu, Shannan, Shigatse, Xinlong, Yajiang and Yunnan Tibetans were used as Tibetan1, which indicated all included reference

Tibetans shared more drifts with them compared with Xunhua Tibetan. Focused on the geographically close Gannan Tibetan, all included Tibetans shared more alleles with Xunhua Tibetan, especially for Nagqqu Tibetan and Chamdo Tibetans (Significant negative values).

Compared with Gangcha Tibetan, it formed a clade with Xunhua Tibetan, as non-significant negative or positive values were observed. Similar patterns of genetic relationship with *$f_4$(Tibetan1, Xunhua Tibetan; Tibetan2, Mbuti)* were identified. Gangcha Tibetan owed fewer ancestry components related to other Tibetans with the exception of Gannan Tibetan. Compared with Gannan Tibetan, Gangcha Tibetan shared relatively increased ancestry related to other Tibetans.

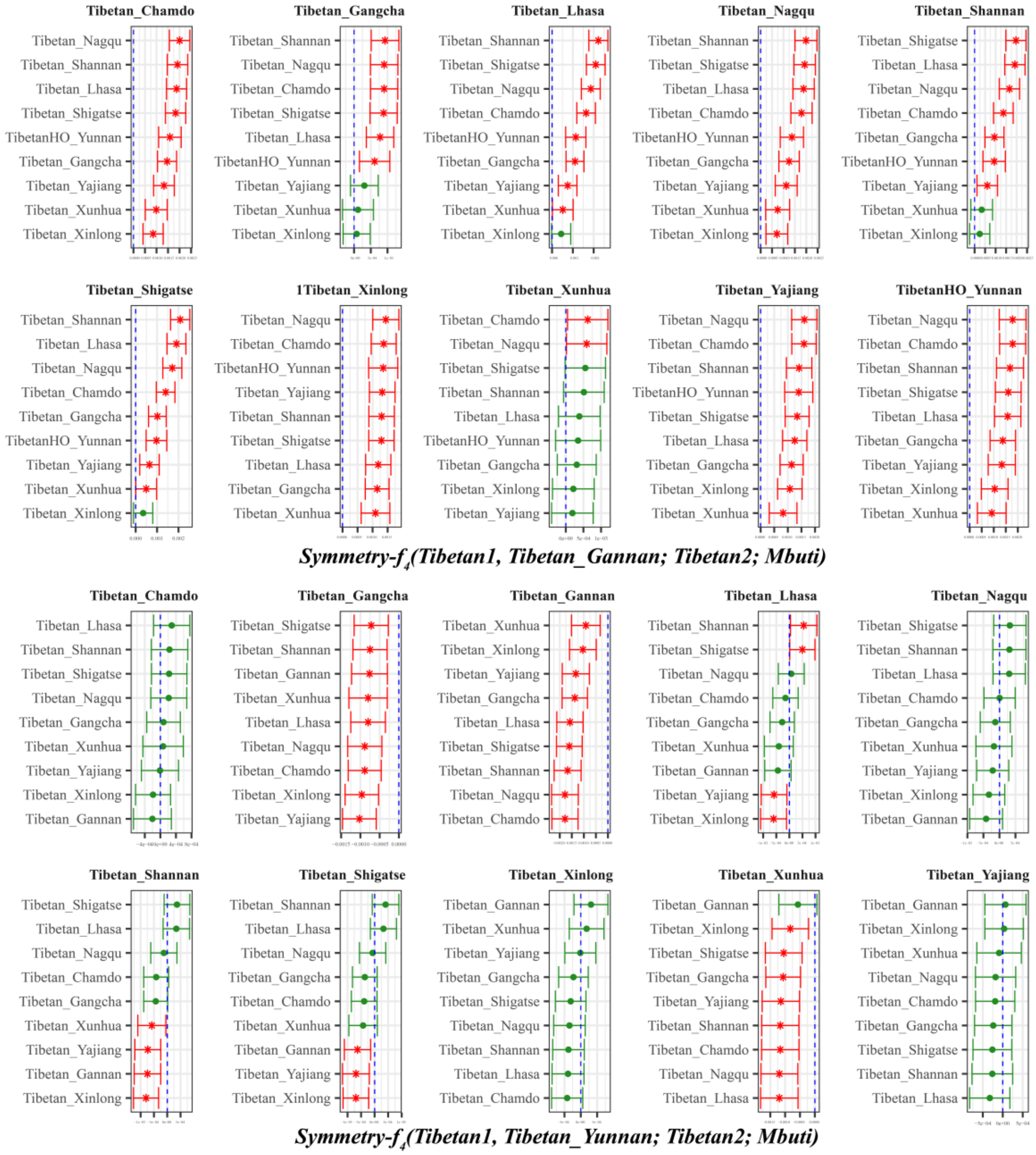

**Figure S14. Genomic similarities and differences among both high-altitude residing Tibetan and low-altitude residing Tibetans inferred from four population symmetry-$f_4$ statistics of the form *$f_4$(Tibetan1, Gannan/Yunnan Tibetan; Tibetan2, Mbuti)*.**

Here, overlapping SNP loci included in the Affymetrix Human Origins platform among four analyzed populations were used. We used the genetic variation of Mbuti as the outgroup. Red asterisk point meant the significant value (Absolute value of Z-scores larger than three or equal to three) observed in the symmetry-$f_4$ statistics and green circle point denoted the non-significant $f_4$-statistic values (Absolute value of Z-scores less than three). All Tibetan2 were listed along the Y-axis and $f_4$ values were labeled along the X-axis. All tested population pairs were faceted or grouped via the Tibetan1. Significant negative $f_4$ values indicated that Tibetan2 shared more alleles with Gannan/Yunnan Tibetan compared with Tibetan1 or Gannan/Yunnan Tibetan harbored increased Tibetan2-related ancestry compared with Tibetan1, and significant positive $f_4$ value indicated that Tibetan2 shared more derived alleles with Tibetan1 compared with Gannan/Yunnan Tibetan, or elucidated as Gannan/Yunnan Tibetan had increased Tibetan2-related ancestry relative to Tibetan1. The value of $f_4$-statistics equal to zero was marked as the blue dash line. The bar indicated three standard errors.

In $f_4$*(Tibetan1, Gannan; Tibetan2, Mbuti)*, all values observed were positive or significative positive, which indicated all included reference Tibetans shared more alleles with source populations related to other Tibetans than with Gannan Tibetan.

Compared with Yunnan Tibetan in the form $f_4$*(Tibetan1, Yunnan Tibetan; Tibetan2, Mbuti),* Chamdo, Nagqu, Xinlong, Yajiang and Yunnan Tibetans formed one clade relative to other included reference populations. All included Tibetans shared more alleles with Yunnan Tibetan than with Gangcha Tibetan, Gannan Tibetan and Xunhua Tibetan (Tebtan1 of Gannan is negative, not significant). Compared with Lhasa Tibetan, Yunnan Tibetan had increased ancestry related to Yajiang and Xinlong Tibetans and decreased ancestry related to Shannan Tibetan and Shigatse Tibetan. Compared with Shannan Tibetan, Yunnan Tibetan owed more ancestry components related to Xunhua Tibetan, Yajiang Tibetan, Gannan Tibetan and Xinlong Tibetan. Compared with Shigatse Tibetan, Yunnan Tibetan harbored increased ancestry components related to Gannan Tibetan, Yajiang Tibetan and Xinlong Tibetan.

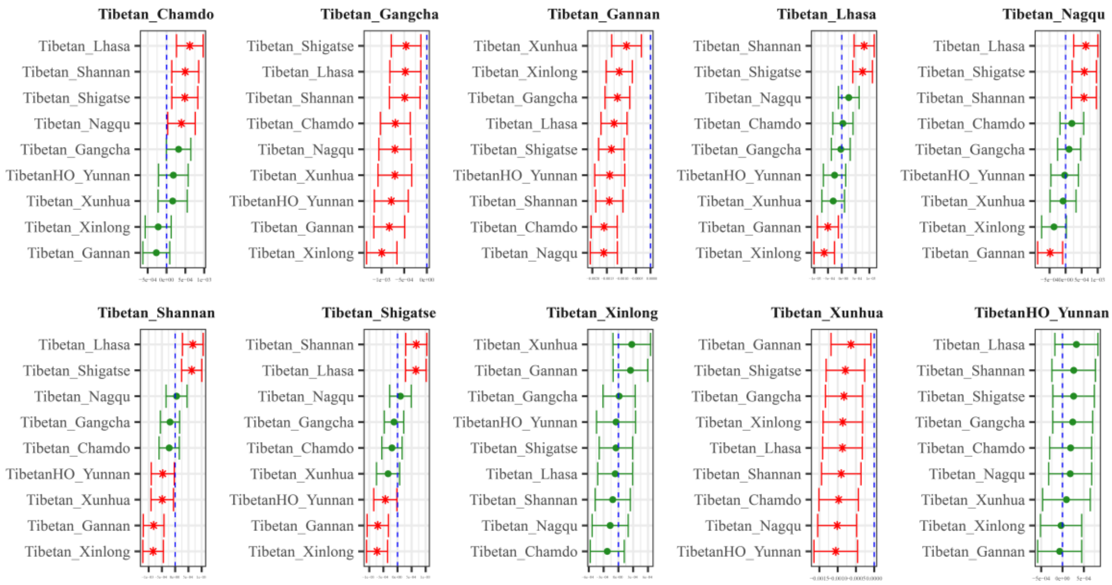

***Symmetry-$f_4$(Tibetan1, Tibetan_Yajiang; Tibetan2; Mbuti)***

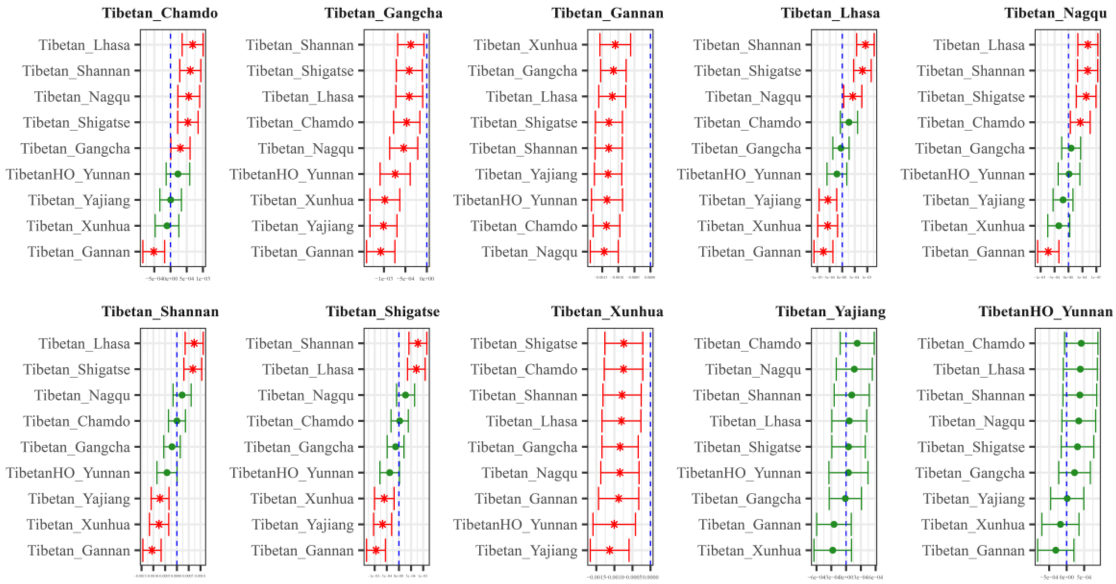

***Symmetry-$f_4$(Tibetan1, Tibetan_Xinlong; Tibetan2; Mbuti)***

**Figure S15. Genomic similarities and differences among both high-altitude residing Tibetan and low-altitude residing Tibetans inferred from four population symmetry-*$f_4$* statistics of the form *$f_4$(Tibetan1, Yajiang/Xinlong Tibetan; Tibetan2, Mbuti)*.**

Here, overlapping SNP loci included in the Affymetrix Human Origins platform among four analyzed populations were used. We used the genetic variation of Mbuti as the outgroup. Red asterisk point meant the significant value (Absolute value of Z-scores larger than three or equal to three) observed in the symmetry-$f_4$ statistics and green circle point denoted the non-significant $f_4$-statistic values (Absolute value of Z-scores less than three). All Tibetan2 were listed along the Y-axis and $f_4$ values were labeled along the X-axis. All tested population pairs were faceted or grouped via the Tibetan1. Significant negative $f_4$ values indicated that Tibetan2 shared more alleles with Yajiang/Xinlong Tibetan compared with Tibetan1 or Yajiang/Xinlong Tibetan harbored increased Tibetan2-related ancestry compared with Tibetan1, and significant positive $f_4$ value indicated that Tibetan2 shared more derived alleles with Tibetan1 compared with Yajiang/Xinlong Tibetan, or elucidated as Yajiang/Xinlong Tibetan had increased Tibetan2-related ancestry relative to Tibetan1. The value of $f_4$ equal to zero was marked as the blue dash line.

In the test of *f4(Tibetan1, Yajiang Tibetan; Tibetan2, Mbuti)*, we found that Tibetans from Sichuan Province (Xinlong and Yajiang Tibetans) and Yunnan Province (Yunnan Tibetan) formed one clade compared with all included Tibetans, as no significant positive or negative values were observed. All $f_4$-statistic values were less than three, which indicated that all included Tibetans shared more alleles with Yajiang Tibetan compared with Gangcha Tibetan, Xunhua Tibetan and Gannan Tibetan. Compared with other high-altitude resided Tibetans from Tibet Province, high-altitude and low-altitude Tibetans each shared with a distinct ancestry associated with their location in high-altitude and low-altitude Tibetan plateau.

In the test of *f4(Tibetan1, Yajiang/Xinlong Tibetan; Tibetan2, Mbuti)*., Yunnan Tibetan, Yajiang Tibetan and Xinlong Tibetan formed one clade related to all included reference Tibetan populations. All significant f4-statistic values observed in the *f4(Gangcha/Gannan/Xunhua Tibetans, Yajiang/Xinlong Tibetan; Tibetan2, Mbuti)* indicated that Xinlong Tibetan harbored more ancestry relative to other included Tibetans. Other tested results showed a geographical affinity associated with the genetic affinity of more shared derived alleles.

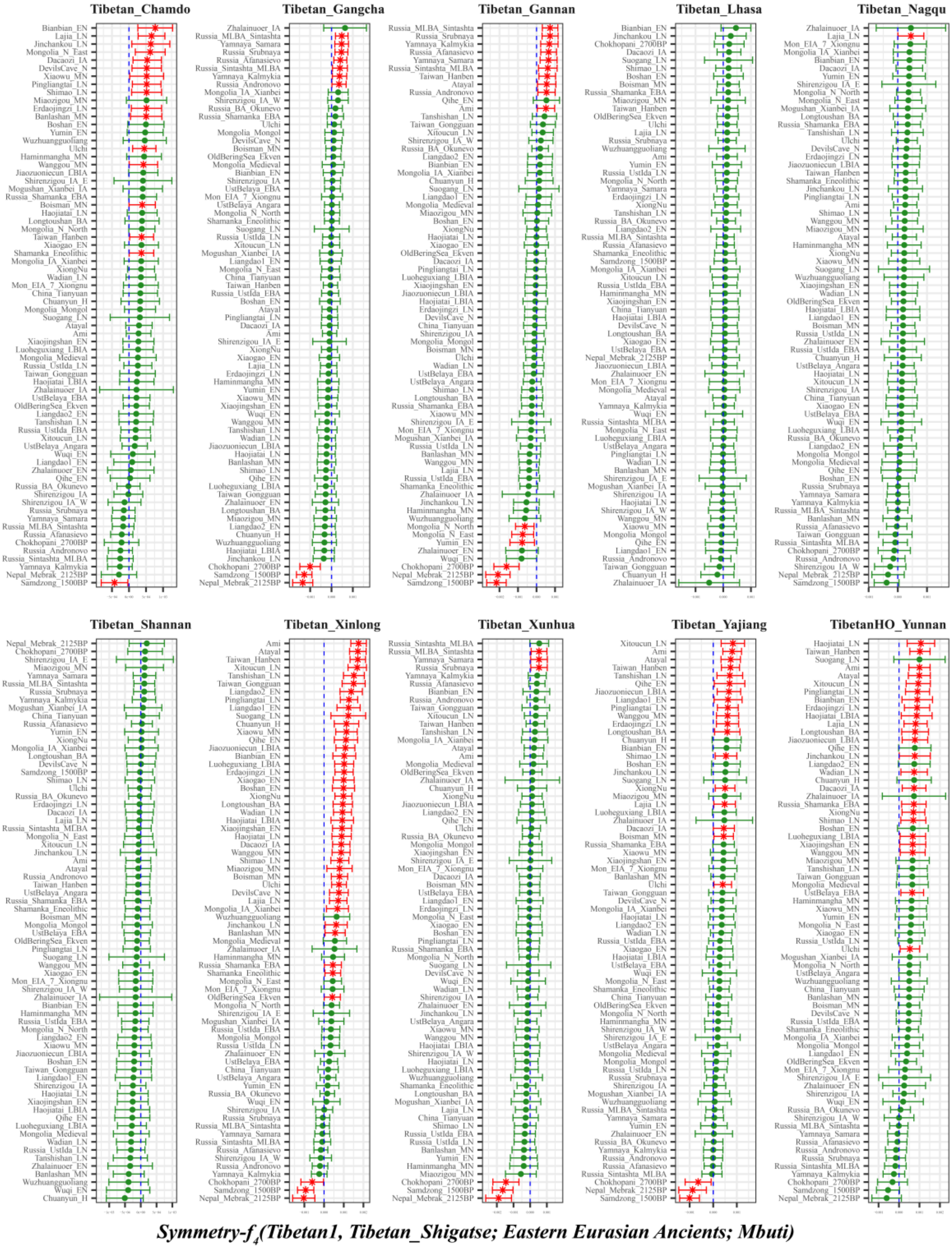

**Figure S16. Genomic affinity between modern Tibetans and eastern Eurasian ancient populations inferred from four population symmetry-$f_4$ statistics of the form $f_4$*(Tibetan1, Shigatse Tibetan; eastern Eurasian ancients, Mbuti)*.**

Here, overlapping SNP loci included in the Affymetrix Human Origins platform among four analyzed populations were used. We used the genetic variation of Mbuti as the outgroup. Red asterisk point meant the significant value (Absolute value of Z-scores larger than three or equal to three) observed in the symmetry-$f_4$ statistics and green circle point denoted the non-significant $f_4$-statistic values (Absolute value of Z-scores less than three). All Tibetan2 were listed along the Y-axis and $f_4$ values were labeled along the X-axis. All tested population pairs were faceted or grouped via the Tibetan1. Significant negative $f_4$ values indicated that the included eastern Eurasian ancient population shared more alleles with Shigatse Tibetan compared with Tibetan1 or Shigatse Tibetan harbored increased eastern Eurasian ancient population-related ancestry compared with Tibetan1, and significant positive $f_4$ value indicated that the included eastern Eurasian ancient population shared more derived alleles with Tibetan1 compared with Shigatse Tibetan, or elucidated as Shigatse Tibetan had increased eastern Eurasian ancient population-related ancestry relative to Tibetan1. The value of $f_4$-statistics equal to zero was marked as the blue dash line. The bar indicated three standard errors.

Compared with all included ancient populations from Yellow River, Yangtze River, West Liao River, Amur River, Russia, Tibet Plateau and others. No significant positive or negative $f_4$-statistic values were identified when we used Lhasa Tibetan, Nagqu Tibetan (except for Lajia_LN with significant $f_4$ value) and Shannan Tibetan as reference Tibetan1, which suggested that Shigatse Tibetan, Lhasa Tibetan, Nagqu Tibetan and Shannan Tibetan formed one clade. Compared with another Tibetan population from Chamdo city in Tibet Province, we found that Shigatse Tibetan harbored increased genetic drift related to the 1500-year-old Samdzong population.

When we used two populations from Qinghai province (Gangcha Tibetan and Xunhua Tibetan) as the reference populations of Tibetan1, $f_4$*(Gangcha/Xunhua Tibetans, Shigatse Tibetan; eastern Eurasian ancients, Mbuti)* showed Shigatse Tibetan harbored increased ancestry component related to ancient populations from Nepal (2700-year-old Chokhopani, 1500-year-old Samdzong and 2125-year-old Mebrak). Similar patterns of genomic affinity between Shigatse Tibetan with Nepal ancient populations were also observed in Tibetans from Sichuan Province of the form $f_4$*(Xinlong/Yajiang Tibetans, Shigatse Tibetan; eastern Eurasian ancients, Mbuti)*, however, compared with Yunnan Tibetan, this pattern of genetic affiliation disappeared, as no significant value was observed.

In the $f_4$*(Gannan Tibetan, Shigatse Tibetan; eastern Eurasian ancients, Mbuti)*, besides of the observed genomic affinity between Shigatse Tibetan and Nepal ancients, we also identified the genomic affinity between the inland Neolithic northern East Asian Yumin, Mongolia Neolithic populations (Mongolia_N_North and Mongolia_N_East).

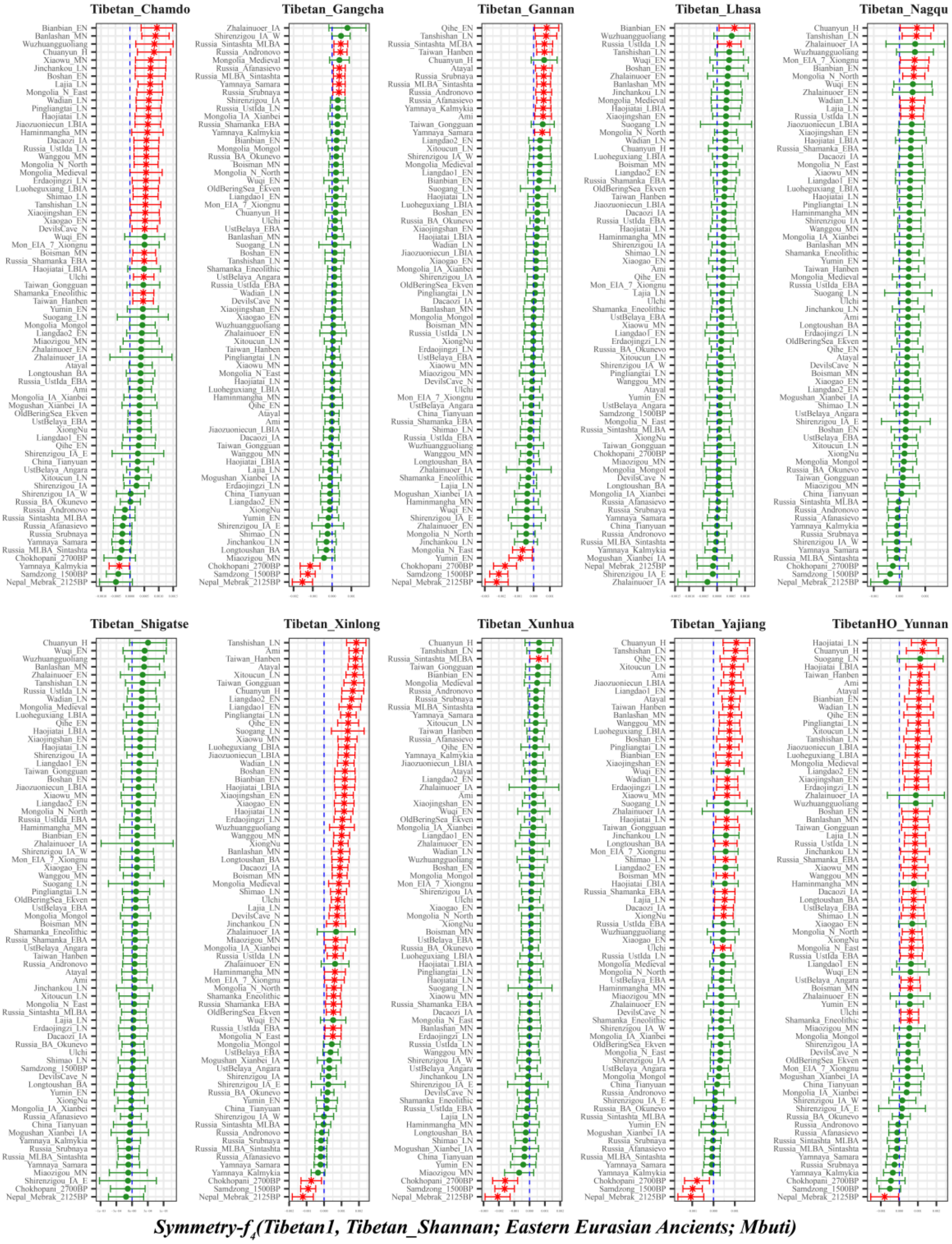

**Figure S17. Genomic affinity between modern Tibetans and eastern Eurasian ancient populations inferred from four population symmetry-$f_4$ statistics of the form $f_4$*(Tibetan1, Shannan Tibetan; eastern Eurasian ancients, Mbuti)*.**

Here, overlapping SNP loci included in the Affymetrix Human Origins platform among four analyzed populations were used. We used the genetic variation of Mbuti as the outgroup. Red asterisk point meant the significant value (Absolute value of Z-scores larger than three or equal to three) observed in the symmetry-$f_4$ statistics and green circle point denoted the non-significant $f_4$-statistic values (Absolute value of Z-scores less than three). All Tibetan2 were listed along the Y-axis and $f_4$ values were labeled along the X-axis. All tested population pairs were faceted or grouped via the Tibetan1. Significant negative $f_4$ values indicated that the included eastern Eurasian ancient population shared more alleles with Shannan Tibetan compared with Tibetan1 or Shannan Tibetan harbored increased eastern Eurasian ancient population-related ancestry compared with Tibetan1, and significant positive $f_4$ value indicated that the included eastern Eurasian ancient population shared more derived alleles with Tibetan1 compared with Shannan Tibetan, or elucidated as Shannan Tibetan had increased eastern Eurasian ancient population-related ancestry relative to Tibetan1. The value of $f_4$-statistics equal to zero was marked as the blue dash line. The bar indicated three standard errors.

No significant $f_4$-statistic values were observed in *$f_4$(Shigatse Tibetan, Shannan Tibetan; eastern Eurasian ancients, Mbuti)* suggested that Shigatse and Shannan Tibetans formed one clade related to all included ancient reference populations. Compared with Lhasa Tibetan and Nagqu Tibetan, only significant positive $f_4$-statistic values were observed in the tested form of *$f_4$(Lhasa Tibetan/Nagqu Tibetan, Shannan Tibetan; eastern Eurasian ancients, Mbuti)*, which indicated that Shannan Tibetan harbored fewer ancestry components related to coastal early Neolithic northern East Asian Bianbian and Late Neolithic Baikal ancient population relative to Lhasa Tibetan, and owned decreased alleles related to coastal southern East Asian from late Neolithic period (Tanshishan-LN) and historic period (Chuanyuan_H), coastal early Neolithic northern East Asian Bianbian and inland Late Neolithic northern East Asian Wadian from middle and lower Yellow River Basin, inland Late Neolithic northern East Asian Lajia from upper Yellow River, Mongolia Neolithic and historic ancients (Mongolia_N_North and Mongolia_EIA_7_Xiongnu) relative to Nagqu Tibetan. Compared with Chamdo Tibetan, relatively increased western Eurasian ancestry of early and late stepped ancestry (Kalmukia Yamnaya-related) was identified in Shannan Tibetan via *$f_4$(Chamdo Tibetan, Shannan Tibetan; Yamnaya_Kalmukia, Mbuti)* with significant negative Z-score. Compared with Tibetans from Sichuan and Qinghai Province, results of *$f_4$(Gangcha/Xinlong/Xunhua/Yajiang Tibetan, Shannan Tibetan; eastern Eurasian ancients of Nepal, Mbuti)* showed significant negative $f_4$-statistic values, which indicated that Shannan Tibetan had increased ancestry components related to Nepal ancients from three different culture backgrounds (Chokhopani, Samdzong and Mebrak). Focused on Yunnan Tibetan, only a significant negative value of $f_4$-statistic was observed when compared with the 2125-year-old Mebrak population, which showed more Mebrak-related ancestry existed in Shannan Tibetan compared with Yunnan Tibetan. Focused on Gannan Tibetan as the comparative subjects, we found Shannan Tibetan had increased ancestry related to both Nepal ancients, and Inner Mongolia (Yumin_EN) and Outer Mongolia (Mongolia_N_East) Neolithic people.

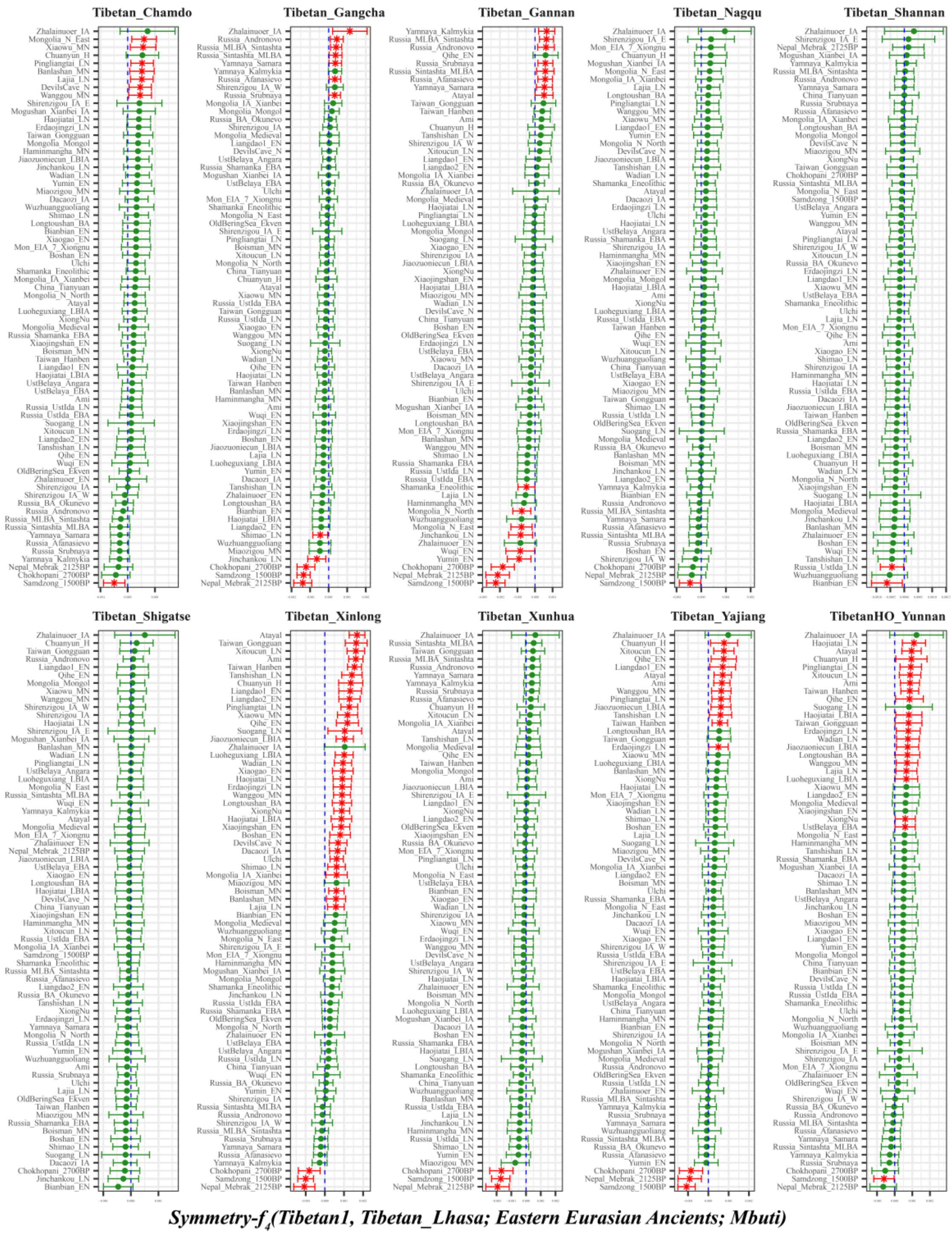

**Figure S18. Genomic affinity between modern Tibetans and eastern Eurasian ancient populations inferred from four population symmetry-$f_4$ statistics of the form *$f_4$(Tibetan1, Lhasa Tibetan; eastern Eurasian ancients, Mbuti)*.**

Here, overlapping SNP loci included in the Affymetrix Human Origins platform among four analyzed populations were used. We used the genetic variation of Mbuti as the outgroup. Red asterisk point meant the significant value (Absolute value of Z-scores larger than three or equal to three) observed in the symmetry-$f_4$ statistics and green circle point denoted the non-significant $f_4$-statistic values (Absolute value of Z-scores less than three). All Tibetan2 were listed along the Y-axis and $f_4$ values were labeled along the X-axis. All tested population pairs were faceted or grouped via the Tibetan1. Significant negative $f_4$ values indicated that the included eastern Eurasian ancient population shared more alleles with Lhasa Tibetan compared with Tibetan1 or Lhasa Tibetan harbored increased eastern Eurasian ancient population-related ancestry compared with Tibetan1, and significant positive $f_4$ value indicated that the included eastern Eurasian ancient population shared more derived alleles with Tibetan1 compared with Lhasa Tibetan or elucidated as Lhasa Tibetan had increased eastern Eurasian ancient population-related ancestry relative to Tibetan1. The value of $f_4$-statistics equal to zero was marked as the blue dash line. The bar indicated three standard errors.

No significant values observed in *$f_4$(Shigatse Tibetan, Lhasa Tibetan; eastern Eurasian ancients, Mbuti)* suggested Shigatse Tibetan and Shigatse Tibetan formed one clade. We also found Nagqu Tibetan and Shannan Tibetan formed one clade with Lhasa Tibetan, with the exception of Nagqu Tibetan with increased Samdzong-related ancestry and Shannan Tibetan with increased early Neolithic Bianbian or Late Neolithic UstIda related ancestry. Significant negative $f_4$-statistic values also observed of the form *$f_4$(Chamdo Tibetan, Lhasa Tibetan; eastern Eurasian ancients of Samdong_1500BP, Mbuti)*. Lhasa Tibetan owned increased Nepal ancient population-related ancestry compared middle and lower altitude Tibetan evidenced via *$f_4$(Xinlong/Yajiang/Xunhua Tibetans, Lhasa Tibetan; eastern Eurasian ancients of Nepal Ancients, Mbuti)* with significant negative $f_4$-statistics values. Compared with Lhasa Tibetan, Xinlong Tibetan and Yajiang Tibetan harbored increased lowland East Asian-related ancestry but lack for Xunhua Tibetan. Besides more Nepal ancients-related ancestry in Lhasa Tibetan compared with Gangcha Tibetan, it also owned more ancestry components related to the Late Neolithic Shimao ancients from Shaanxi Province and Jinchankou people from Qinghai Province. Compared with Gannan Tibetan, Lhasa harbored additional increased ancestry related to inland Early Neolithic northern East Asian of Wuqi from Amur River and Yumin from Inner Mongolia, as well as the ancient populations (Mongolia_N_East and Mongolia_N_North) from Mongolia Plateau in the areas of Outer Mongolia and Eneolithic Shamanka people from Baikal Lake region. Compared with Yunnan Tibetan, only increased Mebrak-related ancestry was observed in Lhasa Tibetan.

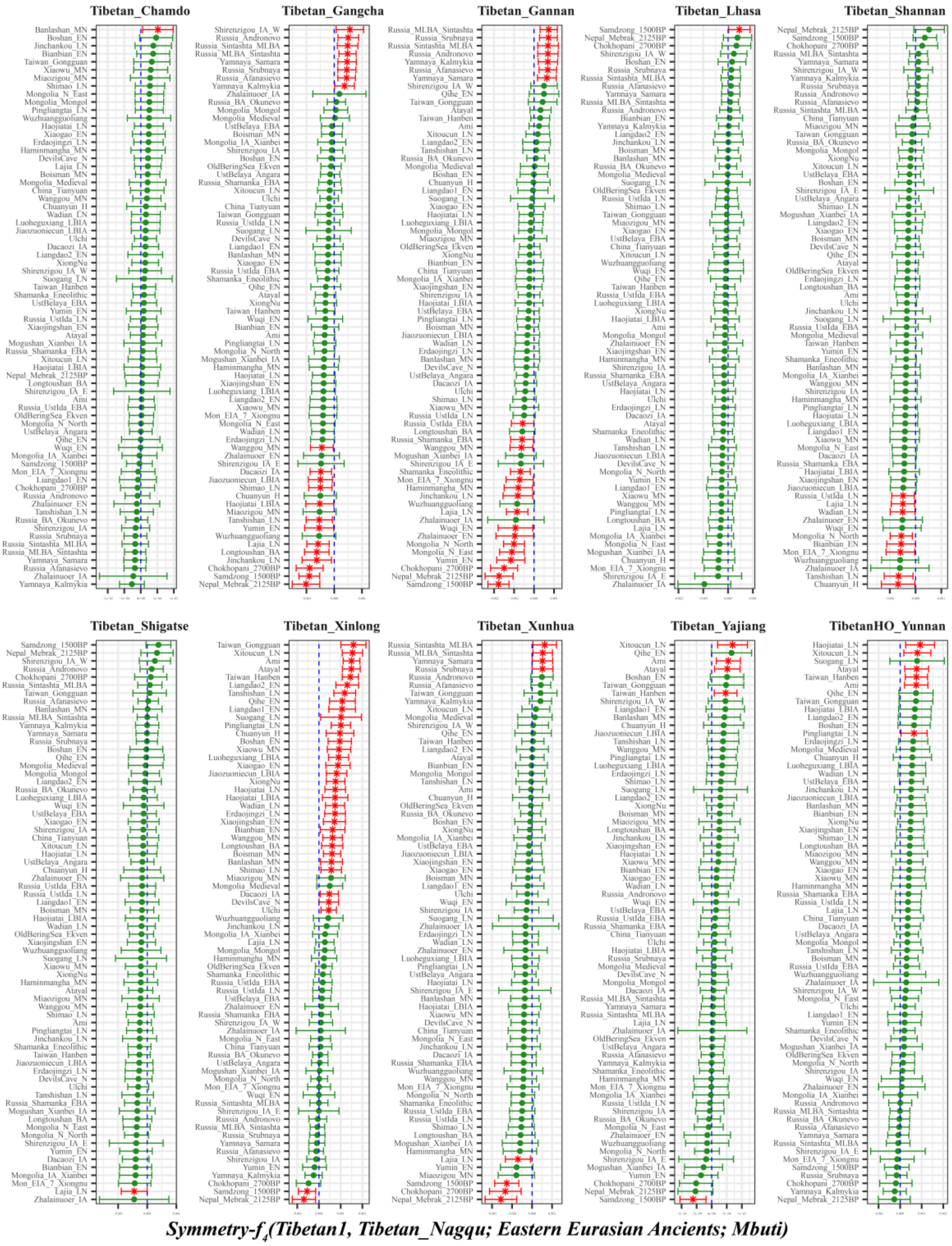

**Figure S19. Genomic affinity between modern Tibetans and eastern Eurasian ancient populations inferred from four population symmetry-$f_4$ statistics of the form $f_4$*(Tibetan1, Nagqu Tibetan; eastern Eurasian ancients, Mbuti)*.**

Here, overlapping SNP loci included in the Affymetrix Human Origins platform among four analyzed populations were used. We used the genetic variation of Mbuti as the outgroup. Red asterisk point meant the significant value (Absolute value of Z-scores larger than three or equal to three) observed in the symmetry-$f_4$ statistics and green circle point denoted the non-significant $f_4$-statistic values (Absolute value of Z-scores less than three). All Tibetan2 were listed along the Y-axis and $f_4$ values were labeled along the X-axis. All tested population pairs were faceted or grouped via the Tibetan1. Significant negative $f_4$ values indicated that the included eastern Eurasian ancient population shared more alleles with Nagqu Tibetan compared with Tibetan1 or Nagqu Tibetan harbored increased eastern Eurasian ancient population-related ancestry compared with Tibetan1, and significant positive $f_4$ value indicated that the included eastern Eurasian ancient population shared more derived alleles with Tibetan1 compared with Nagqu Tibetan or elucidated as Nagqu Tibetan had increased eastern Eurasian ancient population-related ancestry relative to Tibetan1. The value of $f_4$-statistics equal to zero was marked as the blue dash line. The bar indicated three standard errors.

Overall, Chamdo Tibetan, Lhasa Tibetan and Shigatse Tibetan formed one clade with Nagqu Tibetan with the exception of Chamdo Tibetan with increased Middle Neolithic Banlashan-related ancestry, Lhasa Tibetan with more Samdzong-related ancestry and Shigatse Tibetan with decreased Late Neolithic Lajia-related ancestry. Compared with Shannan Tibetan, Nagqu Tibetan harbored increased ancestry associated with lowland ancient populations from south East Asia of the southeast coastal region of southern China (Tanshishan_LN and Chuanyun_H) to as north far as Baikal Region (Russia_UstIda_LN). No increased ancestries related to the included eastern ancient populations were identified in Nagqu Tibetan compared with lowland Yunnan Tibetan, as no significant negative $f_4$-statistic value was observed in $f_4$*(Yunnan Tibetan, Nagqu Tibetan; eastern Eurasian ancients, Mbuti)*. Compared with Tibetans from Sichuan Province, Nagqu Tibetan owned increased Samdzong-related ancestry relative to Yajiang Tibetan and more Samdzong and Mebrak related ancestry relative to Xinlong Tibetan. Compared to Tibetans from Qinghai Province, Nagqu Tibetan owned both increased Nepal ancient related ancestry and increased Late Neolithic Lajia related ancestry relative to Xunhua Tibetan, and it harbored additionally increased ancestry related to coastal Late Neolithic southern East Asian of Tanshishan_LN, middle Yellow River Middle Neolithic to Iron Age ancient populations (Wanggou_MN, Haojiatai_LBIA and Jiaozuoniecun_LBIA), Upper Xiajiadian culture of Longtoushan_BA, inland Neolithic northern East Asian (Yumin_EN and Shimao_LN) and other upper Yellow River Late Neolithic Jinchankou and Iron Age Dacaozi. Here, we found a closer affinity between upper Yellow River ancient populations with Nagqu Tibetan, not with geographically close Gangcha Tibetan, which suggested that ancient populations from Lajia, Jinchankou and Dacaozi may the direct ancestors of modern Nagqu Tibetan. Compared with the Gannan Tibetan with a geographically close relationship with Gangcha Tibetan, Nagqu Tibetan harbored increased ancestry related to ancient populations from Russia (Russia_UstIda_EBA, Russia_Shamanka_EBA and Shamanka_Eneolithic), Mongolia (Mongolia_EIA_7_Xiongnu, Mongolia_N_North and Mongolia_N_East) and early Neolithic northeastern East Asian from Amur River (Zhalainuoer_EN and Wuqi_EN ) and middle Neolithic northeastern East Asian from western Liao River (Haminmangha_MN), which suggested a connection between Nagqu Tibetan and these Siberian-related ancient populations.

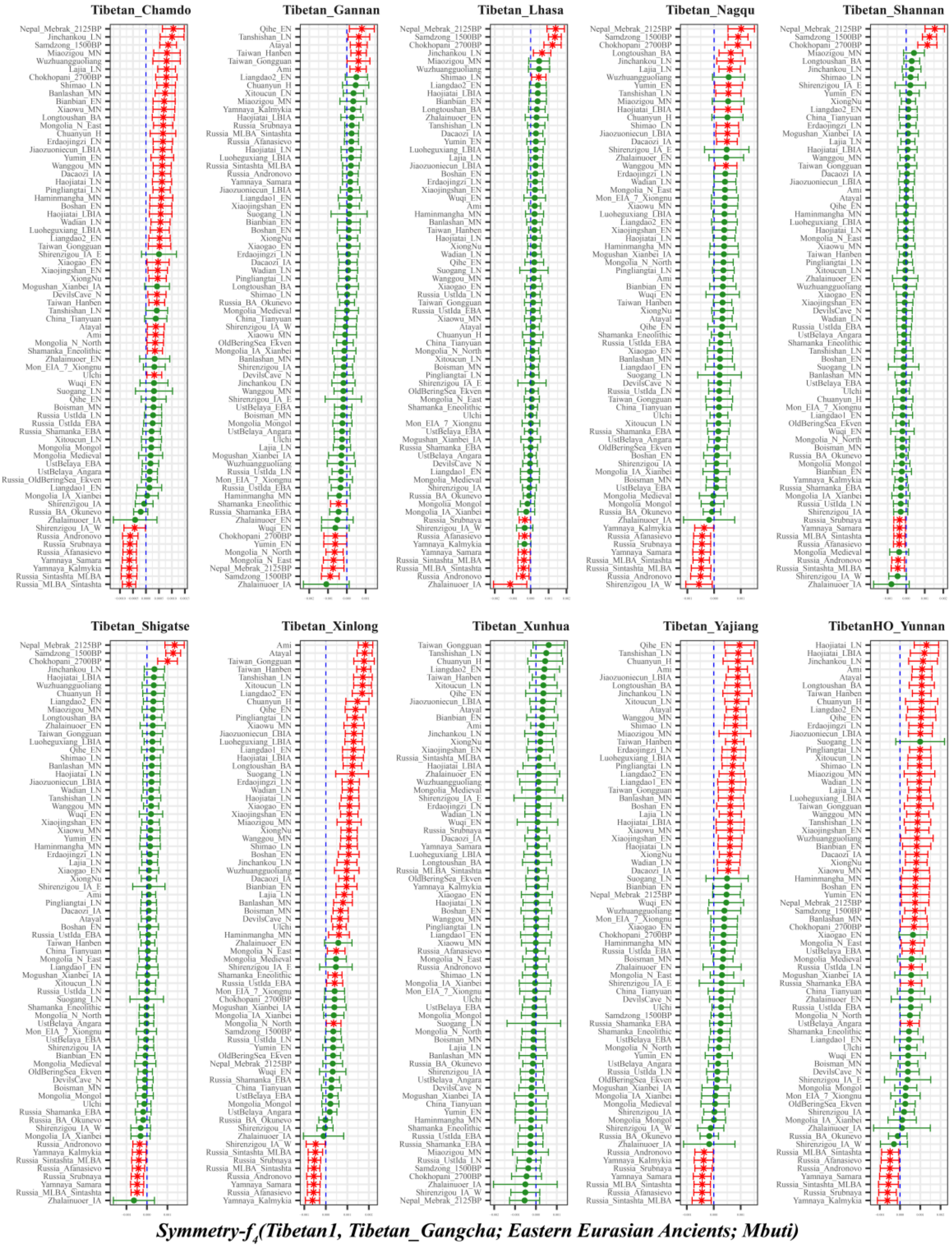

**Figure S20. Genomic affinity between modern Tibetans and eastern Eurasian ancient populations inferred from four population symmetry-$f_4$ statistics of the form $f_4$(*Tibetan1, Gangcha Tibetan; eastern Eurasian ancients, Mbuti*).**

Here, overlapping SNP loci included in the Affymetrix Human Origins platform among four analyzed populations were used. We used the genetic variation of Mbuti as the outgroup. Red asterisk point meant the significant value (Absolute value of Z-scores larger than three or equal to three) observed in the symmetry-$f_4$ statistics and green circle point denoted the non-significant $f_4$-statistic values (Absolute value of Z-scores less than three). All Tibetan2 were listed along the Y-axis and $f_4$ values were labeled along the X-axis. All tested population pairs were faceted or grouped via the Tibetan1. Significant negative $f_4$ values indicated that the included eastern Eurasian ancient population shared more alleles with Gangcha Tibetan compared with Tibetan1 or Gangcha Tibetan harbored increased eastern Eurasian ancient population-related ancestry compared with Tibetan1, and significant positive $f_4$ value indicated that the included eastern Eurasian ancient population shared more derived alleles with Tibetan1 compared with Gangcha Tibetan or elucidated as Gangcha Tibetan had increased eastern Eurasian ancient population-related ancestry relative to Tibetan1. The value of $f_4$-statistics equal to zero was marked as the blue dash line. The bar indicated three standard errors.

Xunhua Tibetan formed one clade with Gangcha Tibetan, as non-significant negative or positive $f_4$-statistics were observed. Compared to other Tibetans, Gangcha Tibetan harbored increased ancestry with Sintashta-related (Russia_MLBA_Sintashta and Russia_Sintashta_MLBA), Yamnaya-related (Yamnaya_Kalmykia and Yamnaya_Samara), and other ancestry related to middle and late Bronze Age Steppe related ancestry (Russia_Afanasievo, Russia_Srubnaya, Russia_Andronovo and Shirenzigou_IA_W).

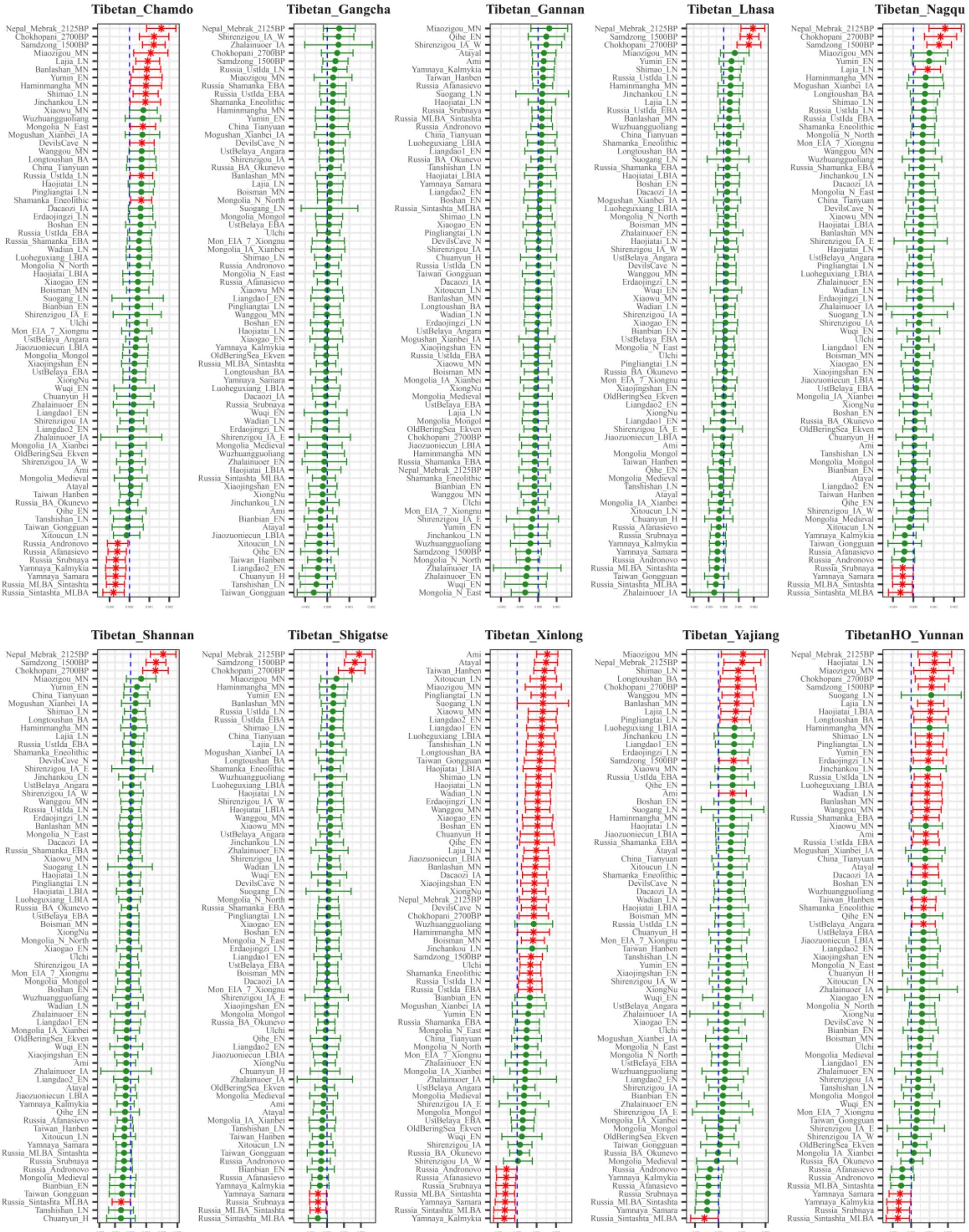

***Symmetry-$f_4$(Tibetan1, Tibetan_Xunhua; Eastern Eurasian Ancients; Mbuti)***

**Figure S21. Genomic affinity between modern Tibetans and eastern Eurasian ancient populations inferred from four population symmetry-$f_4$ statistics of the form $f_4$*(Tibetan1, Xunhua Tibetan; eastern Eurasian ancients, Mbuti)*.**
Here, overlapping SNP loci included in the Affymetrix Human Origins platform among four analyzed populations were used. We used the genetic variation of Mbuti as the outgroup. Red asterisk point meant the significant value (Absolute value of Z-scores larger than three or equal to three) observed in the symmetry-$f_4$ statistics and green circle point denoted the non-significant $f_4$-statistic values (Absolute value of Z-scores less than three). All Tibetan2 were listed along the Y-axis and $f_4$ values were labeled along the X-axis. All tested population pairs were faceted or grouped via the Tibetan1. Significant negative $f_4$ values indicated that the included eastern Eurasian ancient population shared more alleles with Xunhua Tibetan compared with Tibetan1 or Xunhua Tibetan harbored increased eastern Eurasian ancient population-related ancestry compared with Tibetan1, and significant positive $f_4$ value indicated that the included eastern Eurasian ancient population shared more derived alleles with Tibetan1 compared with Xunhua Tibetan or elucidated as Xunhua Tibetan had increased eastern Eurasian ancient population-related ancestry relative to Tibetan1. The value of $f_4$-statistics equal to zero was marked as the blue dash line. The bar indicated three standard errors.
Results of non-significant $f_4$-statistic of the form $f_4$*(Gangcha/Gannan Tibetans, Xunhua Tibetan; eastern Eurasian ancients, Mbuti)* showed that both Gangcha Tibetan and Gannan Tibetan formed one clade with Xunhua Tibetan. Compared with five Tibetans from Tibet Province harboring more Nepal ancient-related ancestry, significant signals of western Eurasian Steppe population affinity with Xunhua Tibetan were observed except for Lhasa Tibetan via $f_4$*(Chamdo/Lhasa/Nagqu/Shannan/Shigatse Tibetans, Xunhua Tibetan; eastern Eurasian ancients, Mbuti)*. Compared with lowland Tibetans, the Steppe-related population affinity with Xunhua Tibetan was also observed.

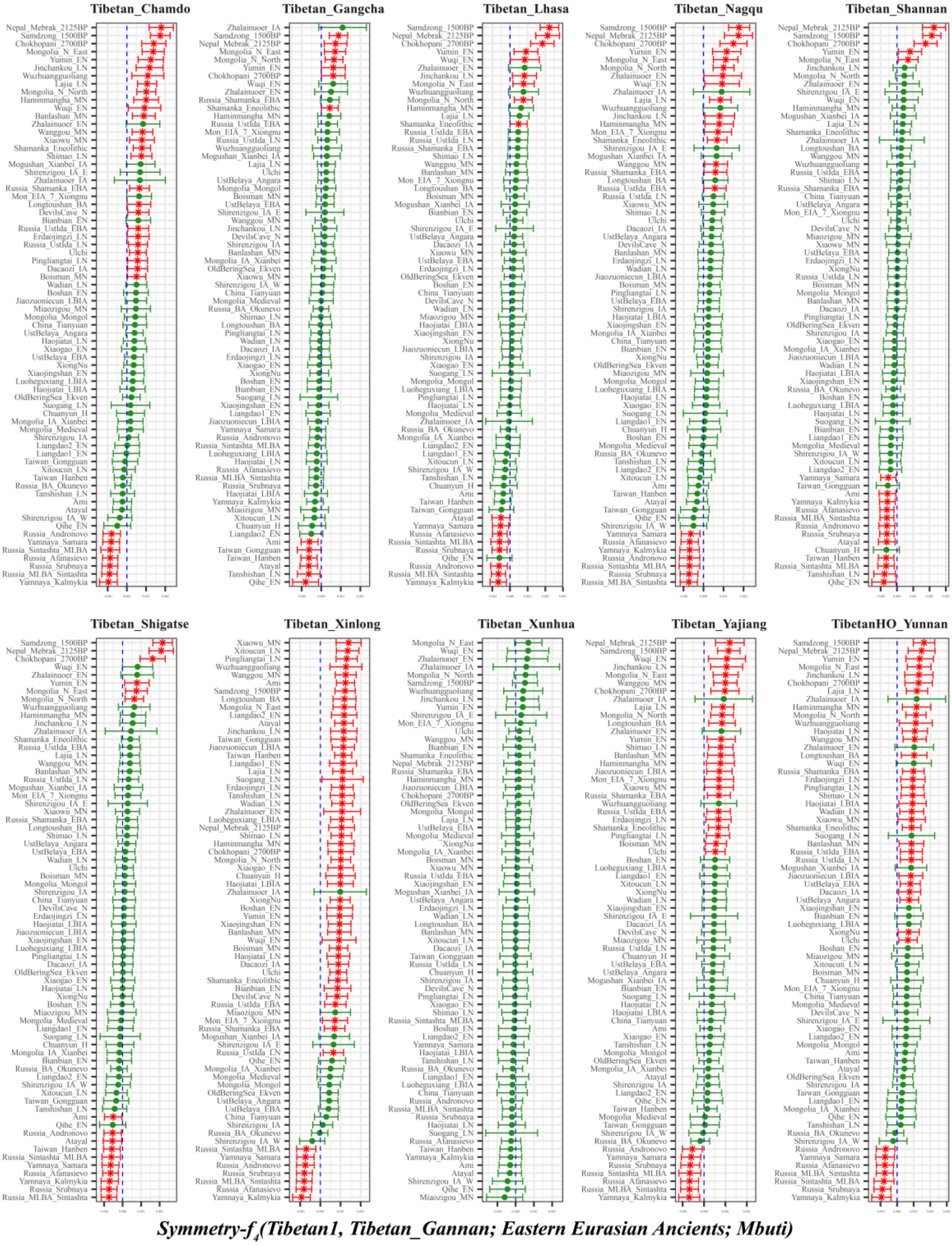

**Figure S22. Genomic affinity between modern Tibetans and eastern Eurasian ancient populations inferred from four population symmetry-$f_4$ statistics of the form $f_4$*(Tibetan1, Gannan Tibetan; eastern Eurasian ancients, Mbuti)*.**

Here, overlapping SNP loci included in the Affymetrix Human Origins platform among four analyzed populations were used. We used the genetic variation of Mbuti as the outgroup. Red asterisk point meant the significant value (Absolute value of Z-scores larger than three or equal to three) observed in the symmetry-$f_4$ statistics and green circle point denoted the non-significant $f_4$-statistic values (Absolute value of Z-scores less than three). All Tibetan2 were listed along the Y-axis and $f_4$ values were labeled along the X-axis. All tested population pairs were faceted or grouped via the Tibetan1. Significant negative $f_4$ values indicated that the included eastern Eurasian ancient population shared more alleles with Gannan Tibetan compared with Tibetan1 or Gannan Tibetan harbored increased eastern Eurasian ancient population-related ancestry compared with Tibetan1, and significant positive $f_4$ value indicated that the included eastern Eurasian ancient population shared more derived alleles with Tibetan1 compared with Gannan Tibetan or elucidated as Gannan Tibetan had increased eastern Eurasian ancient population-related ancestry relative to Tibetan1. The value of $f_4$-statistics equal to zero was marked as the blue dash line. The bar indicated three standard errors.

As non-significant signals were observed in *$f_4$(Xunhua Tibetan, Gannan Tibetan; eastern Eurasian ancients, Mbuti)*, Gannan Tibetan formed one clade with Xunhua Tibetan.

For five Tibetans from Tibet Province, Gannan Tibetan had increased ancestry related to early and middle Bronze Age Steppe Yamnaya ancestry (Yamnaya_Kalmykia, Yamnaya_Samara), and also owned more Sintashta-like ancestry (Russia_MLBA_Sintashta, Russia_Sintashta_MLBA) and Srubnaya-/Afanasievo-/Andronovo-like ancestry (Russia_Srubnaya, Russia_Afanasievo, Russia_Andronovo) relative to Shigatse Tibetan. We also found that Gannan Tibetan had increased ancestry related to coastal Neolithic to Modern southern East Asians (Taiwan_Hanben, Atayal and Qihe_EN) compared with Shigatse Tibetan.

Relative to Shannan Tibetan, Gannan Tibetan harbored increased ancestry related to Bonze Age Steppe Pastoralists (Sintashta: Russia_Sintashta_MLBA and Russia_MLBA_Sintashta, Yamnaya: Yamnaya_Kalmykia, Yamnaya_Samara and others: Russia_Srubnaya, Russia_Andronovo, Russia_Afanasievo) and it also harbored more coastal Neolithic to modern southern East Asian ancestry (Qihe_EN, Tanshishan_LN, Taiwan_Hanben, Atayal and Ami).

Relative to Lhasa Tibetan, Gannan Tibetan shared more coastal early Neolithic southern East Asian Qihe ancestry and modern southern East Asian Atayal ancestry, and it also owned increased ancestry related to the Steppe pastoralists (Yamnaya_Kalmykia, Yamnaya_Samara, Russia_MLBA_Sintashta, Russia_Sintashta_MLBA, Russia_Andronovo, Russia_Srubnaya, Russia_Afanasievo).

Relative to Nagqu Tibetan, Gannan Tibetan harbored more western Eurasian Stepped Pastoralists-related ancestry (Russia_MLBA_Sintashta, Russia_Srubnaya, Russia_Sintashta_MLBA, Russia_Andronovo, Yamnaya_Kalmykia, Russia_Afanasievo and Yamnaya_Samara). Similar patterns of Steppe Pastoralists affinity were observed in *$f_4$(Chamdo Tibetan, Gannan Tibetan; early and middle Bronze Age Steppe Pastoralists, Mbuti)*.

Compared with Gangcha Tibetan from Qinghai Province, Gannan Tibetan harbored increased ancestry related to coastal Neolithic to Bronze Age to modern southern East Asian (Qihe_EN, Tanshishan_LN, Atayal, Taiwan_Hanben, Taiwan_Gongguan and Ami).

The results of *$f_4$(Xinlong/Yajiang Tibetan, Gannan Tibetan; early and middle Bronze Age Steppe Pastoralists, Mbuti)* showed significant negative $f_4$-statistics values, which suggested more Steppe Pastoralists-related ancestry in Gannan Tibetan than in Xinjiang Tibetan and Yajiang Tibetan from Sichuan Province. This western Eurasian Steppe Pastoralist affinity was also identified via *$f_4$(Yunnan Tibetan, Gannan Tibetan; early and middle Bronze Age Steppe Pastoralists, Mbuti)*.

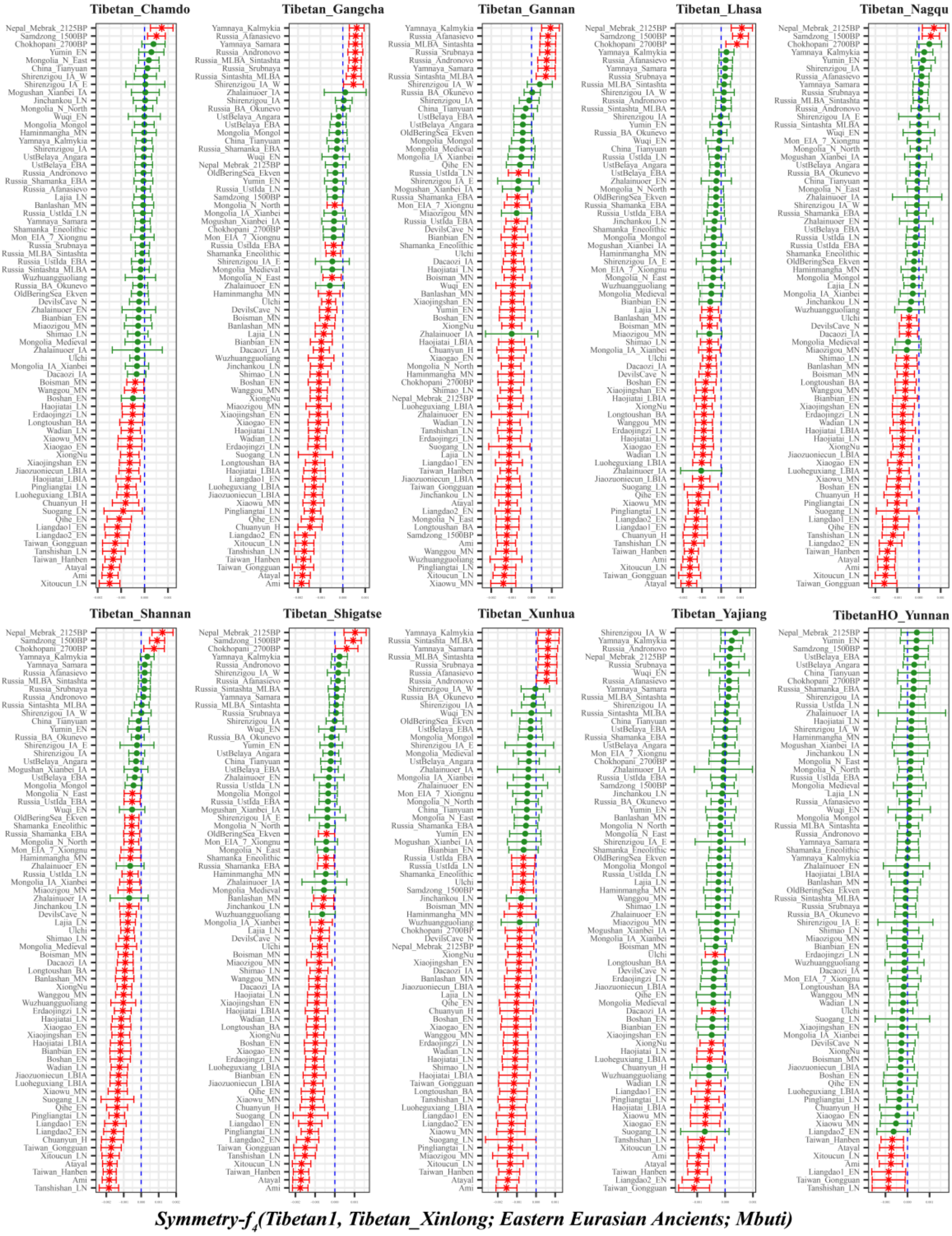

**Figure S23. Genomic affinity between modern Tibetans and eastern Eurasian ancient populations inferred from four population symmetry-*f*$_4$ statistics of the form *f*$_4$*(Tibetan1, Xinlong Tibetan; eastern Eurasian ancients, Mbuti)*.**

Here, overlapping SNP loci included in the Affymetrix Human Origins platform among four analyzed populations were used. We used the genetic variation of Mbuti as the outgroup. Red asterisk point meant the significant value (Absolute value of Z-scores larger than three or equal to three) observed in the symmetry-$f_4$ statistics and green circle point denoted the non-significant $f_4$-statistic values (Absolute value of Z-scores less than three). All Tibetan2 were listed along the Y-axis and $f_4$ values were labeled along the X-axis. All tested population pairs were faceted or grouped via the Tibetan1. Significant negative $f_4$ values indicated that the included eastern Eurasian ancient population shared more alleles with Xinlong Tibetan compared with Tibetan1 or Xinlong Tibetan harbored increased eastern Eurasian ancient population-related ancestry compared with Tibetan1, and significant positive $f_4$ value indicated that the included eastern Eurasian ancient population shared more derived alleles with Tibetan1 compared with Xinlong Tibetan or elucidated as Xinlong Tibetan had increased eastern Eurasian ancient population-related ancestry relative to Tibetan1. The value of $f_4$-statistics equal to zero was marked as the blue dash line. The bar indicated three standard errors.

Compared to Yunnan Tibetan, Xinlong Tibetan harbored increased ancestry related to Coastal/island Neolithic to modern southern East Asian (Tanshishan_LN, Taiwan_Gongguan, Liangdao1_EN, Ami, Xitoucun_LN, Atayal and Taiwan_Hanben). Compared with other Tibetans, Xinlong Tibetan owned more lowland ancient or modern East Asian-related ancestry.

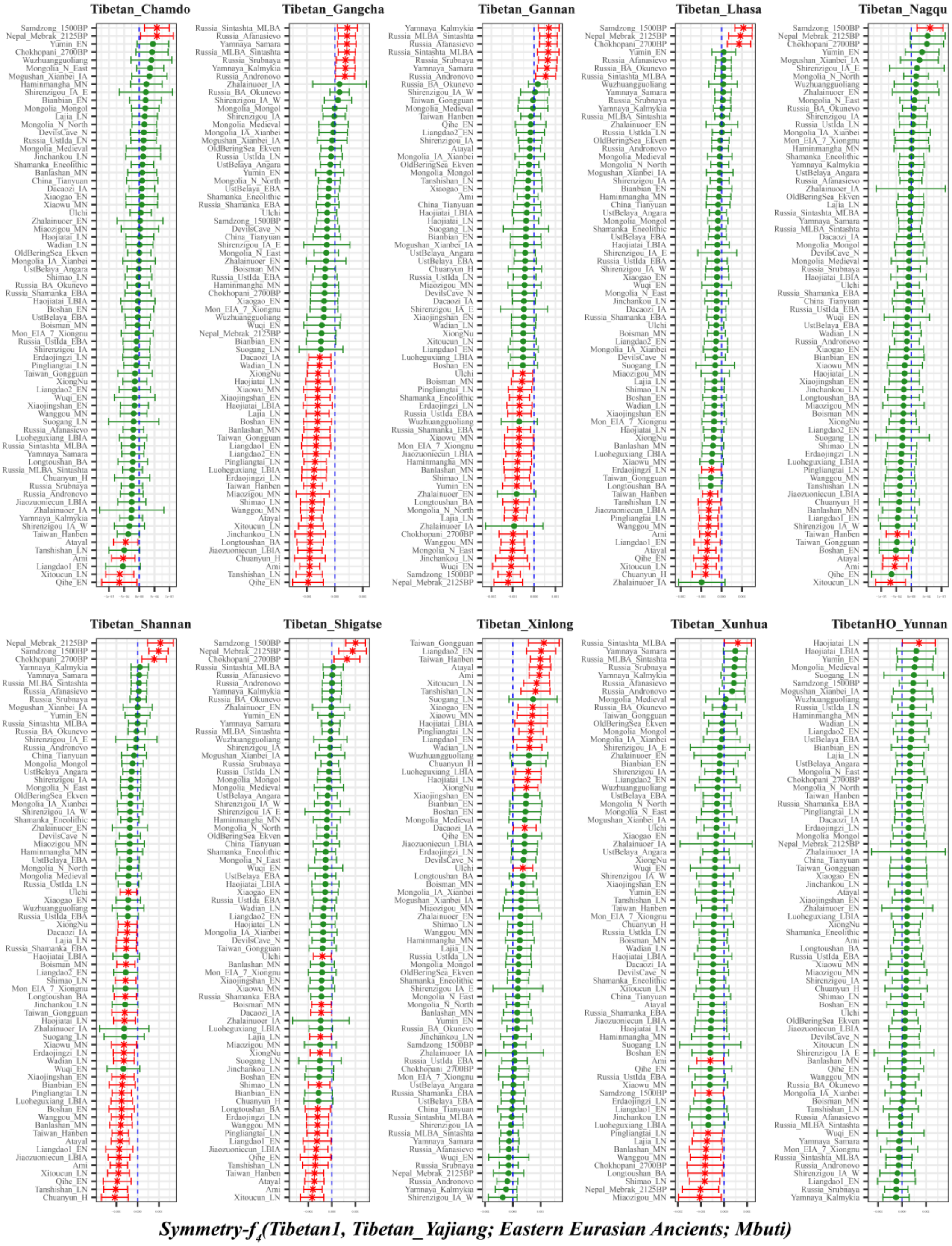

**Figure S24. Genomic affinity between modern Tibetans and eastern Eurasian ancient populations inferred from four population symmetry-$f_4$ statistics of the form $f_4$*(Tibetan1, Yajiang Tibetan; eastern Eurasian ancients, Mbuti)*.**

Here, overlapping SNP loci included in the Affymetrix Human Origins platform among four analyzed populations were used. We used the genetic variation of Mbuti as the outgroup. Red asterisk point meant

the significant value (Absolute value of Z-scores larger than three or equal to three) observed in the symmetry-$f_4$ statistics and green circle point denoted the non-significant $f_4$-statistic values (Absolute value of Z-scores less than three). All Tibetan2 were listed along the Y-axis and $f_4$ values were labeled along the X-axis. All tested population pairs were faceted or grouped via the Tibetan1. Significant negative $f_4$ values indicated that the included eastern Eurasian ancient population shared more alleles with Yajiang Tibetan compared with Tibetan1 or Yajiang Tibetan harbored increased eastern Eurasian ancient population-related ancestry compared with Tibetan1, and significant positive $f_4$ value indicated that the included eastern Eurasian ancient population shared more derived alleles with Tibetan1 compared with Yajiang Tibetan or elucidated as Yajiang Tibetan had increased eastern Eurasian ancient population-related ancestry relative to Tibetan1. The value of $f_4$-statistics equal to zero was marked as the blue dash line. The bar indicated three standard errors.

Compared with Xinlong Tibetan and Yunnan Tibetan, no signals of additional shared ancestry from other source populations into Yajiang Tibetan were identified, as no significant negative $f_4$-statistics were observed in $f_4$*(Xinlong/Yunnan Tibetan, Yajiang Tibetan; eastern Eurasian ancients, Mbuti)*. Compared with other Tibetans, Yajiang harbored increased lowland East Asian ancestry and especially for more coastal southern East Asian ancestry. We also find more highland Nepal ancient-related ancestry in Yajiang Tibetan relative to Gannan Tibetan and Xunhua Tibetan.

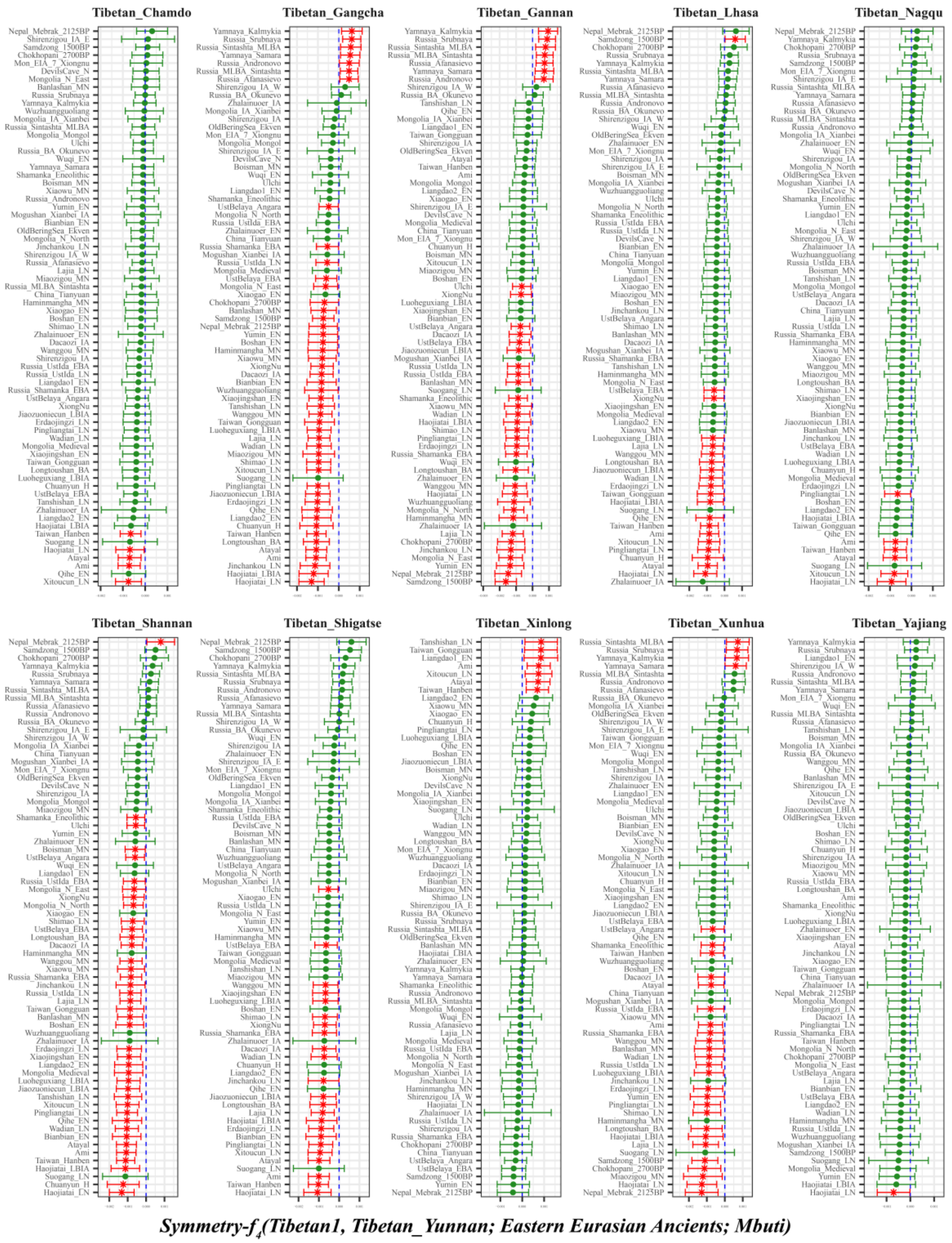

**Figure S25. Genomic affinity between modern Tibetans and eastern Eurasian ancient populations inferred from four population symmetry-$f_4$ statistics of the form $f_4$*(Tibetan1, Yunnan Tibetan; eastern Eurasian ancients, Mbuti)*.**

Here, overlapping SNP loci included in the Affymetrix Human Origins platform among four analyzed populations were used. We used the genetic variation of Mbuti as the outgroup. Red asterisk point meant the significant value (Absolute value of Z-scores larger than three or equal to three) observed in the symmetry-$f_4$ statistics and green circle point denoted the non-significant $f_4$-statistic values (Absolute value of Z-scores less than three). All Tibetan2 were listed along the Y-axis and $f_4$ values were labeled along the X-axis. All tested population pairs were faceted or grouped via the Tibetan1. Significant negative $f_4$ values indicated that the included eastern Eurasian ancient population shared more alleles with Yunnan Tibetan compared with Tibetan1 or Yunnan Tibetan harbored increased eastern Eurasian ancient population-related ancestry compared with Tibetan1, and significant positive $f_4$ value indicated that the included eastern Eurasian ancient population shared more derived alleles with Tibetan1 compared with Yunnan Tibetan or elucidated as Yunnan Tibetan had increased eastern Eurasian ancient population-related ancestry relative to Tibetan1. The value of $f_4$-statistics equal to zero was marked as the blue dash line. The bar indicated three standard errors.

Compared with Yajiang Tibetan and Xinlong Tibetan from Sichuan Province, no signals of additional gene flow into Yunnan Tibetan, except for Yunnan had more Neolithic Haojiatai-related ancestry related to Yajiang Tibetan. Relative to other Tibetans, Yunnan Tibetan harbored more lowland East Asian related ancestry. We also found that compared with Gannan Tibetan, Yunnan Tibetan had more Nepal ancients-related ancestry.

**Shared ancestry with temporally different populations**

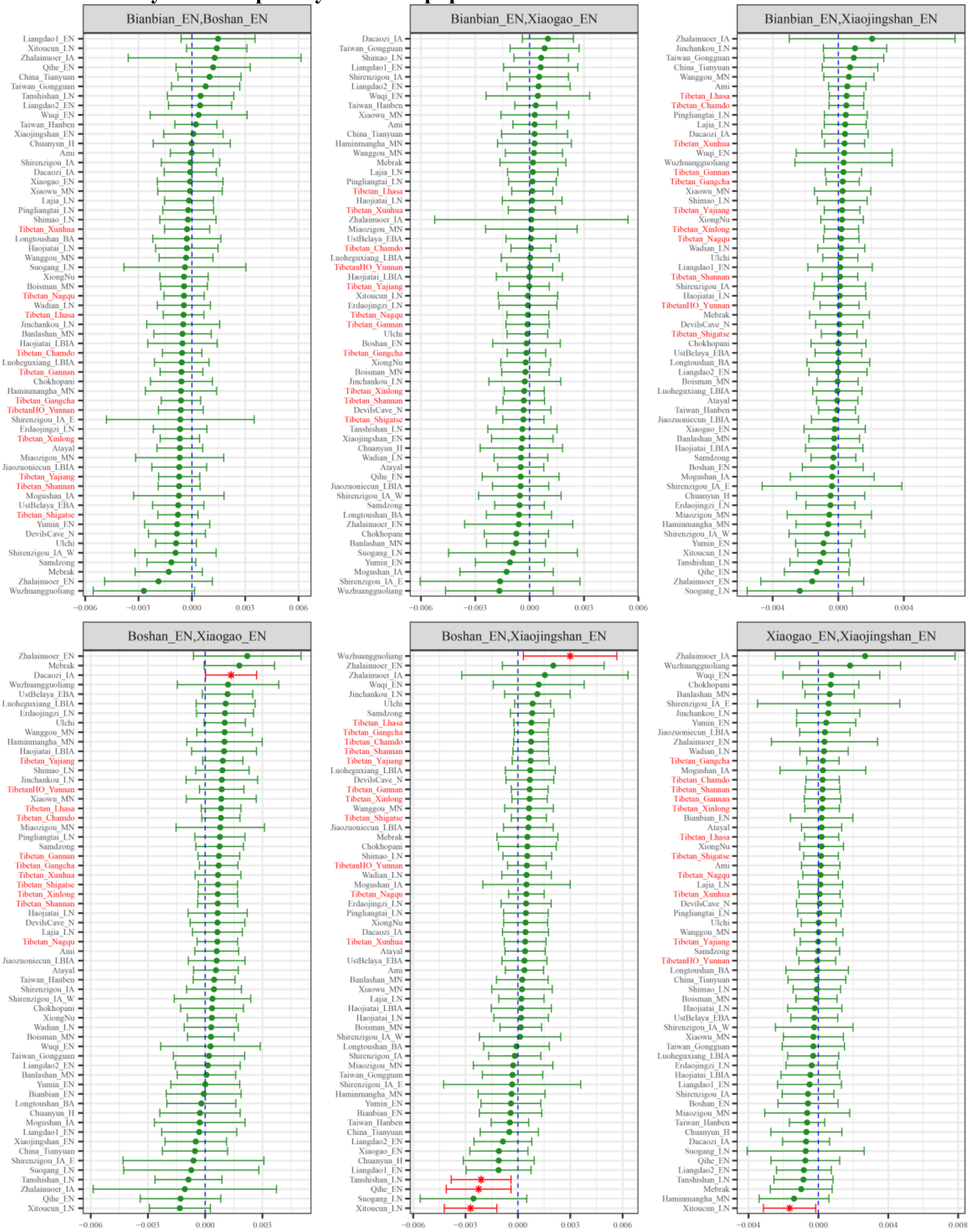

*$f_4$(Coastal Neolithic northern East Asian1, Coastal Neolithic northern East Asian2; Eastern Modern Tibetan/Ancient East Asians, Mbuti)*

**Figure S26. Shared ancestry associated with coastal Neolithic northern East Asian in modern Tibetans and ancient East Asians compared with geographically different coastal Neolithic northern East Asian inferred from four population symmetry-$f_4$-statistics of the form $f_4$*(Coastal Neolithic northern East Asian1, Coastal Neolithic northern East Asian2; Eastern Modern Tibetan/Ancient East Asians, Mbuti)*.**

Here, overlapping SNP loci included in the Affymetrix Human Origins platform among four analyzed populations were used. We used the genetic variation of Mbuti as the outgroup. Red asterisk point meant the significant value (Absolute value of Z-scores larger than three or equal to three) observed in the symmetry-$f_4$ statistics and green circle point denoted the non-significant $f_4$-statistic values (Absolute value of Z-scores less than three). All Eastern Modern Tibetan/Ancient East Asians were listed along the Y-axis and $f_4$ values were labeled along the X-axis. All tested population pairs were faceted or grouped via the combination of Coastal Neolithic northern East Asian1 and Coastal Neolithic northern East

Asian2. Significant negative $f_4$ values indicated that the third population shared more alleles with the second population and significant positive $f_4$ value indicated that the third population shared more derived alleles with the first population. The value of $f_4$-statistics equal to zero was marked as the blue dash line. The bar indicated three standard errors.

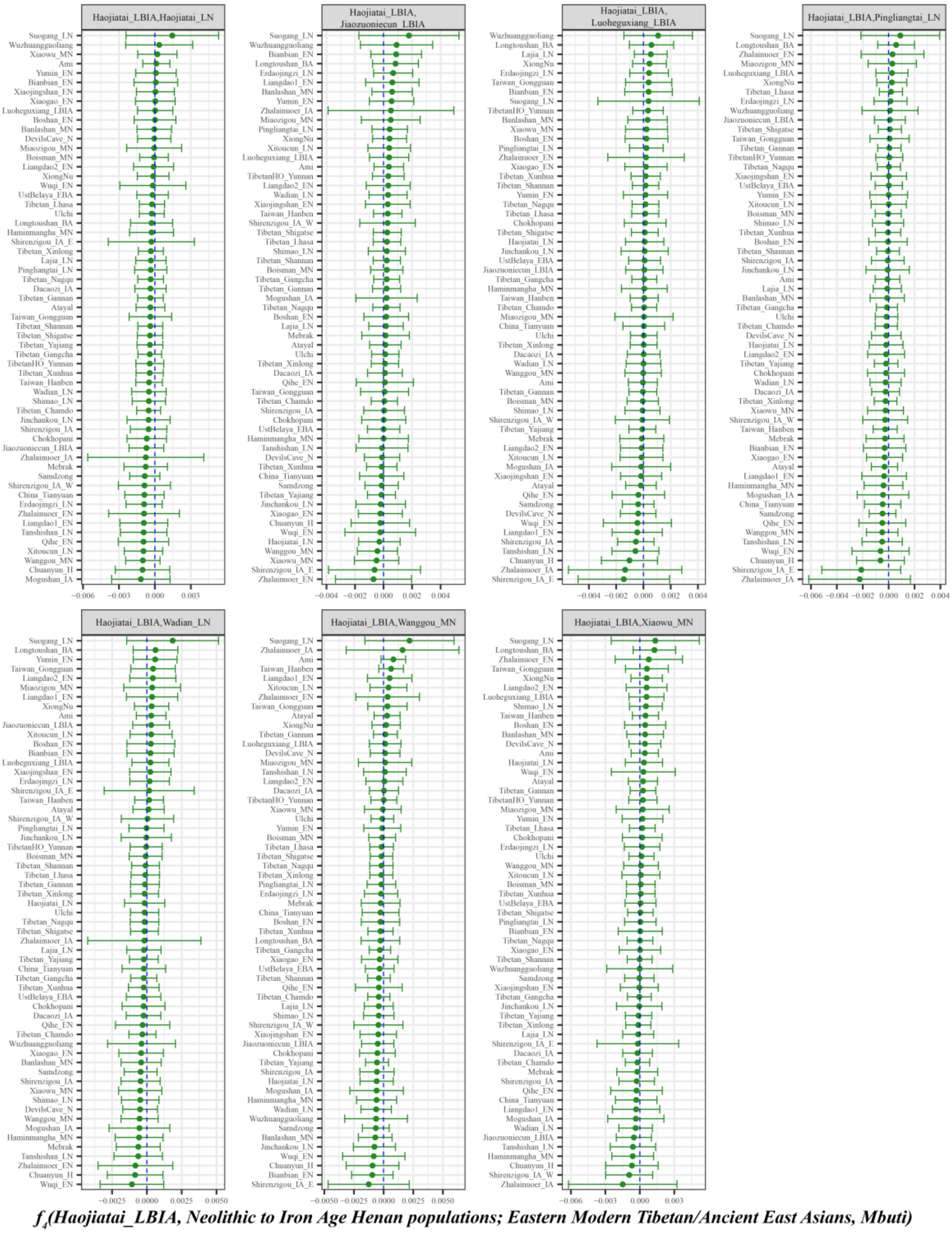

**Figure S27. Shared ancestry associated with inland Neolithic to Iron Age northern East Asian from Henan province in modern Tibetans and ancient East Asians compared with geographically different ancient Henan populations inferred from four population symmetry-$f_4$-statistics of the form *$f_4$(Haojiatai_LBIA, Neolithic to Iron Age Henan populations; Eastern Modern Tibetan/Ancient East Asians, Mbuti)*.**

Here, overlapping SNP loci included in the Affymetrix Human Origins platform among four analyzed populations were used. We used the genetic variation of Mbuti as the outgroup. Red asterisk point meant

the significant value (Absolute value of Z-scores larger than three or equal to three) observed in the symmetry-$f_4$ statistics and green circle point denoted the non-significant $f_4$-statistic values (Absolute value of Z-scores less than three). All Eastern Modern Tibetan/Ancient East Asians were listed along the Y-axis and $f_4$ values were labeled along the X-axis. All tested population pairs were faceted or grouped via the combination of the first and second populations. Significant negative $f_4$ values indicated that the third population shared more alleles with the second population and significant positive $f_4$ value indicated that the third population shared more derived alleles with the first population. The value of $f_4$-statistics equal to zero was marked as the blue dash line. The bar indicated three standard errors.

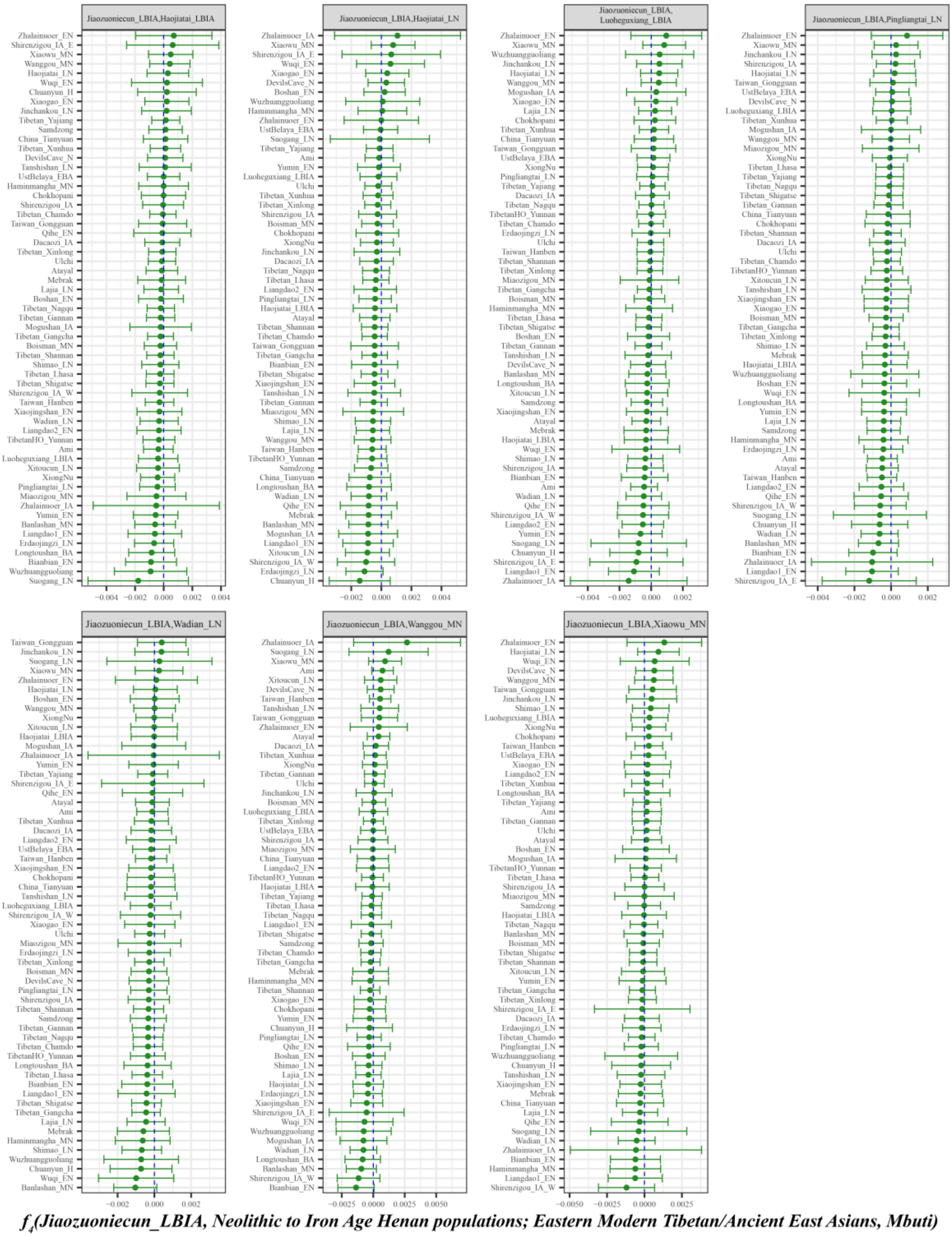

**Figure S28. Shared ancestry associated with inland Neolithic to Iron Age northern East Asian from Henan province in modern Tibetans and ancient East Asians compared with geographically different ancient Henan populations inferred from four population symmetry-$f_4$-statistics of the form $f_4$*(Jiaozuoniecun_LBIA, Neolithic to Iron Age Henan populations; Eastern Modern Tibetan/Ancient East Asians, Mbuti)*.**

Here, overlapping SNP loci included in the Affymetrix Human Origins platform among four analyzed populations were used. We used the genetic variation of Mbuti as the outgroup. Red asterisk point meant the significant value (Absolute value of Z-scores larger than three or equal to three) observed in the symmetry-$f_4$ statistics and green circle point denoted the non-significant $f_4$-statistic values (Absolute value of Z-scores less than three). All Eastern Modern Tibetan/Ancient East Asians were listed along the Y-axis and $f_4$ values were labeled along the X-axis. All tested population pairs were faceted or grouped via the combination of the first and second populations. Significant negative $f_4$ values indicated that the third population shared more alleles with the second population and significant positive $f_4$ value indicated that the third population shared more derived alleles with the first population. The value of $f_4$-statistics equal to zero was marked as the blue dash line. The bar indicated three standard errors.

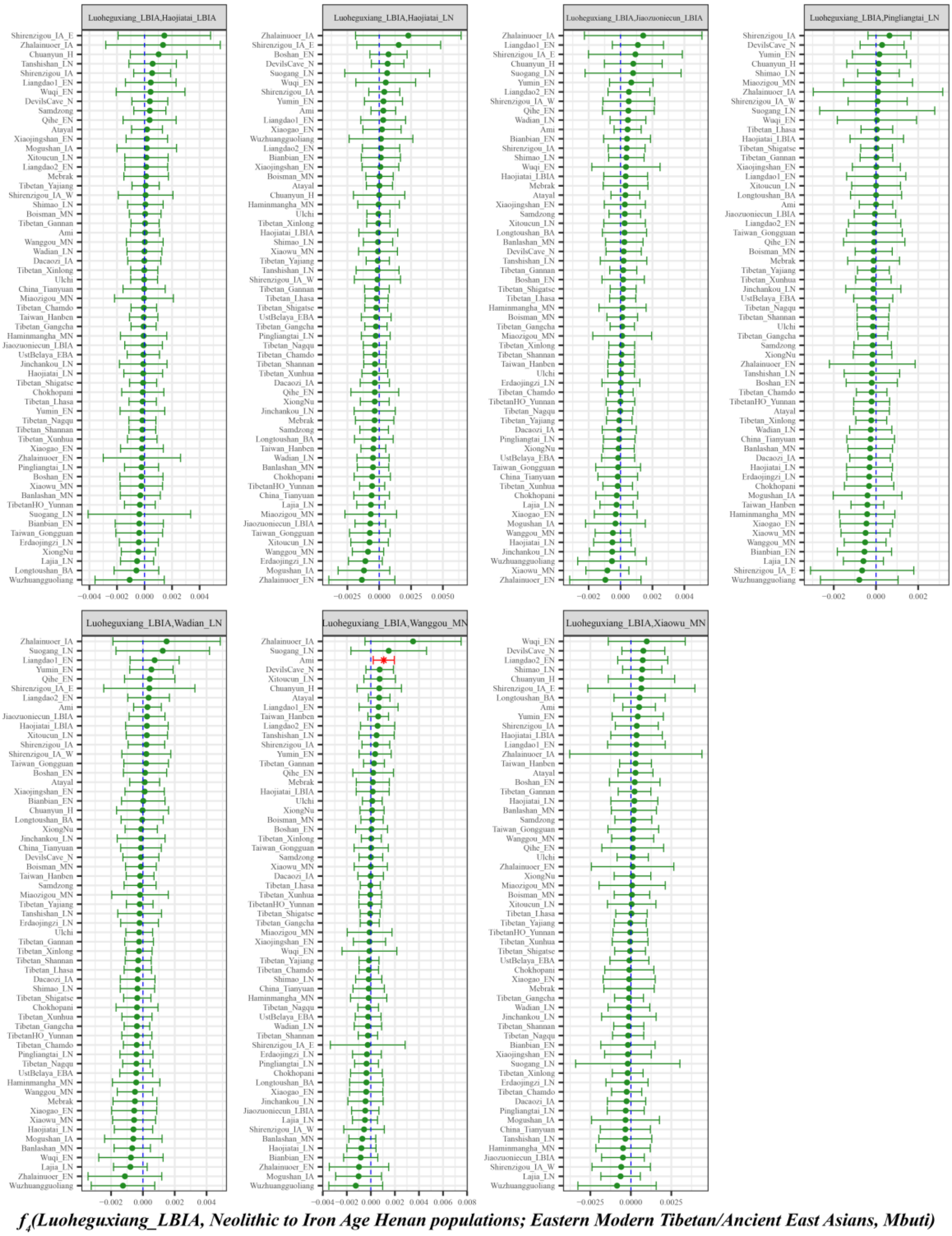

**Figure S29. Shared ancestry associated with inland Neolithic to Iron Age northern East Asian from Henan province in modern Tibetans and ancient East Asians compared with geographically different ancient Henan populations inferred from four population symmetry-$f_4$-statistics of the form $f_4$*(Luoheguxiang_LBIA, Neolithic to Iron Age Henan populations; Eastern Modern Tibetan/Ancient East Asians, Mbuti)*.**

Here, overlapping SNP loci included in the Affymetrix Human Origins platform among four analyzed populations were used. We used the genetic variation of Mbuti as the outgroup. Red asterisk point meant the significant value (Absolute value of Z-scores larger than three or equal to three) observed in the symmetry-$f_4$ statistics and green circle point denoted the non-significant $f_4$-statistic values (Absolute value of Z-scores less than three). All Eastern Modern Tibetan/Ancient East Asians were listed along the Y-axis and $f_4$ values were labeled along the X-axis. All tested population pairs were faceted or grouped via the combination of the first and second populations. Significant negative $f_4$ values indicated that the third population shared more alleles with the second population and significant positive $f_4$ value indicated

that the third population shared more derived alleles with the first population. The value of $f_4$-statistics equal to zero was marked as the blue dash line. The bar indicated three standard errors.

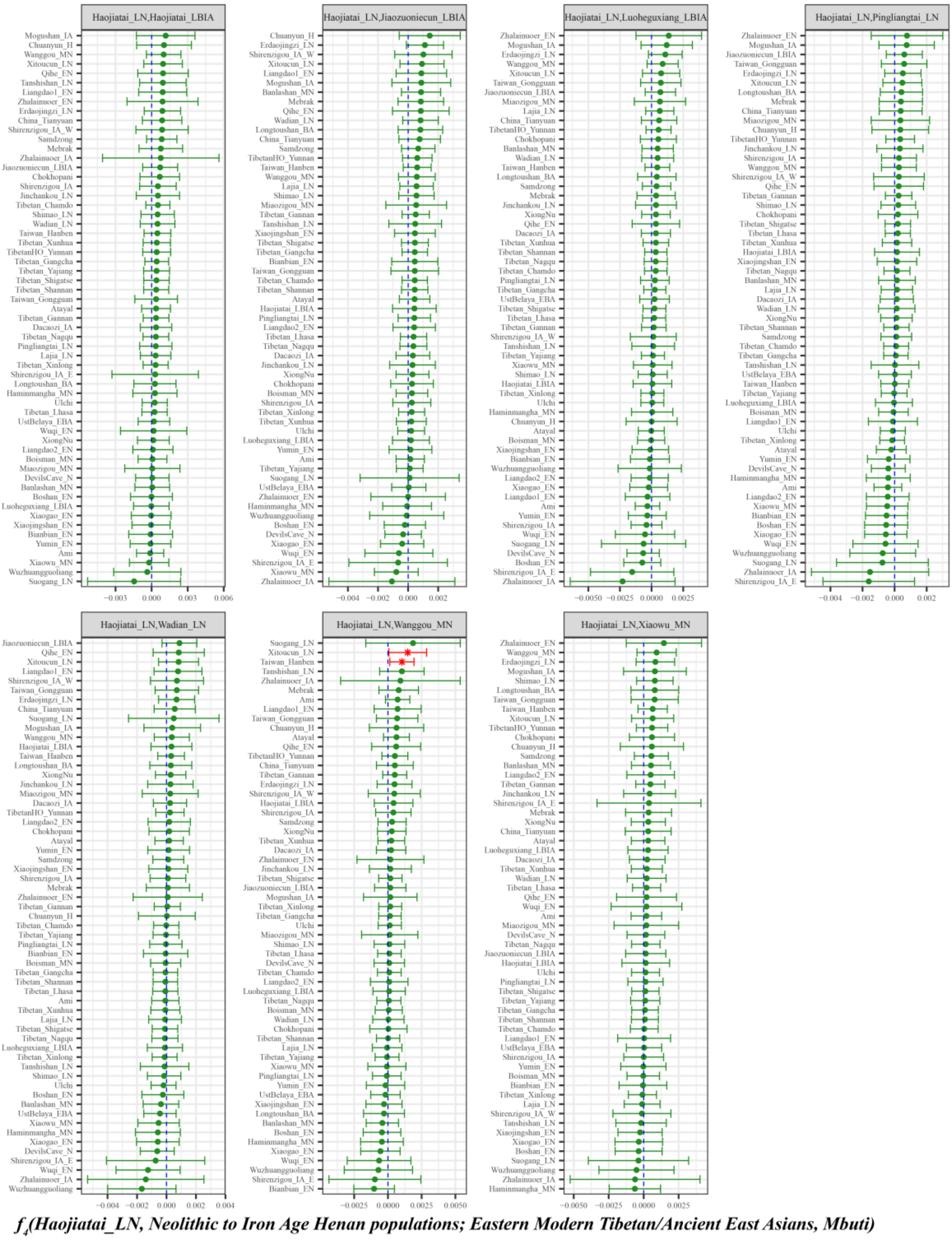

**Figure S30. Shared ancestry associated with inland Neolithic to Iron Age northern East Asian from Henan province in modern Tibetans and ancient East Asians compared with geographically different ancient Henan populations inferred from four population symmetry-$f_4$-statistics of the form $f_4$*(Haojiatai_LN, Neolithic to Iron Age Henan populations; Eastern Modern Tibetan/Ancient East Asians, Mbuti)*.**

Here, overlapping SNP loci included in the Affymetrix Human Origins platform among four analyzed populations were used. We used the genetic variation of Mbuti as the outgroup. Red asterisk point meant the significant value (Absolute value of Z-scores larger than three or equal to three) observed in the symmetry-$f_4$ statistics and green circle point denoted the non-significant $f_4$-statistic values (Absolute

value of Z-scores less than three). All Eastern Modern Tibetan/Ancient East Asians were listed along the Y-axis and $f_4$ values were labeled along the X-axis. All tested population pairs were faceted or grouped via the combination of the first and second populations. Significant negative $f_4$ values indicated that the third population shared more alleles with the second population and significant positive $f_4$ value indicated that the third population shared more derived alleles with the first population. The value of $f_4$-statistics equal to zero was marked as the blue dash line. The bar indicated three standard errors.

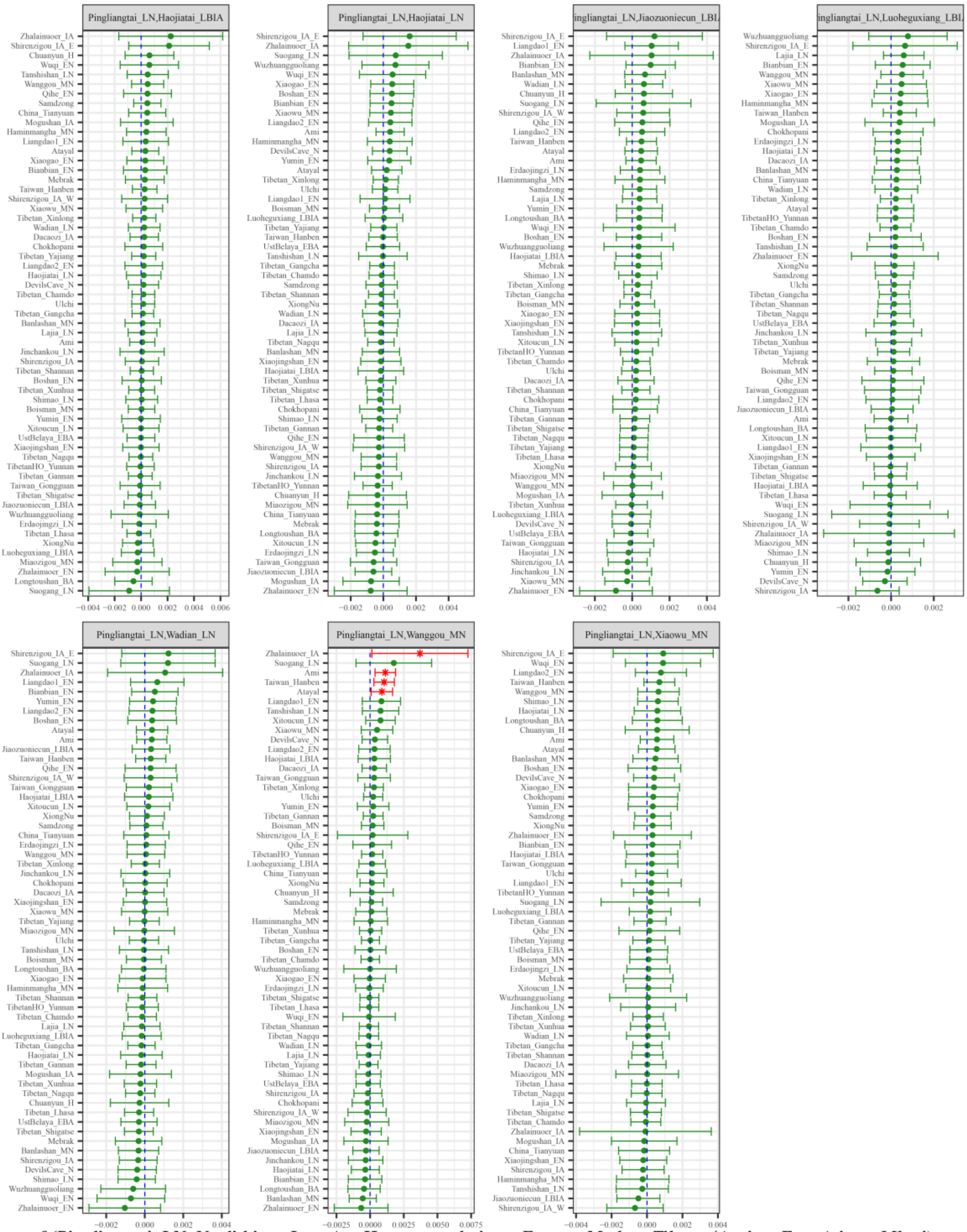

***$f_4$(Pingliangtai_LN, Neolithic to Iron Age Henan populations; Eastern Modern Tibetan/Ancient East Asians, Mbuti)***

Figure S31. Shared ancestry associated with inland Neolithic to Iron Age northern East Asian from Henan province in modern Tibetans and ancient East Asians compared with geographically different ancient Henan populations inferred from four population symmetry-$f_4$-statistics of the form *$f_4$(Pingliangtai_LN, Neolithic to Iron Age Henan populations; Eastern Modern Tibetan/Ancient East Asians, Mbuti).*

Here, overlapping SNP loci included in the Affymetrix Human Origins platform among four analyzed populations were used. We used the genetic variation of Mbuti as the outgroup. Red asterisk point meant the significant value (Absolute value of Z-scores larger than three or equal to three) observed in the symmetry-$f_4$ statistics and green circle point denoted the non-significant $f_4$-statistic values (Absolute value of Z-scores less than three). All Eastern Modern Tibetan/Ancient East Asians were listed along the Y-axis and $f_4$ values were labeled along the X-axis. All tested population pairs were faceted or grouped via the combination of the first and second populations. Significant negative $f_4$ values indicated that the third population shared more alleles with the second population and significant positive $f_4$ value indicated that the third population shared more derived alleles with the first population. The value of $f_4$-statistics equal to zero was marked as the blue dash line. The bar indicated three standard errors.

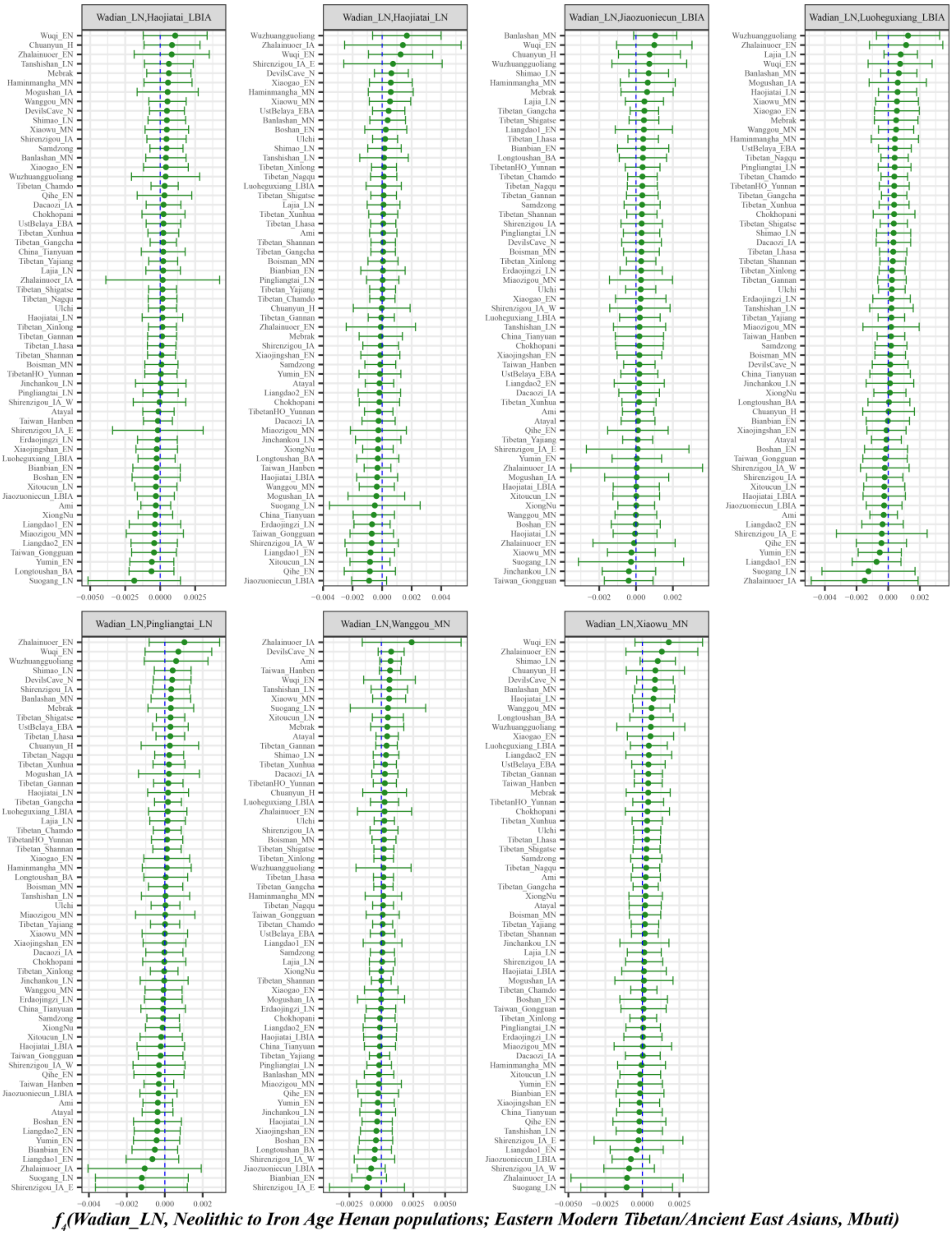

**Figure S32. Shared ancestry associated with inland Neolithic to Iron Age northern East Asian from Henan province in modern Tibetans and ancient East Asians compared with geographically different ancient Henan populations inferred from four population symmetry-$f_4$-statistics of the form $f_4$*(Wadian_LN, Neolithic to Iron Age Henan populations; Eastern Modern Tibetan/Ancient East Asians, Mbuti)*.**

Here, overlapping SNP loci included in the Affymetrix Human Origins platform among four analyzed populations were used. We used the genetic variation of Mbuti as the outgroup. Red asterisk point meant the significant value (Absolute value of Z-scores larger than three or equal to three) observed in the symmetry-$f_4$ statistics and green circle point denoted the non-significant $f_4$-statistic values (Absolute value of Z-scores less than three). All Eastern Modern Tibetan/Ancient East Asians were listed along the Y-axis and $f_4$ values were labeled along the X-axis. All tested population pairs were faceted or grouped via the combination of the first and second populations. Significant negative $f_4$ values indicated that the third population shared more alleles with the second population and significant positive $f_4$ value indicated

that the third population shared more derived alleles with the first population. The value of $f_4$-statistics equal to zero was marked as the blue dash line. The bar indicated three standard errors.

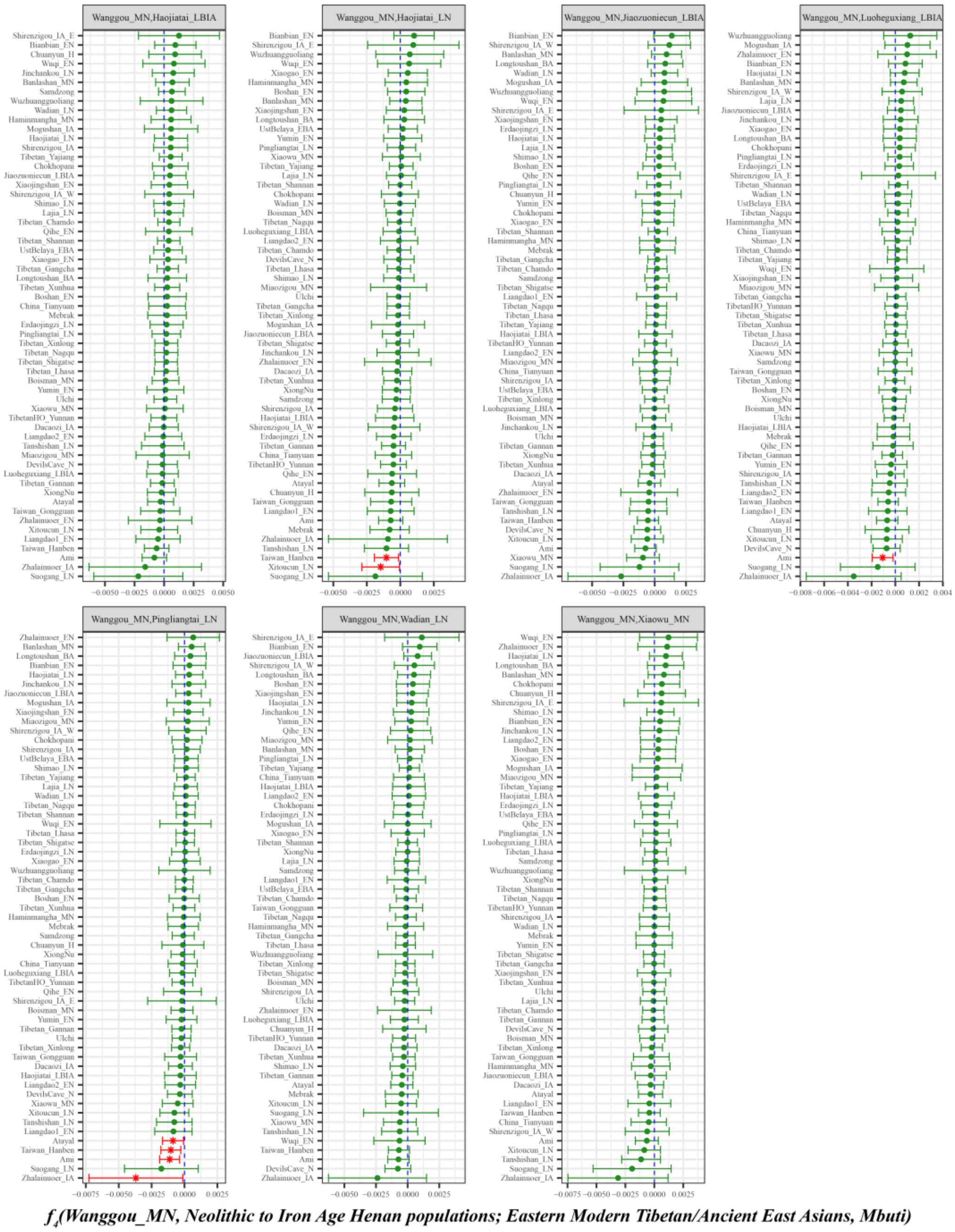

**Figure S33. Shared ancestry associated with inland Neolithic to Iron Age northern East Asian from Henan province in modern Tibetans and ancient East Asians compared with geographically different ancient Henan populations inferred from four population symmetry-$f_4$-statistics of the form $f_4$*(Wanggou_MN, Neolithic to Iron Age Henan populations; Eastern Modern Tibetan/Ancient East Asians, Mbuti)*.**

Here, overlapping SNP loci included in the Affymetrix Human Origins platform among four analyzed populations were used. We used the genetic variation of Mbuti as the outgroup. Red asterisk point meant the significant value (Absolute value of Z-scores larger than three or equal to three) observed in the symmetry-$f_4$ statistics and green circle point denoted the non-significant $f_4$-statistic values (Absolute value of Z-scores less than three). All Eastern Modern Tibetan/Ancient East Asians were listed along the Y-axis and $f_4$ values were labeled along the X-axis. All tested population pairs were faceted or grouped via the combination of the first and second populations. Significant negative $f_4$ values indicated that the third population shared more alleles with the second population and significant positive $f_4$ value indicated that the third population shared more derived alleles with the first population. The value of $f_4$-statistics equal to zero was marked as the blue dash line. The bar indicated three standard errors.

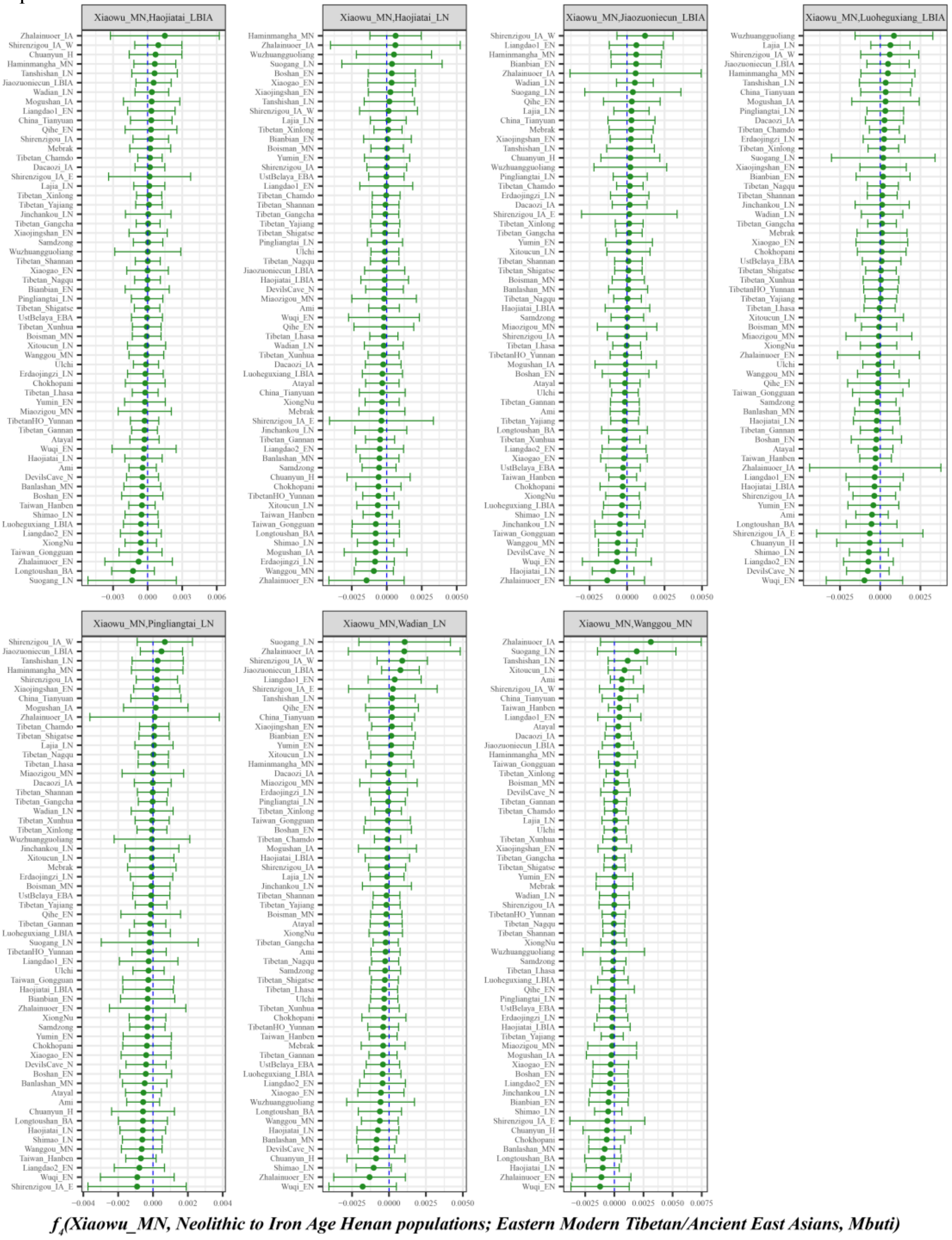

**Figure S34. Shared ancestry associated with inland Neolithic to Iron Age northern East Asian from Henan province in modern Tibetans and ancient East Asians compared with geographically different ancient Henan populations inferred from four population symmetry-$f_4$-statistics of the form $f_4$*(Xiaowu_MN, Neolithic to Iron Age Henan populations; Eastern Modern Tibetan/Ancient East Asians, Mbuti)*.**

Here, overlapping SNP loci included in the Affymetrix Human Origins platform among four analyzed populations were used. We used the genetic variation of Mbuti as the outgroup. Red asterisk point meant the significant value (Absolute value of Z-scores larger than three or equal to three) observed in the symmetry-$f_4$ statistics and green circle point denoted the non-significant $f_4$-statistic values (Absolute value of Z-scores less than three). All Eastern Modern Tibetan/Ancient East Asians were listed along the Y-axis and $f_4$ values were labeled along the X-axis. All tested population pairs were faceted or grouped via the combination of the first and second populations. Significant negative $f_4$ values indicated that the third population shared more alleles with the second population and significant positive $f_4$ value indicated

that the third population shared more derived alleles with the first population. The value of $f_4$-statistics equal to zero was marked as the blue dash line. The bar indicated three standard errors.

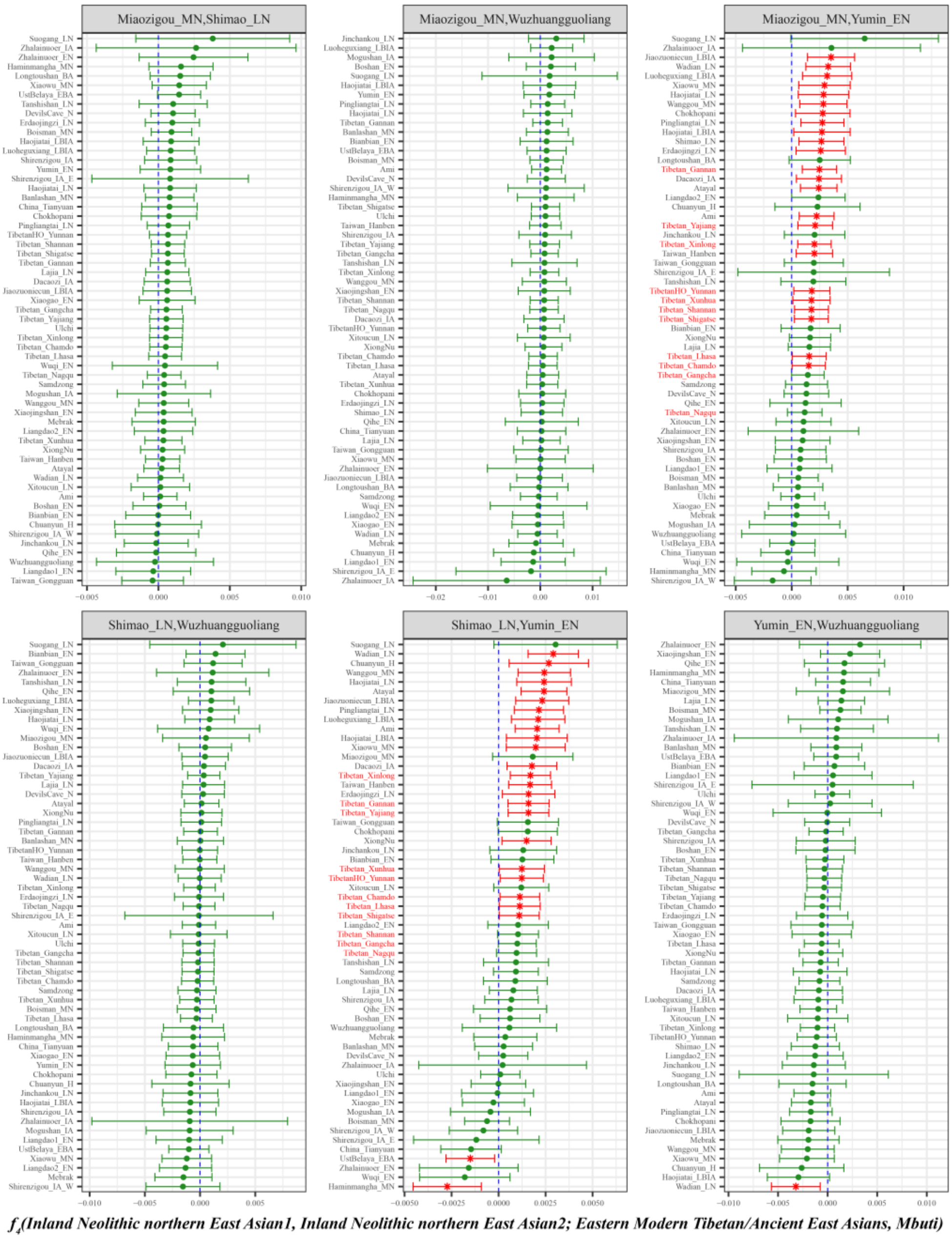

**Figure S35. Shared ancestry associated with inland Neolithic northern East Asian from Shaanxi and Inner Mongolia provinces in modern Tibetans and ancient East Asians compared with geographically different ancient populations inferred from four population symmetry-$f_4$-statistics of the form *$f_4$(Inland Neolithic northern East Asian1, Inland Neolithic northern East Asian2; Eastern Modern Tibetan/Ancient East Asians, Mbuti)*.**

Here, overlapping SNP loci included in the Affymetrix Human Origins platform among four analyzed populations were used. We used the genetic variation of Mbuti as the outgroup. Red asterisk point meant the significant value (Absolute value of Z-scores larger than three or equal to three) observed in the symmetry-$f_4$ statistics and green circle point denoted the non-significant $f_4$-statistic values (Absolute value of Z-scores less than three). All Eastern Modern Tibetan/Ancient East Asians were listed along the Y-axis and $f_4$ values were labeled along the X-axis. All tested population pairs were faceted or grouped via the combination of the first and second populations. Significant negative $f_4$ values indicated that the third population shared more alleles with the second population and significant positive $f_4$ value indicated that the third population shared more derived alleles with the first population. The value of $f_4$-statistics equal to zero was marked as the blue dash line. The bar indicated three standard errors.

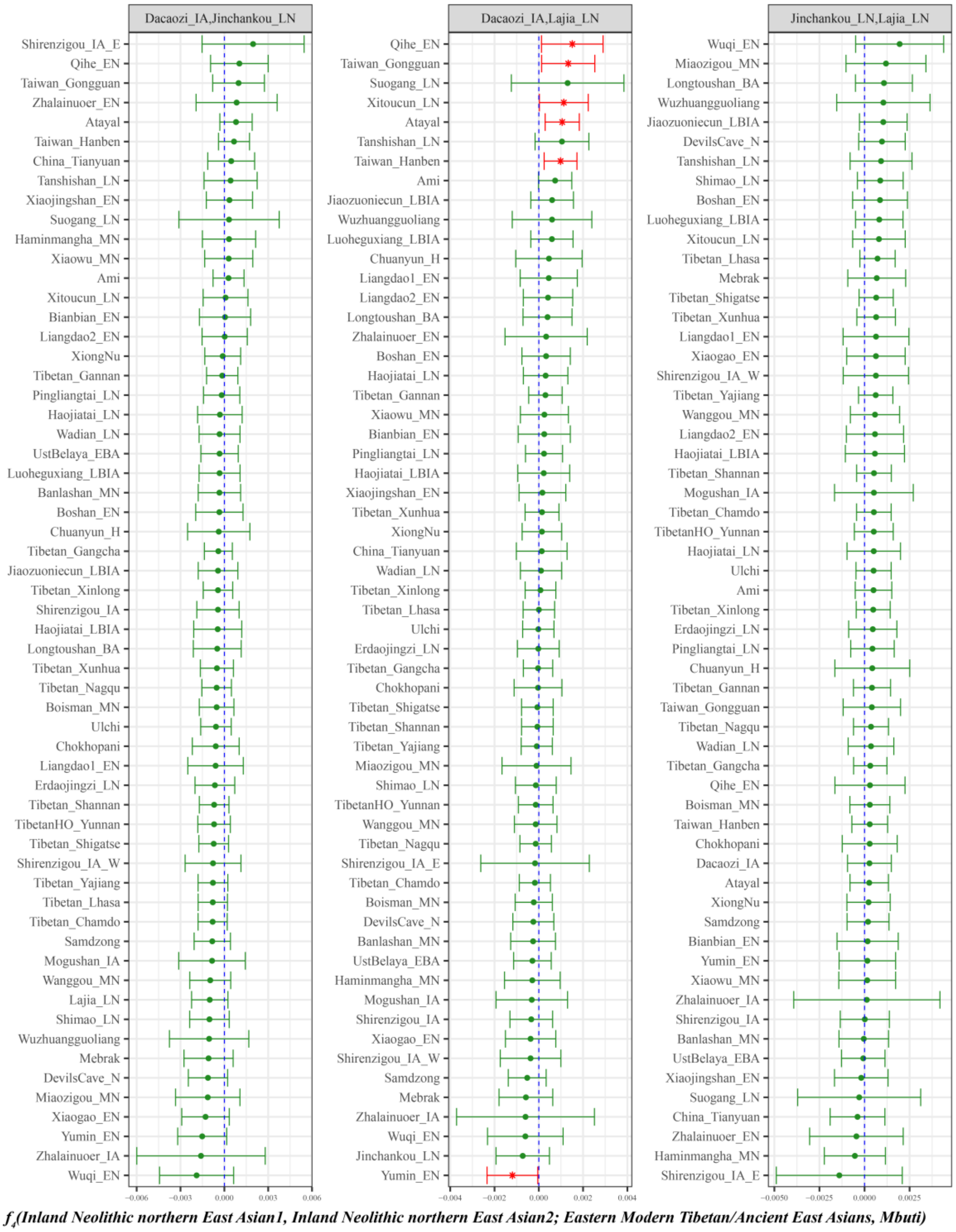

**Figure S36. Shared ancestry associated with inland Neolithic northern East Asian from Qinghai province in modern Tibetans and ancient East Asians compared with geographically different ancient populations inferred from four population symmetry-$f_4$-statistics of the form $f_4$*(Inland Neolithic northern East Asian1, Inland Neolithic northern East Asian2; Eastern Modern Tibetan/Ancient East Asians, Mbuti)*.**

Here, overlapping SNP loci included in the Affymetrix Human Origins platform among four analyzed populations were used. We used the genetic variation of Mbuti as the outgroup. Red asterisk point meant the significant value (Absolute value of Z-scores larger than three or equal to three) observed in the symmetry-$f_4$ statistics and green circle point denoted the non-significant $f_4$-statistic values (Absolute value of Z-scores less than three). All Eastern Modern Tibetan/Ancient East Asians were listed along the Y-axis and $f_4$ values were labeled along the X-axis. All tested population pairs were faceted or grouped via the combination of the first and second populations. Significant negative $f_4$ values indicated that the third population shared more alleles with the second population and significant positive $f_4$ value indicated that the third population shared more derived alleles with the first population. The value of $f_4$-statistics equal to zero was marked as the blue dash line. The bar indicated three standard errors.

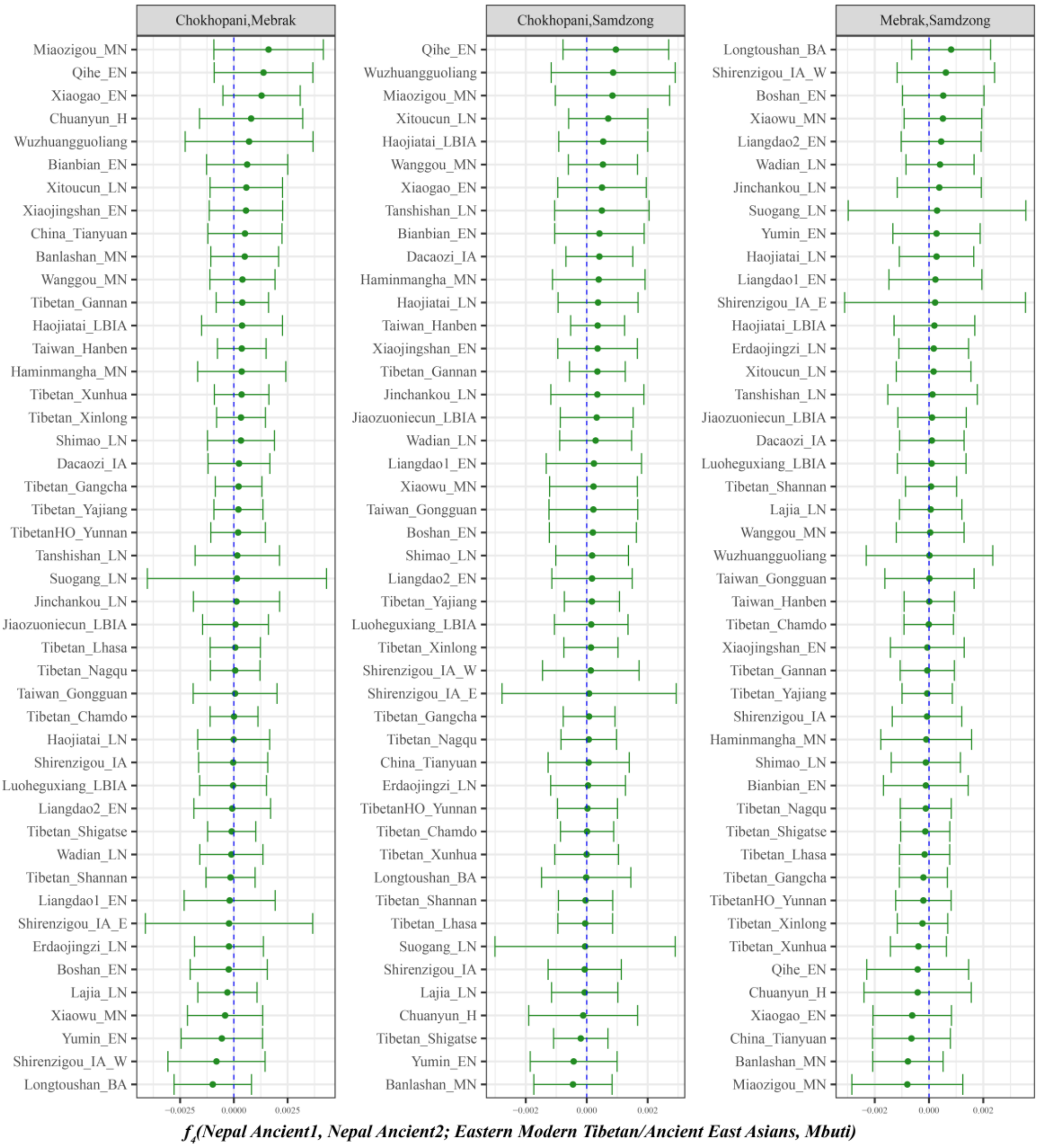

**Figure S37. Shared ancestry associated with Bronze Age to historic period from Nepal in modern Tibetans and ancient East Asians compared with geographically different ancient populations inferred from four population symmetry-$f_4$-statistics of the form $f_4$*(Nepal Ancient1, Nepal Ancient2; Eastern Modern Tibetan/Ancient East Asians, Mbuti)*.**

Here, overlapping SNP loci included in the Affymetrix Human Origins platform among four analyzed populations were used. We used the genetic variation of Mbuti as the outgroup. Red asterisk point meant the significant value (Absolute value of Z-scores larger than three or equal to three) observed in the symmetry-$f_4$ statistics and green circle point denoted the non-significant $f_4$-statistic values (Absolute value of Z-scores less than three). All Eastern Modern Tibetan/Ancient East Asians were listed along the Y-axis and $f_4$ values were labeled along the X-axis. All tested population pairs were faceted or grouped via the combination of the first and second populations. Significant negative $f_4$ values indicated that the third population shared more alleles with the second population and significant positive $f_4$ value indicated that the third population shared more derived alleles with the first population. The value of $f_4$-statistics equal to zero was marked as the blue dash line. The bar indicated three standard errors.

**Similarities and differences of the shared genetic profiles related to northern Neolithic East Asians via the spatial comparison analysis in modern Tibetans and all available ancient East Asians.**

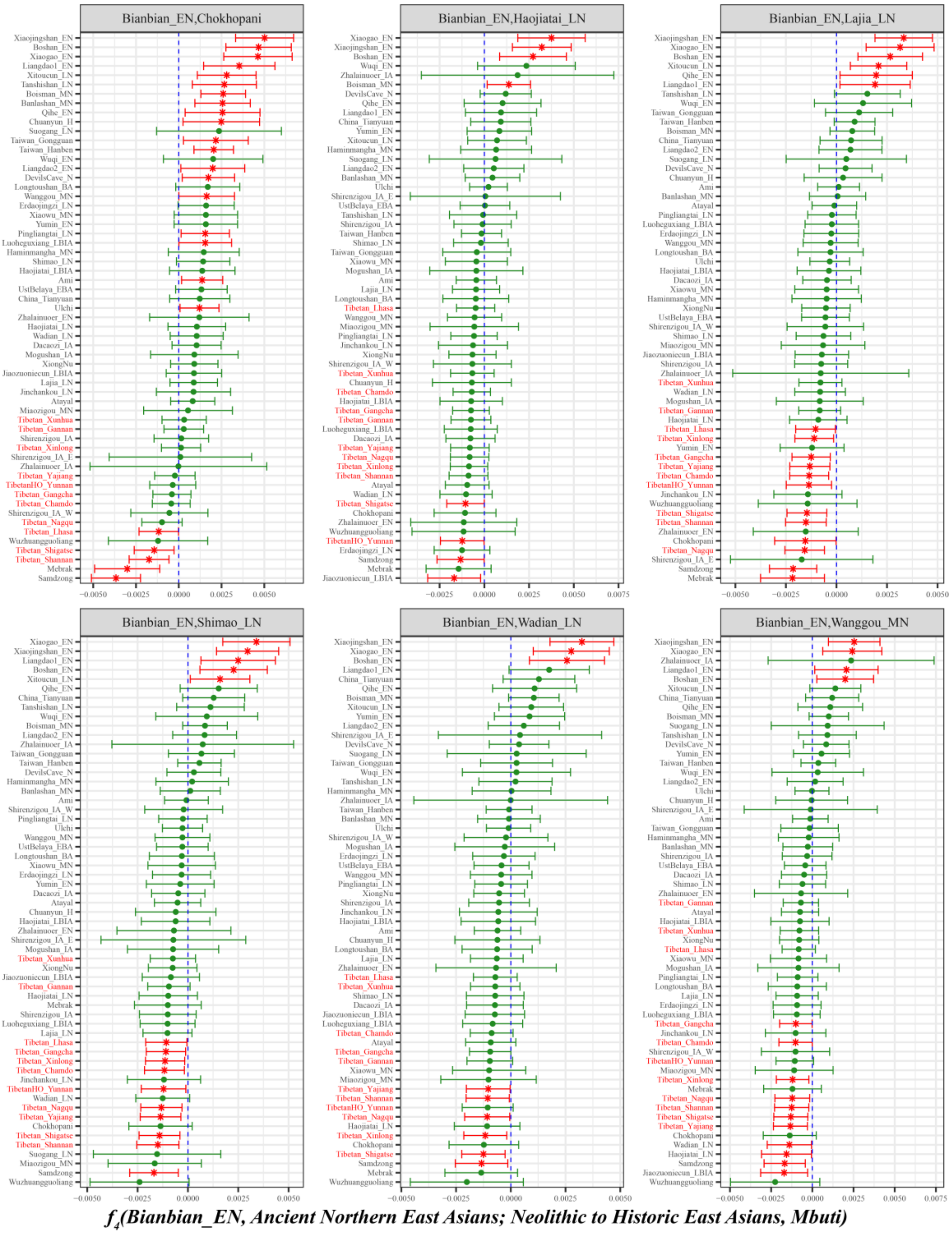

**Figure S38. Spatial comparison analysis showed the shared ancestry related to early northern Neolithic East Asians in modern Tibetans and all available ancient East Asians inferred from four population symmetry-$f_4$-statistics of the form $f_4$*(Bianbian_EN, Ancient Northern East Asians; Modern Tibetan/Neolithic to Historic East Asians, Mbuti)*.**

Here, overlapping SNP loci included in the Affymetrix Human Origins platform among four analyzed populations were used. We used the genetic variation of Mbuti as the outgroup. Red asterisk point meant the significant value (Absolute value of Z-scores larger than three or equal to three) observed in the symmetry-$f_4$ statistics and green circle point denoted the non-significant $f_4$-statistic values (Absolute value of Z-scores less than three). All Eastern Modern Tibetan/Ancient East Asians were listed along the

Y-axis and $f_4$ values were labeled along the X-axis. All tested population pairs were faceted or grouped via the combination of the first and second populations. Significant negative $f_4$ values indicated that the third population shared more alleles with the second population and significant positive $f_4$ value indicated that the third population shared more derived alleles with the first population. The value of $f_4$-statistics equal to zero was marked as the blue dash line. The bar indicated three standard errors.

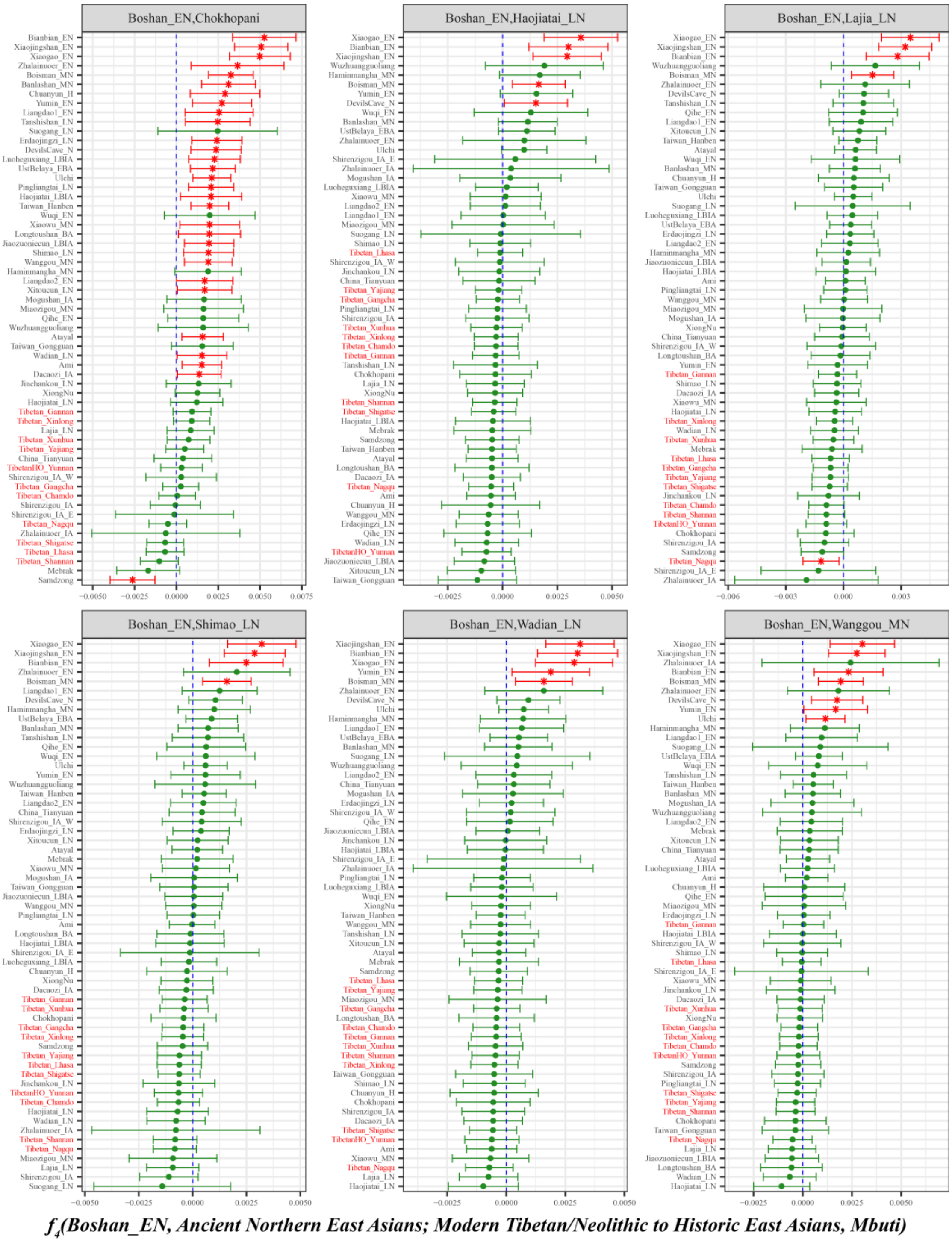

**Figure S39. Spatial comparison analysis showed the shared ancestry related to early northern Neolithic East Asians in modern Tibetans and all available ancient East Asians inferred from four population symmetry-$f_4$-statistics of the form *$f_4$(Boshan_EN, Ancient Northern East Asians; Modern Tibetan/Neolithic to Historic East Asians, Mbuti)*.**

Here, overlapping SNP loci included in the Affymetrix Human Origins platform among four analyzed populations were used. We used the genetic variation of Mbuti as the outgroup. Red asterisk point meant

the significant value (Absolute value of Z-scores larger than three or equal to three) observed in the symmetry-$f_4$ statistics and green circle point denoted the non-significant $f_4$-statistic values (Absolute value of Z-scores less than three). All Eastern Modern Tibetan/Ancient East Asians were listed along the Y-axis and $f_4$ values were labeled along the X-axis. All tested population pairs were faceted or grouped via the combination of the first and second populations. Significant negative $f_4$ values indicated that the third population shared more alleles with the second population and significant positive $f_4$ value indicated that the third population shared more derived alleles with the first population. The value of $f_4$-statistics equal to zero was marked as the blue dash line. The bar indicated three standard errors.

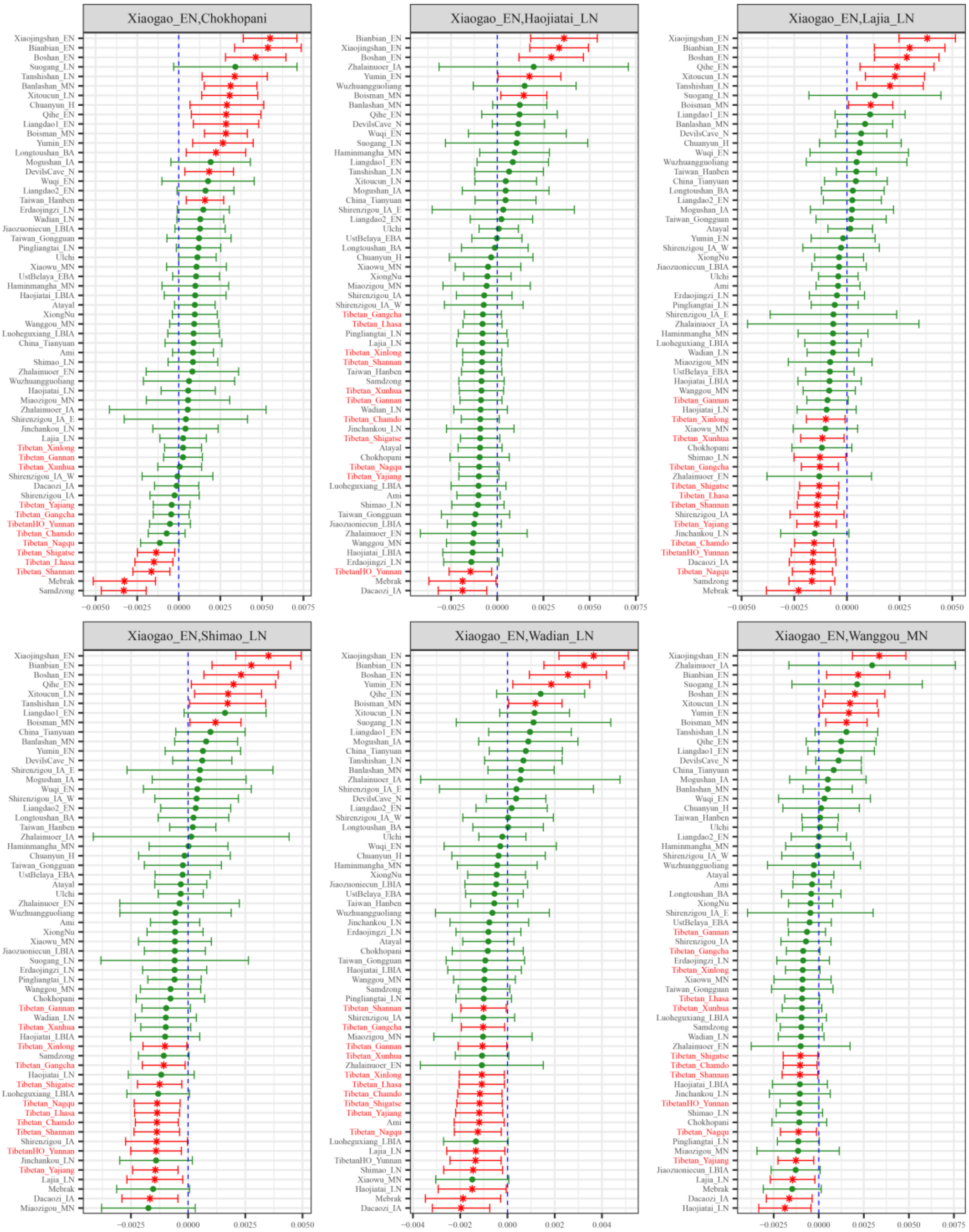

***$f_4$(Xiaogao_EN, Ancient Northern East Asians; Modern Tibetan/Neolithic to Historic East Asians, Mbuti)***

**Figure S40. Spatial comparison analysis showed the shared ancestry related to early northern Neolithic East Asians in modern Tibetans and all available ancient East Asians inferred from four population symmetry-$f_4$-statistics of the form $f_4$*(Xiaogao_EN, Ancient Northern East Asians; Modern Tibetan/Neolithic to Historic East Asians, Mbuti)*.**

Here, overlapping SNP loci included in the Affymetrix Human Origins platform among four analyzed populations were used. We used the genetic variation of Mbuti as the outgroup. Red asterisk point meant the significant value (Absolute value of Z-scores larger than three or equal to three) observed in the symmetry-$f_4$ statistics and green circle point denoted the non-significant $f_4$-statistic values (Absolute value of Z-scores less than three). All Eastern Modern Tibetan/Ancient East Asians were listed along the Y-axis and $f_4$ values were labeled along the X-axis. All tested population pairs were faceted or grouped via the combination of the first and second populations. Significant negative $f_4$ values indicated that the third population shared more alleles with the second population and significant positive $f_4$ value indicated that the third population shared more derived alleles with the first population. The value of $f_4$-statistics equal to zero was marked

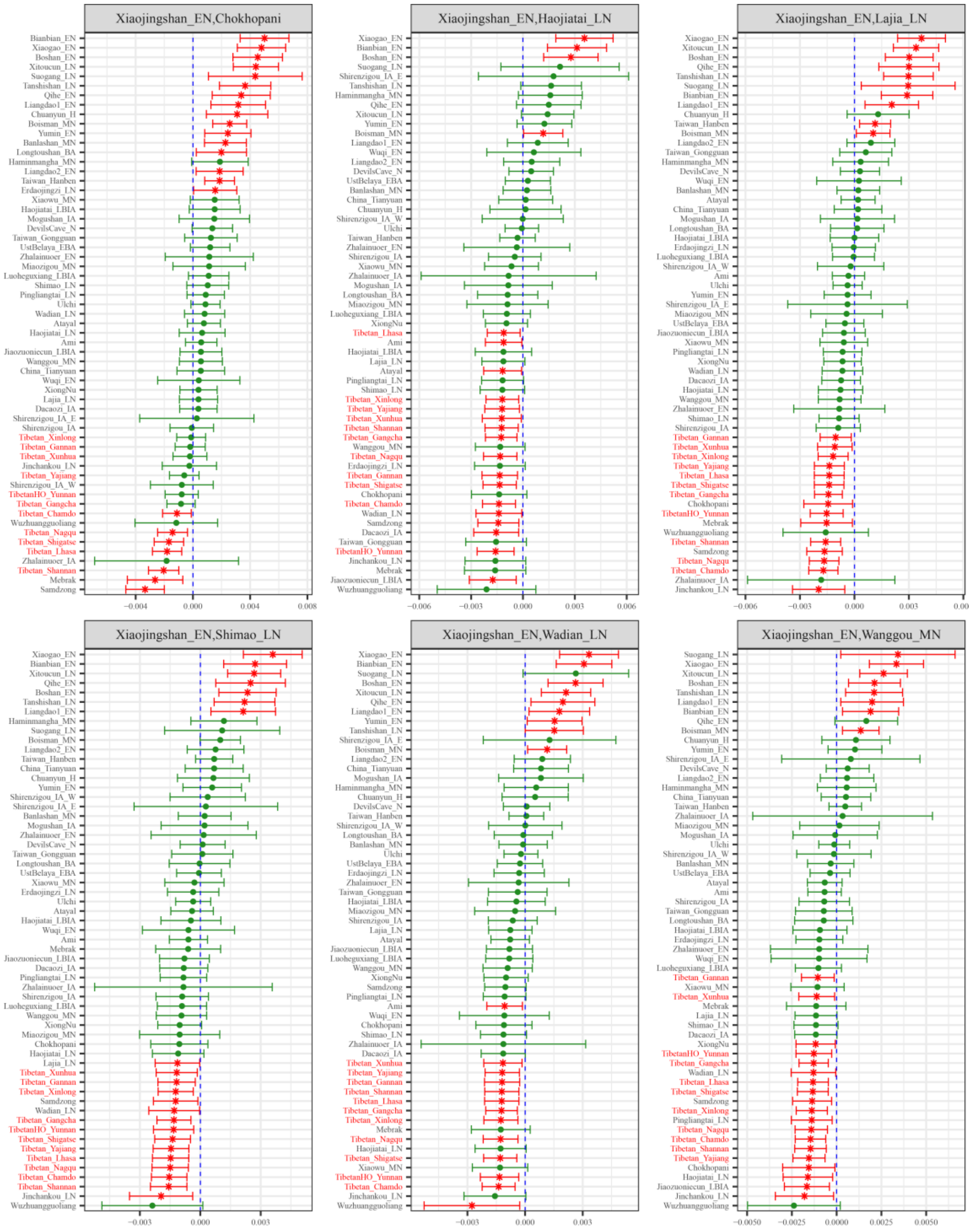

***$f_4$(Xiaojingshan_EN, Ancient Northern East Asians; Modern Tibetan/Neolithic to Historic East Asians, Mbuti)***

as the blue dash line. The bar indicated three standard errors.

**Figure S41. Spatial comparison analysis showed the shared ancestry related to early northern Neolithic East Asians in modern Tibetans and all available ancient East Asians inferred from four population symmetry-$f_4$-statistics of the form $f_4$*(Xiaojingshan_EN, Ancient Northern East Asians; Modern Tibetan/Neolithic to Historic East Asians, Mbuti)*.**

Here, overlapping SNP loci included in the Affymetrix Human Origins platform among four analyzed populations were used. We used the genetic variation of Mbuti as the outgroup. Red asterisk point meant the significant value (Absolute value of Z-scores larger than three or equal to three) observed in the symmetry-$f_4$ statistics and green circle point denoted the non-significant $f_4$-statistic values (Absolute value of Z-scores less than three). All Eastern Modern Tibetan/Ancient East Asians were listed along the Y-axis and $f_4$ values were labeled along the X-axis. All tested population pairs were faceted or grouped via the combination of the first and second populations. Significant negative $f_4$ values indicated that the third population shared more alleles with the second population and significant positive $f_4$ value indicated that the third population shared more derived alleles with the first population. The value of $f_4$-statistics equal to zero was marked as the blue dash line. The bar indicated three standard errors.

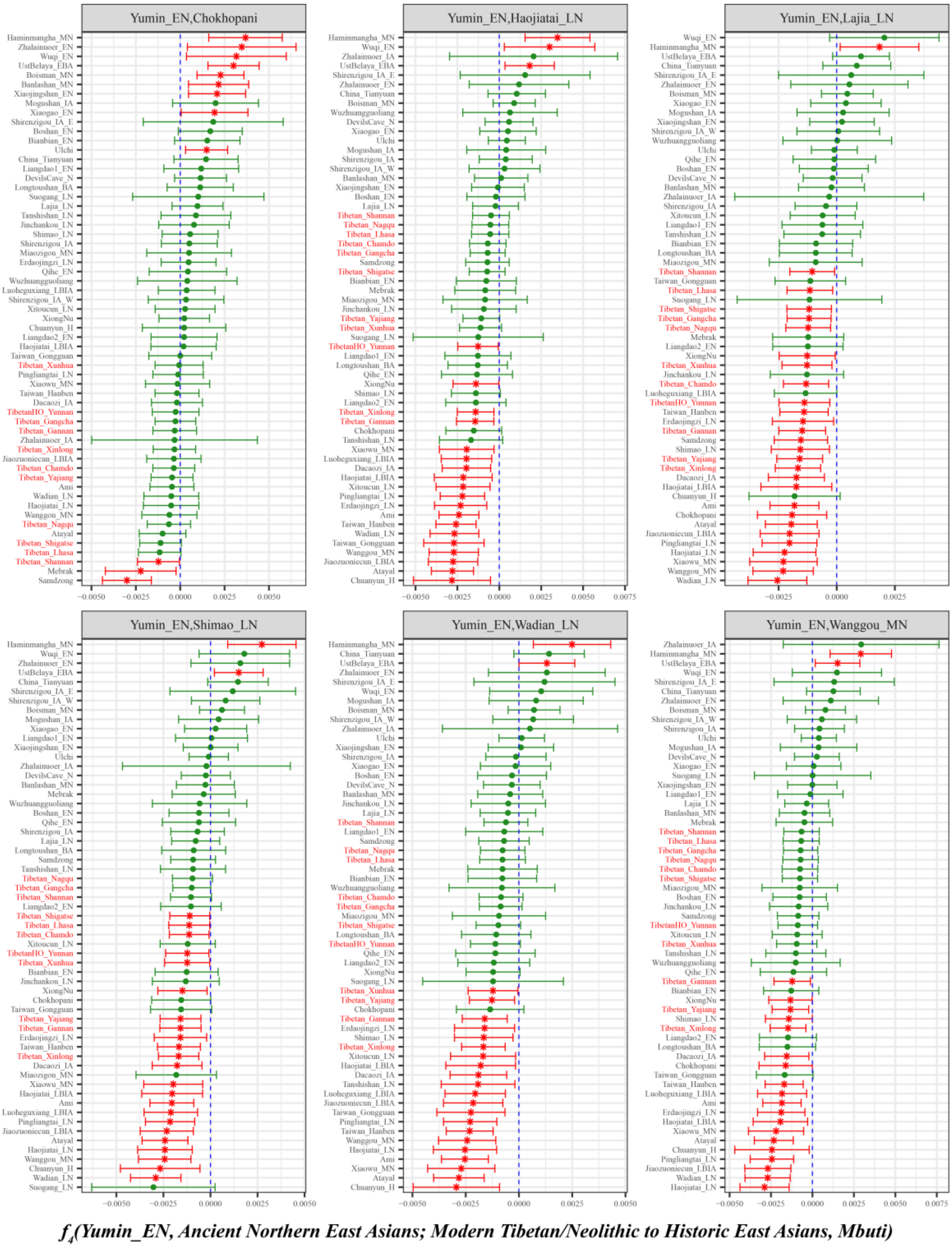

**Figure S42. Spatial comparison analysis showed the shared ancestry related to early northern Neolithic East Asians in modern Tibetans and all available ancient East Asians inferred from four population symmetry-$f_4$-statistics of the form $f_4$*(Yumin_EN, Ancient Northern East Asians; Modern Tibetan/Neolithic to Historic East Asians, Mbuti)*.**

Here, overlapping SNP loci included in the Affymetrix Human Origins platform among four analyzed populations were used. We used the genetic variation of Mbuti as the outgroup. Red asterisk point meant the significant value (Absolute value of Z-scores larger than three or equal to three) observed in the symmetry-$f_4$ statistics and green circle point denoted the non-significant $f_4$-statistic values (Absolute

value of Z-scores less than three). All Eastern Modern Tibetan/Ancient East Asians were listed along the Y-axis and $f_4$ values were labeled along the X-axis. All tested population pairs were faceted or grouped via the combination of the first and second populations. Significant negative $f_4$ values indicated that the third population shared more alleles with the second population and significant positive $f_4$ value indicated that the third population shared more derived alleles with the first population. The value of $f_4$-statistics equal to zero was marked as the blue dash line. The bar indicated three standard errors.

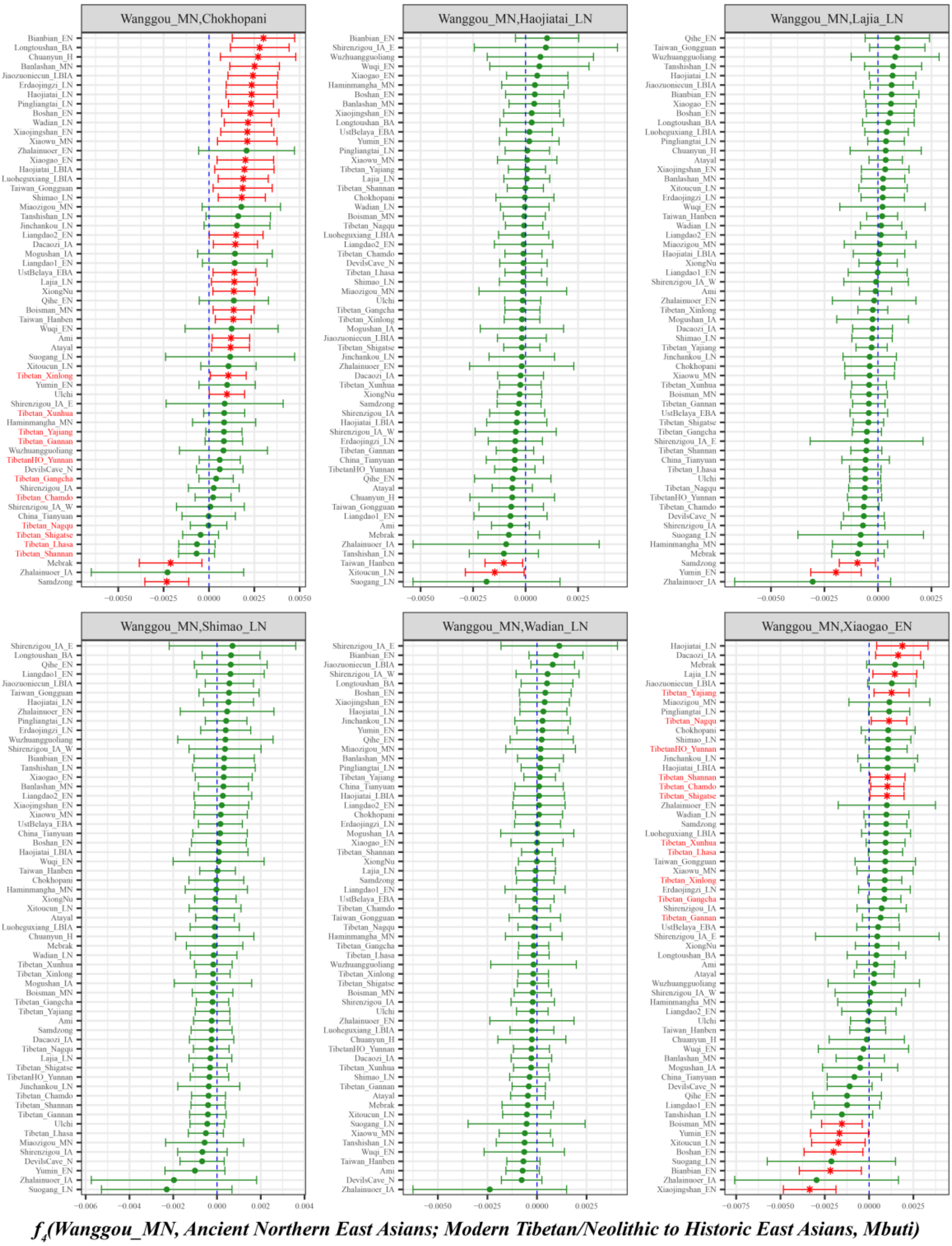

**Figure S43. Spatial comparison analysis showed the shared ancestry related to middle northern Neolithic East Asians in modern Tibetans and all available ancient East Asians inferred from four population symmetry-$f_4$-statistics of the form $f_4$*(Wanggou_MN, Ancient Northern East Asians; Modern Tibetan/Neolithic to Historic East Asians, Mbuti)*.**

Here, overlapping SNP loci included in the Affymetrix Human Origins platform among four analyzed populations were used. We used the genetic variation of Mbuti as the outgroup. Red asterisk point meant the significant value (Absolute value of Z-scores larger than three or equal to three) observed in the symmetry-$f_4$ statistics and green circle point denoted the non-significant $f_4$-statistic values (Absolute value of Z-scores less than three). All Eastern Modern Tibetan/Ancient East Asians were listed along the Y-axis and $f_4$ values were labeled along the X-axis. All tested population pairs were faceted or grouped via the combination of the first and second populations. Significant negative $f_4$ values indicated that the third population shared more alleles with the second population and significant positive $f_4$ value indicated that the third population shared more derived alleles with the first population. The value of $f_4$-statistics equal to zero was marked as the blue dash line. The bar indicated three standard errors.

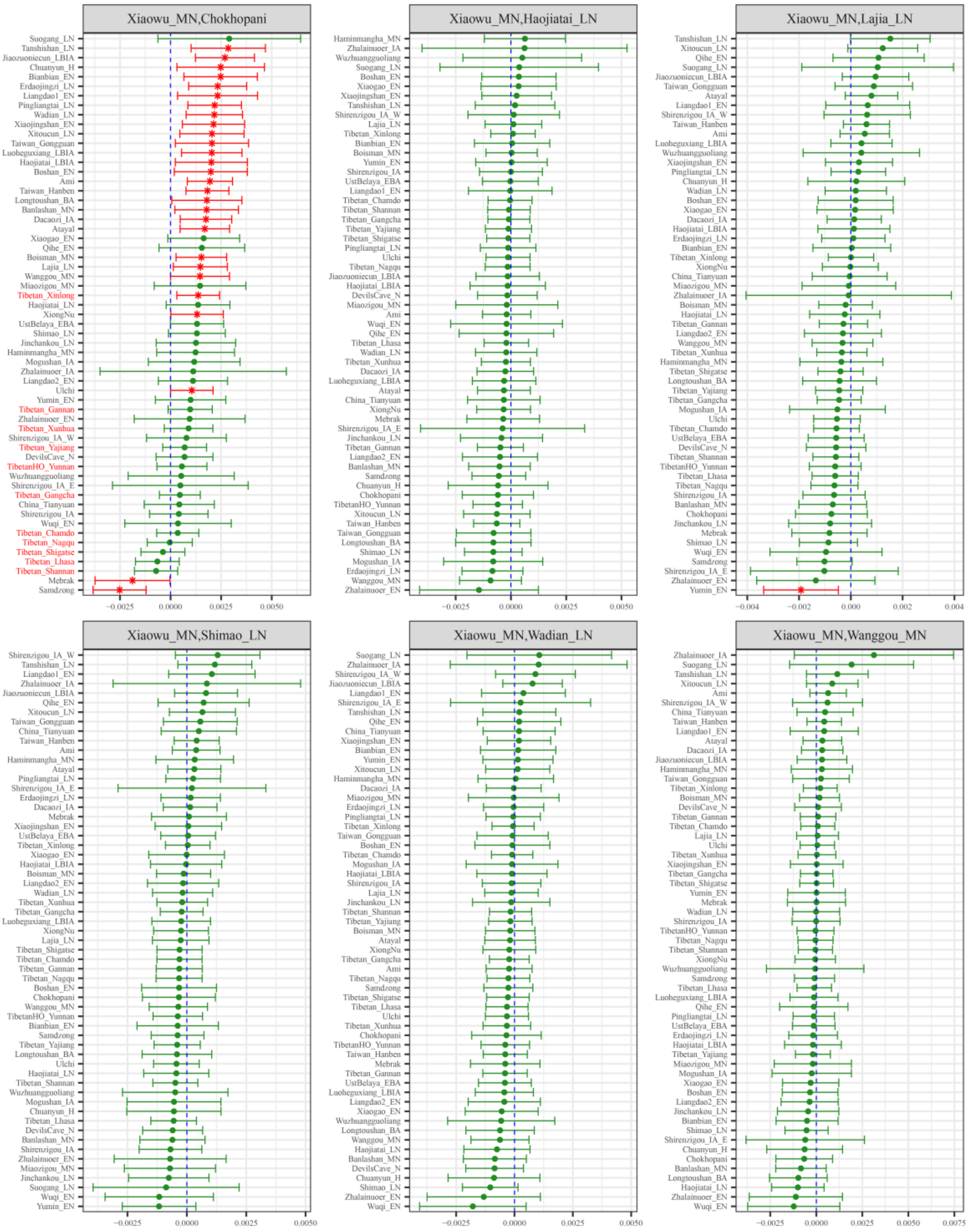

***$f_4$(Xiaowu_MN, Ancient Northern East Asians; Modern Tibetan/Neolithic to Historic East Asians, Mbuti)***

**Figure S44. Spatial comparison analysis showed the shared ancestry related to middle northern Neolithic East Asians in modern Tibetans and all available ancient East Asians inferred from four population symmetry-$f_4$-statistics of the form *$f_4$(Xiaowu_MN, Ancient Northern East Asians; Modern Tibetan/Neolithic to Historic East Asians, Mbuti)*.**

Here, overlapping SNP loci included in the Affymetrix Human Origins platform among four analyzed populations were used. We used the genetic variation of Mbuti as the outgroup. Red asterisk point meant the significant value (Absolute value of Z-scores larger than three or equal to three) observed in the symmetry-$f_4$ statistics and green circle point denoted the non-significant $f_4$-statistic values (Absolute value of Z-scores less than three). All Eastern Modern Tibetan/Ancient East Asians were listed along the Y-axis and $f_4$ values were labeled along the X-axis. All tested population pairs were faceted or grouped via the combination of the first and second populations. Significant negative $f_4$ values indicated that the third population shared more alleles with the second population and significant positive $f_4$ value indicated that the third population shared more derived alleles with the first population. The value of $f_4$-statistics equal to zero was marked as the blue dash line. The bar indicated three standard errors.

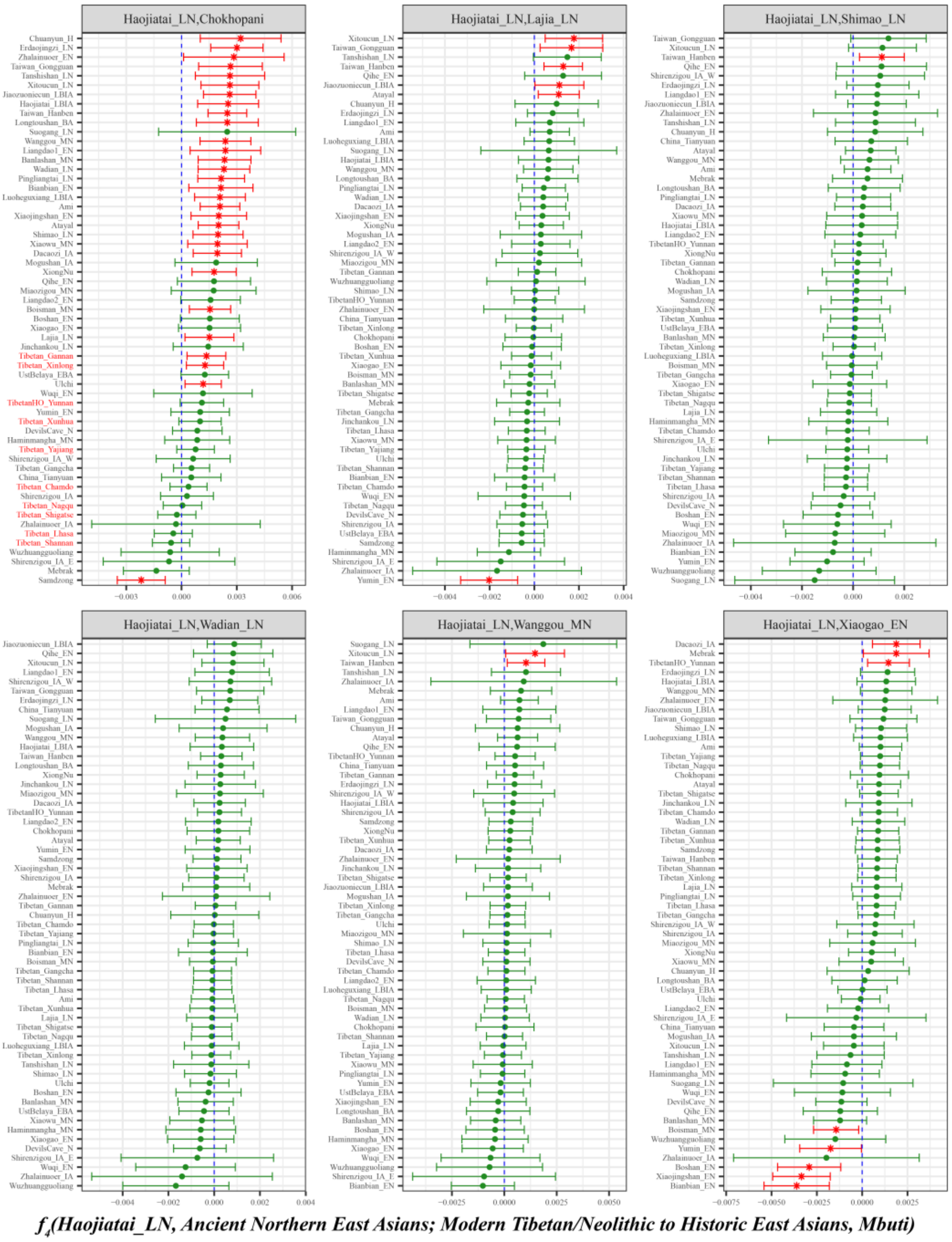

**Figure S45. Spatial comparison analysis showed the shared ancestry related to late northern Neolithic East Asians in modern Tibetans and all available ancient East Asians inferred from four population symmetry-$f_4$-statistics of the form *$f_4$(Haojiatai_LN, Ancient Northern East Asians; Modern Tibetan/Neolithic to Historic East Asians, Mbuti)*.**

Here, overlapping SNP loci included in the Affymetrix Human Origins platform among four analyzed populations were used. We used the genetic variation of Mbuti as the outgroup. Red asterisk point meant the significant value (Absolute value of Z-scores larger than three or equal to three) observed in the symmetry-$f_4$ statistics and green circle point denoted the non-significant $f_4$-statistic values (Absolute value of Z-scores less than three). All Eastern Modern Tibetan/Ancient East Asians were listed along the Y-axis and $f_4$ values were labeled along the X-axis. All tested population pairs were faceted or grouped via the combination of the first and second populations. Significant negative $f_4$ values indicated that the third population shared more alleles with the second population and significant positive $f_4$ value indicated that the third population shared more derived alleles with the first population. The value of $f_4$-statistics equal to zero was marked as the blue dash line. The bar indicated three standard errors.

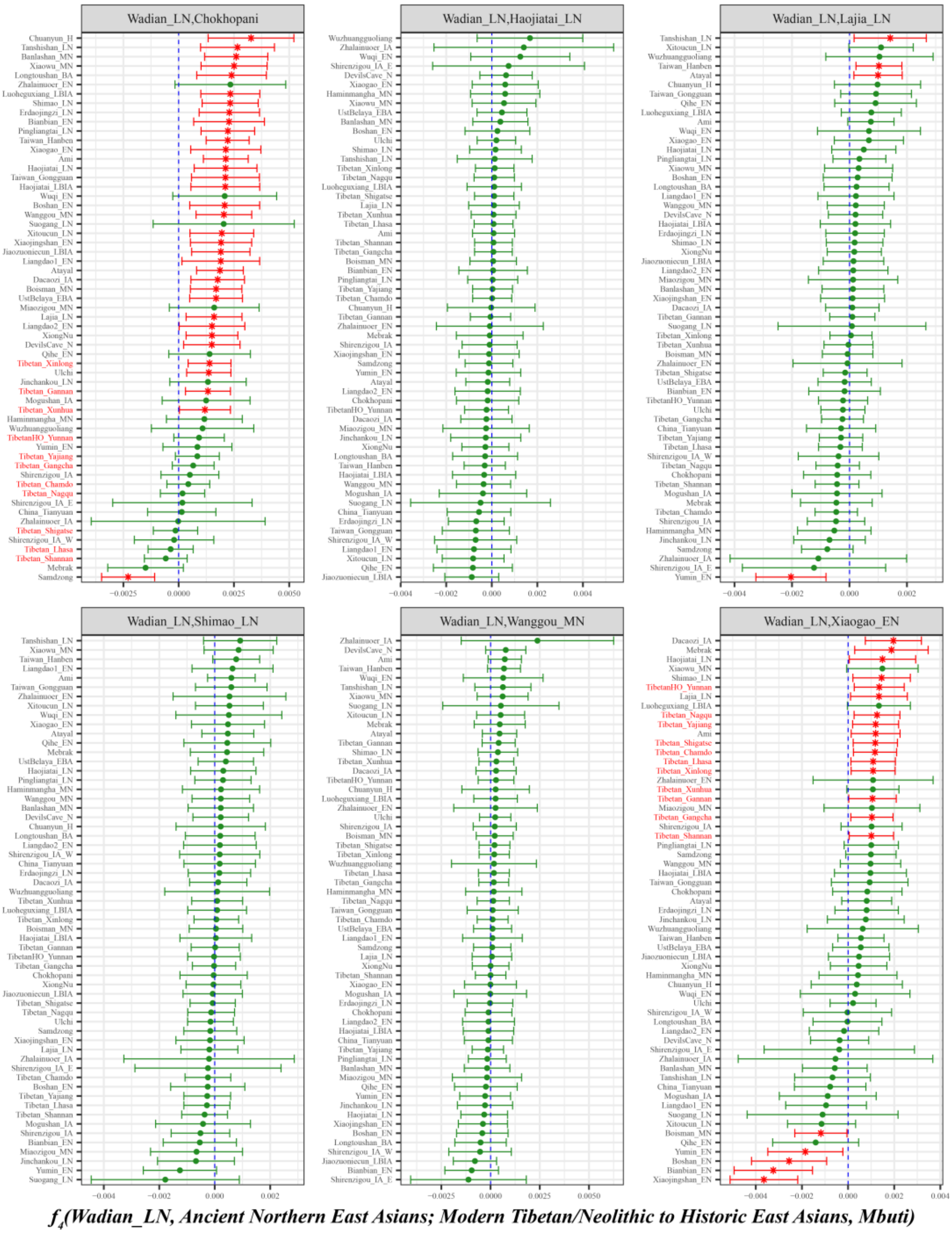

**Figure S46. Spatial comparison analysis showed the shared ancestry related to late northern Neolithic East Asians in modern Tibetans and all available ancient East Asians inferred from four population symmetry-$f_4$-statistics of the form $f_4$*(Wadian_LN, Ancient Northern East Asians; Modern Tibetan/Neolithic to Historic East Asians, Mbuti)*.**
Here, overlapping SNP loci included in the Affymetrix Human Origins platform among four analyzed populations were used. We used the genetic variation of Mbuti as the outgroup. Red asterisk point meant the significant value (Absolute value of Z-scores larger than three or equal to three) observed in the symmetry-$f_4$ statistics and green circle point denoted the non-significant $f_4$-statistic values (Absolute value of Z-scores less than three). All Eastern Modern Tibetan/Ancient East Asians were listed along the Y-axis and $f_4$ values were labeled along the X-axis. All tested population pairs were faceted or grouped via the combination of the first and second populations. Significant negative $f_4$ values indicated that the third population shared more alleles with the second population and significant positive $f_4$ value indicated that the third population shared more derived alleles with the first population. The value of $f_4$-statistics equal to zero was marked as the blue dash line. The bar indicated three standard errors.

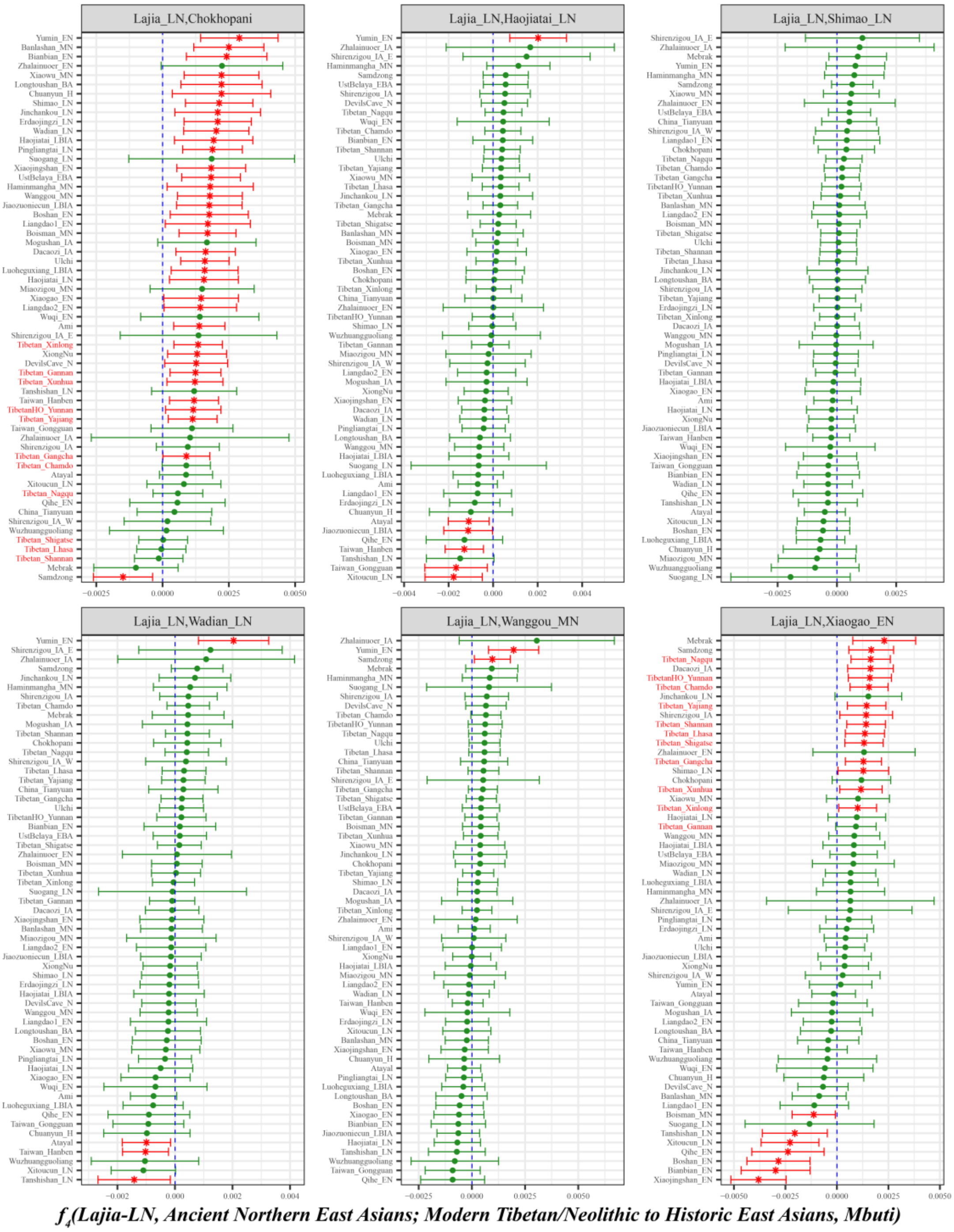

**Figure S47. Spatial comparison analysis showed the shared ancestry related to late northern Neolithic East Asians in modern Tibetans and all available ancient East Asians inferred from four population symmetry-$f_4$-statistics of the form *$f_4$(Lajia_LN, Ancient Northern East Asians; Modern Tibetan/Neolithic to Historic East Asians, Mbuti)*.**

Here, overlapping SNP loci included in the Affymetrix Human Origins platform among four analyzed populations were used. We used the genetic variation of Mbuti as the outgroup. Red asterisk point meant the significant value (Absolute value of Z-scores larger than three or equal to three) observed in the symmetry-$f_4$ statistics and green circle point denoted the non-significant $f_4$-statistic values (Absolute value of Z-scores less than three). All Eastern Modern Tibetan/Ancient East Asians were listed along the Y-axis and $f_4$ values were labeled along the X-axis. All tested population pairs were faceted or grouped via the combination of the first and second populations. Significant negative $f_4$ values indicated that the third population shared more alleles with the second population and significant positive $f_4$ value indicated that the third population shared more derived alleles with the first population. The value of $f_4$-statistics equal to zero was marked as the blue dash line. The bar indicated three standard errors.

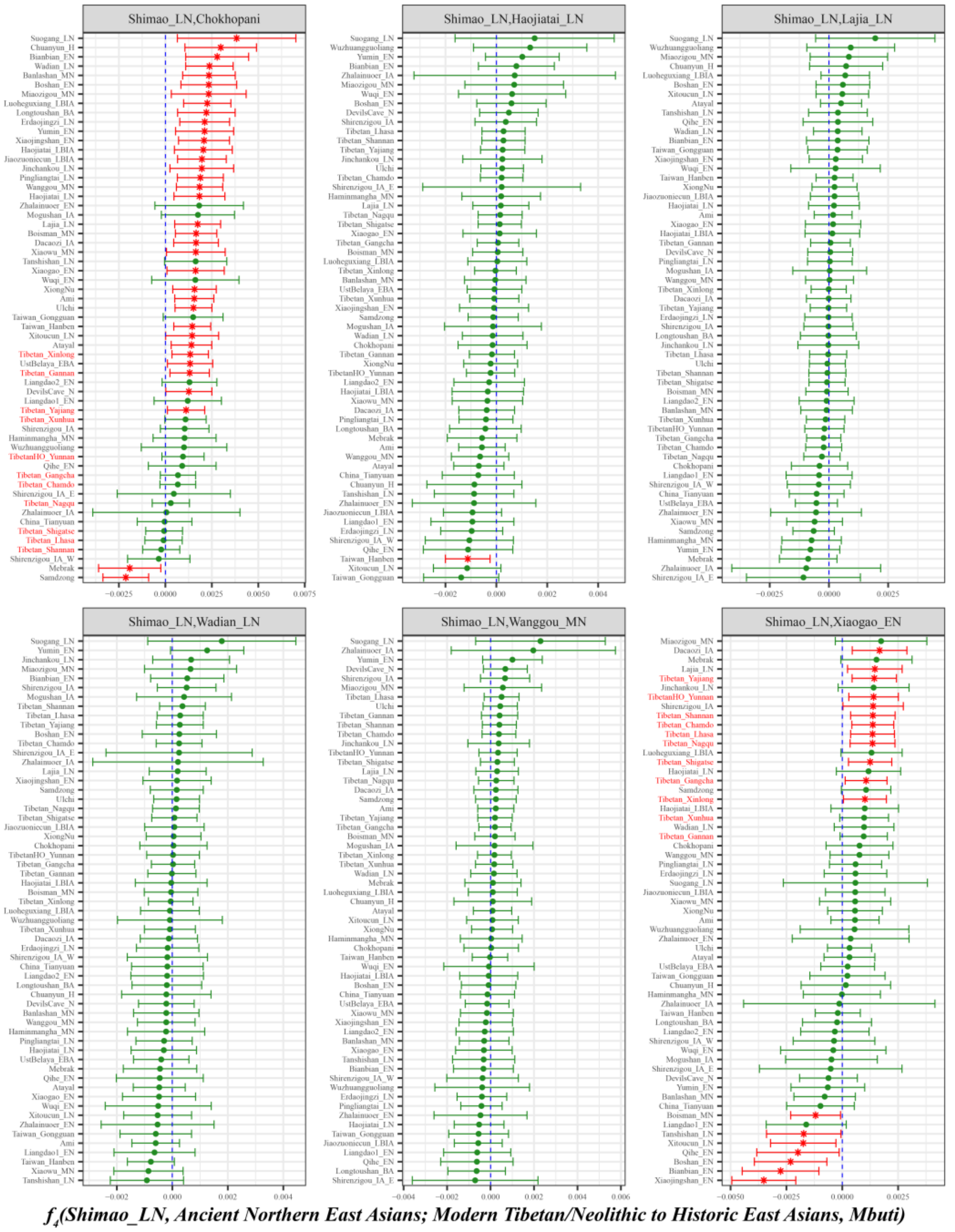

Figure S48. Spatial comparison analysis showed the shared ancestry related to late northern Neolithic East Asians in modern Tibetans and all available ancient East Asians inferred from four population symmetry-$f_4$-statistics of the form $f_4$*(Shimao_LN, Ancient Northern East Asians; Modern Tibetan/Neolithic to Historic East Asians, Mbuti)*.

Here, overlapping SNP loci included in the Affymetrix Human Origins platform among four analyzed populations were used. We used the genetic variation of Mbuti as the outgroup. Red asterisk point meant the significant value (Absolute value of Z-scores larger than three or equal to three) observed in the symmetry-$f_4$ statistics and green circle point denoted the non-significant $f_4$-statistic values (Absolute value of Z-scores less than three). All Eastern Modern Tibetan/Ancient East Asians were listed along the Y-axis and $f_4$ values were labeled along the X-axis. All tested population pairs were faceted or grouped via the combination of the first and second populations. Significant negative $f_4$ values indicated that the third population shared more alleles with the second population and significant positive $f_4$ value indicated that the third population shared more derived alleles with the first population. The value of $f_4$-statistics equal to zero was marked as the blue dash line. The bar indicated three standard errors.

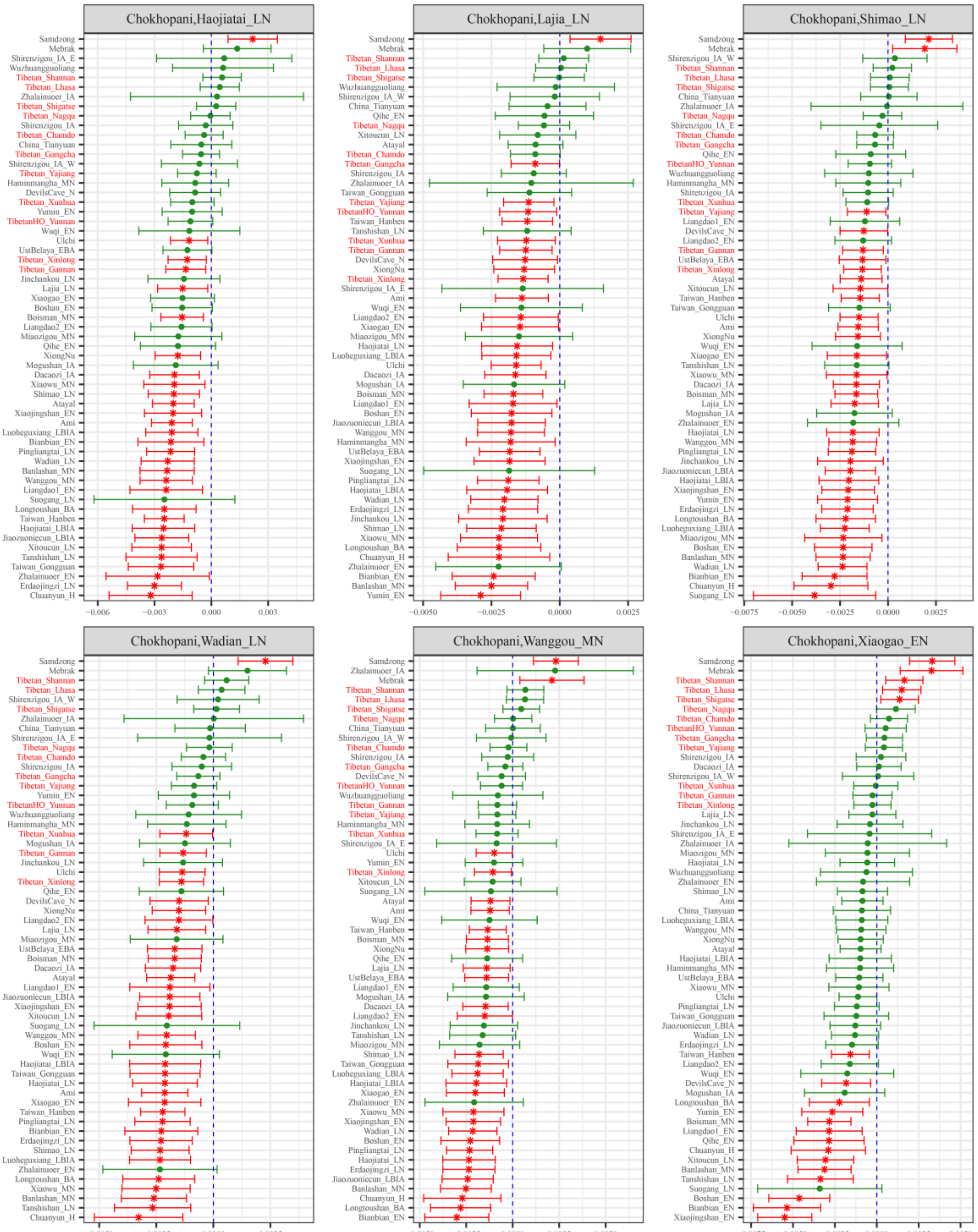

***$f_4$(Chokhopani, Ancient Northern East Asians; Modern Tibetan/Neolithic to Historic East Asians, Mbuti)***

**Figure S49. Spatial comparison analysis showed the shared ancestry related to the Ancient Nepal population in modern Tibetans and all available ancient East Asians inferred from four population symmetry-$f_4$-statistics of the form $f_4$*(Chokhopani, Ancient Northern East Asians; Modern Tibetan/Neolithic to Historic East Asians, Mbuti)*.**

Here, overlapping SNP loci included in the Affymetrix Human Origins platform among four analyzed populations were used. We used the genetic variation of Mbuti as the outgroup. Red asterisk point meant the significant value (Absolute value of Z-scores larger than three or equal to three) observed in the symmetry-$f_4$ statistics and green circle point denoted the non-significant $f_4$-statistic values (Absolute value of Z-scores less than three). All Eastern Modern Tibetan/Ancient East Asians were listed along the Y-axis and $f_4$ values were labeled along the X-axis. All tested population pairs were faceted or grouped via the combination of the first and second populations. Significant negative $f_4$ values indicated that the third population shared more alleles with the second population and significant positive $f_4$ value indicated that the third population shared more derived alleles with the first population. The value of $f_4$-statistics equal to zero was marked as the blue dash line. The bar indicated three standard errors.
