## Supplementary Figures S49-107 for "Peopling of Tibet Plateau and multiple waves of admixture of Tibetans inferred from both modern and ancient genome-wide data": Supplementary Figures S49-107.pdf

**Supplementary Figures S50-S107**

### Peopling of Tibet Plateau and multiple waves of admixture of Tibetans inferred from both modern and ancient genome-wide data

Mengge Wang[1,*], Xing Zou[1,*], Hui-Yuan Ye[2,*], Zheng Wang[1], Yan Liu[3], Jing Liu[1], Fei Wang[1], Hongbin Yao[4], Pengyu Chen[5], Ruiyang Tao[1], Shouyu Wang[1], Lan-Hai Wei[6], Renkuan Tang[7,#], Chuan-Chao Wang[6,#] , Guanglin He[1,6,#]

[1]Institute of Forensic Medicine, West China School of Basic Science and Forensic Medicine, Sichuan University, Chengdu, China
[2]School of Humanities, Nanyang Technological University, Nanyang, 639798, Singapore
[3]College of Basic Medicine, Chuanbei Medical University
[4]Belt and Road Research Center for Forensic Molecular Anthropology, Key Laboratory of Evidence Science of Gansu Province, Gansu University of Political Science and Law, Lanzhou 730070, China
[5]Center of Forensic Expertise, Affiliated hospital of Zunyi Medical University, Zunyi, Guizhou, China
[6]Department of Anthropology and Ethnology, Institute of Anthropology, National Institute for Data Science in Health and Medicine, and School of Life Sciences, Xiamen University, Xiamen, China
[7]Department of Forensic Medicine, College of Basic Medicine, Chongqing Medical University, Chongqing, China

*These authors contributed equally to this work and should be considered co-first authors.

#Corresponding author
Renkuan Tang
Department of Forensic Medicine, College of Basic Medicine, Chongqing Medical University, Chongqing, China

Chuan-Chao Wang
Affiliation: Department of Anthropology and Ethnology, Institute of Anthropology, National Institute for Data Science in Health and Medicine, Xiamen University,
Xiamen, China.

Guanglin He
Affiliation: Department of Anthropology and Ethnology, Institute of Anthropology, National Institute for Data Science in Health and Medicine, Xiamen University,
Xiamen, China.


**Contents of Supplementary Figures S50-S107**

**Additional gene flow events when we assumed that Tibetans' direct ancestor is Henan**

**Neolithic to Iron Age East Asian related ancestral populations** 15
**Figure S63**. Results of affinity-$f_4$ statistics for the form *$f_4$(Xiaowu_LN, Modern Tibetan; Neolithic to Historic East Asians, Mbuti)* showed the additional shared derived alleles from source populations except for Henan ancient population or their related ancestral populations. 15
**Figure S64**. Results of affinity-$f_4$ statistics for the form *$f_4$(Wanggou_LN, Modern Tibetan; Neolithic to Historic East Asians, Mbuti)* showed the additional shared derived alleles from source populations except for Henan ancient population or their related ancestral populations. 16
**Figure S65**. Results of affinity-$f_4$ statistics for the form *$f_4$(Wadian_LN, Modern Tibetan; Neolithic to Historic East Asians, Mbuti)* showed the additional shared derived alleles from source populations except for Henan ancient population or their related ancestral populations. 17
**Figure S66**. Results of affinity-$f_4$ statistics for the form *$f_4$(Wadian_LN, Modern Tibetan; Neolithic to Historic East Asians, Mbuti)* showed the additional shared derived alleles from source populations except for Henan ancient population or their related ancestral populations. 18
**Figure S67**. Results of affinity-$f_4$ statistics for the form *$f_4$(Pingliangtai_LN, Modern Tibetan; Neolithic to Historic East Asians, Mbuti)* showed the additional shared derived alleles from source populations except for Henan ancient population or their related ancestral populations. 19
**Figure S68**. Results of affinity-$f_4$ statistics for the form *$f_4$(Haojiatai_LN, Modern Tibetan; Neolithic to Historic East Asians, Mbuti)* showed the additional shared derived alleles from source populations except for Henan ancient population or their related ancestral populations. 20
**Figure S69**. Results of affinity-$f_4$ statistics for the form *$f_4$(Haojiatai_LBIA, Modern Tibetan; Neolithic to Historic East Asians, Mbuti)* showed the additional shared derived alleles from source populations except for Henan ancient population or their related ancestral populations. 21
**Figure S70**. Results of affinity-$f_4$ statistics for the form *$f_4$(Jiaozuoniecun_LBIA, Modern Tibetan; Neolithic to Historic East Asians, Mbuti)* showed the additional shared derived alleles from source populations except for Henan ancient population or their related ancestral populations. 22
**Figure S71**. Results of affinity-$f_4$ statistics for the form *$f_4$(Luoheguxiang_LBIA, Modern Tibetan; Neolithic to Historic East Asians, Mbuti)* showed the additional shared derived alleles from source populations except for Henan ancient population or their related ancestral populations. 23
Additional gene flow events when we assumed that Tibetans' direct ancestor is inland Neolithic northern East Asian from Shaanxi or Inner Mongolia or their related ancestral populations 25
**Figure S72**. Results of affinity-$f_4$ statistics for the form *$f_4$(Wuzhuangguoliang, Modern Tibetan; Neolithic to Historic East Asians, Mbuti)* showed the additional shared derived alleles from source populations except for inland Neolithic northern East Asian from Shaanxi or Inner Mongolia or their related ancestral populations. 25
**Figure S73**. Results of affinity-$f_4$ statistics for the form *$f_4$(Shimao_LN, Modern Tibetan; Neolithic to Historic East Asians, Mbuti)* showed the additional shared derived alleles from source populations except for inland Neolithic northern East Asian from Shaanxi or Inner Mongolia or their related ancestral populations. 26
**Figure S74**. Results of affinity-$f_4$ statistics for the form *$f_4$(Miaozigou_MN, Modern Tibetan; Neolithic to Historic East Asians, Mbuti)* showed the additional shared derived alleles from source populations except for inland Neolithic northern East Asian from Shaanxi or Inner Mongolia or their related ancestral populations. 27
**Figure S75**. Results of affinity-$f_4$ statistics for the form *$f_4$(Yumin_EN, Modern Tibetan; Neolithic to Historic East Asians, Mbuti)* showed the additional shared derived alleles from source populations

except for inland Neolithic northern East Asian from Shaanxi or Inner Mongolia or their related ancestral populations. 28
**Additional gene flow events when we assumed that Tibetans' direct ancestor is inland Neolithic to Iron Age northern East Asian from upper Yellow River Basin or their related ancestral populations** 30
**Figure S76**. Results of affinity-$f_4$ statistics for the form *$f_4$(Lajia_LN, Modern Tibetan; Neolithic to Historic East Asians, Mbuti)* showed the additional shared derived alleles from source populations except for inland Neolithic northern East Asian from Shaanxi or Inner Mongolia or their related ancestral populations. 30
**Figure S77**. Results of affinity-$f_4$ statistics for the form *$f_4$(Jinchankou_LN, Modern Tibetan; Neolithic to Historic East Asians, Mbuti)* showed the additional shared derived alleles from source populations except for inland Neolithic northern East Asian from Shaanxi or Inner Mongolia or their related ancestral populations. 31
**Figure S78**. Results of affinity-$f_4$ statistics for the form *$f_4$(Dacaozi_IA, Modern Tibetan; Neolithic to Historic East Asians, Mbuti)* showed the additional shared derived alleles from source populations except for inland Neolithic northern East Asian from Shaanxi or Inner Mongolia or their related ancestral populations. 32
**Additional gene flow events when we assumed that Tibetans' direct ancestor is Liao River ancients or their related ancestral populations** 33
**Figure S79**. Results of affinity-$f_4$ statistics for the form *$f_4$(Banhashan_MN, Modern Tibetan; Neolithic to Historic East Asians, Mbuti)* showed the additional shared derived alleles from source populations except for ancestral populations from Liao River Basin. 33
**Figure S80**. Results of affinity-$f_4$ statistics for the form *$f_4$(Haminmangha_MN, Modern Tibetan; Neolithic to Historic East Asians, Mbuti)* showed the additional shared derived alleles from source populations except for ancestral populations from Liao River Basin. 34
**Figure S81**. Results of affinity-$f_4$ statistics for the form *$f_4$(Erdaojingzi_LN, Modern Tibetan; Neolithic to Historic East Asians, Mbuti)* showed the additional shared derived alleles from source populations except for ancestral populations from Liao River Basin. 35
**Figure S82**. Results of affinity-$f_4$ statistics for the form *$f_4$(Longtoushan_BA, Modern Tibetan; Neolithic to Historic East Asians, Mbuti)* showed the additional shared derived alleles from source populations except for ancestral populations from Liao River Basin. 36
**Additional gene flow events when we assumed that Tibetans' direct ancestor is ancestral populations from Amur River Basin** 38
**Figure S83**. Results of affinity-$f_4$ statistics for the form *$f_4$(Wuqi_EN, Modern Tibetan; Neolithic to Historic East Asians, Mbuti)* showed the additional shared derived alleles from source populations except for ancestral populations from Amur River Basin. 38
**Figure S84**. Results of affinity-$f_4$ statistics for the form *$f_4$(Zhalainuoer_EN, Modern Tibetan; Neolithic to Historic East Asians, Mbuti)* showed the additional shared derived alleles from source populations except for ancestral populations from Amur River Basin. 39
**Figure S85**. Results of affinity-$f_4$ statistics for the form *$f_4$(Boisman_MN, Modern Tibetan; Neolithic to Historic East Asians, Mbuti)* showed the additional shared derived alleles from source populations except for ancestral populations from Amur River Basin. 40
**Figure S86**. Results of affinity-$f_4$ statistics for the form *$f_4$(DevilsCave_N, Modern Tibetan; Neolithic to Historic East Asians, Mbuti)* showed the additional shared derived alleles from source

populations except for ancestral populations from Amur River Basin. 41
**Figure S87**. Results of affinity-$f_4$ statistics for the form *$f_4$(Mogushan_IA, Modern Tibetan; Neolithic to Historic East Asians, Mbuti)* showed the additional shared derived alleles from source populations except for ancestral populations from Amur River Basin. 42
**Figure S88**. Results of affinity-$f_4$ statistics for the form *$f_4$(Ulchi, Modern Tibetan; Neolithic to Historic East Asians, Mbuti)* showed the additional shared derived alleles from source populations except for ancestral populations from Amur River Basin. 43
**Additional gene flow events when we assumed that Tibetans' direct ancestor is ancestral populations from Nepal** 44
**Figure S89**. Results of affinity-$f_4$ statistics for the form *$f_4$(Chokhopani, Modern Tibetan; Neolithic to Historic East Asians, Mbuti)* showed the additional shared derived alleles from source populations except for ancestral populations from Nepal. 44
**Figure S90**. Results of affinity-$f_4$ statistics for the form *$f_4$(Mebrak, Modern Tibetan; Neolithic to Historic East Asians, Mbuti)* showed the additional shared derived alleles from source populations except for ancestral populations from Nepal. 45
**Figure S91**. Results of affinity-$f_4$ statistics for the form *$f_4$(Samdzong, Modern Tibetan; Neolithic to Historic East Asians, Mbuti)* showed the additional shared derived alleles from source populations except for ancestral populations from Nepal. 46
**Additional gene flow events when we assumed that Tibetans' direct ancestor is ancestral populations from Mongolia Plateau or Baikal Lake Region** 48
**Figure S92**. Results of affinity-$f_4$ statistics for the form *$f_4$(Mongolia_N_East, Modern Tibetan; Neolithic to Historic East Asians, Mbuti)* showed the additional shared derived alleles from source populations except for ancestral populations from Mongolia Plateau or Baikal Lake Region. 48
**Figure S93**. Results of affinity-$f_4$ statistics for the form *$f_4$(Mongolia_N_North, Modern Tibetan; Neolithic to Historic East Asians, Mbuti)* showed the additional shared derived alleles from source populations except for ancestral populations from Mongolia Plateau or Baikal Lake Region. 49
**Figure S94**. Results of affinity-$f_4$ statistics for the form *$f_4$(Russia_OldBeringSea_Ekven, Modern Tibetan; Neolithic to Historic East Asians, Mbuti)* showed the additional shared derived alleles from source populations except for ancestral populations from Mongolia Plateau or Baikal Lake Region. 50
**Figure S95**. Results of affinity-$f_4$ statistics for the form *$f_4$(Russia_Shamanka_EBA, Modern Tibetan; Neolithic to Historic East Asians, Mbuti)* showed the additional shared derived alleles from source populations except for ancestral populations from Mongolia Plateau or Baikal Lake Region. 51
**Figure S96**. Results of affinity-$f_4$ statistics for the form *$f_4$(Russia_Shamanka_Eneolithic, Modern Tibetan; Neolithic to Historic East Asians, Mbuti)* showed the additional shared derived alleles from source populations except for ancestral populations from Mongolia Plateau or Baikal Lake Region. 52
**Figure S97**. Results of affinity-$f_4$ statistics for the form *$f_4$(Russia_UstBelaya_Angara, Modern Tibetan; Neolithic to Historic East Asians, Mbuti)* showed the additional shared derived alleles from source populations except for ancestral populations from Mongolia Plateau or Baikal Lake Region. 53
**Figure S98**. Results of affinity-$f_4$ statistics for the form *$f_4$(Russia_UstIda_EBA, Modern Tibetan; Neolithic to Historic East Asians, Mbuti)* showed the additional shared derived alleles from source populations except for ancestral populations from Mongolia Plateau or Baikal Lake Region. 54

**Additional gene flow events when we assumed that Tibetans' direct ancestor is Yangtze rice farmer related ancestral populations**

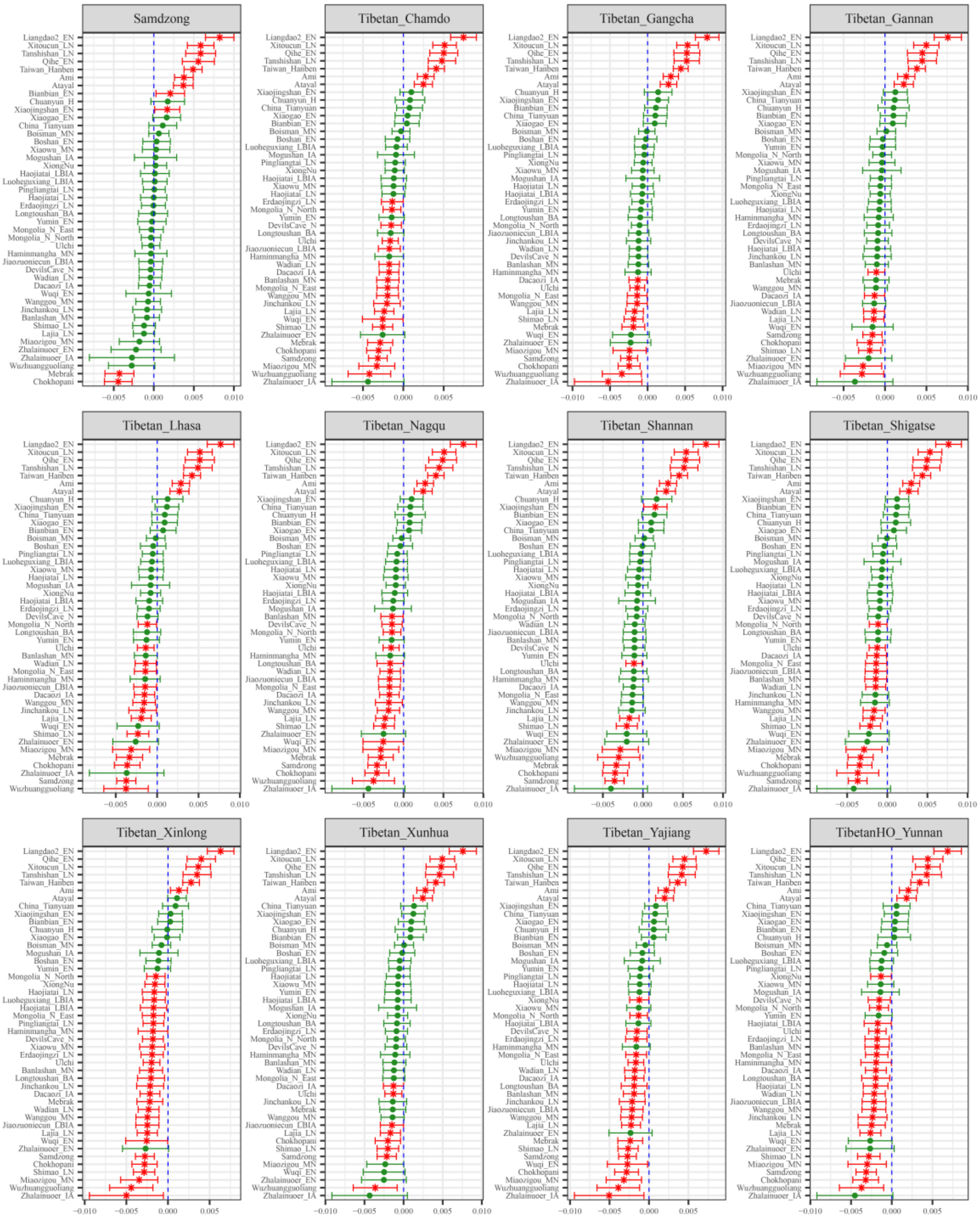

**Figure S50. Results of affinity-$f_4$ statistics for the form *$f_4$(Liaodao1_EN, Modern Tibetan; Neolithic to Historic East Asians, Mbuti)* showed the additional shared derived alleles from source populations except for Yangtze rice farmer related ancestral populations.**

Here, overlapping SNP loci included in the Affymetrix Human Origins platform among four analyzed populations were used. We used the genetic variation of Mbuti as the outgroup. Red asterisk point meant the significant value (Absolute value of Z-scores larger than three or equal to three) observed in the symmetry-$f_4$ statistics and green circle point denoted the non-significant $f_4$-statistic values (Absolute value of Z-scores less than three). All ancient East Asians were listed along the Y-axis and $f_4$ values were labeled along the X-axis. All results were faceted or grouped via Tibetan populations. Significant negative $f_4$ values indicated that the third population shared more alleles with the second population, and also means Tibetan obtained additional gene flow from the third source population (or related

populations). And the significant positive $f_4$ value indicated that the third population shared more derived alleles with the first population. The value of $f_4$-statistics equal to zero was marked as the blue dash line. The bar indicated three standard errors.

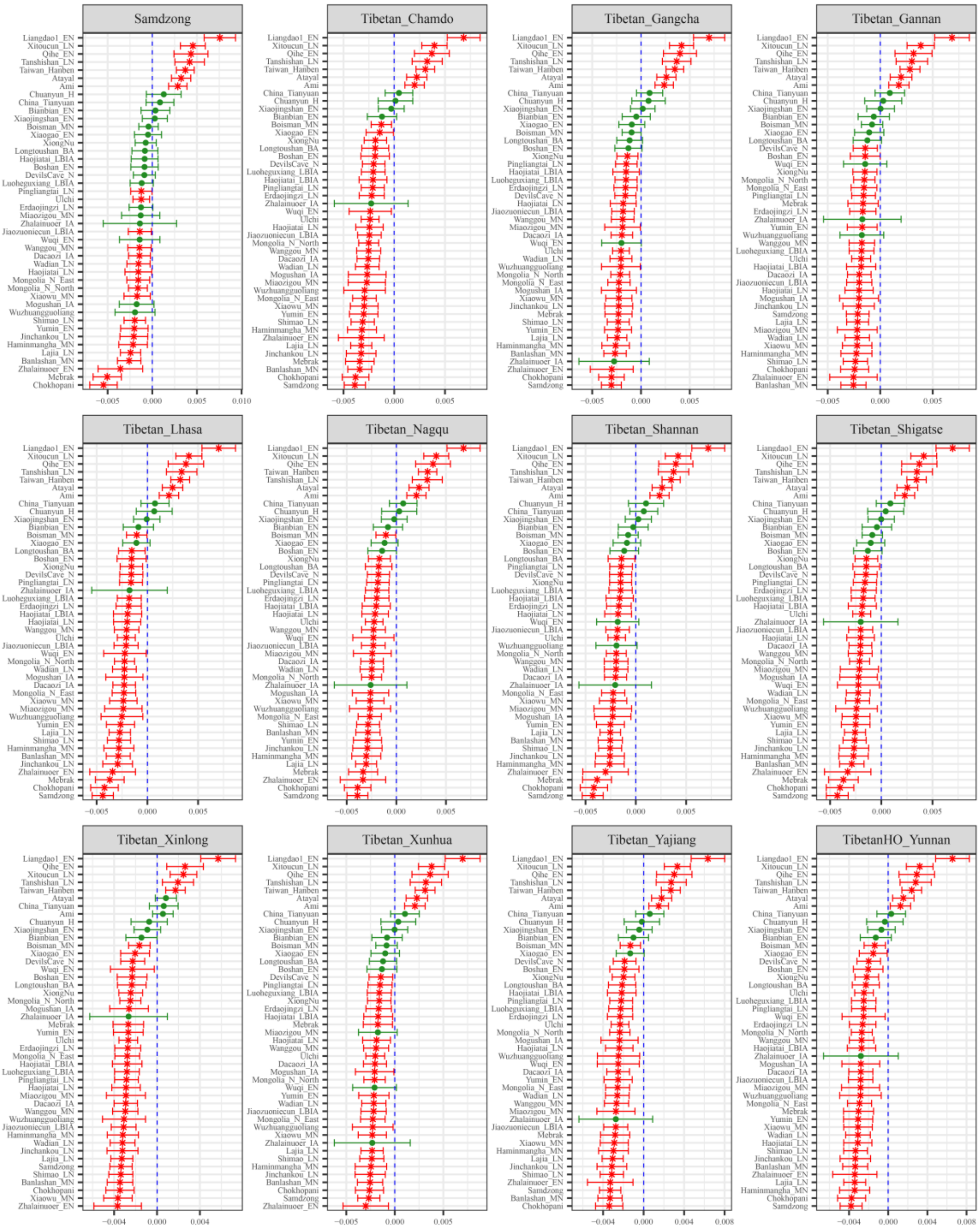

**Figure S51. Results of affinity-$f_4$ statistics for the form $f_4$*(Liaodao2_EN, Modern Tibetan; Neolithic to Historic East Asians, Mbuti)* showed the additional shared derived alleles from source populations except for Yangtze rice farmer related ancestral populations.**

Here, overlapping SNP loci included in the Affymetrix Human Origins platform among four analyzed populations were used. We used the genetic variation of Mbuti as the outgroup. Red asterisk point meant the significant value (Absolute value of Z-scores larger than three or equal to three) observed in the symmetry-$f_4$ statistics and green circle point denoted the non-significant $f_4$-statistic values (Absolute value of Z-scores less than three). All ancient East Asians were listed along the Y-axis and $f_4$ values were labeled along the X-axis. All results were faceted or grouped via Tibetan populations. Significant negative $f_4$ values indicated that the third population shared more alleles with the second population, and also means Tibetan obtained additional gene flow from the third source population (or related

populations). And the significant positive $f_4$ value indicated that the third population shared more derived alleles with the first population. The value of $f_4$-statistics equal to zero was marked as the blue dash line. The bar indicated three standard errors.

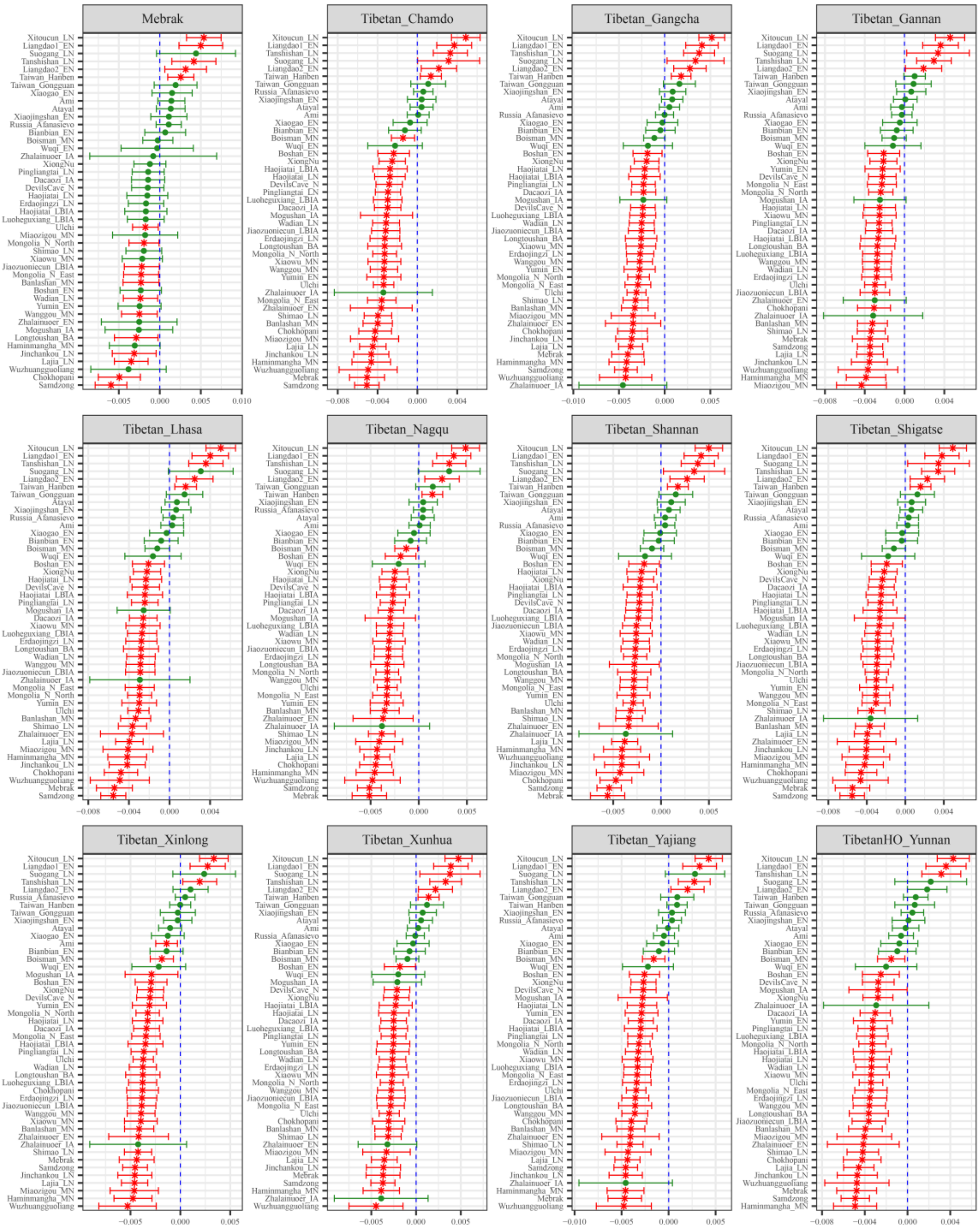

**Figure S52. Results of affinity-$f_4$ statistics for the form $f_4$*(Qihe_EN, Modern Tibetan; Neolithic to Historic East Asians, Mbuti)* showed the additional shared derived alleles from source populations except for Yangtze rice farmer related ancestral populations.**

Here, overlapping SNP loci included in the Affymetrix Human Origins platform among four analyzed populations were used. We used the genetic variation of Mbuti as the outgroup. Red asterisk point meant the significant value (Absolute value of Z-scores larger than three or equal to three) observed in the symmetry-$f_4$ statistics and green circle point denoted the non-significant $f_4$-statistic values (Absolute value of Z-scores less than three). All ancient East Asians were listed along the Y-axis and $f_4$ values were labeled along the X-axis. All results were faceted or grouped via Tibetan populations. Significant negative $f_4$ values indicated that the third population shared more alleles with the second population, and also means Tibetan obtained additional gene flow from the third source population (or related

populations). And the significant positive $f_4$ value indicated that the third population shared more derived alleles with the first population. The value of $f_4$-statistics equal to zero was marked as the blue dash line. The bar indicated three standard errors.

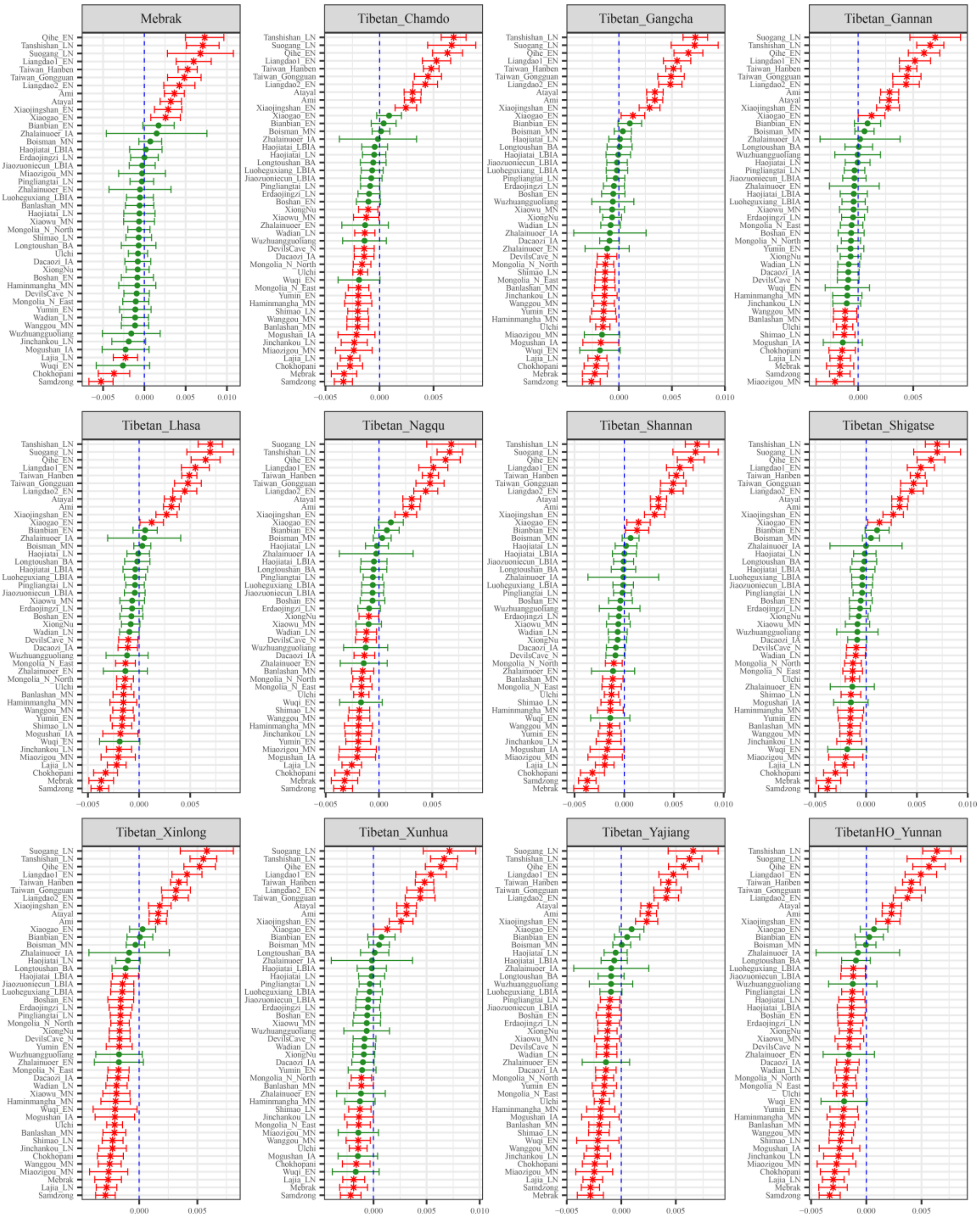

**Figure S53. Results of affinity-$f_4$ statistics for the form $f_4$*(Xitoucun_LN, Modern Tibetan; Neolithic to Historic East Asians, Mbuti)* showed the additional shared derived alleles from source populations except for Yangtze rice farmer related ancestral populations.**

Here, overlapping SNP loci included in the Affymetrix Human Origins platform among four analyzed populations were used. We used the genetic variation of Mbuti as the outgroup. Red asterisk point meant the significant value (Absolute value of Z-scores larger than three or equal to three) observed in the symmetry-$f_4$ statistics and green circle point denoted the non-significant $f_4$-statistic values (Absolute value of Z-scores less than three). All ancient East Asians were listed along the Y-axis and $f_4$ values were labeled along the X-axis. All results were faceted or grouped via Tibetan populations. Significant negative $f_4$ values indicated that the third population shared more alleles with the second population, and also means Tibetan obtained additional gene flow from the third source population (or related

populations). And the significant positive $f_4$ value indicated that the third population shared more derived alleles with the first population. The value of $f_4$-statistics equal to zero was marked as the blue dash line. The bar indicated three standard errors.

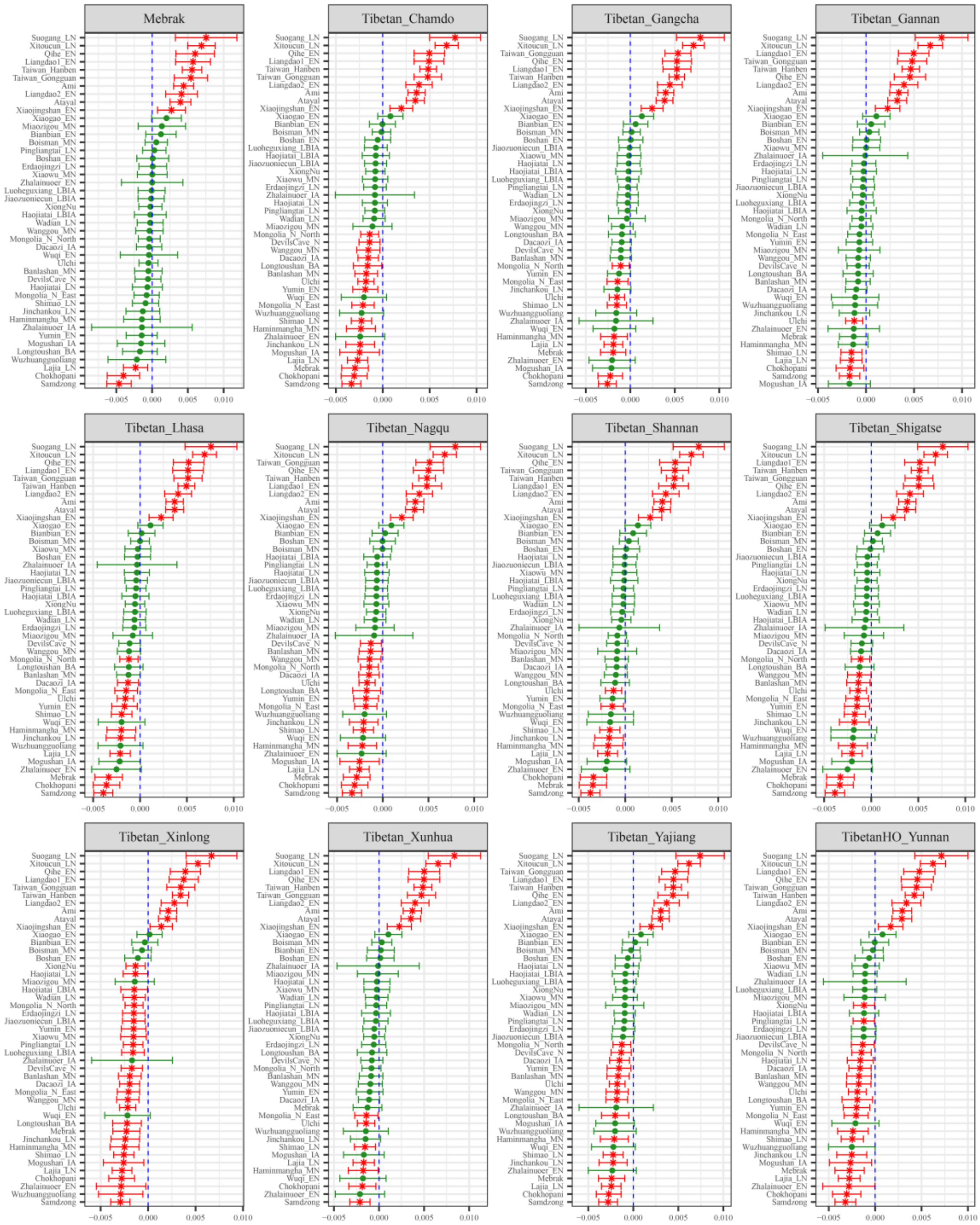

**Figure S54. Results of affinity-$f_4$ statistics for the form $f_4$*(Tanshishan_LN, Modern Tibetan; Neolithic to Historic East Asians, Mbuti)* showed the additional shared derived alleles from source populations except for Yangtze rice farmer related ancestral populations.**

Here, overlapping SNP loci included in the Affymetrix Human Origins platform among four analyzed populations were used. We used the genetic variation of Mbuti as the outgroup. Red asterisk point meant the significant value (Absolute value of Z-scores larger than three or equal to three) observed in the symmetry-$f_4$ statistics and green circle point denoted the non-significant $f_4$-statistic values (Absolute value of Z-scores less than three). All ancient East Asians were listed along the Y-axis and $f_4$ values were labeled along the X-axis. All results were faceted or grouped via Tibetan populations. Significant negative $f_4$ values indicated that the third population shared more alleles with the second population, and also means Tibetan obtained additional gene flow from the third source population (or related

populations). And the significant positive $f_4$ value indicated that the third population shared more derived alleles with the first population. The value of $f_4$-statistics equal to zero was marked as the blue dash line. The bar indicated three standard errors.

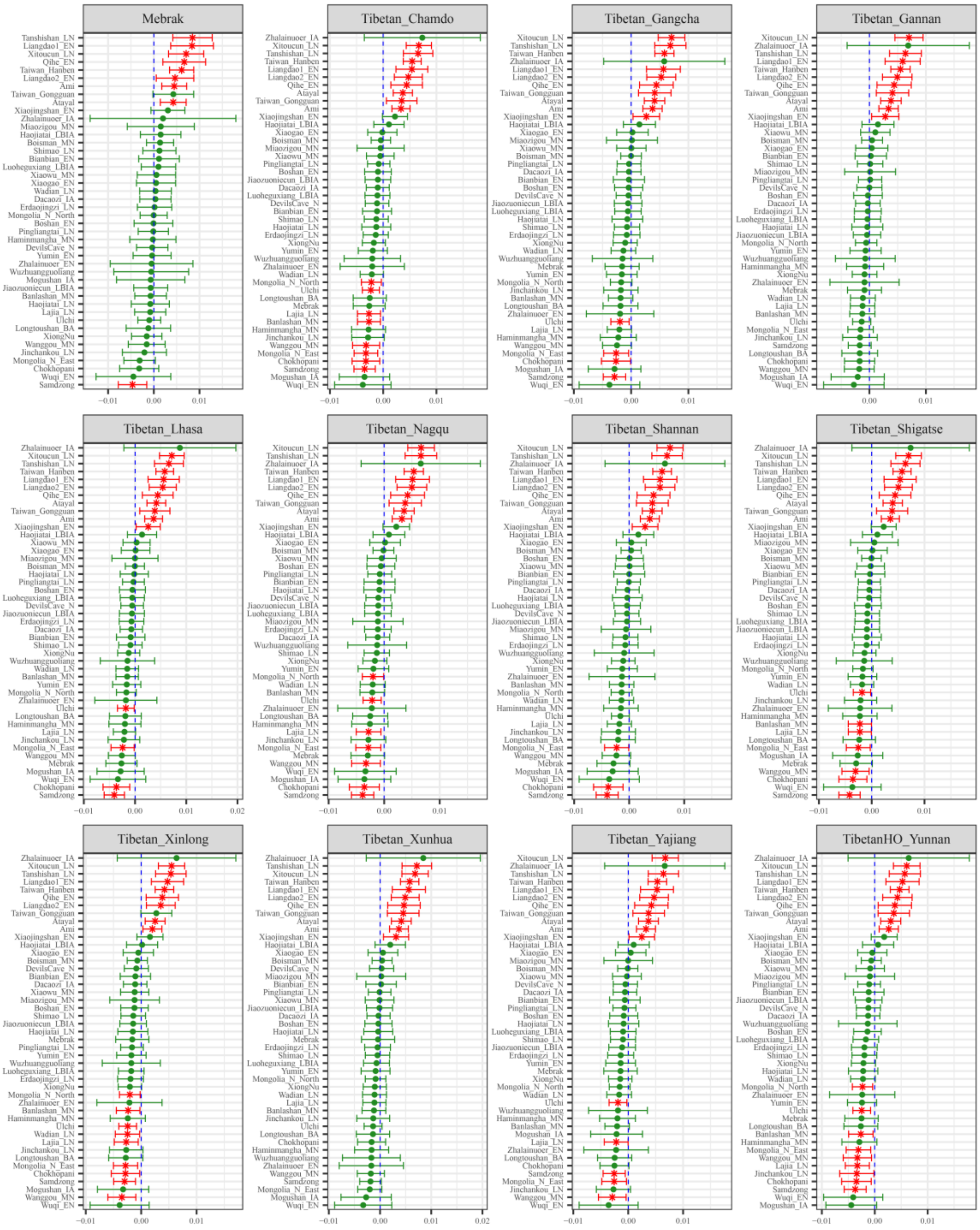

**Figure S55. Results of affinity-$f_4$ statistics for the form $f_4$*(Suogang_LN, Modern Tibetan; Neolithic to Historic East Asians, Mbuti)* showed the additional shared derived alleles from source populations except for Yangtze rice farmer related ancestral populations.**

Here, overlapping SNP loci included in the Affymetrix Human Origins platform among four analyzed populations were used. We used the genetic variation of Mbuti as the outgroup. Red asterisk point meant the significant value (Absolute value of Z-scores larger than three or equal to three) observed in the symmetry-$f_4$ statistics and green circle point denoted the non-significant $f_4$-statistic values (Absolute value of Z-scores less than three). All ancient East Asians were listed along the Y-axis and $f_4$ values were labeled along the X-axis. All results were faceted or grouped via Tibetan populations. Significant negative $f_4$ values indicated that the third population shared more alleles with the second population, and also means Tibetan obtained additional gene flow from the third source population (or related

populations). And the significant positive $f_4$ value indicated that the third population shared more derived alleles with the first population. The value of $f_4$-statistics equal to zero was marked as the blue dash line. The bar indicated three standard errors.

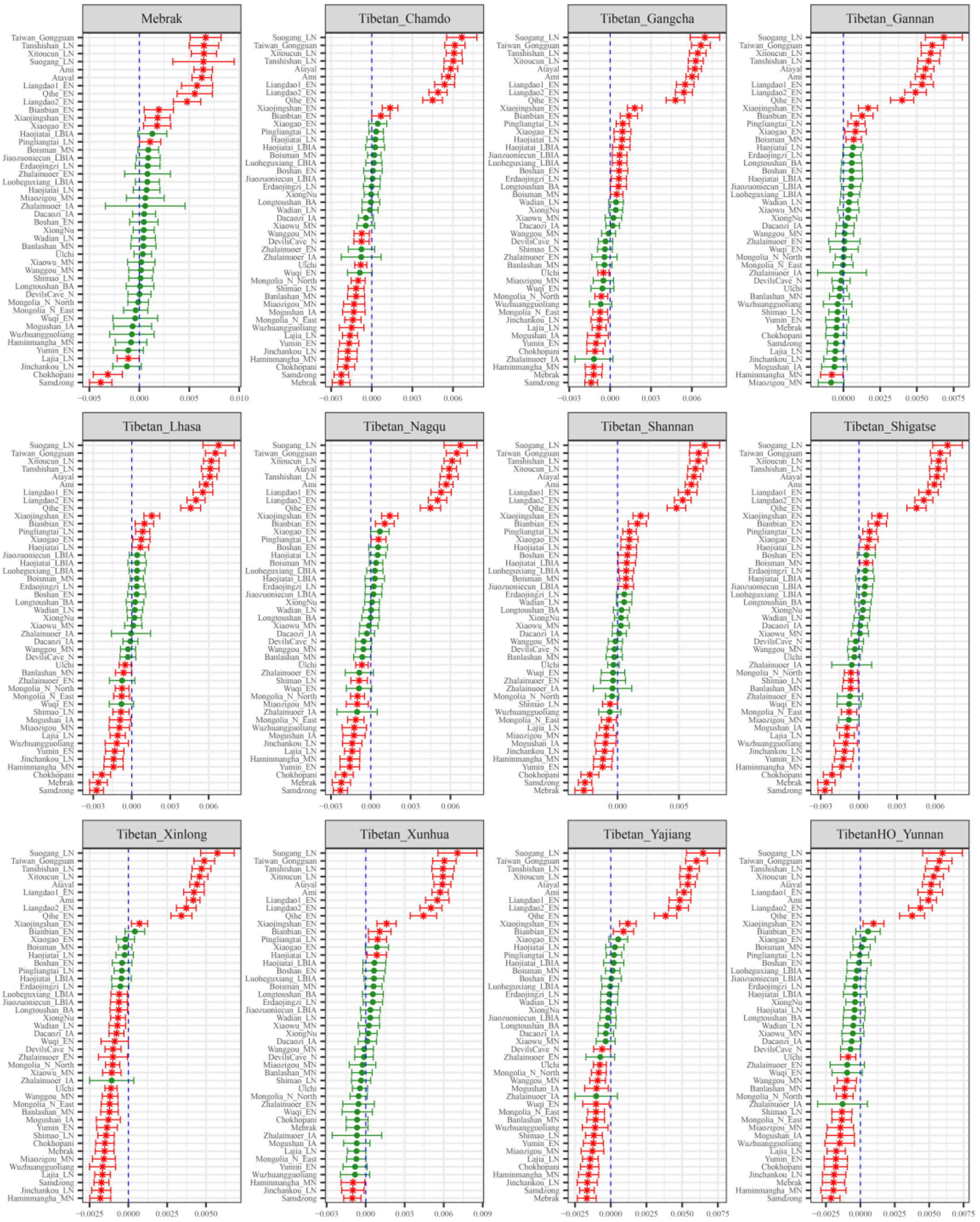

**Figure S56. Results of affinity-$f_4$ statistics for the form $f_4$*(Taiwan_Hanben, Modern Tibetan; Neolithic to Historic East Asians, Mbuti)* showed the additional shared derived alleles from source populations except for Yangtze rice farmer related ancestral populations.**

Here, overlapping SNP loci included in the Affymetrix Human Origins platform among four analyzed populations were used. We used the genetic variation of Mbuti as the outgroup. Red asterisk point meant the significant value (Absolute value of Z-scores larger than three or equal to three) observed in the symmetry-$f_4$ statistics and green circle point denoted the non-significant $f_4$-statistic values (Absolute value of Z-scores less than three). All ancient East Asians were listed along the Y-axis and $f_4$ values were labeled along the X-axis. All results were faceted or grouped via Tibetan populations. Significant negative $f_4$ values indicated that the third population shared more alleles with the second population, and

also means Tibetan obtained additional gene flow from the third source population (or related populations). And the significant positive $f_4$ value indicated that the third population shared more derived alleles with the first population. The value of $f_4$-statistics equal to zero was marked as the blue dash line. The bar indicated three standard errors.

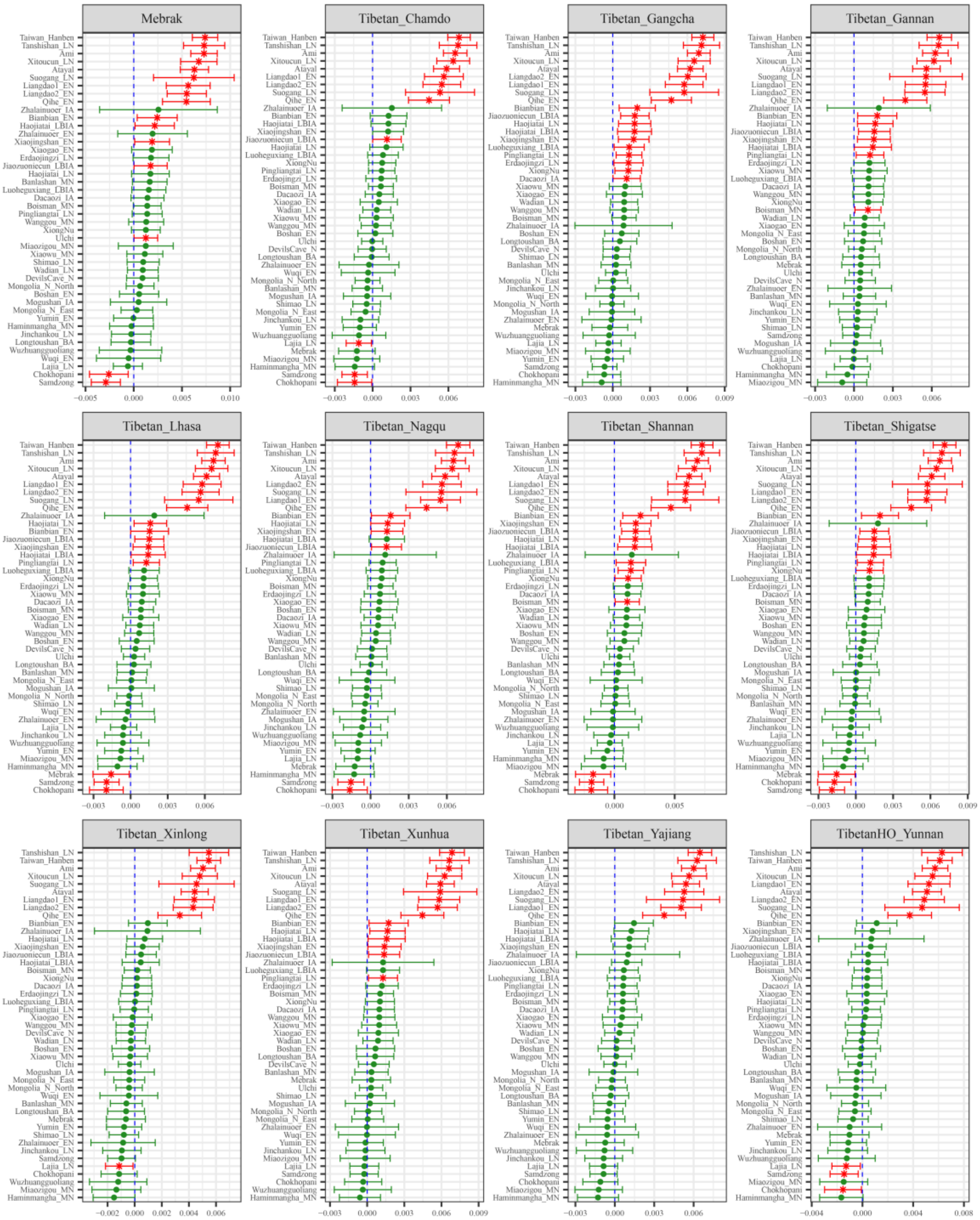

**Figure S57. Results of affinity-$f_4$ statistics for the form $f_4$*(Taiwan_Gongguan, Modern Tibetan; Neolithic to Historic East Asians, Mbuti)* showed the additional shared derived alleles from source populations except for Yangtze rice farmer related ancestral populations.**

Here, overlapping SNP loci included in the Affymetrix Human Origins platform among four analyzed populations were used. We used the genetic variation of Mbuti as the outgroup. Red asterisk point meant the significant value (Absolute value of Z-scores larger than three or equal to three) observed in the symmetry-$f_4$ statistics and green circle point denoted the non-significant $f_4$-statistic values (Absolute value of Z-scores less than three). All ancient East Asians were listed along the Y-axis and $f_4$ values were labeled along the X-axis. All results were faceted or grouped via Tibetan populations. Significant negative $f_4$ values indicated that the third population shared more alleles with the second population, and

also means Tibetan obtained additional gene flow from the third source population (or related populations). And the significant positive $f_4$ value indicated that the third population shared more derived alleles with the first population. The value of $f_4$-statistics equal to zero was marked as the blue dash line. The bar indicated three standard errors.

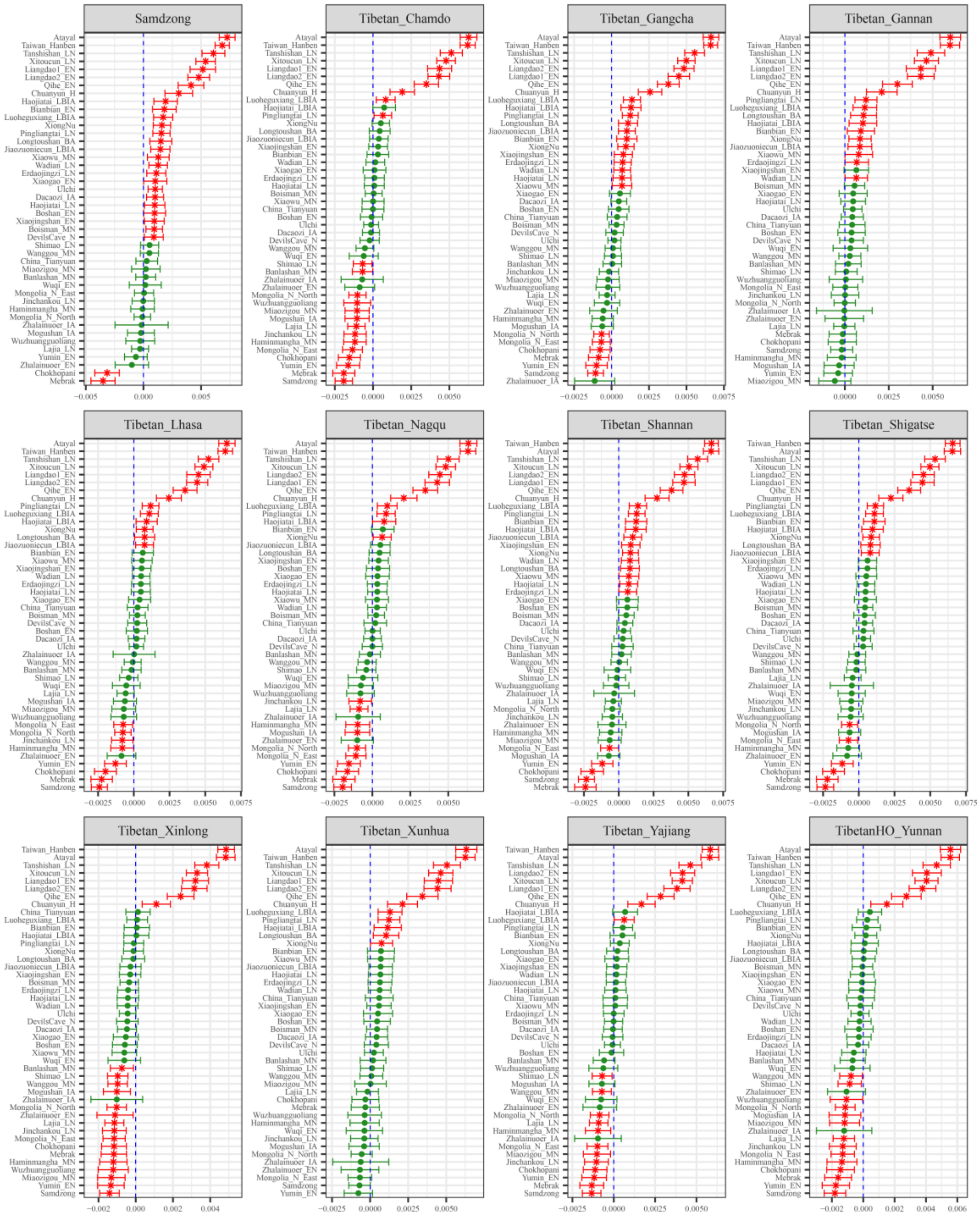

**Figure S58. Results of affinity-$f_4$ statistics for the form $f_4$*(Ami, Modern Tibetan; Neolithic to Historic East Asians, Mbuti)* showed the additional shared derived alleles from source populations except for Yangtze rice farmer related ancestral populations.**

Here, overlapping SNP loci included in the Affymetrix Human Origins platform among four analyzed populations were used. We used the genetic variation of Mbuti as the outgroup. Red asterisk point meant the significant value (Absolute value of Z-scores larger than three or equal to three) observed in the symmetry-$f_4$ statistics and green circle point denoted the non-significant $f_4$-statistic values (Absolute value of Z-scores less than three). All ancient East Asians were listed along the Y-axis and $f_4$ values were labeled along the X-axis. All results were faceted or grouped via Tibetan populations. Significant negative $f_4$ values indicated that the third population shared more alleles with the second population, and

also means Tibetan obtained additional gene flow from the third source population (or related populations). And the significant positive $f_4$ value indicated that the third population shared more derived alleles with the first population. The value of $f_4$-statistics equal to zero was marked as the blue dash line. The bar indicated three standard errors.

**Additional gene flow events when we assumed that Tibetans' direct ancestor is coastal Neolithic northern East Asian related ancestral populations**

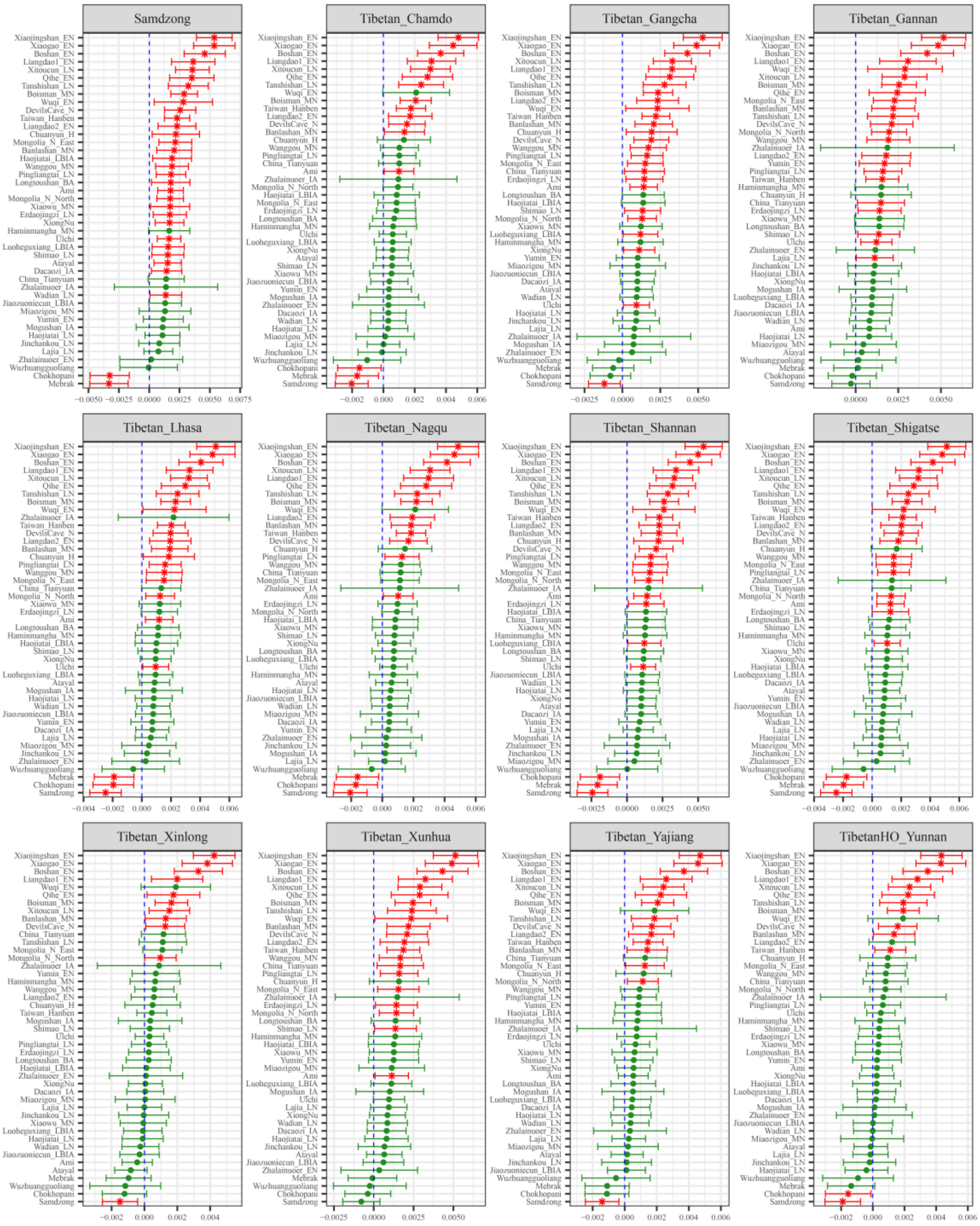

**Figure S59. Results of affinity-$f_4$ statistics for the form $f_4$*(Bianbian_EN, Modern Tibetan; Neolithic to Historic East Asians, Mbuti)* showed the additional shared derived alleles from source populations except for coastal Neolithic northern East Asian related ancestral populations.** Here, overlapping SNP loci included in the Affymetrix Human Origins platform among four analyzed populations were used. We used the genetic variation of Mbuti as the outgroup. Red asterisk point meant the significant value (Absolute value of Z-scores larger than three or equal to three) observed in the symmetry-$f_4$ statistics and green circle point denoted the non-significant $f_4$-statistic values (Absolute value of Z-scores less than three). All ancient East Asians were listed along the Y-axis and $f_4$ values were labeled along the X-axis. All results were faceted or grouped via Tibetan populations. Significant negative $f_4$ values indicated that the third population shared more alleles with the second population, and also means Tibetan obtained additional gene flow from the third source population (or related populations). And the significant positive $f_4$ value indicated that the third population shared more derived alleles with the first population. The value of $f_4$-statistics equal to zero was marked as the blue dash line.

The bar indicated three standard errors.

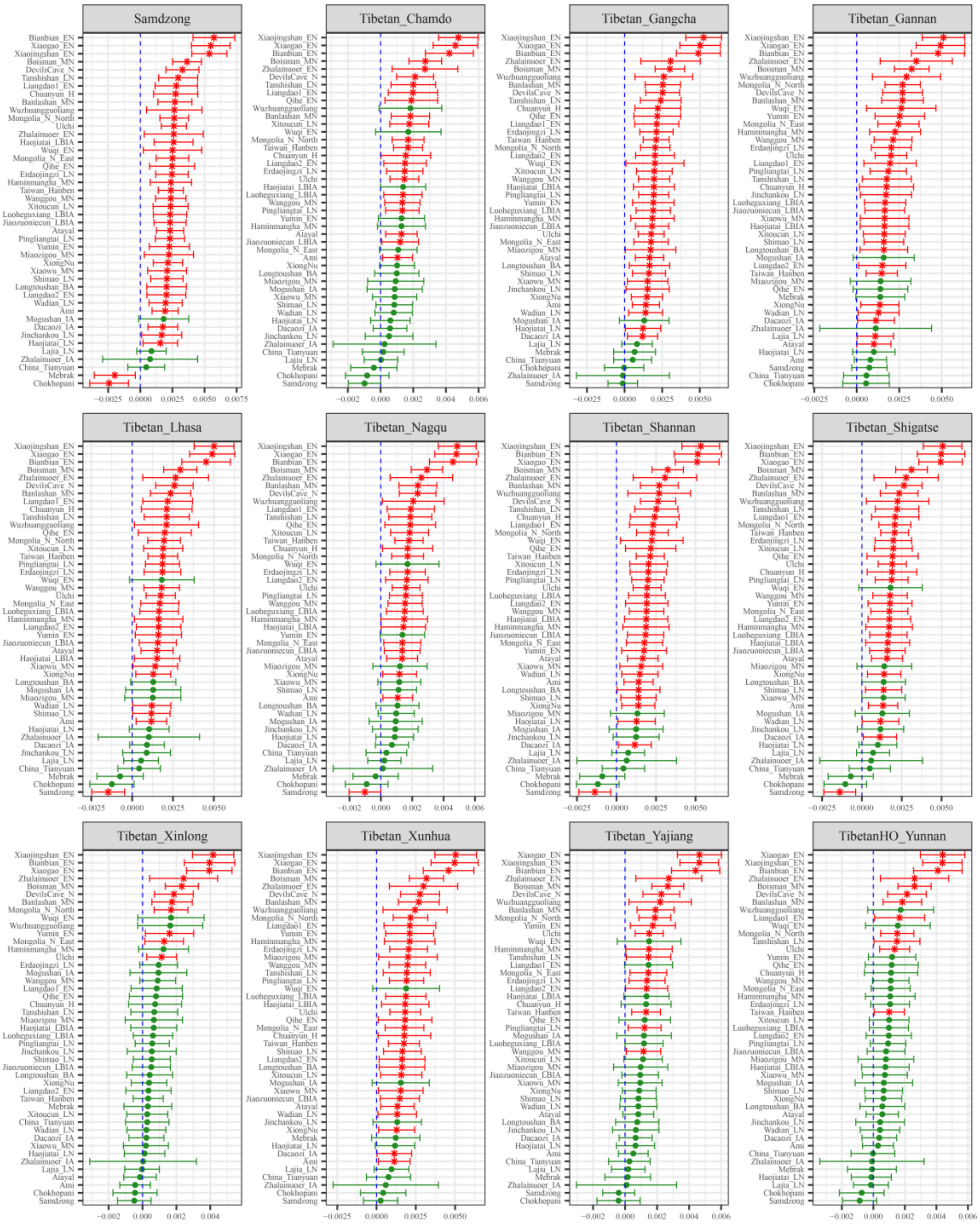

**Figure S60. Results of affinity-$f_4$ statistics for the form $f_4$*(Boshan_EN, Modern Tibetan; Neolithic to Historic East Asians, Mbuti)* showed the additional shared derived alleles from source populations except for coastal Neolithic northern East Asian related ancestral populations.** Here, overlapping SNP loci included in the Affymetrix Human Origins platform among four analyzed populations were used. We used the genetic variation of Mbuti as the outgroup. Red asterisk point meant the significant value (Absolute value of Z-scores larger than three or equal to three) observed in the symmetry-$f_4$ statistics and green circle point denoted the non-significant $f_4$-statistic values (Absolute value of Z-scores less than three). All ancient East Asians were listed along the Y-axis and $f_4$ values were labeled along the X-axis. All results were faceted or grouped via Tibetan populations. Significant negative $f_4$ values indicated that the third population shared more alleles with the second population, and also means Tibetan obtained additional gene flow from the third source population (or related populations). And the significant positive $f_4$ value indicated that the third population shared more derived alleles with the first population. The value of $f_4$-statistics equal to zero was marked as the blue dash line. The bar indicated three standard errors.

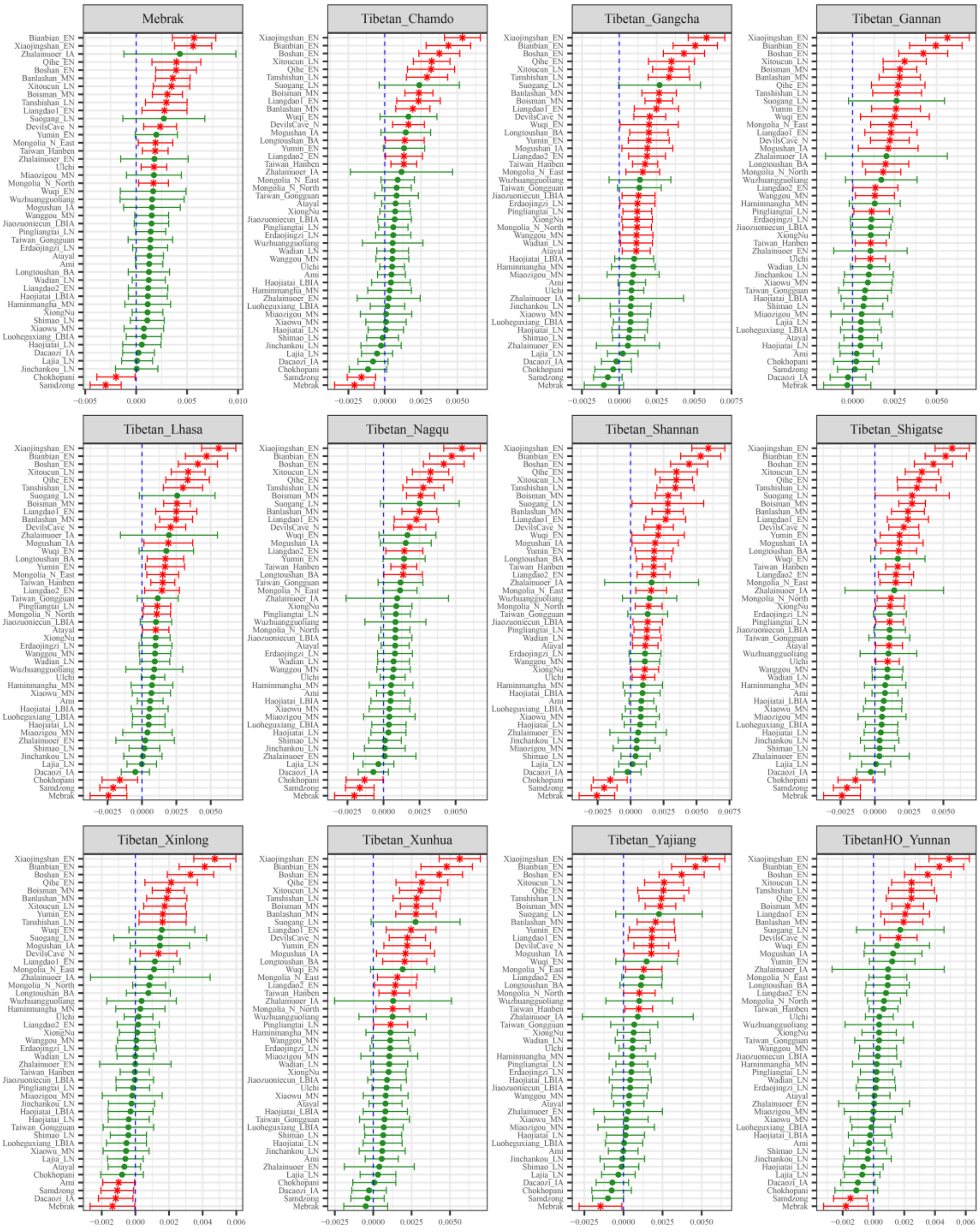

**Figure S61. Results of affinity-$f_4$ statistics for the form $f_4$*(Xiaogao_EN, Modern Tibetan; Neolithic to Historic East Asians, Mbuti)* showed the additional shared derived alleles from source populations except for coastal Neolithic northern East Asian related ancestral populations.**

Here, overlapping SNP loci included in the Affymetrix Human Origins platform among four analyzed populations were used. We used the genetic variation of Mbuti as the outgroup. Red asterisk point meant the significant value (Absolute value of Z-scores larger than three or equal to three) observed in the symmetry-$f_4$ statistics and green circle point denoted the non-significant $f_4$-statistic values (Absolute value of Z-scores less than three). All ancient East Asians were listed along the Y-axis and $f_4$ values were labeled along the X-axis. All results were faceted or grouped via Tibetan populations. Significant negative $f_4$ values indicated that the third population shared more alleles with the second population, and also means Tibetan obtained additional gene flow from the third source population (or related populations). And the significant positive $f_4$ value indicated that the third population shared more derived alleles with the first population. The value of $f_4$-statistics equal to zero was marked as the blue dash line.

The bar indicated three standard errors.

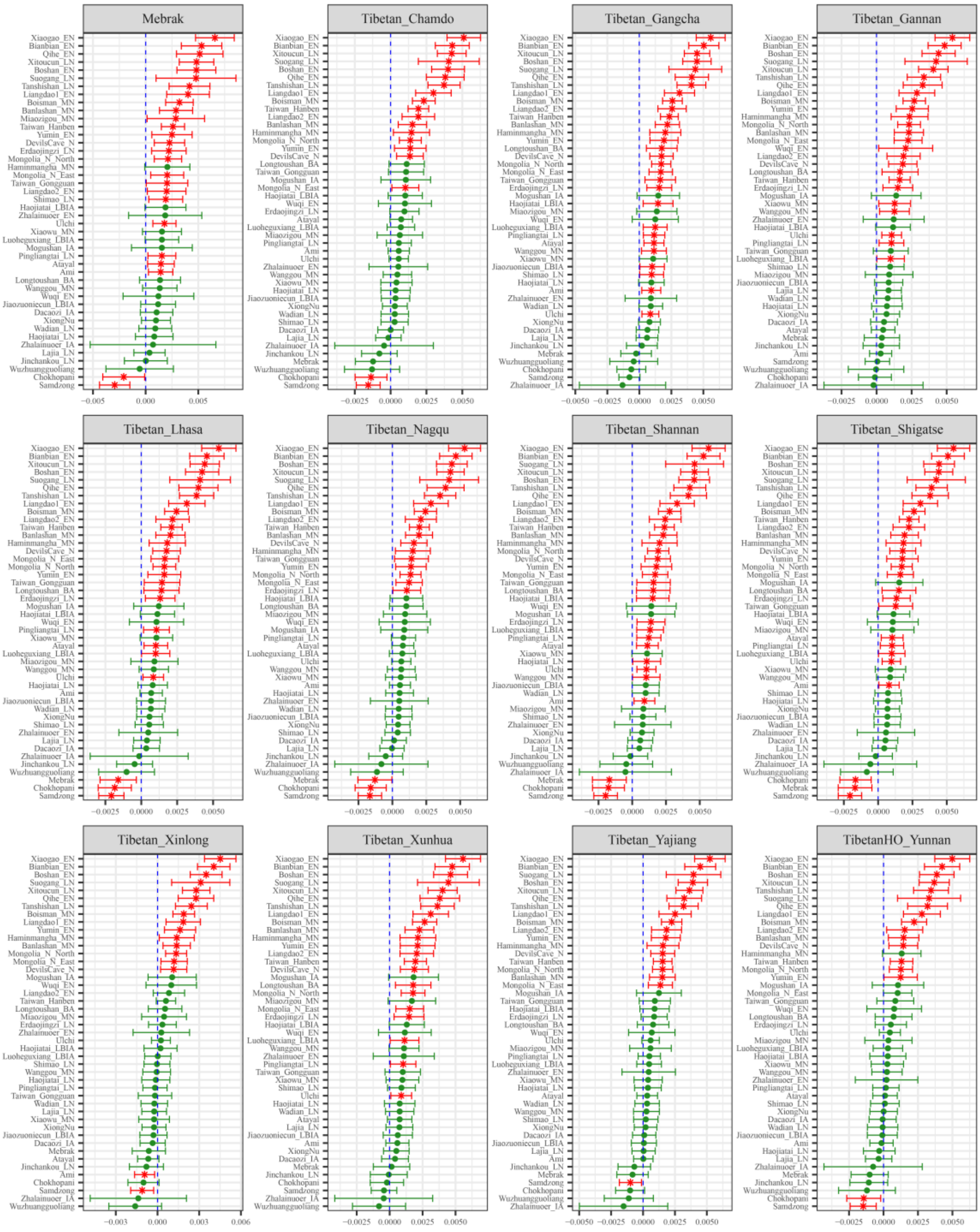

**Figure S62. Results of affinity-$f_4$ statistics for the form $f_4$*(Xiaojingshan_EN, Modern Tibetan; Neolithic to Historic East Asians, Mbuti)* showed the additional shared derived alleles from source populations except for coastal Neolithic northern East Asian related ancestral populations.**

Here, overlapping SNP loci included in the Affymetrix Human Origins platform among four analyzed populations were used. We used the genetic variation of Mbuti as the outgroup. Red asterisk point meant the significant value (Absolute value of Z-scores larger than three or equal to three) observed in the symmetry-$f_4$ statistics and green circle point denoted the non-significant $f_4$-statistic values (Absolute value of Z-scores less than three). All ancient East Asians were listed along the Y-axis and $f_4$ values were labeled along the X-axis. All results were faceted or grouped via Tibetan populations. Significant negative $f_4$ values indicated that the third population shared more alleles with the second population, and also means Tibetan obtained additional gene flow from the third source population (or related populations). And the significant positive $f_4$ value indicated that the third population shared more derived alleles with the first population. The value of $f_4$-statistics equal to zero was marked as the blue dash line.

The bar indicated three standard errors.

**Additional gene flow events when we assumed that Tibetans' direct ancestor is Henan Neolithic to Iron Age East Asian related ancestral populations**

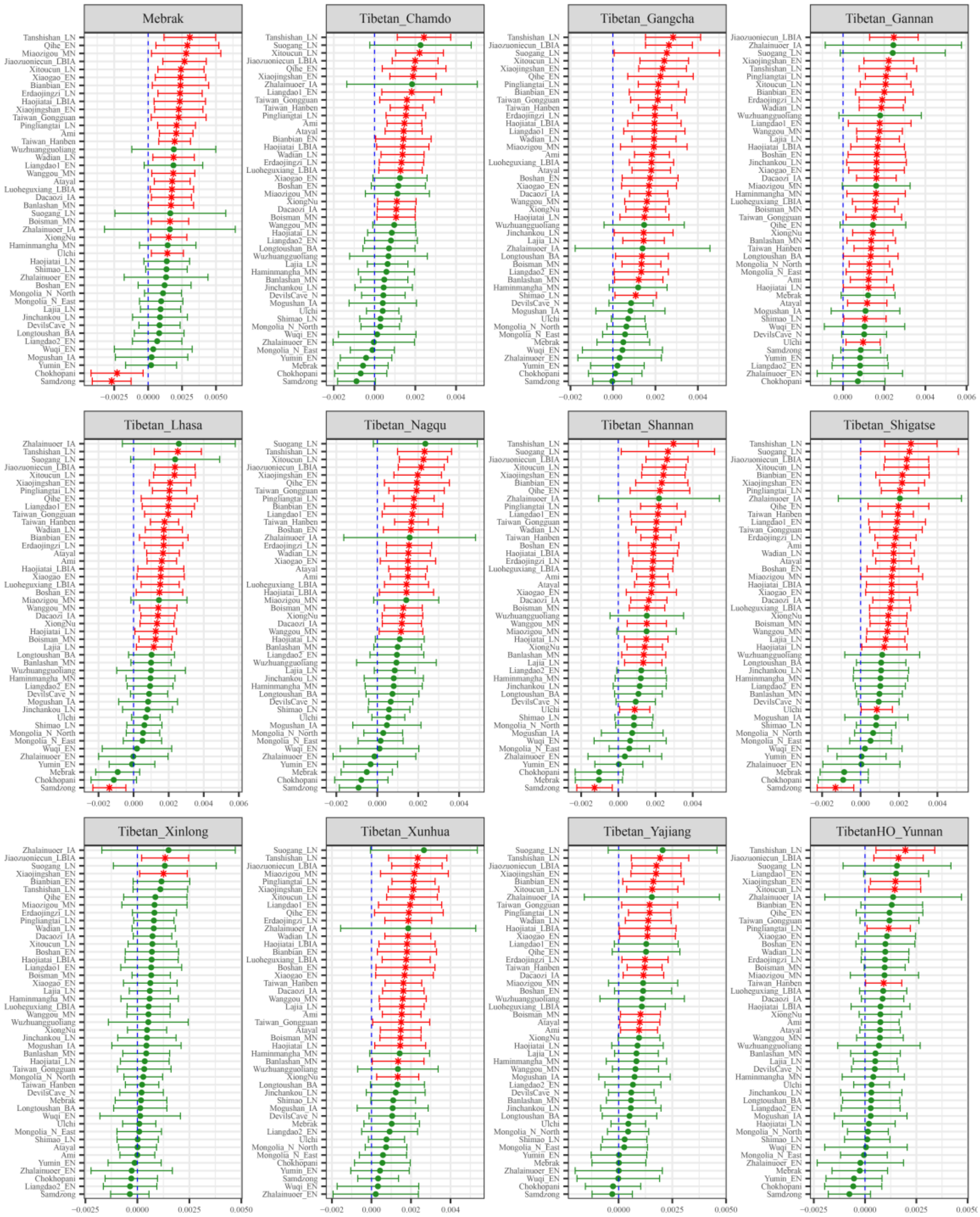

**Figure S63. Results of affinity-$f_4$ statistics for the form $f_4$*(Xiaowu_LN, Modern Tibetan; Neolithic to Historic East Asians, Mbuti)* showed the additional shared derived alleles from source populations except for Henan ancient population or their related ancestral populations.**
Here, overlapping SNP loci included in the Affymetrix Human Origins platform among four analyzed populations were used. We used the genetic variation of Mbuti as the outgroup. Red asterisk point meant the significant value (Absolute value of Z-scores larger than three or equal to three) observed in the symmetry-$f_4$ statistics and green circle point denoted the non-significant $f_4$-statistic values (Absolute value of Z-scores less than three). All ancient East Asians were listed along the Y-axis and $f_4$ values were labeled along the X-axis. All results were faceted or grouped via Tibetan populations. Significant negative $f_4$ values indicated that the third population shared more alleles with the second population, and

also means Tibetan obtained additional gene flow from the third source population (or related populations). And the significant positive $f_4$ value indicated that the third population shared more derived alleles with the first population. The value of $f_4$-statistics equal to zero was marked as the blue dash line. The bar indicated three standard errors.

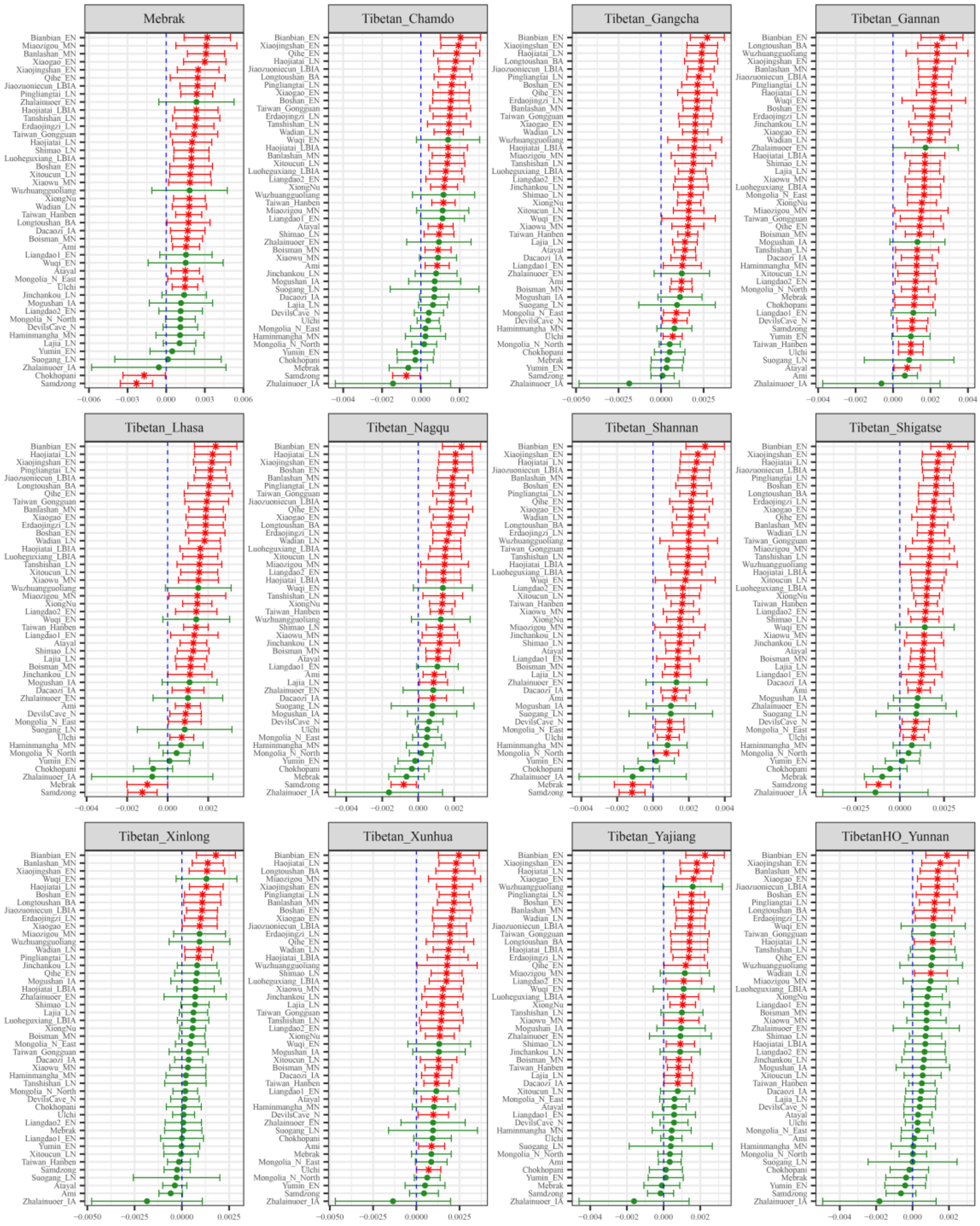

**Figure S64. Results of affinity-$f_4$ statistics for the form $f_4$(Wanggou_LN, *Modern Tibetan; Neolithic to Historic East Asians, Mbuti*) showed the additional shared derived alleles from source populations except for Henan ancient population or their related ancestral populations.**
Here, overlapping SNP loci included in the Affymetrix Human Origins platform among four analyzed populations were used. We used the genetic variation of Mbuti as the outgroup. Red asterisk point meant the significant value (Absolute value of Z-scores larger than three or equal to three) observed in the symmetry-$f_4$ statistics and green circle point denoted the non-significant $f_4$-statistic values (Absolute value of Z-scores less than three). All ancient East Asians were listed along the Y-axis and $f_4$ values were labeled along the X-axis. All results were faceted or grouped via Tibetan populations. Significant negative $f_4$ values indicated that the third population shared more alleles with the second population, and

also means Tibetan obtained additional gene flow from the third source population (or related populations). And the significant positive $f_4$ value indicated that the third population shared more derived alleles with the first population. The value of $f_4$-statistics equal to zero was marked as the blue dash line. The bar indicated three standard errors.

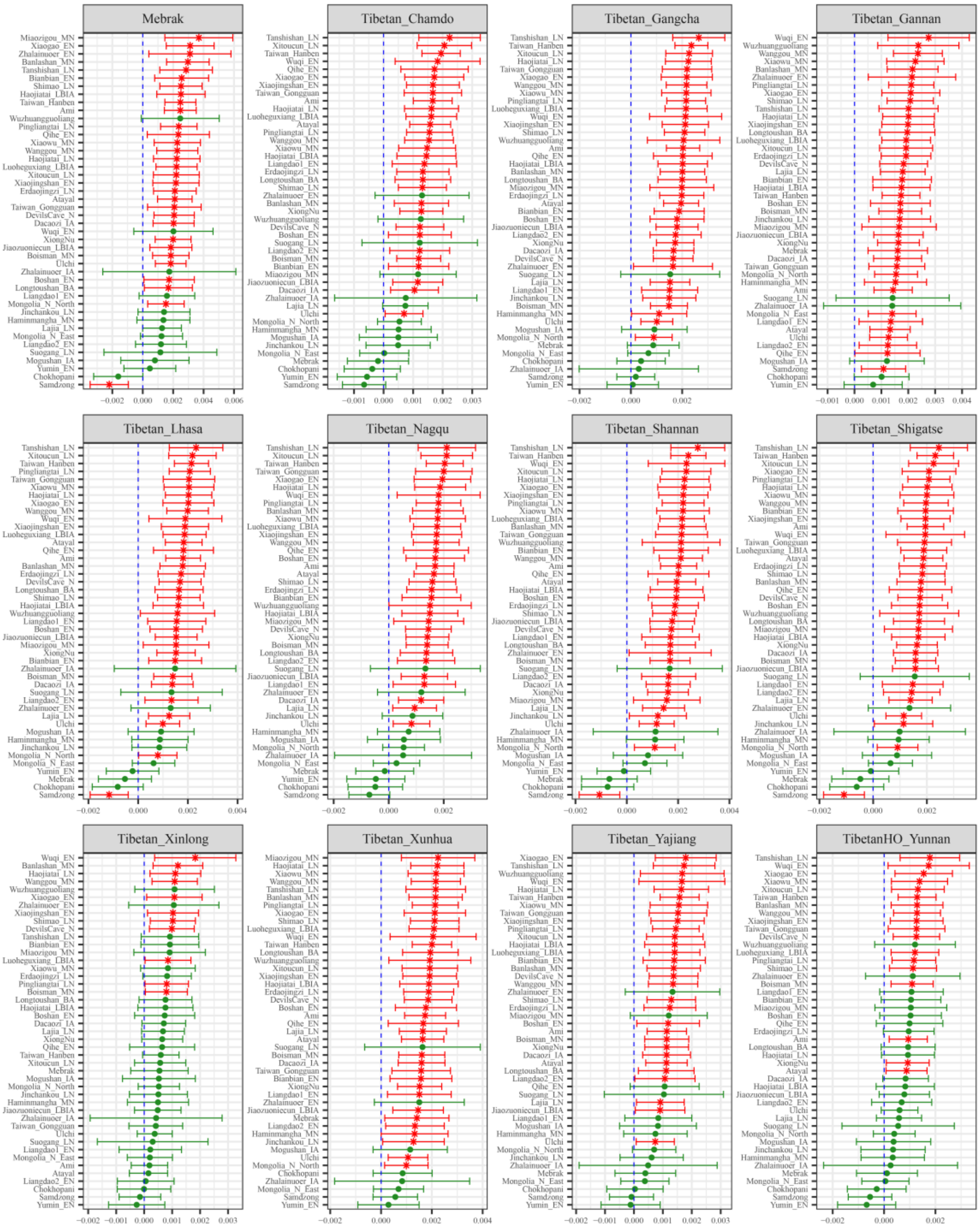

**Figure S65. Results of affinity-$f_4$ statistics for the form $f_4$*(Wadian_LN, Modern Tibetan; Neolithic to Historic East Asians, Mbuti)* showed the additional shared derived alleles from source populations except for Henan ancient population or their related ancestral populations.**

Here, overlapping SNP loci included in the Affymetrix Human Origins platform among four analyzed populations were used. We used the genetic variation of Mbuti as the outgroup. Red asterisk point meant the significant value (Absolute value of Z-scores larger than three or equal to three) observed in the symmetry-$f_4$ statistics and green circle point denoted the non-significant $f_4$-statistic values (Absolute value of Z-scores less than three). All ancient East Asians were listed along the Y-axis and $f_4$ values were labeled along the X-axis. All results were faceted or grouped via Tibetan populations. Significant

negative $f_4$ values indicated that the third population shared more alleles with the second population, and also means Tibetan obtained additional gene flow from the third source population (or related populations). And the significant positive $f_4$ value indicated that the third population shared more derived alleles with the first population. The value of $f_4$-statistics equal to zero was marked as the blue dash line. The bar indicated three standard errors.

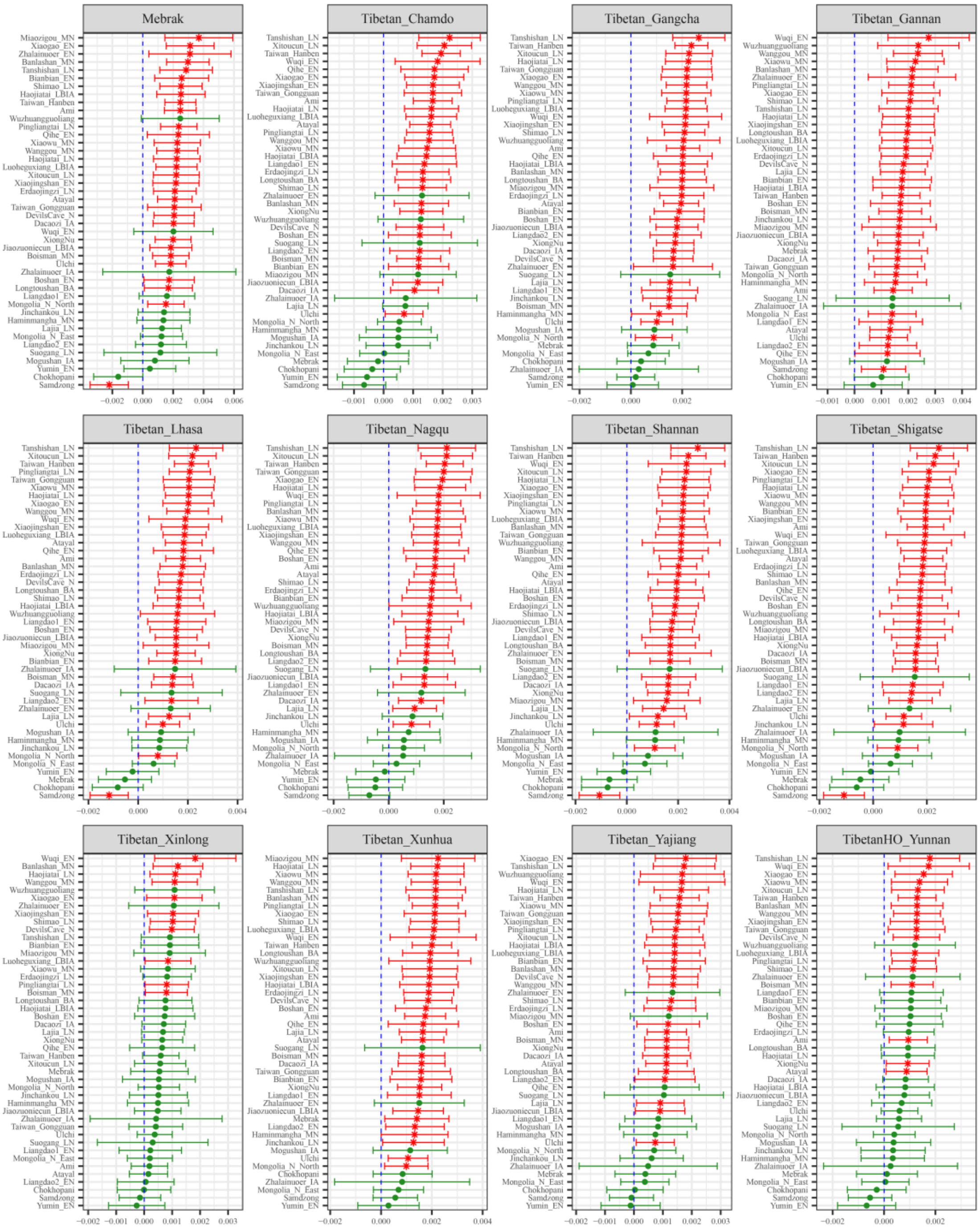

**Figure S66. Results of affinity-$f_4$ statistics for the form $f_4$*(Wadian_LN, Modern Tibetan; Neolithic to Historic East Asians, Mbuti)* showed the additional shared derived alleles from source populations except for Henan ancient population or their related ancestral populations.**

Here, overlapping SNP loci included in the Affymetrix Human Origins platform among four analyzed populations were used. We used the genetic variation of Mbuti as the outgroup. Red asterisk point meant the significant value (Absolute value of Z-scores larger than three or equal to three) observed in the symmetry-$f_4$ statistics and green circle point denoted the non-significant $f_4$-statistic values (Absolute value of Z-scores less than three). All ancient East Asians were listed along the Y-axis and $f_4$ values were labeled along the X-axis. All results were faceted or grouped via Tibetan populations. Significant

negative $f_4$ values indicated that the third population shared more alleles with the second population, and also means Tibetan obtained additional gene flow from the third source population (or related populations). And the significant positive $f_4$ value indicated that the third population shared more derived alleles with the first population. The value of $f_4$-statistics equal to zero was marked as the blue dash line. The bar indicated three standard errors.

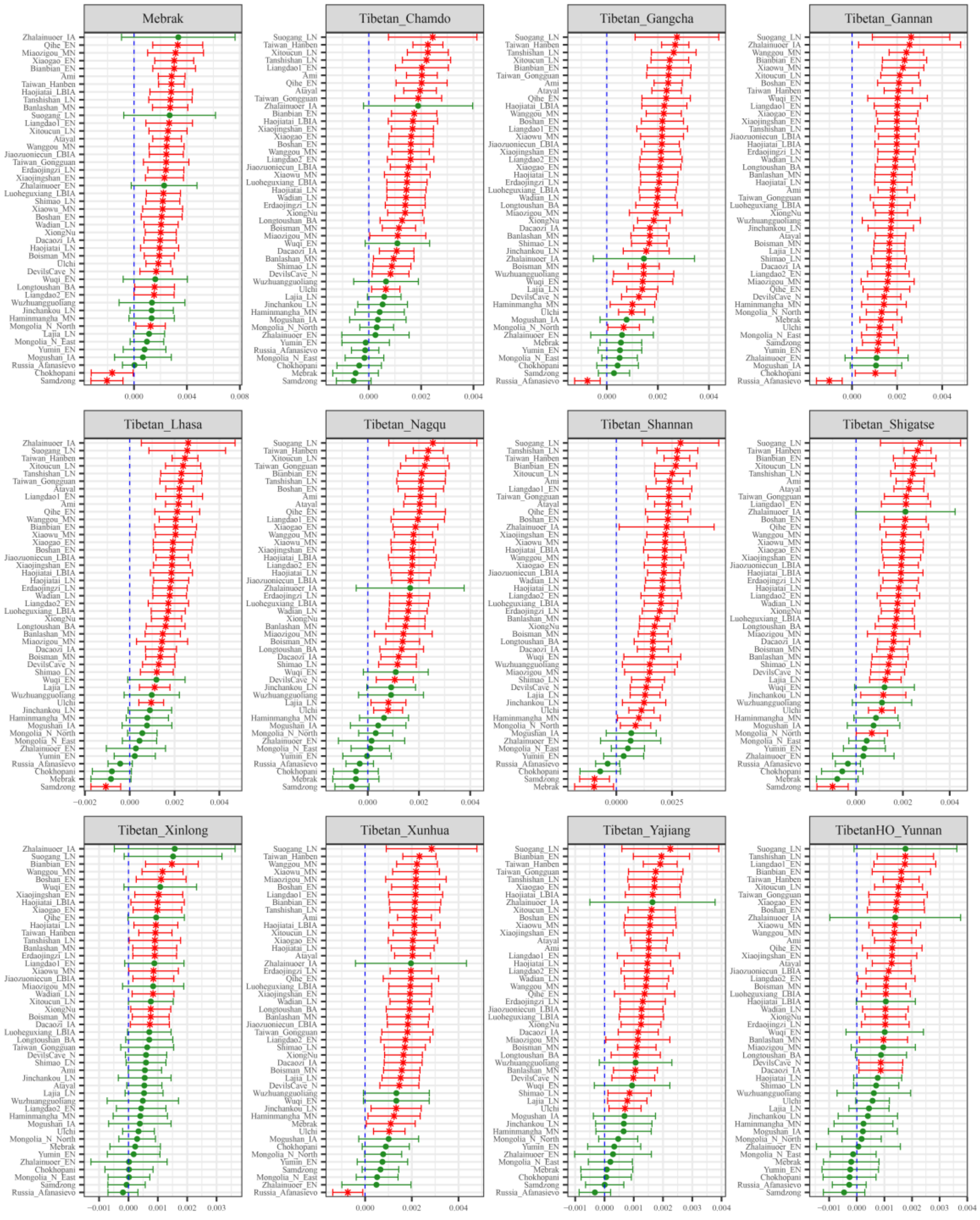

**Figure S67. Results of affinity-$f_4$ statistics for the form $f_4$*(Pingliangtai_LN, Modern Tibetan; Neolithic to Historic East Asians, Mbuti)* showed the additional shared derived alleles from source populations except for Henan ancient population or their related ancestral populations.**

Here, overlapping SNP loci included in the Affymetrix Human Origins platform among four analyzed populations were used. We used the genetic variation of Mbuti as the outgroup. Red asterisk point meant the significant value (Absolute value of Z-scores larger than three or equal to three) observed in the symmetry-$f_4$ statistics and green circle point denoted the non-significant $f_4$-statistic values (Absolute value of Z-scores less than three). All ancient East Asians were listed along the Y-axis and $f_4$ values were labeled along the X-axis. All results were faceted or grouped via Tibetan populations. Significant

negative $f_4$ values indicated that the third population shared more alleles with the second population, and also means Tibetan obtained additional gene flow from the third source population (or related populations). And the significant positive $f_4$ value indicated that the third population shared more derived alleles with the first population. The value of $f_4$-statistics equal to zero was marked as the blue dash line. The bar indicated three standard errors.

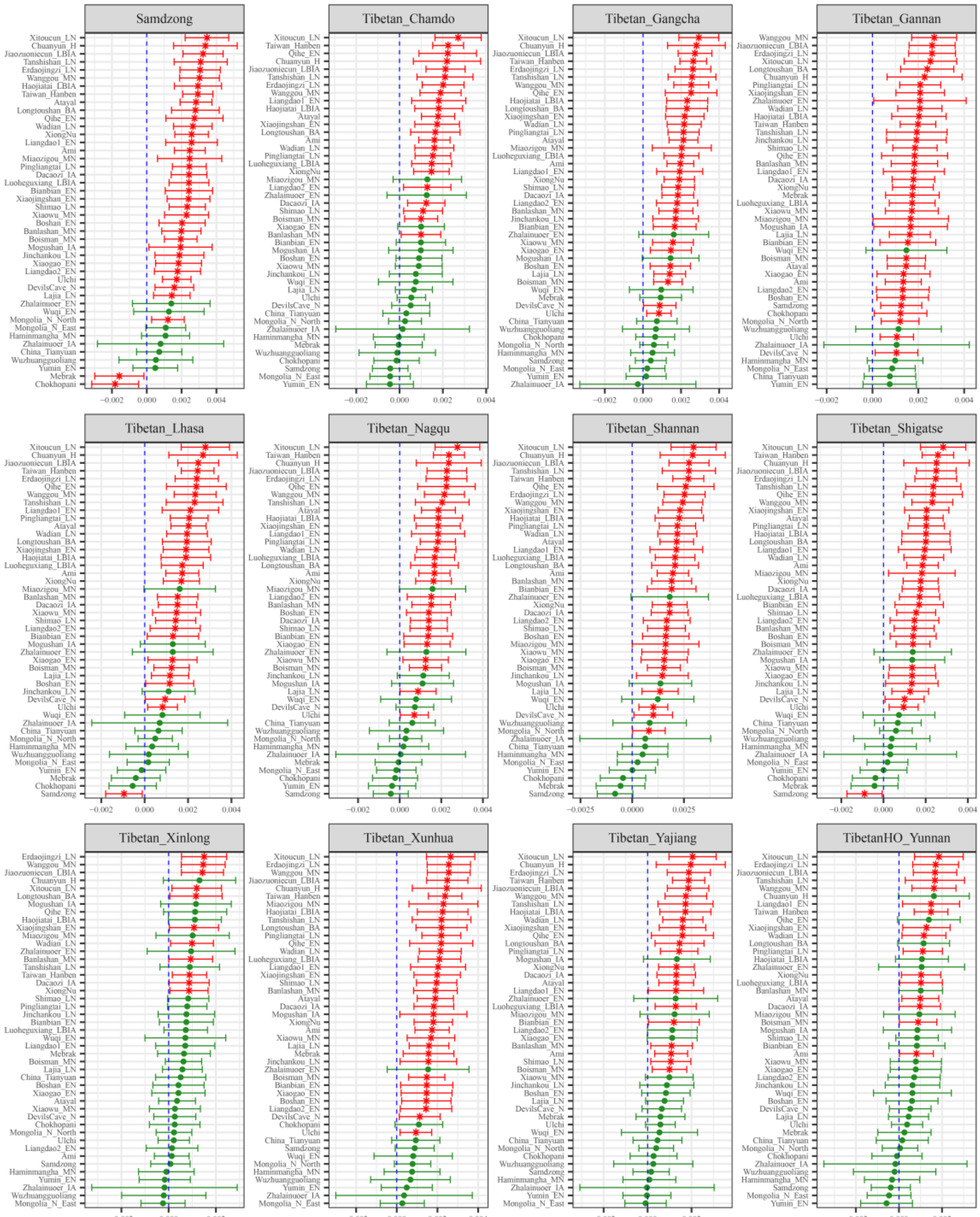

**Figure S68. Results of affinity-$f_4$ statistics for the form $f_4$*(Haojiatai_LN, Modern Tibetan; Neolithic to Historic East Asians, Mbuti)* showed the additional shared derived alleles from source populations except for Henan ancient population or their related ancestral populations.**

Here, overlapping SNP loci included in the Affymetrix Human Origins platform among four analyzed populations were used. We used the genetic variation of Mbuti as the outgroup. Red asterisk point meant the significant value (Absolute value of Z-scores larger than three or equal to three) observed in the symmetry-$f_4$ statistics and green circle point denoted the non-significant $f_4$-statistic values (Absolute value of Z-scores less than three). All ancient East Asians were listed along the Y-axis and $f_4$ values were

labeled along the X-axis. All results were faceted or grouped via Tibetan populations. Significant negative $f_4$ values indicated that the third population shared more alleles with the second population, and also means Tibetan obtained additional gene flow from the third source population (or related populations). And the significant positive $f_4$ value indicated that the third population shared more derived alleles with the first population. The value of $f_4$-statistics equal to zero was marked as the blue dash line. The bar indicated three standard errors.

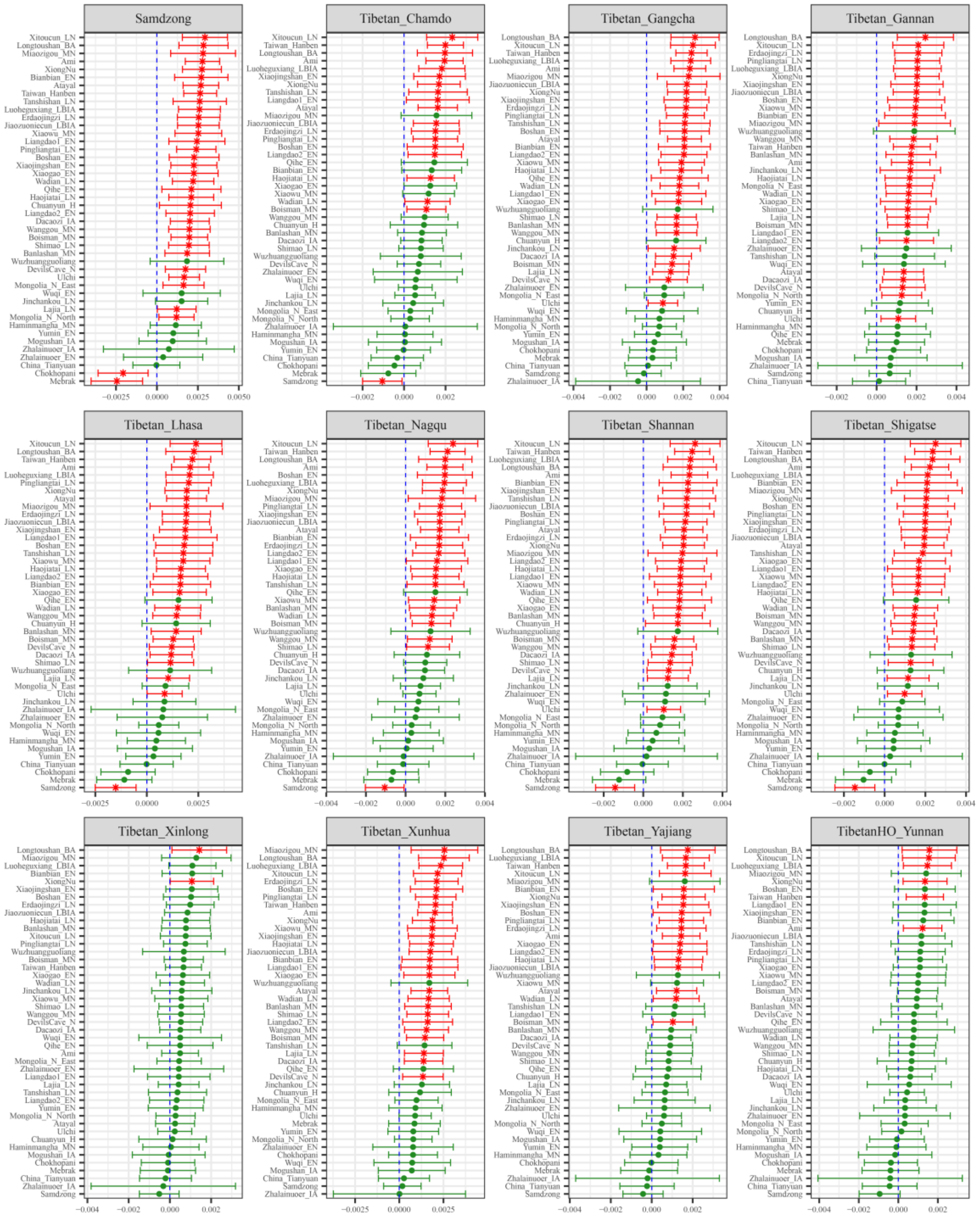

**Figure S69. Results of affinity-$f_4$ statistics for the form $f_4$*(Haojiatai_LBIA, Modern Tibetan; Neolithic to Historic East Asians, Mbuti)* showed the additional shared derived alleles from source populations except for Henan ancient population or their related ancestral populations.**

Here, overlapping SNP loci included in the Affymetrix Human Origins platform among four analyzed populations were used. We used the genetic variation of Mbuti as the outgroup. Red asterisk point meant the significant value (Absolute value of Z-scores larger than three or equal to three) observed in the symmetry-$f_4$ statistics and green circle point denoted the non-significant $f_4$-statistic values (Absolute value of Z-scores less than three). All ancient East Asians were listed along the Y-axis and $f_4$ values were

labeled along the X-axis. All results were faceted or grouped via Tibetan populations. Significant negative $f_4$ values indicated that the third population shared more alleles with the second population, and also means Tibetan obtained additional gene flow from the third source population (or related populations). And the significant positive $f_4$ value indicated that the third population shared more derived alleles with the first population. The value of $f_4$-statistics equal to zero was marked as the blue dash line. The bar indicated three standard errors.

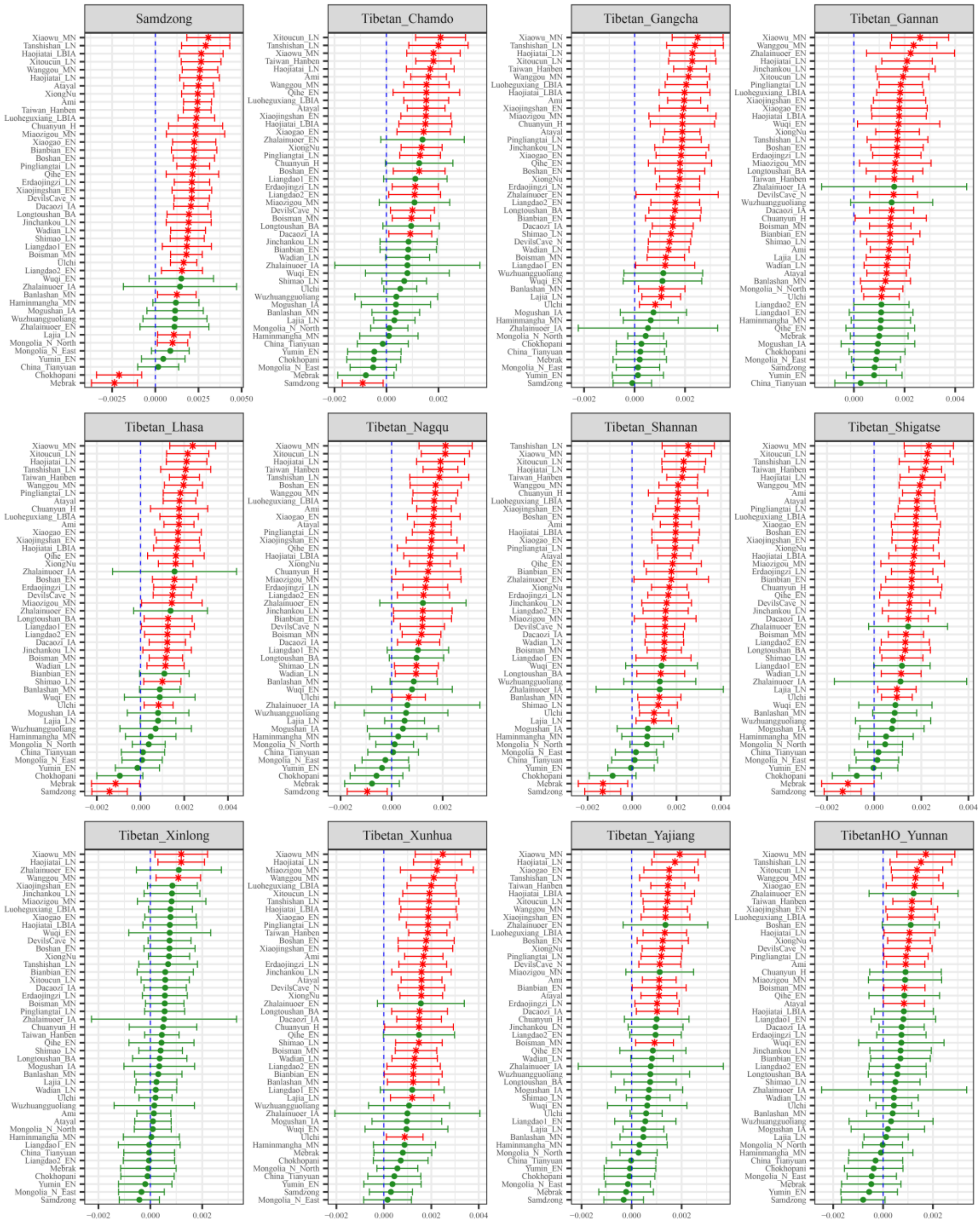

**Figure S70. Results of affinity-$f_4$ statistics for the form $f_4$(*Jiaozuoniecun_LBIA, Modern Tibetan; Neolithic to Historic East Asians, Mbuti*) showed the additional shared derived alleles from source populations except for Henan ancient population or their related ancestral populations.**

Here, overlapping SNP loci included in the Affymetrix Human Origins platform among four analyzed populations were used. We used the genetic variation of Mbuti as the outgroup. Red asterisk point meant the significant value (Absolute value of Z-scores larger than three or equal to three) observed in the symmetry-$f_4$ statistics and green circle point denoted the non-significant $f_4$-statistic values (Absolute value of Z-scores less than three). All ancient East Asians were listed along the Y-axis and $f_4$ values were

labeled along the X-axis. All results were faceted or grouped via Tibetan populations. Significant negative $f_4$ values indicated that the third population shared more alleles with the second population, and also means Tibetan obtained additional gene flow from the third source population (or related populations). And the significant positive $f_4$ value indicated that the third population shared more derived alleles with the first population. The value of $f_4$-statistics equal to zero was marked as the blue dash line. The bar indicated three standard errors.

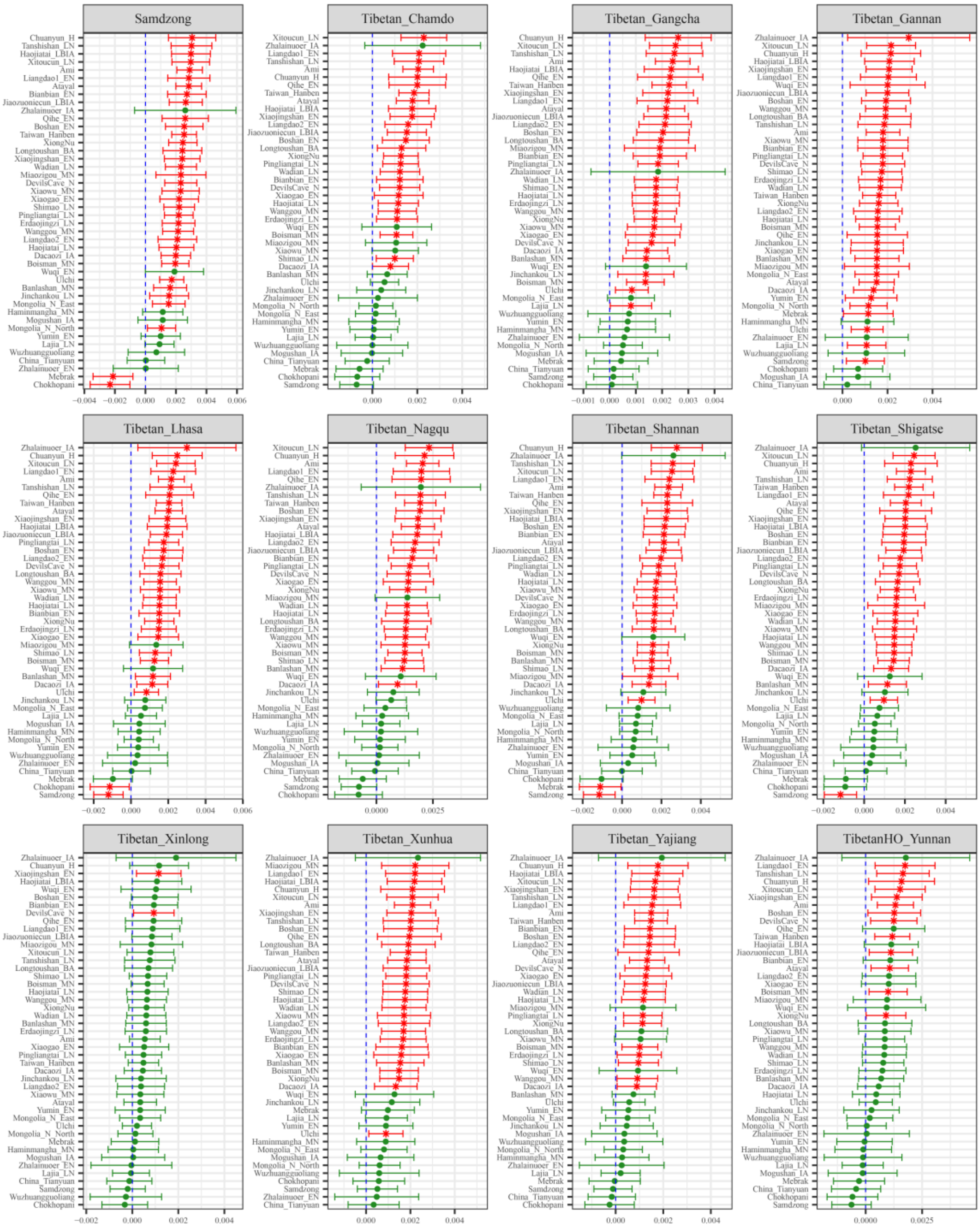

**Figure S71. Results of affinity-$f_4$ statistics for the form $f_4$*(Luoheguxiang_LBIA, Modern Tibetan; Neolithic to Historic East Asians, Mbuti)* showed the additional shared derived alleles from source populations except for Henan ancient population or their related ancestral populations.**
Here, overlapping SNP loci included in the Affymetrix Human Origins platform among four analyzed populations were used. We used the genetic variation of Mbuti as the outgroup. Red asterisk point meant the significant value (Absolute value of Z-scores larger than three or equal to three) observed in the symmetry-$f_4$ statistics and green circle point denoted the non-significant $f_4$-statistic values (Absolute value of Z-scores less than three). All ancient East Asians were listed along the Y-axis and $f_4$ values were

labeled along the X-axis. All results were faceted or grouped via Tibetan populations. Significant negative $f_4$ values indicated that the third population shared more alleles with the second population, and also means Tibetan obtained additional gene flow from the third source population (or related populations). And the significant positive $f_4$ value indicated that the third population shared more derived alleles with the first population. The value of $f_4$-statistics equal to zero was marked as the blue dash line. The bar indicated three standard errors.

**Additional gene flow events when we assumed that Tibetans' direct ancestor is inland Neolithic northern East Asian from Shaanxi or Inner Mongolia or their related ancestral populations**

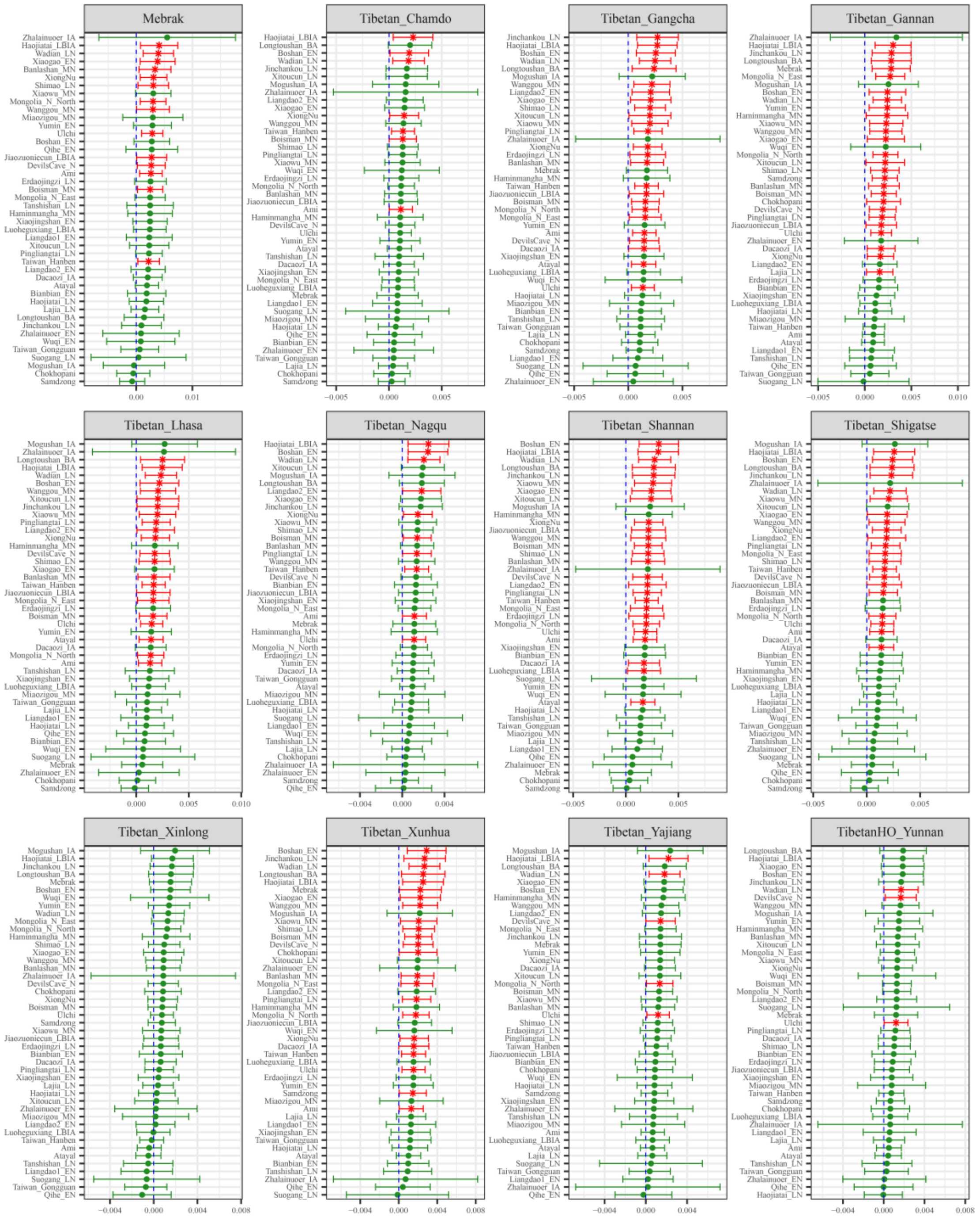

**Figure S72. Results of affinity-$f_4$ statistics for the form $f_4$*(Wuzhuangguoliang, Modern Tibetan; Neolithic to Historic East Asians, Mbuti)* showed the additional shared derived alleles from source populations except for inland Neolithic northern East Asian from Shaanxi or Inner Mongolia or their related ancestral populations.**

Here, overlapping SNP loci included in the Affymetrix Human Origins platform among four analyzed populations were used. We used the genetic variation of Mbuti as the outgroup. Red asterisk point meant the significant value (Absolute value of Z-scores larger than three or equal to three) observed in the symmetry-$f_4$ statistics and green circle point denoted the non-significant $f_4$-statistic values (Absolute value of Z-scores less than three). All ancient East Asians were listed along the Y-axis and $f_4$ values were labeled along the X-axis. All results were faceted or grouped via Tibetan populations. Significant negative $f_4$ values indicated that the third population shared more alleles with the second population, and also means Tibetan obtained additional gene flow from the third source population (or related

populations). And the significant positive $f_4$ value indicated that the third population shared more derived alleles with the first population. The value of $f_4$-statistics equal to zero was marked as the blue dash line. The bar indicated three standard errors.

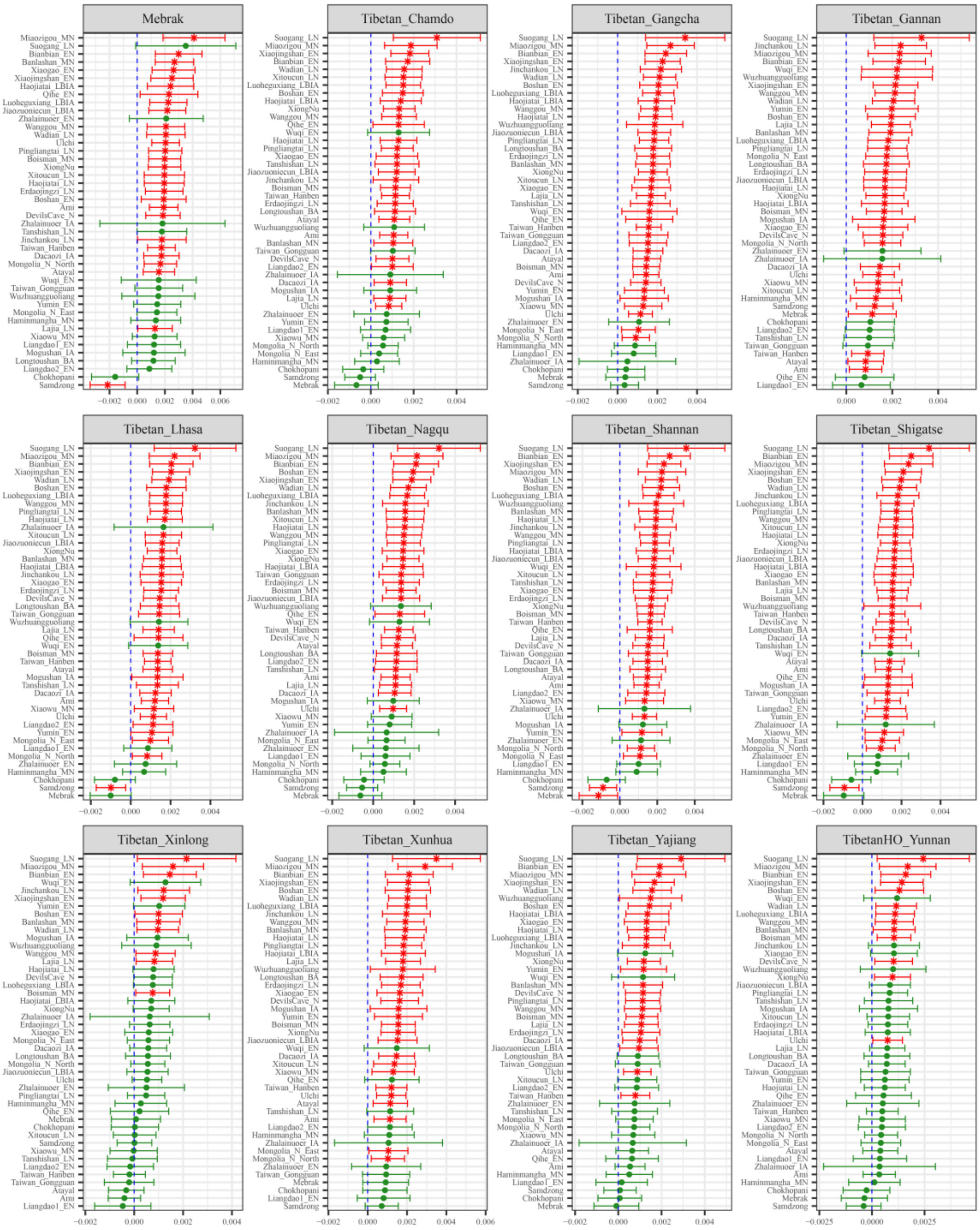

**Figure S73. Results of affinity-$f_4$ statistics for the form $f_4$*(Shimao_LN, Modern Tibetan; Neolithic to Historic East Asians, Mbuti)* showed the additional shared derived alleles from source populations except for inland Neolithic northern East Asian from Shaanxi or Inner Mongolia or their related ancestral populations.**

Here, overlapping SNP loci included in the Affymetrix Human Origins platform among four analyzed populations were used. We used the genetic variation of Mbuti as the outgroup. Red asterisk point meant the significant value (Absolute value of Z-scores larger than three or equal to three) observed in the symmetry-$f_4$ statistics and green circle point denoted the non-significant $f_4$-statistic values (Absolute value of Z-scores less than three). All ancient East Asians were listed along the Y-axis and $f_4$ values were labeled along the X-axis. All results were faceted or grouped via Tibetan populations. Significant negative $f_4$ values indicated that the third population shared more alleles with the second population, and

also means Tibetan obtained additional gene flow from the third source population (or related populations). And the significant positive $f_4$ value indicated that the third population shared more derived alleles with the first population. The value of $f_4$-statistics equal to zero was marked as the blue dash line. The bar indicated three standard errors.

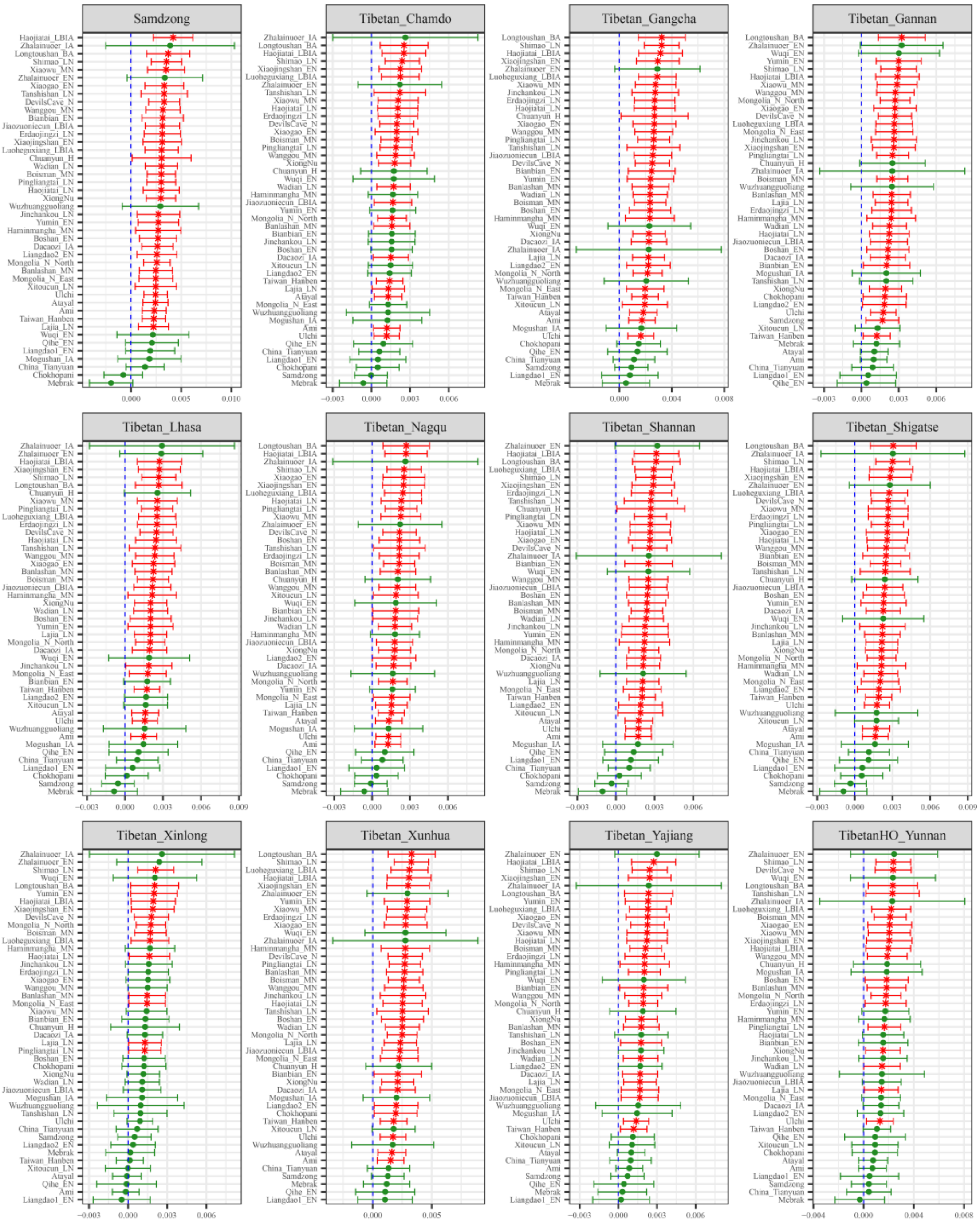

**Figure S74. Results of affinity-$f_4$ statistics for the form $f_4$(*Miaozigou_MN, Modern Tibetan; Neolithic to Historic East Asians, Mbuti*) showed the additional shared derived alleles from source populations except for inland Neolithic northern East Asian from Shaanxi or Inner Mongolia or their related ancestral populations.**

Here, overlapping SNP loci included in the Affymetrix Human Origins platform among four analyzed populations were used. We used the genetic variation of Mbuti as the outgroup. Red asterisk point meant the significant value (Absolute value of Z-scores larger than three or equal to three) observed in the symmetry-$f_4$ statistics and green circle point denoted the non-significant $f_4$-statistic values (Absolute value of Z-scores less than three). All ancient East Asians were listed along the Y-axis and $f_4$ values were labeled along the X-axis. All results were faceted or grouped via Tibetan populations. Significant

negative $f_4$ values indicated that the third population shared more alleles with the second population, and also means Tibetan obtained additional gene flow from the third source population (or related populations). And the significant positive $f_4$ value indicated that the third population shared more derived alleles with the first population. The value of $f_4$-statistics equal to zero was marked as the blue dash line. The bar indicated three standard errors.

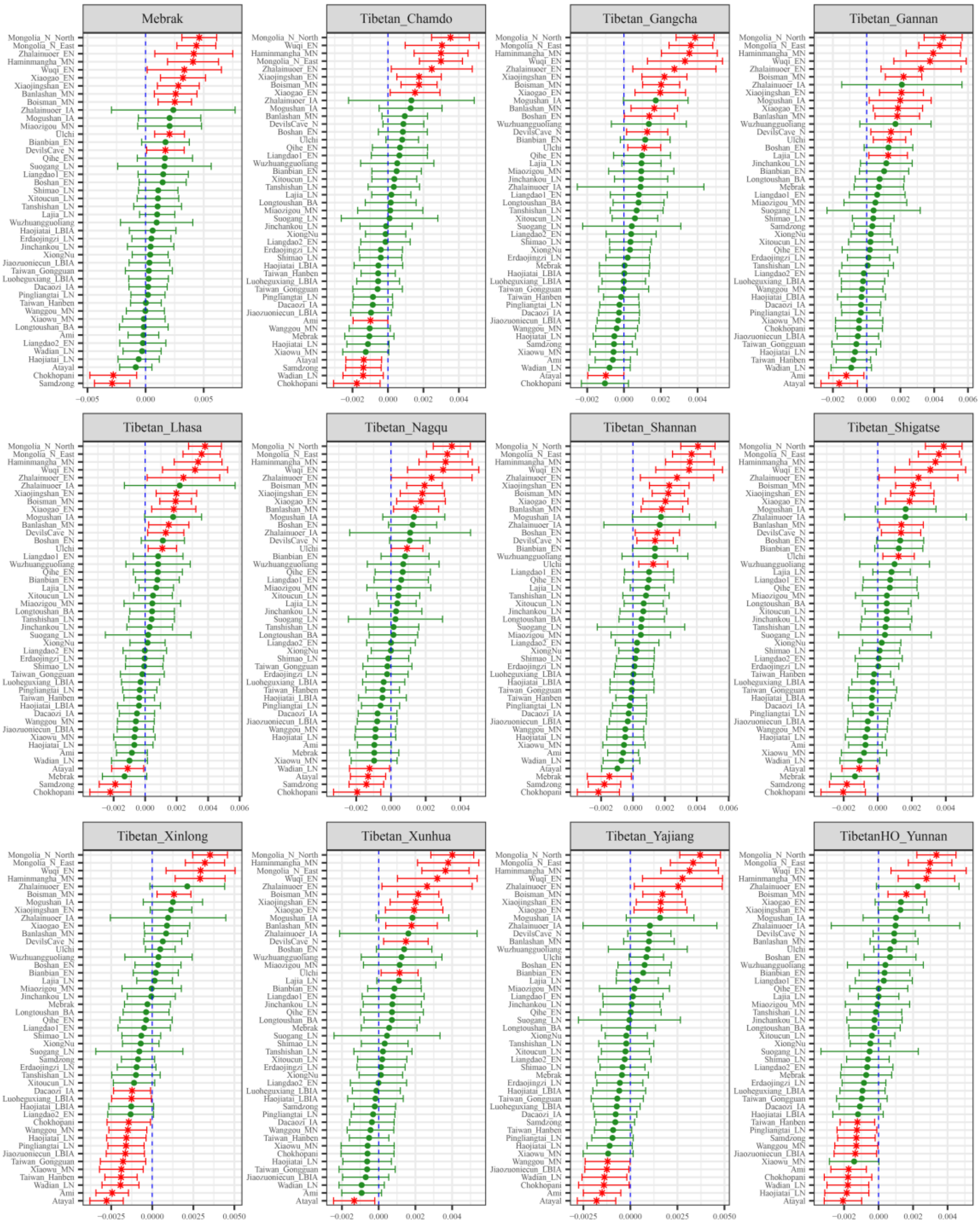

**Figure S75. Results of affinity-$f_4$ statistics for the form $f_4$(*Yumin_EN, Modern Tibetan; Neolithic to Historic East Asians, Mbuti*) showed the additional shared derived alleles from source populations except for inland Neolithic northern East Asian from Shaanxi or Inner Mongolia or their related ancestral populations.**

Here, overlapping SNP loci included in the Affymetrix Human Origins platform among four analyzed populations were used. We used the genetic variation of Mbuti as the outgroup. Red asterisk point meant the significant value (Absolute value of Z-scores larger than three or equal to three) observed in the symmetry-$f_4$ statistics and green circle point denoted the non-significant $f_4$-statistic values (Absolute value of Z-scores less than three). All ancient East Asians were listed along the Y-axis and $f_4$ values were

labeled along the X-axis. All results were faceted or grouped via Tibetan populations. Significant negative $f_4$ values indicated that the third population shared more alleles with the second population, and also means Tibetan obtained additional gene flow from the third source population (or related populations). And the significant positive $f_4$ value indicated that the third population shared more derived alleles with the first population. The value of $f_4$-statistics equal to zero was marked as the blue dash line. The bar indicated three standard errors.

**Additional gene flow events when we assumed that Tibetans' direct ancestor is inland Neolithic to Iron Age northern East Asian from upper Yellow River Basin or their related ancestral populations**

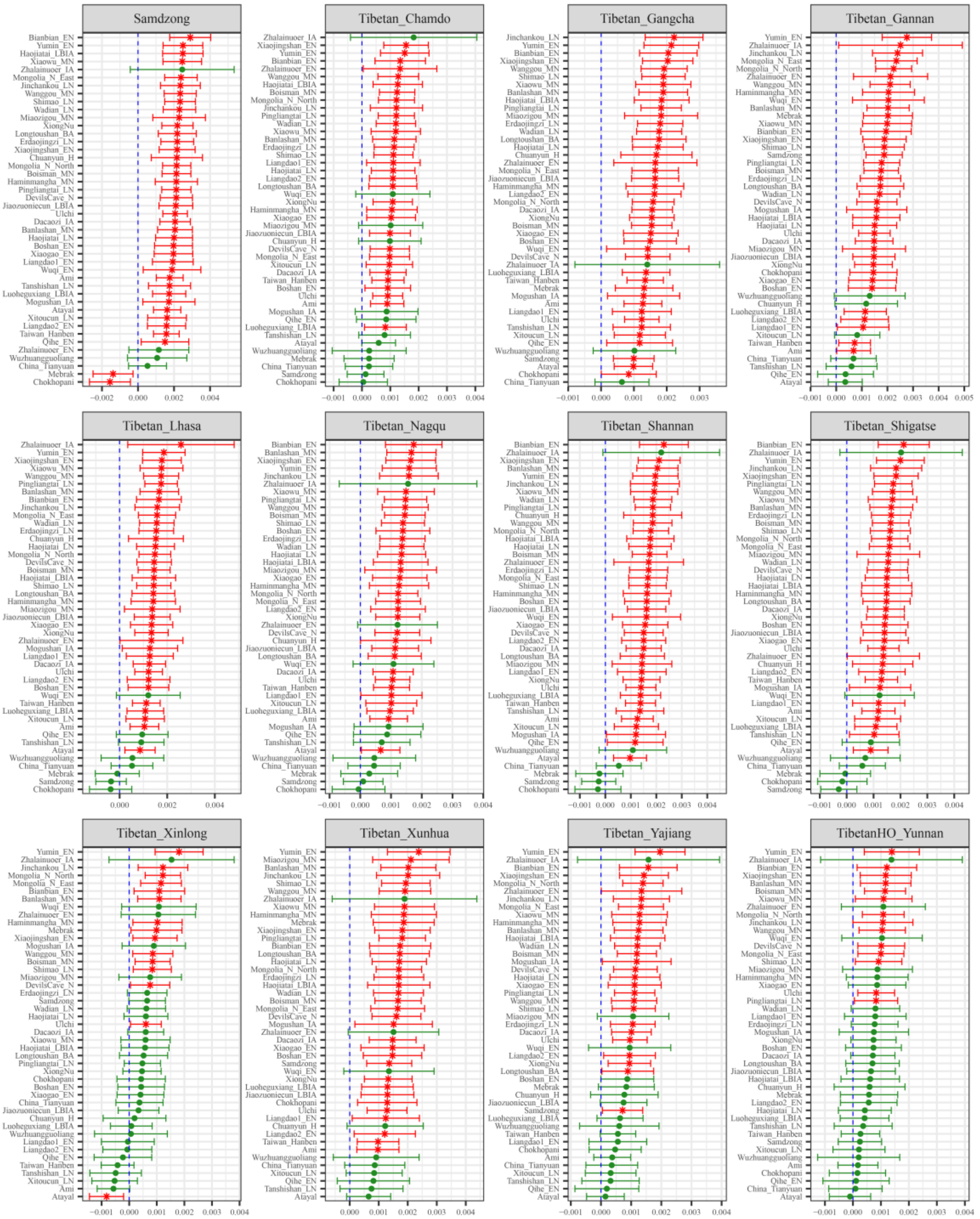

**Figure S76. Results of affinity-$f_4$ statistics for the form *$f_4$(Lajia_LN, Modern Tibetan; Neolithic to Historic East Asians, Mbuti)* showed the additional shared derived alleles from source populations except for inland Neolithic northern East Asian from Shaanxi or Inner Mongolia or their related ancestral populations.**

Here, overlapping SNP loci included in the Affymetrix Human Origins platform among four analyzed populations were used. We used the genetic variation of Mbuti as the outgroup. Red asterisk point meant the significant value (Absolute value of Z-scores larger than three or equal to three) observed in the symmetry-$f_4$ statistics and green circle point denoted the non-significant $f_4$-statistic values (Absolute value of Z-scores less than three). All ancient East Asians were listed along the Y-axis and $f_4$ values were labeled along the X-axis. All results were faceted or grouped via Tibetan populations. Significant negative $f_4$ values indicated that the third population shared more alleles with the second population, and also means Tibetan obtained additional gene flow from the third source population (or related

populations). And the significant positive $f_4$ value indicated that the third population shared more derived alleles with the first population. The value of $f_4$-statistics equal to zero was marked as the blue dash line. The bar indicated three standard errors.

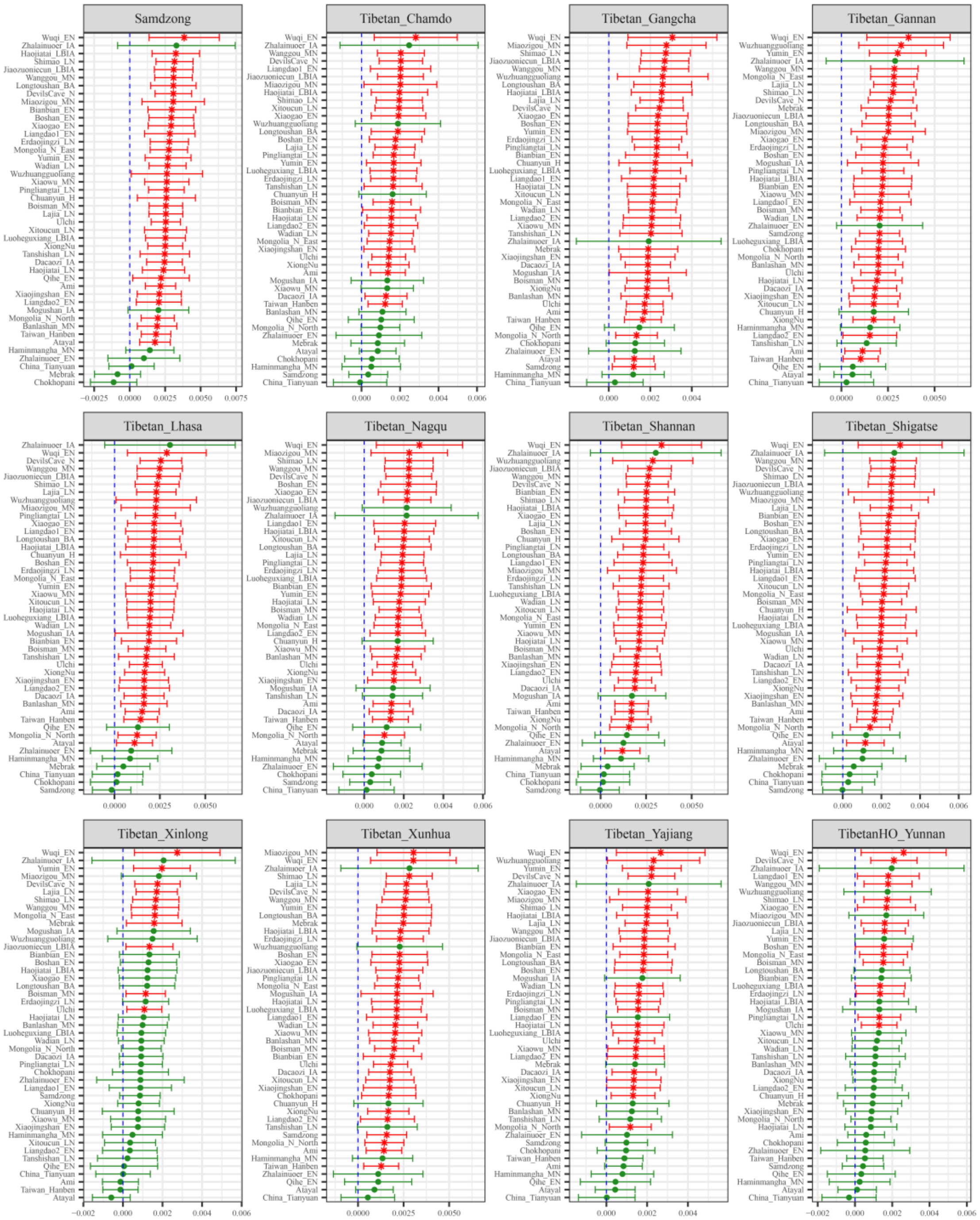

**Figure S77. Results of affinity-$f_4$ statistics for the form *$f_4$(Jinchankou_LN, Modern Tibetan; Neolithic to Historic East Asians, Mbuti)* showed the additional shared derived alleles from source populations except for inland Neolithic northern East Asian from Shaanxi or Inner Mongolia or their related ancestral populations.**

Here, overlapping SNP loci included in the Affymetrix Human Origins platform among four analyzed populations were used. We used the genetic variation of Mbuti as the outgroup. Red asterisk point meant the significant value (Absolute value of Z-scores larger than three or equal to three) observed in the symmetry-$f_4$ statistics and green circle point denoted the non-significant $f_4$-statistic values (Absolute value of Z-scores less than three). All ancient East Asians were listed along the Y-axis and $f_4$ values were labeled along the X-axis. All results were faceted or grouped via Tibetan populations. Significant negative $f_4$ values indicated that the third population shared more alleles with the second population, and also means Tibetan obtained additional gene flow from the third source population (or related

populations). And the significant positive $f_4$ value indicated that the third population shared more derived alleles with the first population. The value of $f_4$-statistics equal to zero was marked as the blue dash line. The bar indicated three standard errors.

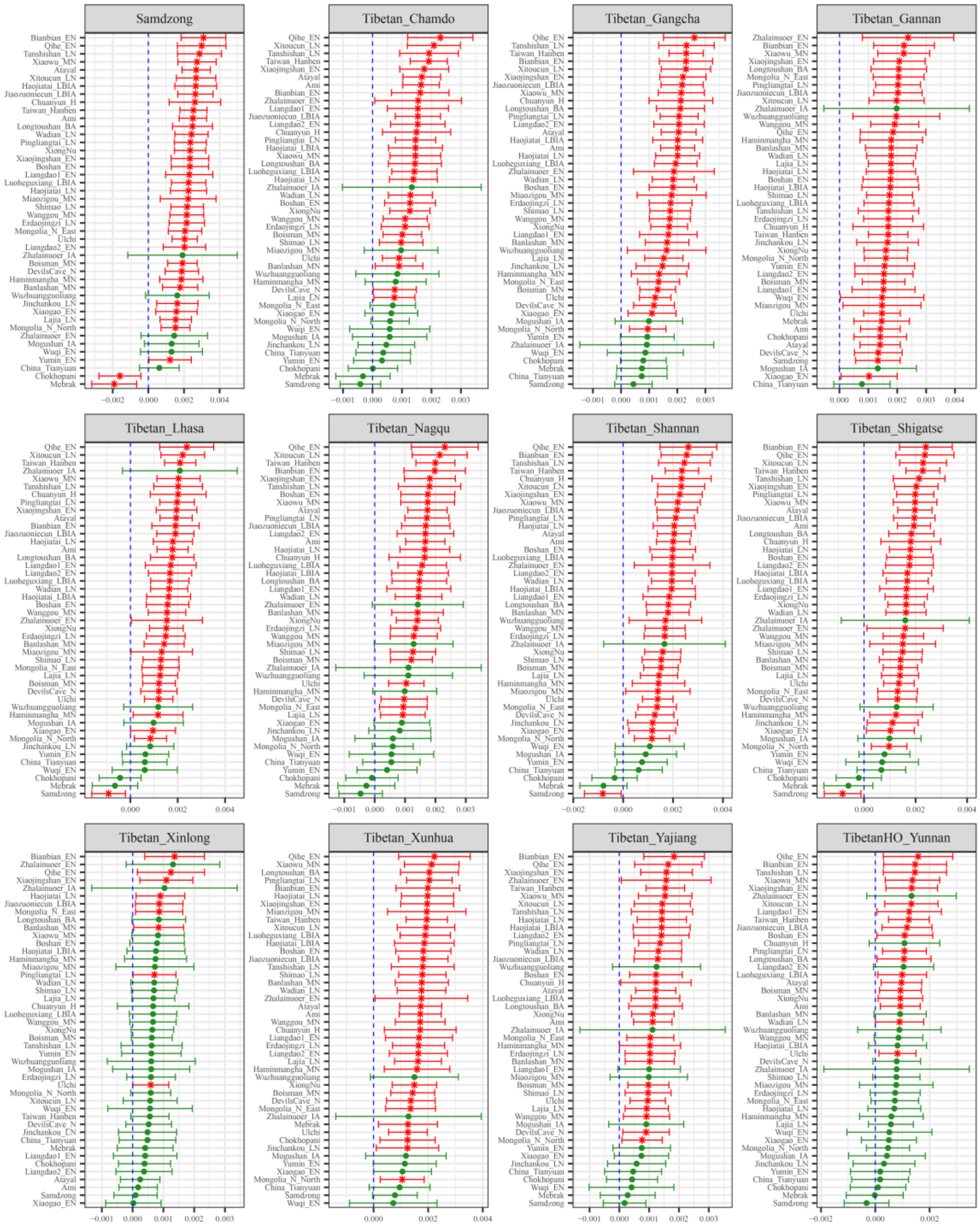

**Figure S78. Results of affinity-$f_4$ statistics for the form $f_4$*(Dacaozi_IA, Modern Tibetan; Neolithic to Historic East Asians, Mbuti)* showed the additional shared derived alleles from source populations except for inland Neolithic northern East Asian from Shaanxi or Inner Mongolia or their related ancestral populations.**

Here, overlapping SNP loci included in the Affymetrix Human Origins platform among four analyzed populations were used. We used the genetic variation of Mbuti as the outgroup. Red asterisk point meant the significant value (Absolute value of Z-scores larger than three or equal to three) observed in the symmetry-$f_4$ statistics and green circle point denoted the non-significant $f_4$-statistic values (Absolute value of Z-scores less than three). All ancient East Asians were listed along the Y-axis and $f_4$ values were labeled along the X-axis. All results were faceted or grouped via Tibetan populations. Significant negative $f_4$ values indicated that the third population shared more alleles with the second population, and

also means Tibetan obtained additional gene flow from the third source population (or related populations). And the significant positive $f_4$ value indicated that the third population shared more derived alleles with the first population. The value of $f_4$-statistics equal to zero was marked as the blue dash line. The bar indicated three standard errors.

**Additional gene flow events when we assumed that Tibetans' direct ancestor is Liao River ancients or their related ancestral populations**

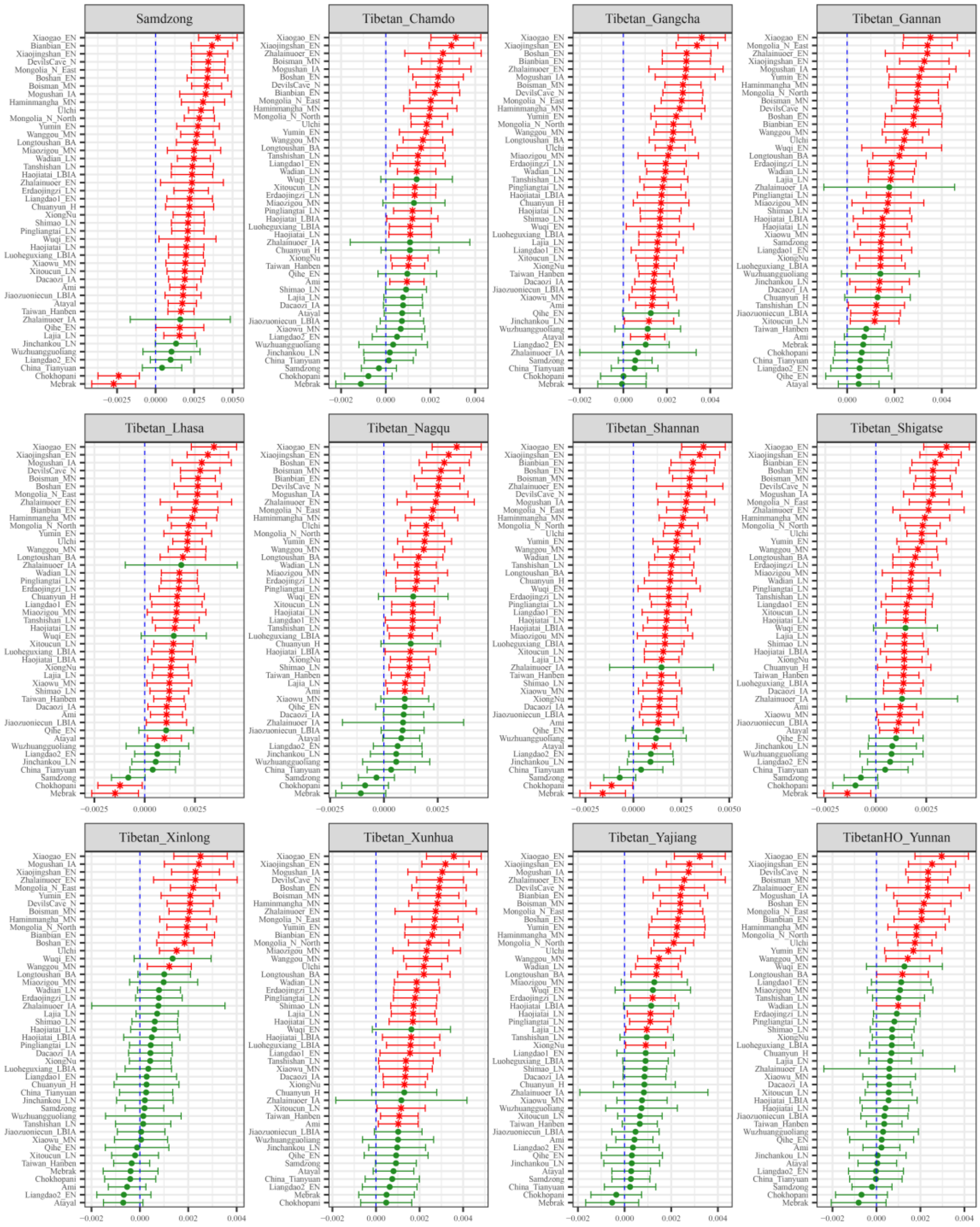

**Figure S79. Results of affinity-$f_4$ statistics for the form $f_4$*(Banhashan_MN, Modern Tibetan; Neolithic to Historic East Asians, Mbuti)* showed the additional shared derived alleles from source populations except for ancestral populations from Liao River Basin.**

Here, overlapping SNP loci included in the Affymetrix Human Origins platform among four analyzed populations were used. We used the genetic variation of Mbuti as the outgroup. Red asterisk point meant the significant value (Absolute value of Z-scores larger than three or equal to three) observed in the symmetry-$f_4$ statistics and green circle point denoted the non-significant $f_4$-statistic values (Absolute

value of Z-scores less than three). All ancient East Asians were listed along the Y-axis and $f_4$ values were labeled along the X-axis. All results were faceted or grouped via Tibetan populations. Significant negative $f_4$ values indicated that the third population shared more alleles with the second population, and also means Tibetan obtained additional gene flow from the third source population (or related populations). And the significant positive $f_4$ value indicated that the third population shared more derived alleles with the first population. The value of $f_4$-statistics equal to zero was marked as the blue dash line. The bar indicated three standard errors.

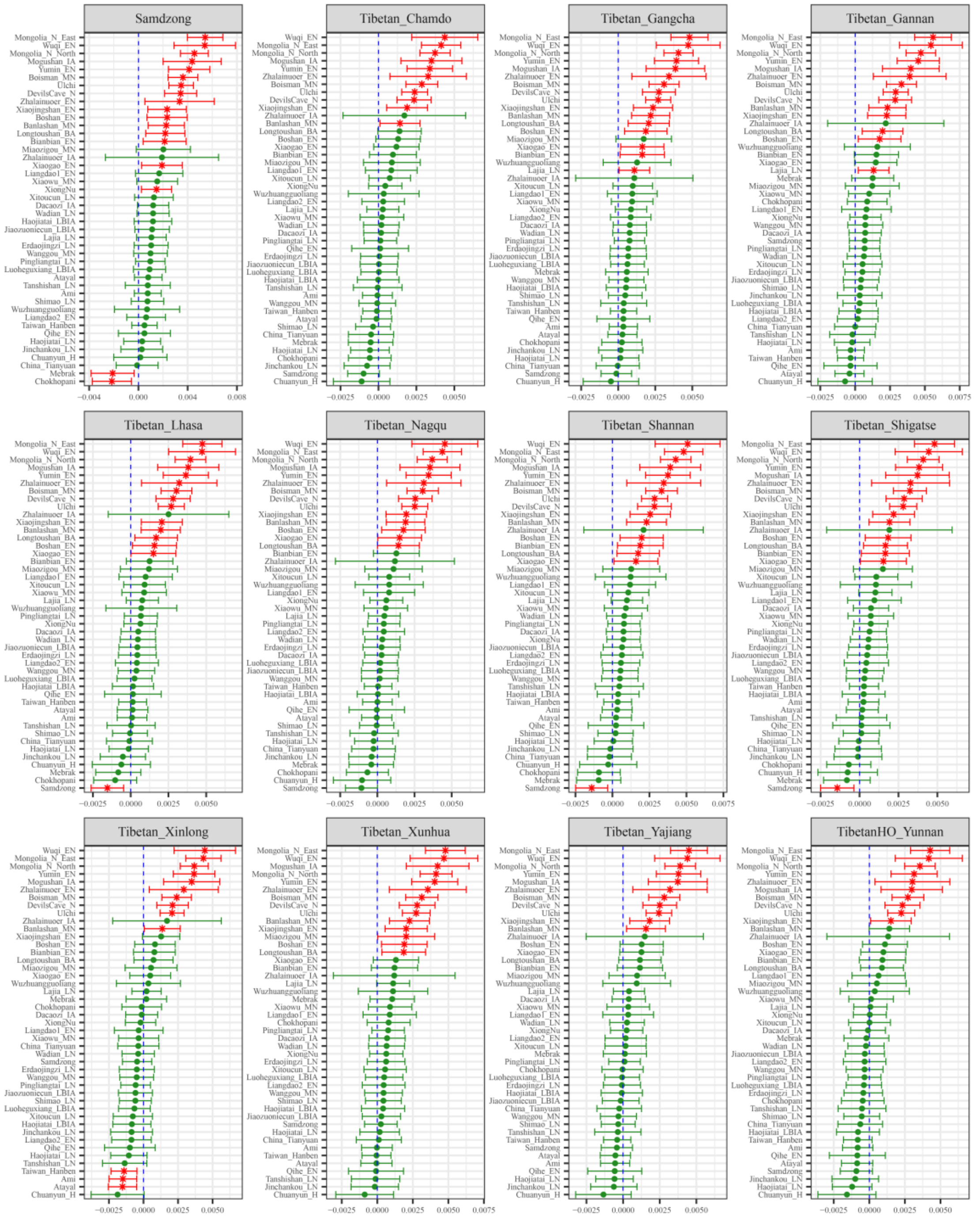

**Figure S80. Results of affinity-$f_4$ statistics for the form $f_4$*(Haminmangha_MN, Modern Tibetan; Neolithic to Historic East Asians, Mbuti)* showed the additional shared derived alleles from source populations except for ancestral populations from Liao River Basin.**

Here, overlapping SNP loci included in the Affymetrix Human Origins platform among four analyzed populations were used. We used the genetic variation of Mbuti as the outgroup. Red asterisk point meant the significant value (Absolute value of Z-scores larger than three or equal to three) observed in the symmetry-$f_4$ statistics and green circle point denoted the non-significant $f_4$-statistic values (Absolute

value of Z-scores less than three). All ancient East Asians were listed along the Y-axis and $f_4$ values were labeled along the X-axis. All results were faceted or grouped via Tibetan populations. Significant negative $f_4$ values indicated that the third population shared more alleles with the second population, and also means Tibetan obtained additional gene flow from the third source population (or related populations). And the significant positive $f_4$ value indicated that the third population shared more derived alleles with the first population. The value of $f_4$-statistics equal to zero was marked as the blue dash line. The bar indicated three standard errors.

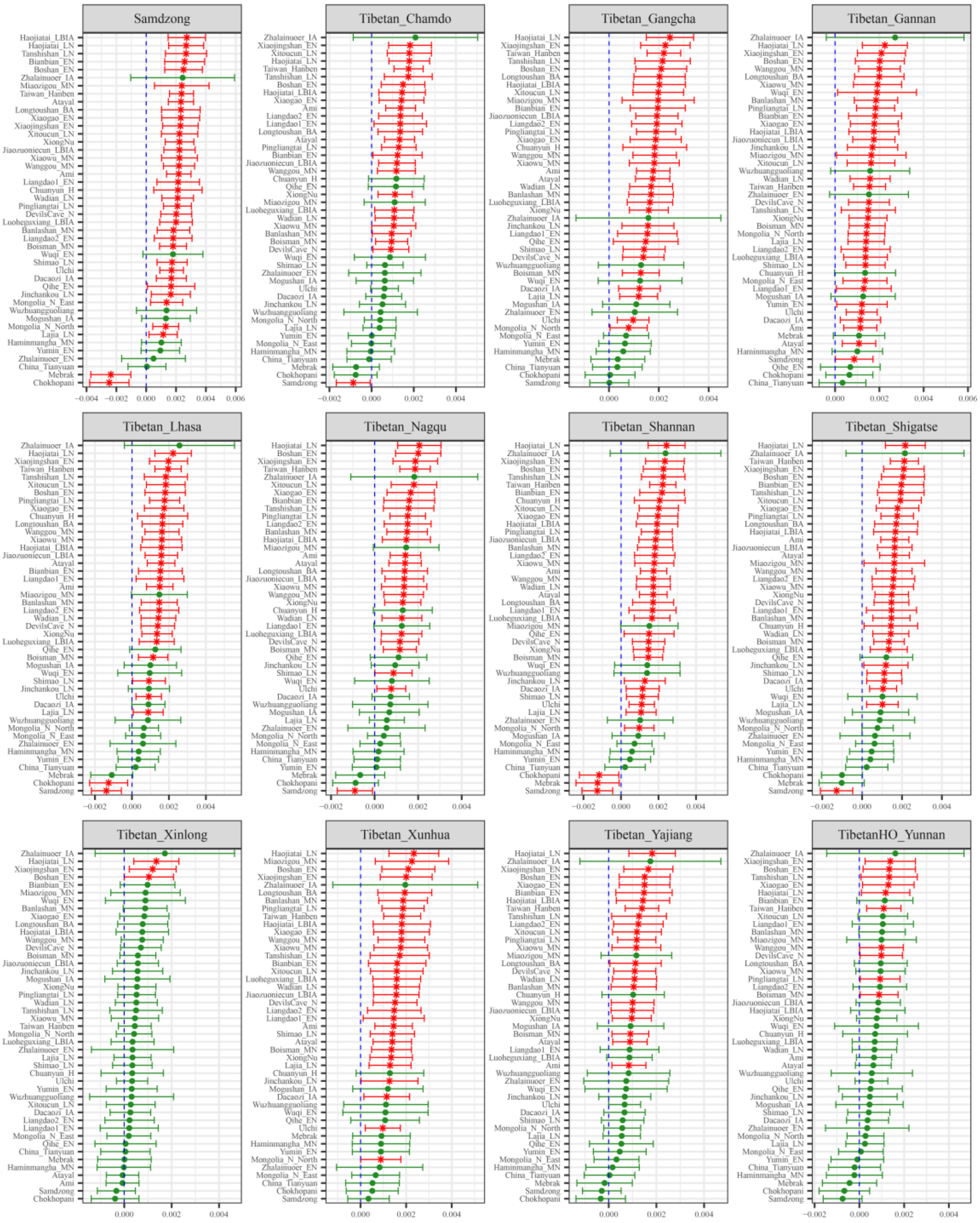

**Figure S81. Results of affinity-$f_4$ statistics for the form $f_4$(Erdaojingzi_LN, Modern Tibetan; Neolithic to Historic East Asians, Mbuti) showed the additional shared derived alleles from source populations except for ancestral populations from Liao River Basin.**

Here, overlapping SNP loci included in the Affymetrix Human Origins platform among four analyzed populations were used. We used the genetic variation of Mbuti as the outgroup. Red asterisk point meant the significant value (Absolute value of Z-scores larger than three or equal to three) observed in the symmetry-$f_4$ statistics and green circle point denoted the non-significant $f_4$-statistic values (Absolute

value of Z-scores less than three). All ancient East Asians were listed along the Y-axis and $f_4$ values were labeled along the X-axis. All results were faceted or grouped via Tibetan populations. Significant negative $f_4$ values indicated that the third population shared more alleles with the second population, and also means Tibetan obtained additional gene flow from the third source population (or related populations). And the significant positive $f_4$ value indicated that the third population shared more derived alleles with the first population. The value of $f_4$-statistics equal to zero was marked as the blue dash line. The bar indicated three standard errors.

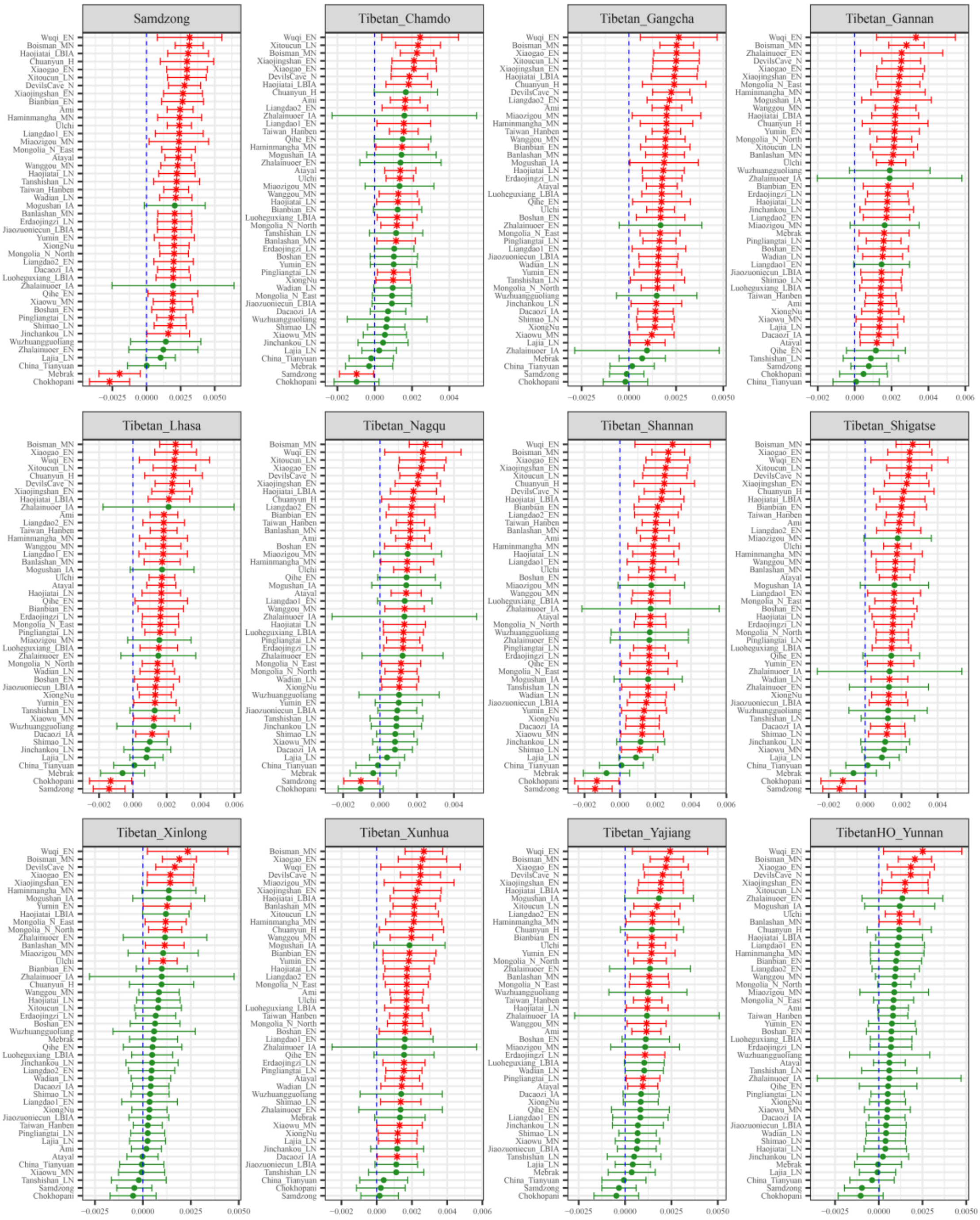

**Figure S82. Results of affinity-$f_4$ statistics for the form $f_4$*(Longtoushan_BA, Modern Tibetan; Neolithic to Historic East Asians, Mbuti)* showed the additional shared derived alleles from source populations except for ancestral populations from Liao River Basin.**

Here, overlapping SNP loci included in the Affymetrix Human Origins platform among four analyzed populations were used. We used the genetic variation of Mbuti as the outgroup. Red asterisk point meant the significant value (Absolute value of Z-scores larger than three or equal to three) observed in the symmetry-$f_4$ statistics and green circle point denoted the non-significant $f_4$-statistic values (Absolute

value of Z-scores less than three). All ancient East Asians were listed along the Y-axis and $f_4$ values were labeled along the X-axis. All results were faceted or grouped via Tibetan populations. Significant negative $f_4$ values indicated that the third population shared more alleles with the second population, and also means Tibetan obtained additional gene flow from the third source population (or related populations). And the significant positive $f_4$ value indicated that the third population shared more derived alleles with the first population. The value of $f_4$-statistics equal to zero was marked as the blue dash line. The bar indicated three standard errors.

**Additional gene flow events when we assumed that Tibetans' direct ancestor is ancestral populations from Amur River Basin**

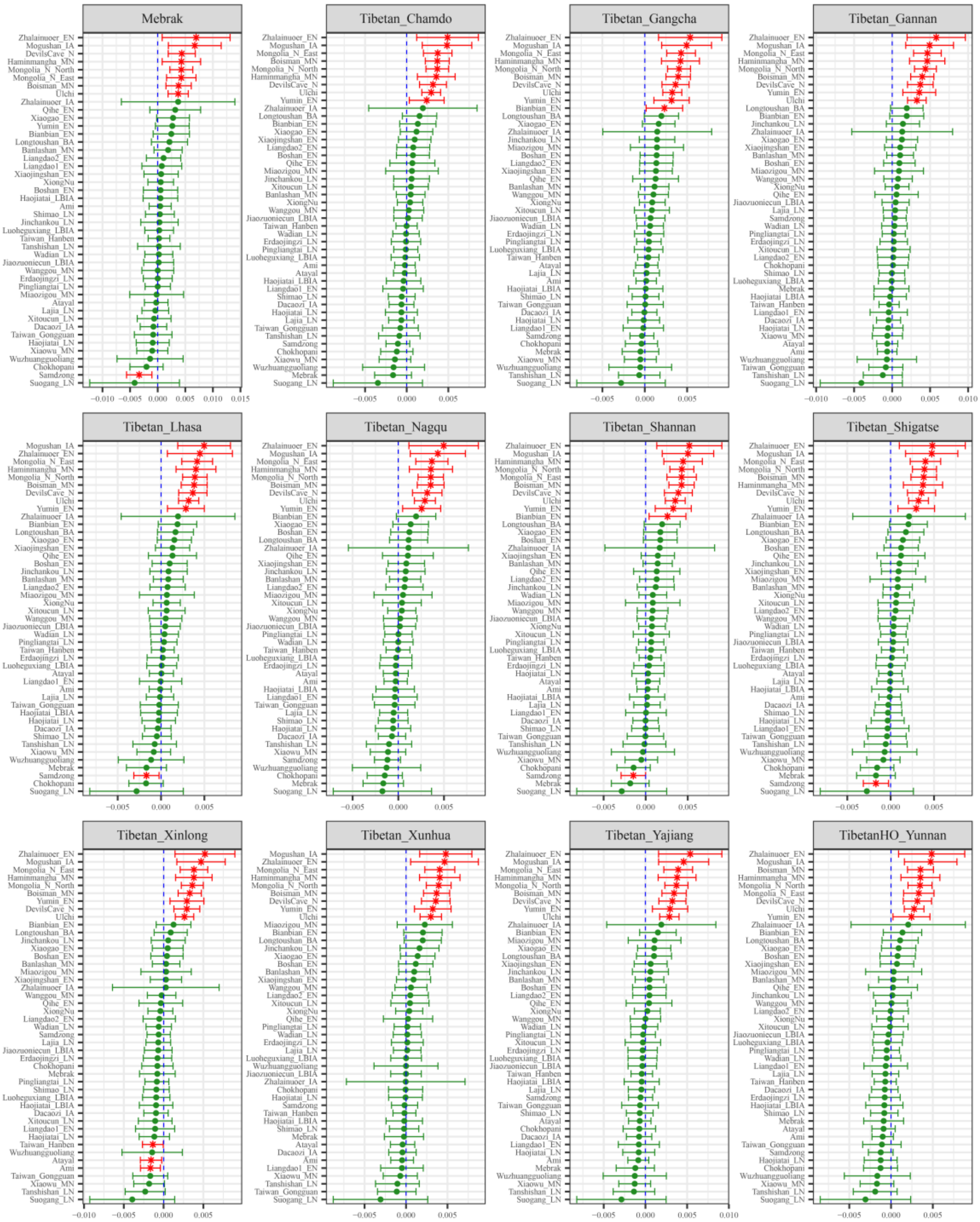

**Figure S83. Results of affinity-$f_4$ statistics for the form $f_4$*(Wuqi_EN, Modern Tibetan; Neolithic to Historic East Asians, Mbuti)* showed the additional shared derived alleles from source populations except for ancestral populations from Amur River Basin.**

Here, overlapping SNP loci included in the Affymetrix Human Origins platform among four analyzed populations were used. We used the genetic variation of Mbuti as the outgroup. Red asterisk point meant the significant value (Absolute value of Z-scores larger than three or equal to three) observed in the symmetry-$f_4$ statistics and green circle point denoted the non-significant $f_4$-statistic values (Absolute value of Z-scores less than three). All ancient East Asians were listed along the Y-axis and $f_4$ values were labeled along the X-axis. All results were faceted or grouped via Tibetan populations. Significant negative $f_4$ values indicated that the third population shared more alleles with the second population, and also means Tibetan obtained additional gene flow from the third source population (or related populations). And the significant positive $f_4$ value indicated that the third population shared more derived alleles with the first population. The value of $f_4$-statistics equal to zero was marked as the blue dash line.

The bar indicated three standard errors.

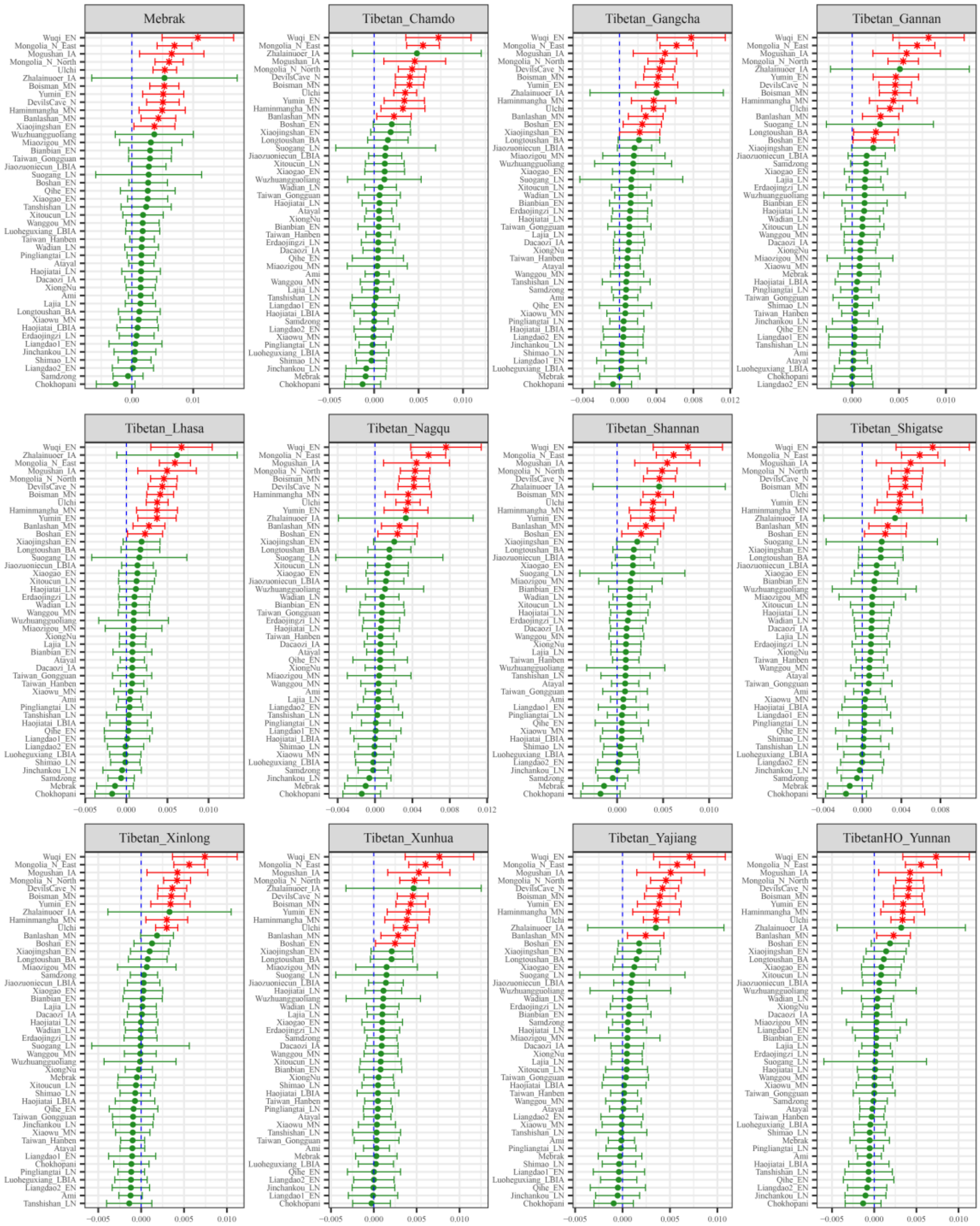

**Figure S84. Results of affinity-$f_4$ statistics for the form $f_4$(*Zhalainuoer_EN, Modern Tibetan; Neolithic to Historic East Asians, Mbuti*) showed the additional shared derived alleles from source populations except for ancestral populations from Amur River Basin.**

Here, overlapping SNP loci included in the Affymetrix Human Origins platform among four analyzed populations were used. We used the genetic variation of Mbuti as the outgroup. Red asterisk point meant the significant value (Absolute value of Z-scores larger than three or equal to three) observed in the symmetry-$f_4$ statistics and green circle point denoted the non-significant $f_4$-statistic values (Absolute value of Z-scores less than three). All ancient East Asians were listed along the Y-axis and $f_4$ values were labeled along the X-axis. All results were faceted or grouped via Tibetan populations. Significant negative $f_4$ values indicated that the third population shared more alleles with the second population, and also means Tibetan obtained additional gene flow from the third source population (or related populations). And the significant positive $f_4$ value indicated that the third population shared more derived alleles with the first population. The value of $f_4$-statistics equal to zero was marked as the blue dash line.

The bar indicated three standard errors.

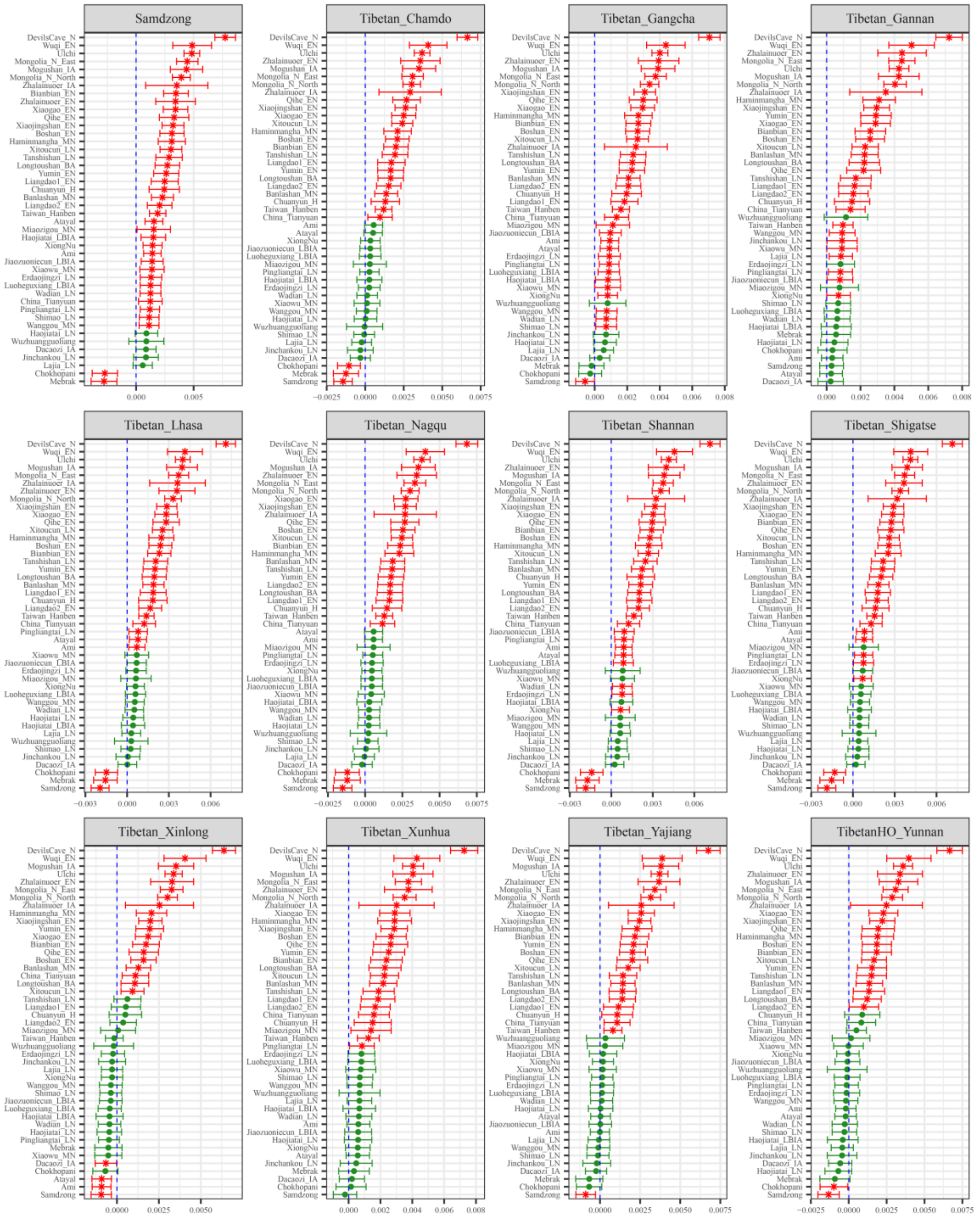

**Figure S85. Results of affinity-$f_4$ statistics for the form $f_4$(*Boisman_MN, Modern Tibetan; Neolithic to Historic East Asians, Mbuti*) showed the additional shared derived alleles from source populations except for ancestral populations from Amur River Basin.**
Here, overlapping SNP loci included in the Affymetrix Human Origins platform among four analyzed populations were used. We used the genetic variation of Mbuti as the outgroup. Red asterisk point meant the significant value (Absolute value of Z-scores larger than three or equal to three) observed in the symmetry-$f_4$ statistics and green circle point denoted the non-significant $f_4$-statistic values (Absolute value of Z-scores less than three). All ancient East Asians were listed along the Y-axis and $f_4$ values were labeled along the X-axis. All results were faceted or grouped via Tibetan populations. Significant negative $f_4$ values indicated that the third population shared more alleles with the second population, and also means Tibetan obtained additional gene flow from the third source population (or related populations). And the significant positive $f_4$ value indicated that the third population shared more derived alleles with the first population. The value of $f_4$-statistics equal to zero was marked as the blue dash line.

The bar indicated three standard errors.

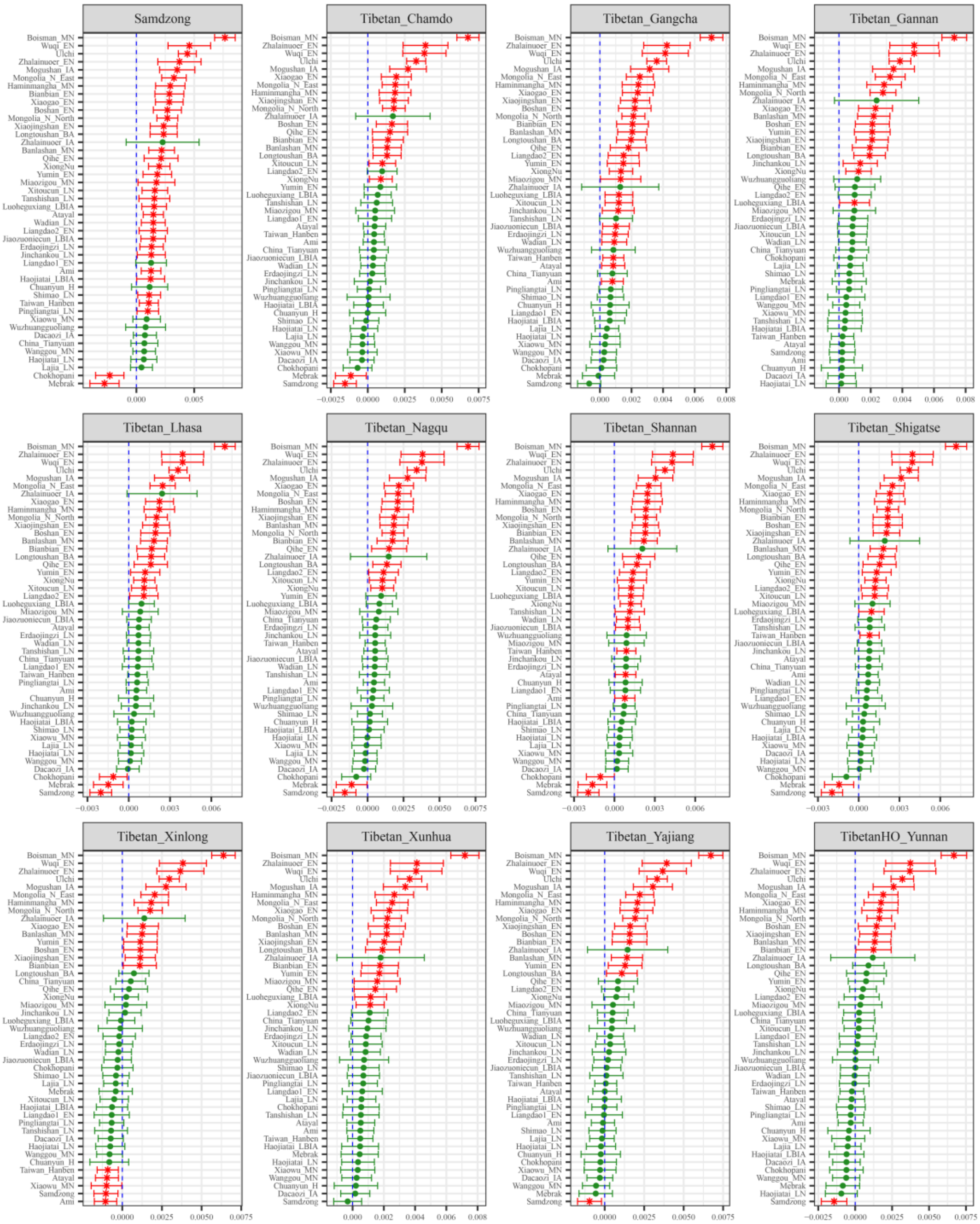

**Figure S86. Results of affinity-$f_4$ statistics for the form $f_4$*(DevilsCave_N, Modern Tibetan; Neolithic to Historic East Asians, Mbuti)* showed the additional shared derived alleles from source populations except for ancestral populations from Amur River Basin.**

Here, overlapping SNP loci included in the Affymetrix Human Origins platform among four analyzed populations were used. We used the genetic variation of Mbuti as the outgroup. Red asterisk point meant the significant value (Absolute value of Z-scores larger than three or equal to three) observed in the symmetry-$f_4$ statistics and green circle point denoted the non-significant $f_4$-statistic values (Absolute value of Z-scores less than three). All ancient East Asians were listed along the Y-axis and $f_4$ values were labeled along the X-axis. All results were faceted or grouped via Tibetan populations. Significant negative $f_4$ values indicated that the third population shared more alleles with the second population, and also means Tibetan obtained additional gene flow from the third source population (or related populations). And the significant positive $f_4$ value indicated that the third population shared more derived alleles with the first population. The value of $f_4$-statistics equal to zero was marked as the blue dash line.

The bar indicated three standard errors.

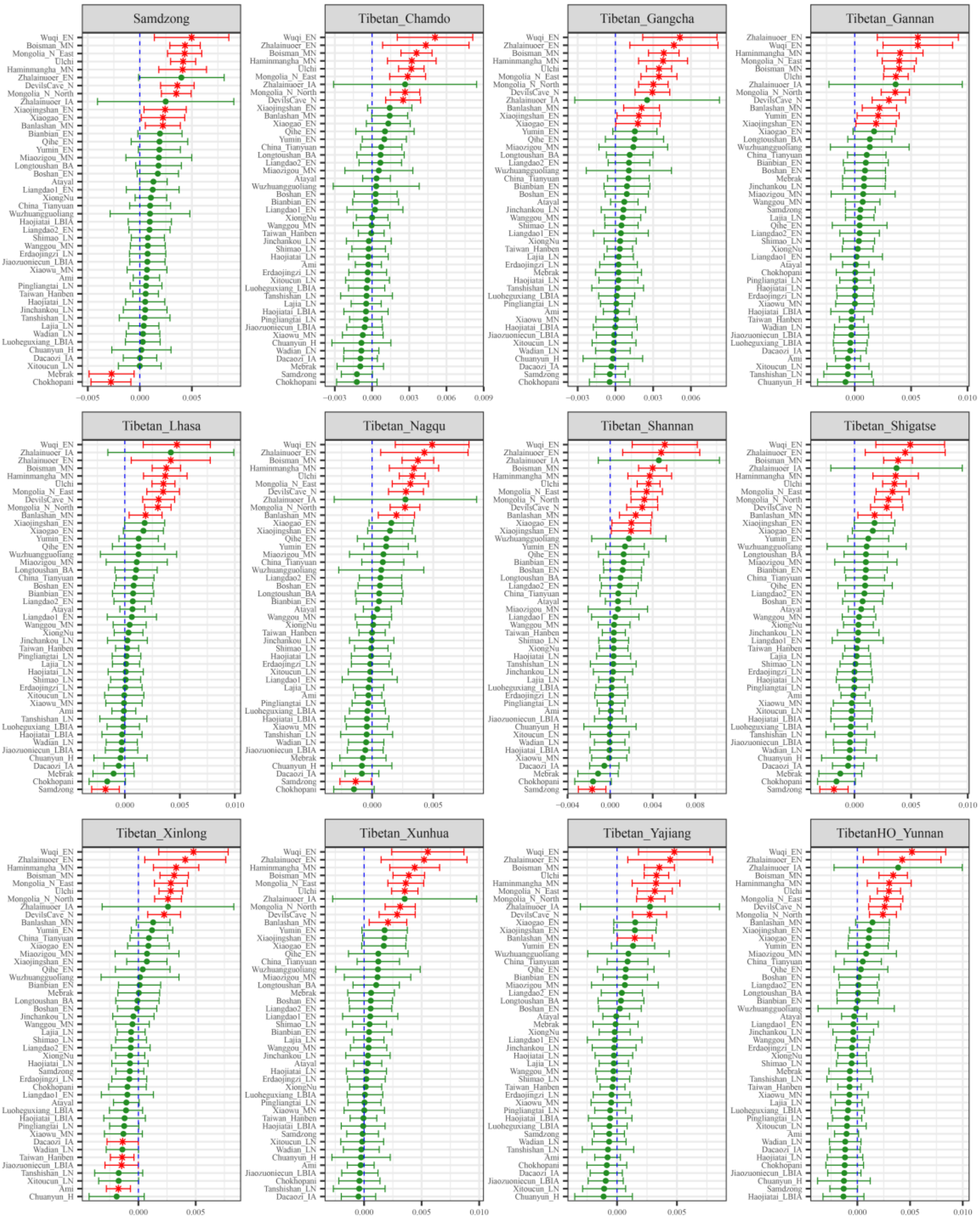

**Figure S87. Results of affinity-$f_4$ statistics for the form $f_4$*(Mogushan_IA, Modern Tibetan; Neolithic to Historic East Asians, Mbuti)* showed the additional shared derived alleles from source populations except for ancestral populations from Amur River Basin.**

Here, overlapping SNP loci included in the Affymetrix Human Origins platform among four analyzed populations were used. We used the genetic variation of Mbuti as the outgroup. Red asterisk point meant the significant value (Absolute value of Z-scores larger than three or equal to three) observed in the symmetry-$f_4$ statistics and green circle point denoted the non-significant $f_4$-statistic values (Absolute value of Z-scores less than three). All ancient East Asians were listed along the Y-axis and $f_4$ values were labeled along the X-axis. All results were faceted or grouped via Tibetan populations. Significant negative $f_4$ values indicated that the third population shared more alleles with the second population, and also means Tibetan obtained additional gene flow from the third source population (or related populations). And the significant positive $f_4$ value indicated that the third population shared more derived alleles with the first population. The value of $f_4$-statistics equal to zero was marked as the blue dash line.

The bar indicated three standard errors.

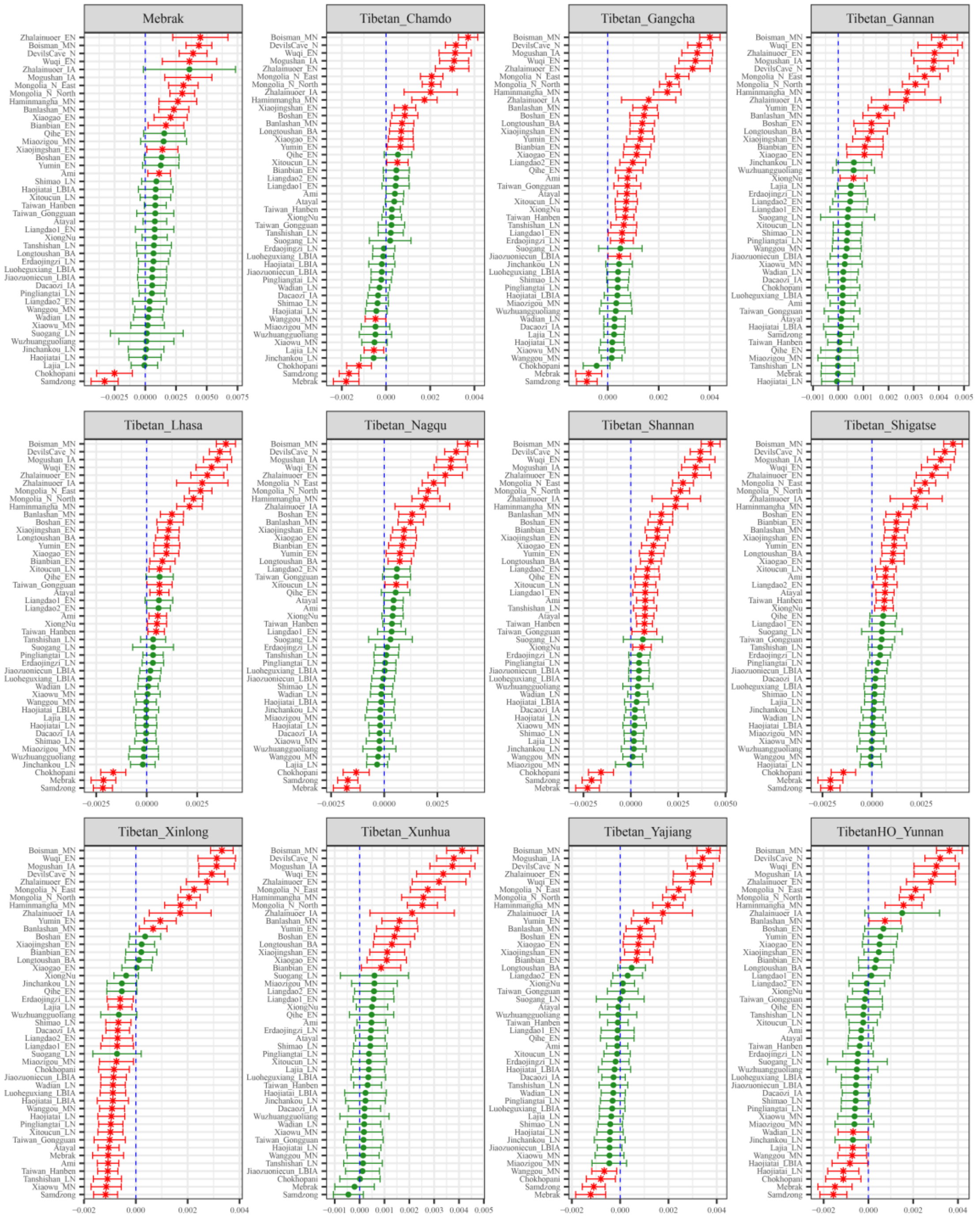

**Figure S88. Results of affinity-$f_4$ statistics for the form $f_4$*(Ulchi, Modern Tibetan; Neolithic to Historic East Asians, Mbuti)* showed the additional shared derived alleles from source populations except for ancestral populations from Amur River Basin.**

Here, overlapping SNP loci included in the Affymetrix Human Origins platform among four analyzed populations were used. We used the genetic variation of Mbuti as the outgroup. Red asterisk point meant the significant value (Absolute value of Z-scores larger than three or equal to three) observed in the symmetry-$f_4$ statistics and green circle point denoted the non-significant $f_4$-statistic values (Absolute value of Z-scores less than three). All ancient East Asians were listed along the Y-axis and $f_4$ values were labeled along the X-axis. All results were faceted or grouped via Tibetan populations. Significant negative $f_4$ values indicated that the third population shared more alleles with the second population, and also means Tibetan obtained additional gene flow from the third source population (or related populations). And the significant positive $f_4$ value indicated that the third population shared more derived alleles with the first population. The value of $f_4$-statistics equal to zero was marked as the blue dash line.

The bar indicated three standard errors.

**Additional gene flow events when we assumed that Tibetans' direct ancestor is ancestral populations from Nepal**

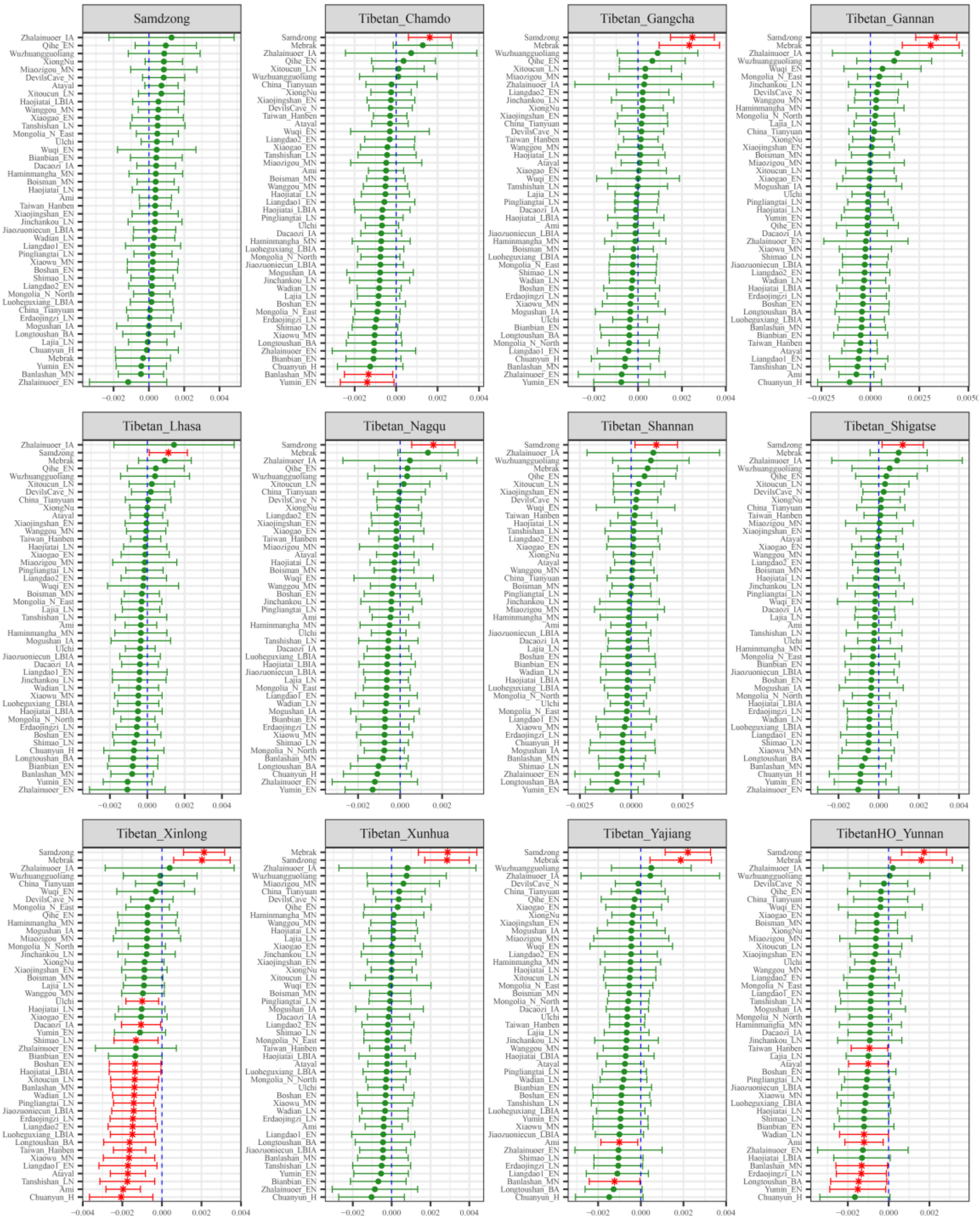

**Figure S89. Results of affinity-$f_4$ statistics for the form $f_4$*(Chokhopani, Modern Tibetan; Neolithic to Historic East Asians, Mbuti)* showed the additional shared derived alleles from source populations except for ancestral populations from Nepal.**

Here, overlapping SNP loci included in the Affymetrix Human Origins platform among four analyzed populations were used. We used the genetic variation of Mbuti as the outgroup. Red asterisk point meant the significant value (Absolute value of Z-scores larger than three or equal to three) observed in the symmetry-$f_4$ statistics and green circle point denoted the non-significant $f_4$-statistic values (Absolute value of Z-scores less than three). All ancient East Asians were listed along the Y-axis and $f_4$ values were labeled along the X-axis. All results were faceted or grouped via Tibetan populations. Significant negative $f_4$ values indicated that the third population shared more alleles with the second population, and also means Tibetan obtained additional gene flow from the third source population (or related

populations). And the significant positive $f_4$ value indicated that the third population shared more derived alleles with the first population. The value of $f_4$-statistics equal to zero was marked as the blue dash line. The bar indicated three standard errors.

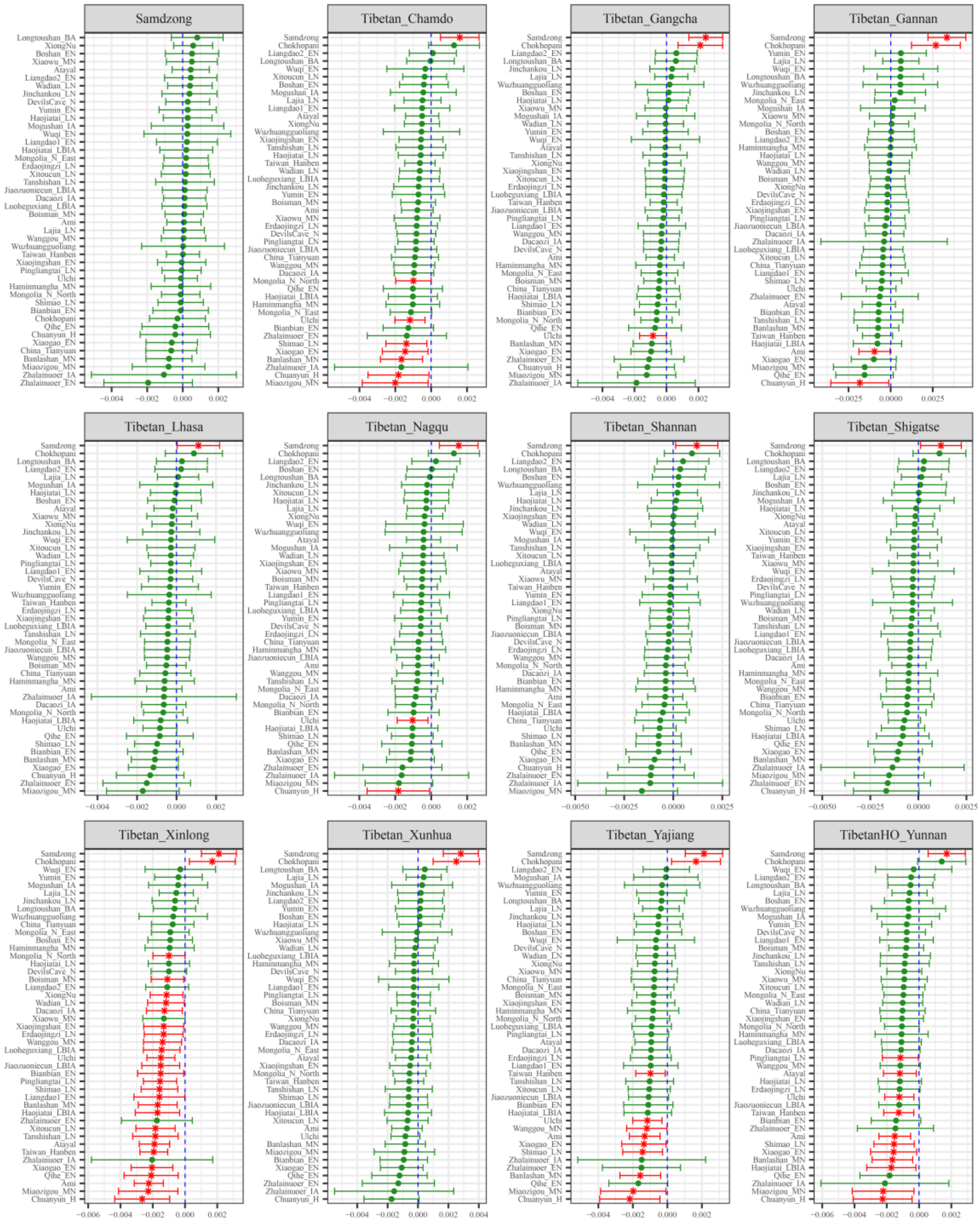

**Figure S90. Results of affinity-$f_4$ statistics for the form $f_4$*(Mebrak, Modern Tibetan; Neolithic to Historic East Asians, Mbuti)* showed the additional shared derived alleles from source populations except for ancestral populations from Nepal.**

Here, overlapping SNP loci included in the Affymetrix Human Origins platform among four analyzed populations were used. We used the genetic variation of Mbuti as the outgroup. Red asterisk point meant the significant value (Absolute value of Z-scores larger than three or equal to three) observed in the symmetry-$f_4$ statistics and green circle point denoted the non-significant $f_4$-statistic values (Absolute value of Z-scores less than three). All ancient East Asians were listed along the Y-axis and $f_4$ values were labeled along the X-axis. All results were faceted or grouped via Tibetan populations. Significant negative $f_4$ values indicated that the third population shared more alleles with the second population, and also means Tibetan obtained additional gene flow from the third source population (or related

populations). And the significant positive $f_4$ value indicated that the third population shared more derived alleles with the first population. The value of $f_4$-statistics equal to zero was marked as the blue dash line. The bar indicated three standard errors.

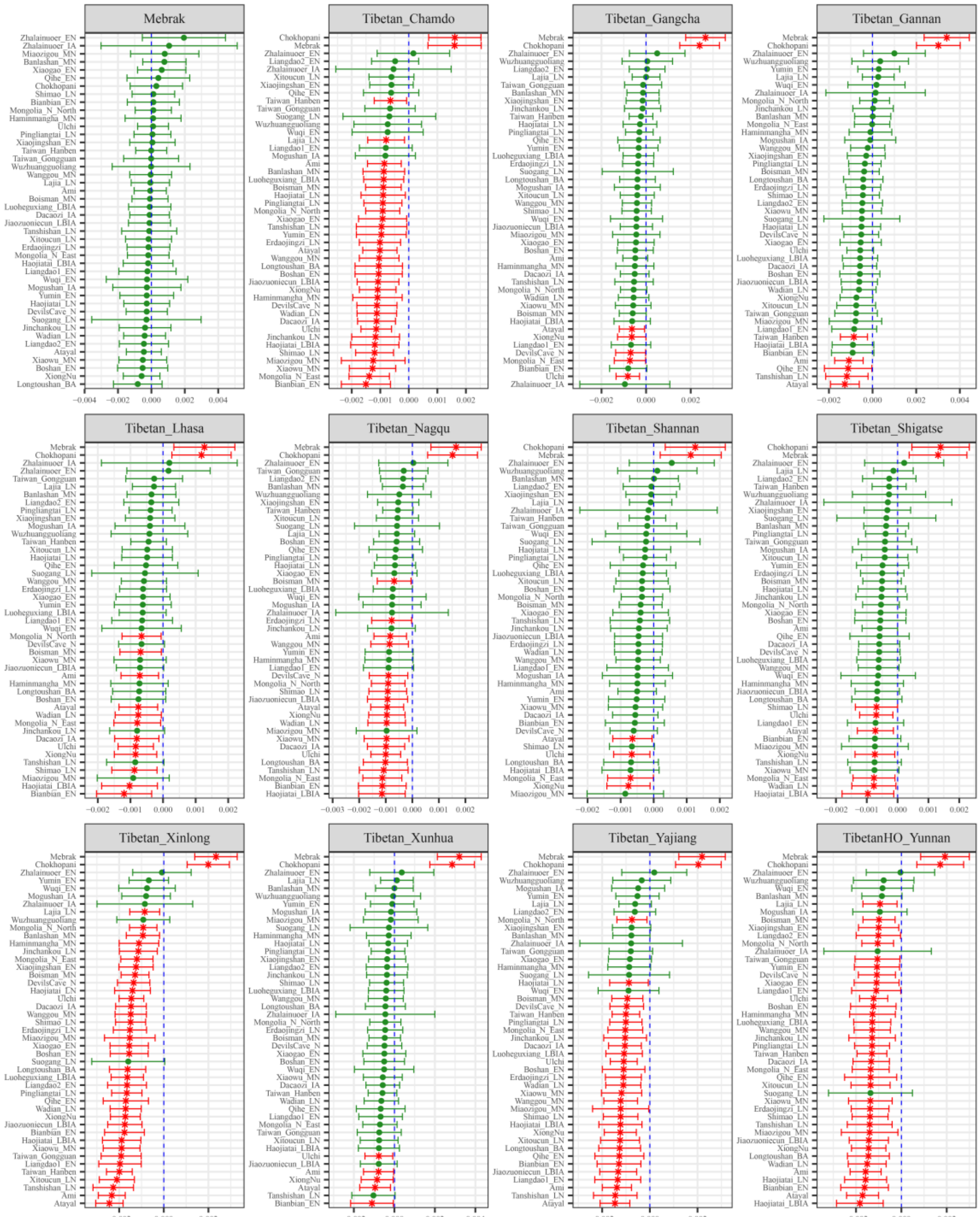

**Figure S91. Results of affinity-$f_4$ statistics for the form $f_4$*(Samdzong, Modern Tibetan; Neolithic to Historic East Asians, Mbuti)* showed the additional shared derived alleles from source populations except for ancestral populations from Nepal.**

Here, overlapping SNP loci included in the Affymetrix Human Origins platform among four analyzed populations were used. We used the genetic variation of Mbuti as the outgroup. Red asterisk point meant the significant value (Absolute value of Z-scores larger than three or equal to three) observed in the symmetry-$f_4$ statistics and green circle point denoted the non-significant $f_4$-statistic values (Absolute value of Z-scores less than three). All ancient East Asians were listed along the Y-axis and $f_4$ values were labeled along the X-axis. All results were faceted or grouped via Tibetan populations. Significant negative $f_4$ values indicated that the third population shared more alleles with the second population, and also means Tibetan obtained additional gene flow from the third source population (or related

populations). And the significant positive $f_4$ value indicated that the third population shared more derived alleles with the first population. The value of $f_4$-statistics equal to zero was marked as the blue dash line. The bar indicated three standard errors.

**Additional gene flow events when we assumed that Tibetans' direct ancestor is ancestral populations from Mongolia Plateau or Baikal Lake Region**

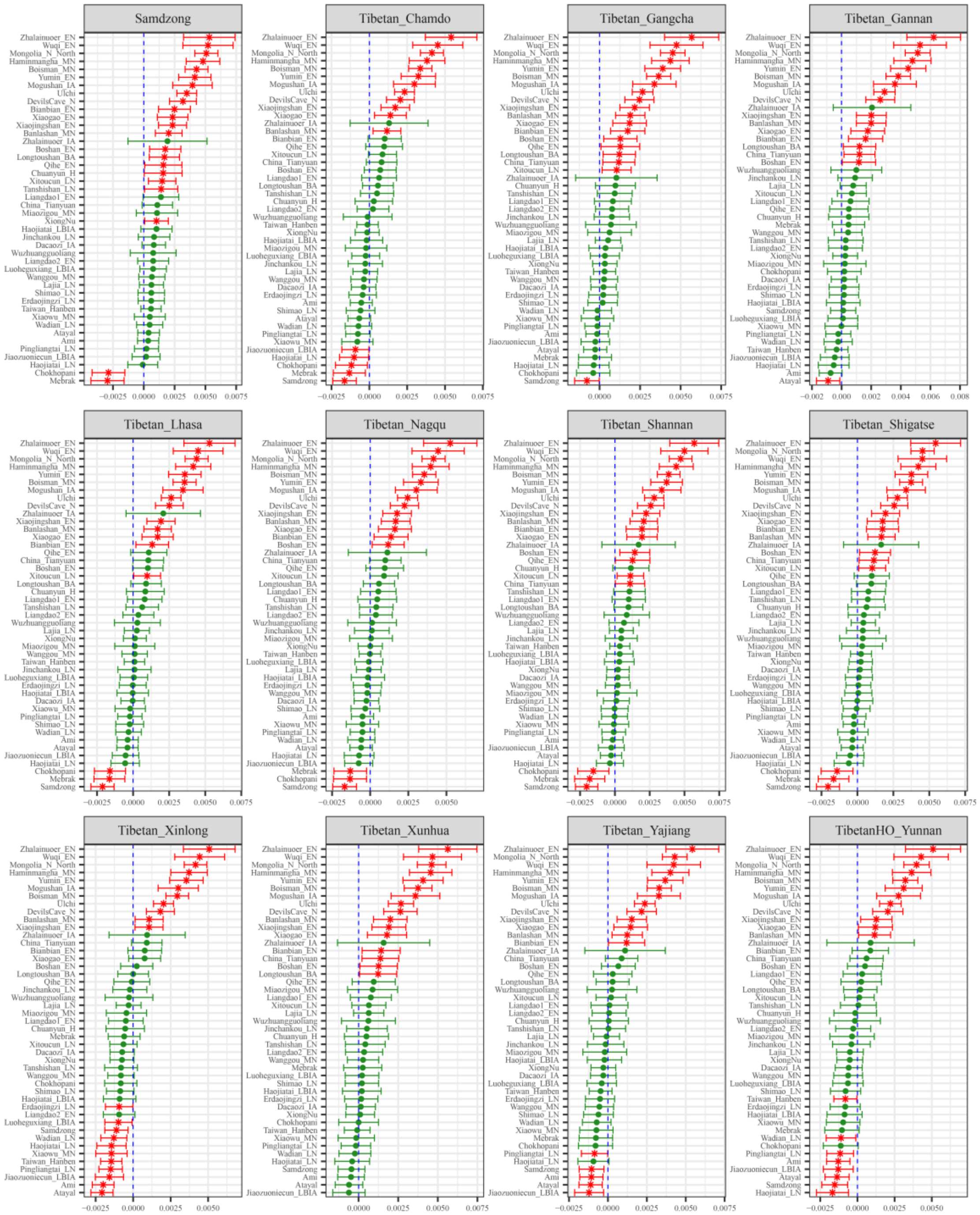

**Figure S92. Results of affinity-$f_4$ statistics for the form $f_4$*(Mongolia_N_East, Modern Tibetan; Neolithic to Historic East Asians, Mbuti)* showed the additional shared derived alleles from source populations except for ancestral populations from Mongolia Plateau or Baikal Lake Region.** Here, overlapping SNP loci included in the Affymetrix Human Origins platform among four analyzed populations were used. We used the genetic variation of Mbuti as the outgroup. Red asterisk point meant the significant value (Absolute value of Z-scores larger than three or equal to three) observed in the symmetry-$f_4$ statistics and green circle point denoted the non-significant $f_4$-statistic values (Absolute value of Z-scores less than three). All ancient East Asians were listed along the Y-axis and $f_4$ values were labeled along the X-axis. All results were faceted or grouped via Tibetan populations. Significant negative $f_4$ values indicated that the third population shared more alleles with the second population, and also means Tibetan obtained additional gene flow from the third source population (or related populations). And the significant positive $f_4$ value indicated that the third population shared more derived alleles with the first population. The value of $f_4$-statistics equal to zero was marked as the blue dash line.

The bar indicated three standard errors.

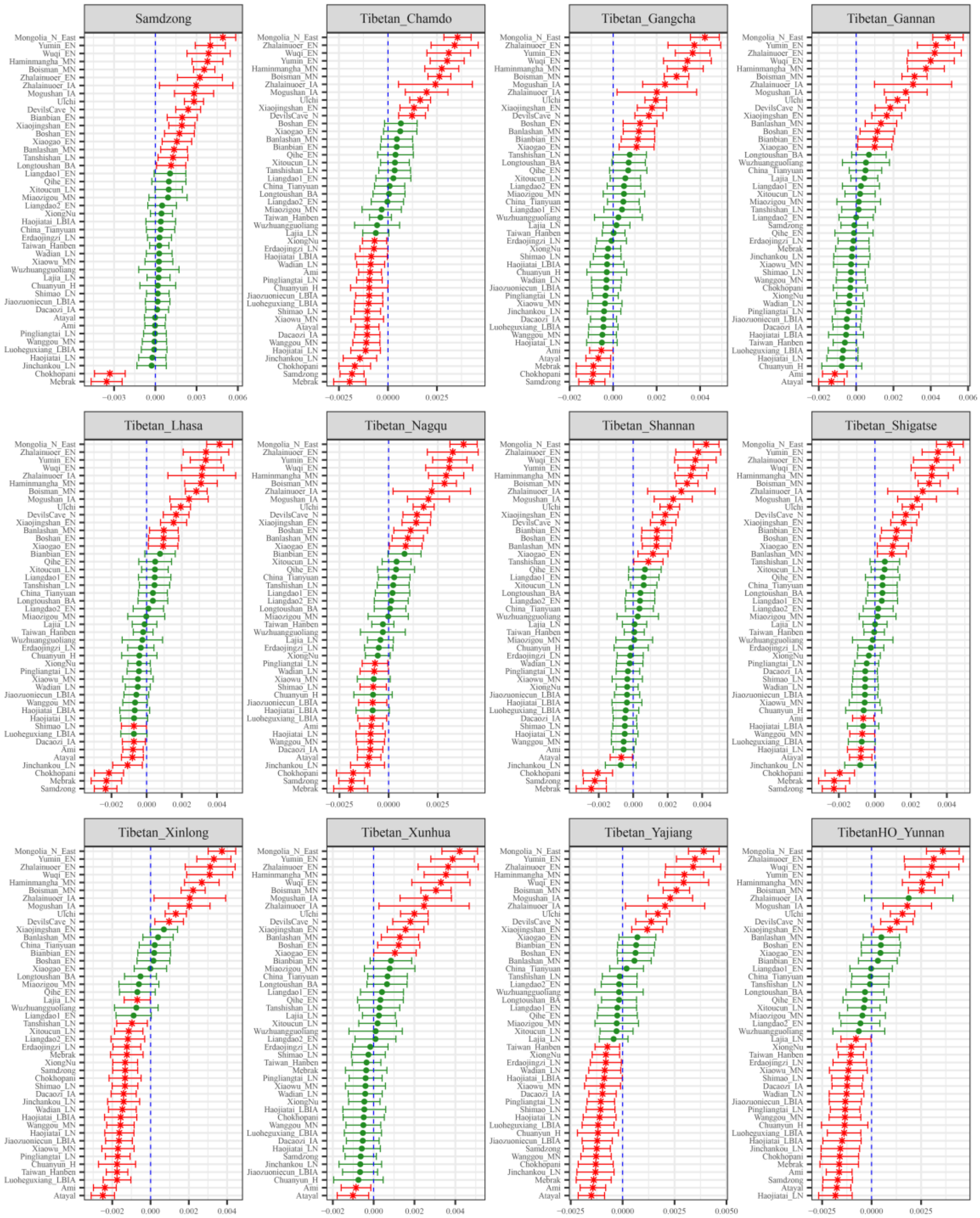

**Figure S93. Results of affinity-$f_4$ statistics for the form $f_4$*(Mongolia_N_North, Modern Tibetan; Neolithic to Historic East Asians, Mbuti)* showed the additional shared derived alleles from source populations except for ancestral populations from Mongolia Plateau or Baikal Lake Region.**
Here, overlapping SNP loci included in the Affymetrix Human Origins platform among four analyzed populations were used. We used the genetic variation of Mbuti as the outgroup. Red asterisk point meant the significant value (Absolute value of Z-scores larger than three or equal to three) observed in the symmetry-$f_4$ statistics and green circle point denoted the non-significant $f_4$-statistic values (Absolute value of Z-scores less than three). All ancient East Asians were listed along the Y-axis and $f_4$ values were labeled along the X-axis. All results were faceted or grouped via Tibetan populations. Significant negative $f_4$ values indicated that the third population shared more alleles with the second population, and also means Tibetan obtained additional gene flow from the third source population (or related populations). And the significant positive $f_4$ value indicated that the third population shared more derived alleles with the first population. The value of $f_4$-statistics equal to zero was marked as the blue dash line.

The bar indicated three standard errors.

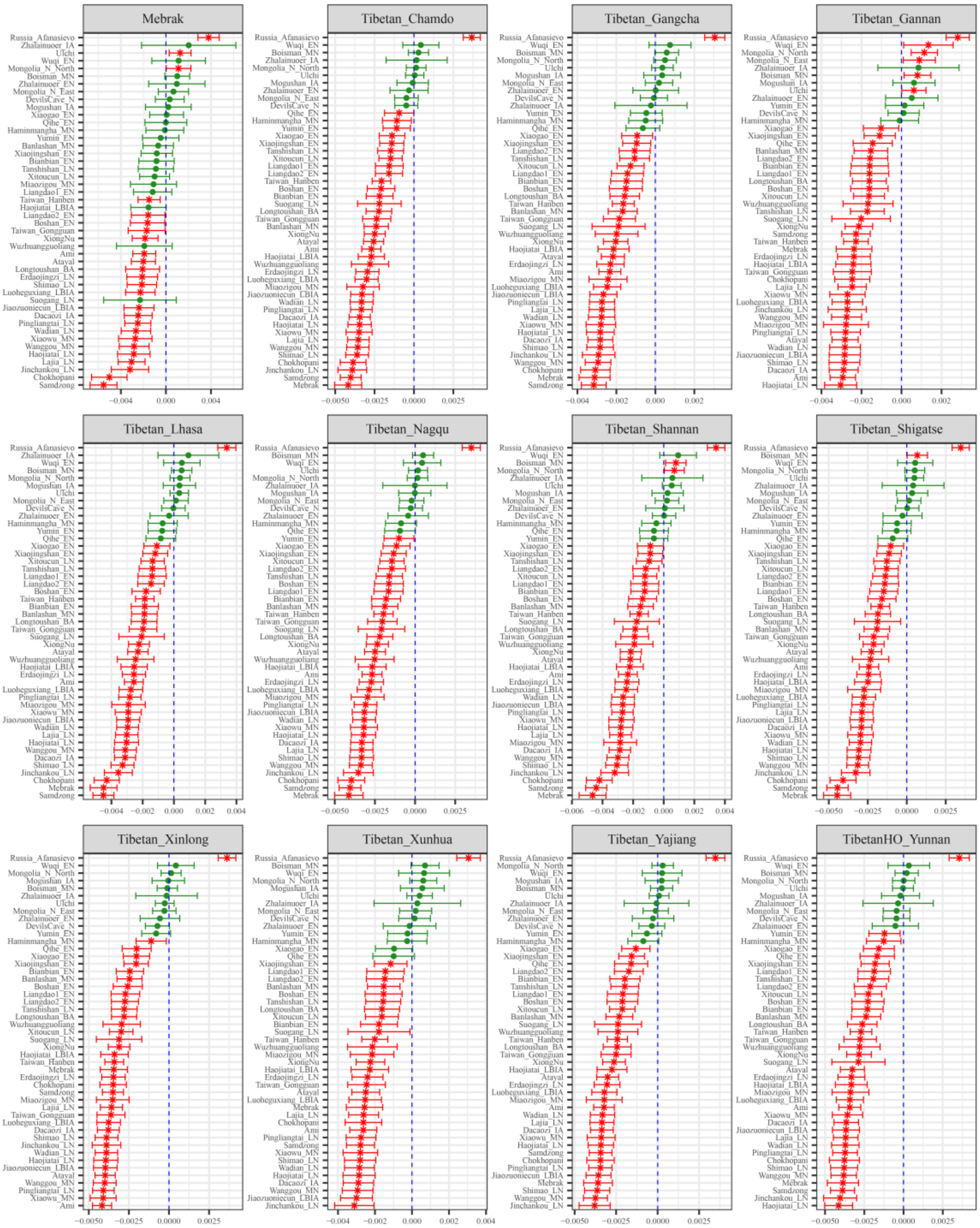

**Figure S94. Results of affinity-$f_4$ statistics for the form $f_4$*(Russia_OldBeringSea_Ekven, Modern Tibetan; Neolithic to Historic East Asians, Mbuti)* showed the additional shared derived alleles from source populations except for ancestral populations from Mongolia Plateau or Baikal Lake Region.** Here, overlapping SNP loci included in the Affymetrix Human Origins platform among four analyzed populations were used. We used the genetic variation of Mbuti as the outgroup. Red asterisk point meant the significant value (Absolute value of Z-scores larger than three or equal to three) observed in the symmetry-$f_4$ statistics and green circle point denoted the non-significant $f_4$-statistic values (Absolute value of Z-scores less than three). All ancient East Asians were listed along the Y-axis and $f_4$ values were labeled along the X-axis. All results were faceted or grouped via Tibetan populations. Significant negative $f_4$ values indicated that the third population shared more alleles with the second population, and also means Tibetan obtained additional gene flow from the third source population (or related populations). And the significant positive $f_4$ value indicated that the third population shared more derived alleles with the first population. The value of $f_4$-statistics equal to zero was marked as the blue dash line.

The bar indicated three standard errors.

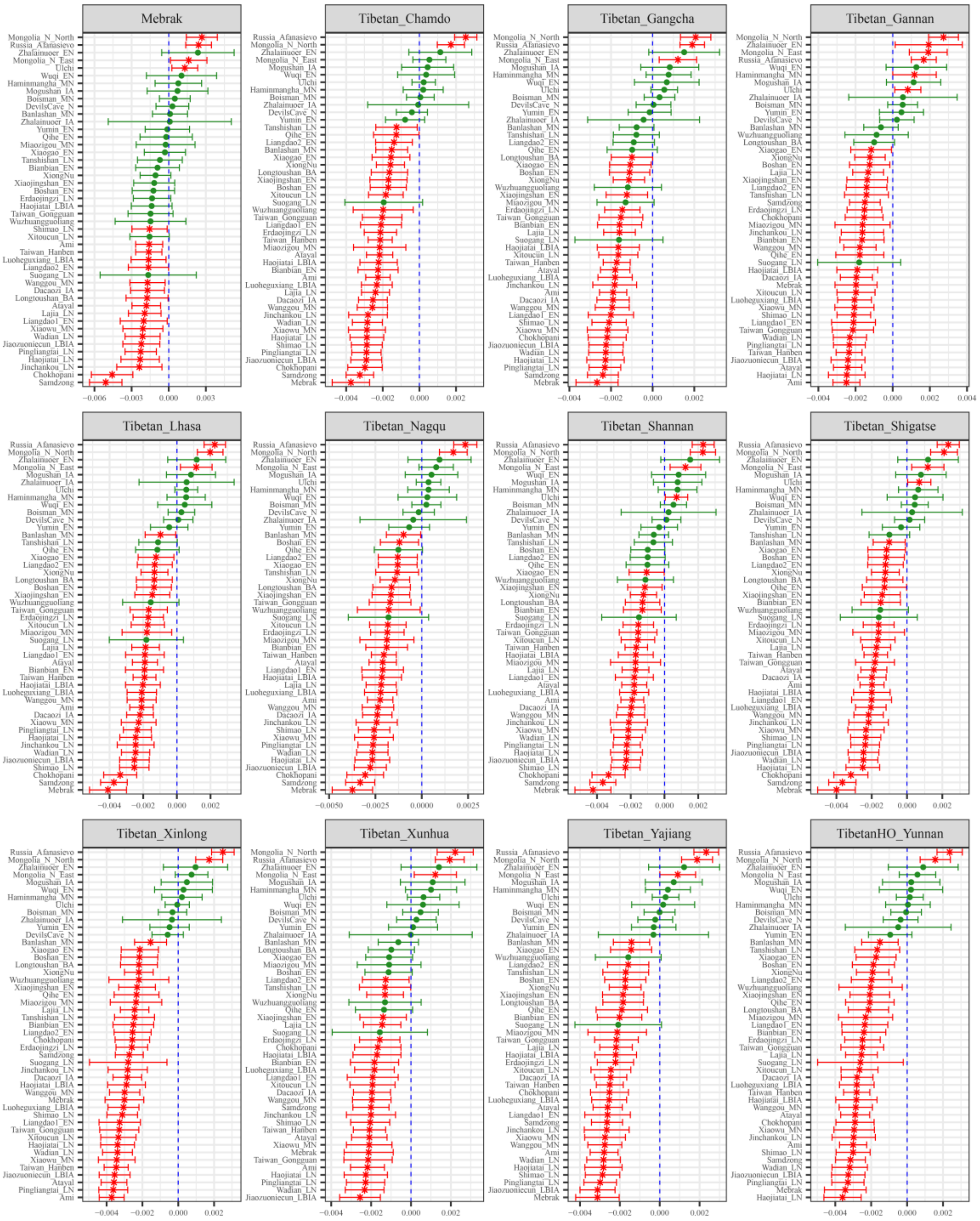

**Figure S95. Results of affinity-$f_4$ statistics for the form $f_4$*(Russia_Shamanka_EBA, Modern Tibetan; Neolithic to Historic East Asians, Mbuti)* showed the additional shared derived alleles from source populations except for ancestral populations from Mongolia Plateau or Baikal Lake Region.**
Here, overlapping SNP loci included in the Affymetrix Human Origins platform among four analyzed populations were used. We used the genetic variation of Mbuti as the outgroup. Red asterisk point meant the significant value (Absolute value of Z-scores larger than three or equal to three) observed in the symmetry-$f_4$ statistics and green circle point denoted the non-significant $f_4$-statistic values (Absolute value of Z-scores less than three). All ancient East Asians were listed along the Y-axis and $f_4$ values were labeled along the X-axis. All results were faceted or grouped via Tibetan populations. Significant negative $f_4$ values indicated that the third population shared more alleles with the second population, and also means Tibetan obtained additional gene flow from the third source population (or related populations). And the significant positive $f_4$ value indicated that the third population shared more derived alleles with the first population. The value of $f_4$-statistics equal to zero was marked as the blue dash line. The bar indicated three standard errors.

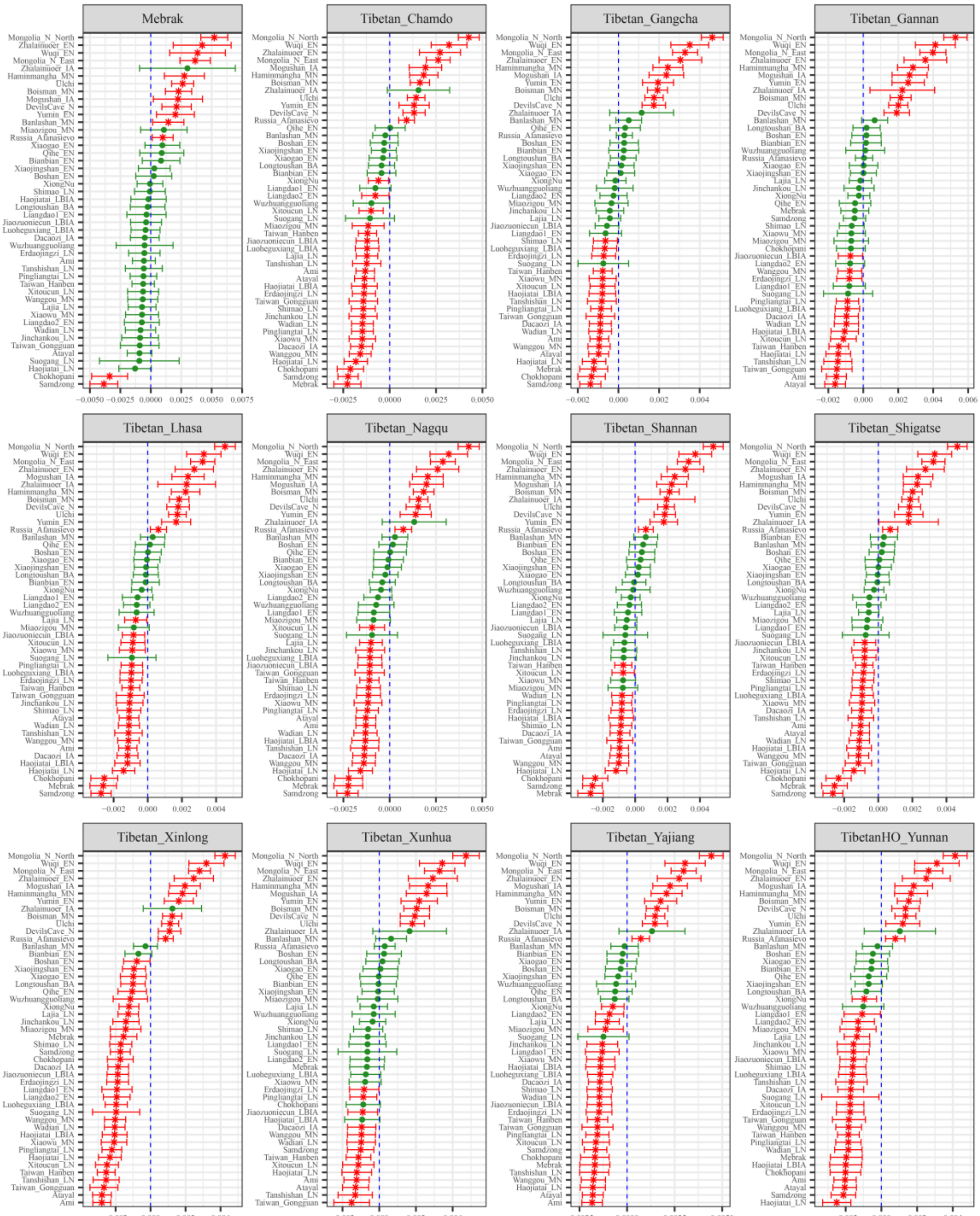

**Figure S96. Results of affinity-$f_4$ statistics for the form $f_4$*(Russia_Shamanka_Eneolithic, Modern Tibetan; Neolithic to Historic East Asians, Mbuti)* showed the additional shared derived alleles from source populations except for ancestral populations from Mongolia Plateau or Baikal Lake Region.** Here, overlapping SNP loci included in the Affymetrix Human Origins platform among four analyzed populations were used. We used the genetic variation of Mbuti as the outgroup. Red asterisk point meant the significant value (Absolute value of Z-scores larger than three or equal to three) observed in the symmetry-$f_4$ statistics and green circle point denoted the non-significant $f_4$-statistic values (Absolute value of Z-scores less than three). All ancient East Asians were listed along the Y-axis and $f_4$ values were labeled along the X-axis. All results were faceted or grouped via Tibetan populations. Significant negative $f_4$ values indicated that the third population shared more alleles with the second population, and also means Tibetan obtained additional gene flow from the third source population (or related populations). And the significant positive $f_4$ value indicated that the third population shared more derived alleles with the first population. The value of $f_4$-statistics equal to zero was marked as the blue dash line. The bar indicated three standard errors.

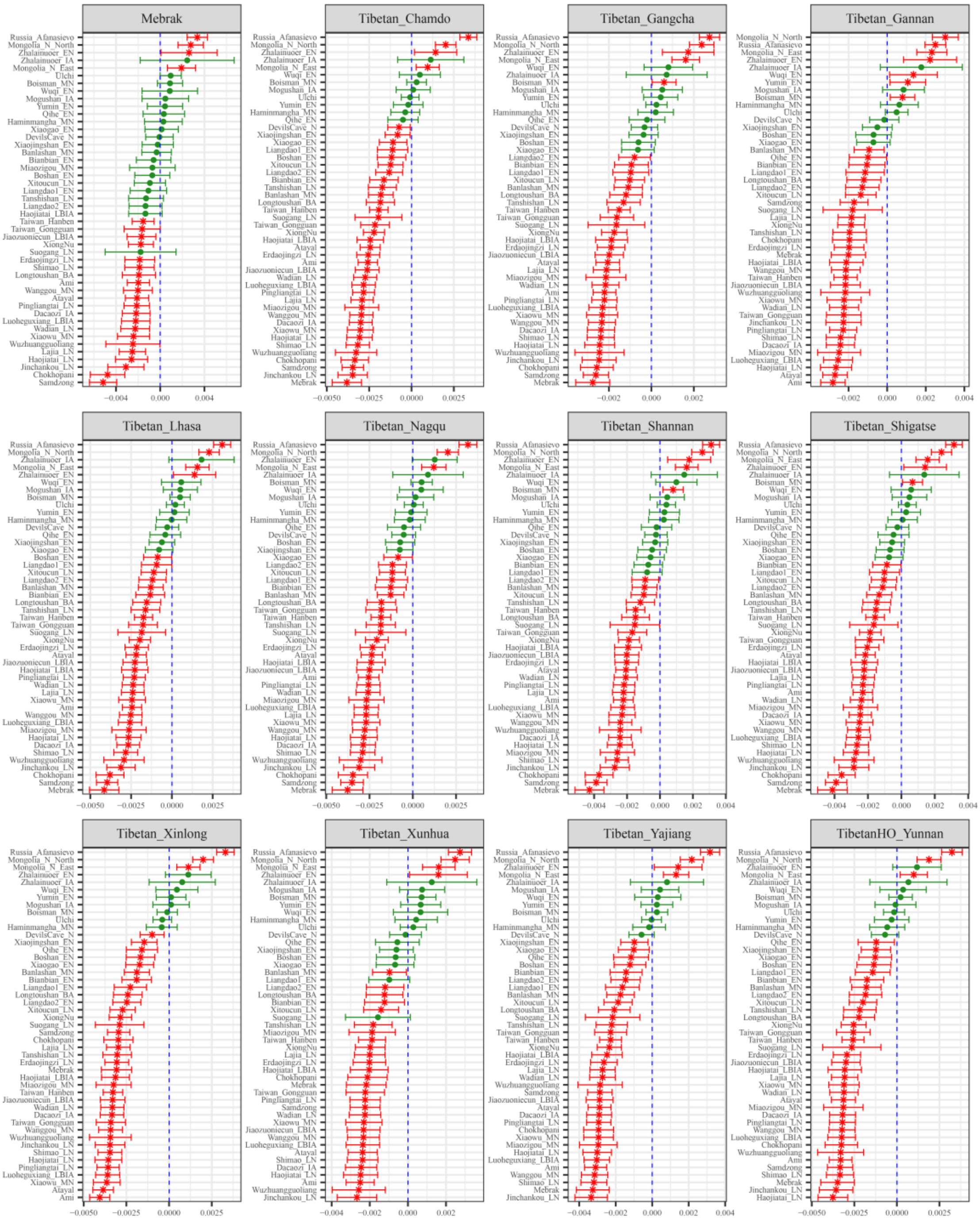

**Figure S97. Results of affinity-$f_4$ statistics for the form $f_4$*(Russia_UstBelaya_Angara, Modern Tibetan; Neolithic to Historic East Asians, Mbuti)* showed the additional shared derived alleles from source populations except for ancestral populations from Mongolia Plateau or Baikal Lake Region.** Here, overlapping SNP loci included in the Affymetrix Human Origins platform among four analyzed populations were used. We used the genetic variation of Mbuti as the outgroup. Red asterisk point meant the significant value (Absolute value of Z-scores larger than three or equal to three) observed in the symmetry-$f_4$ statistics and green circle point denoted the non-significant $f_4$-statistic values (Absolute value of Z-scores less than three). All ancient East Asians were listed along the Y-axis and $f_4$ values were labeled along the X-axis. All results were faceted or grouped via Tibetan populations. Significant negative $f_4$ values indicated that the third population shared more alleles with the second population, and also means Tibetan obtained additional gene flow from the third source population (or related populations). And the significant positive $f_4$ value indicated that the third population shared more derived alleles with the first population. The value of $f_4$-statistics equal to zero was marked as the blue dash line. The bar indicated three standard errors.

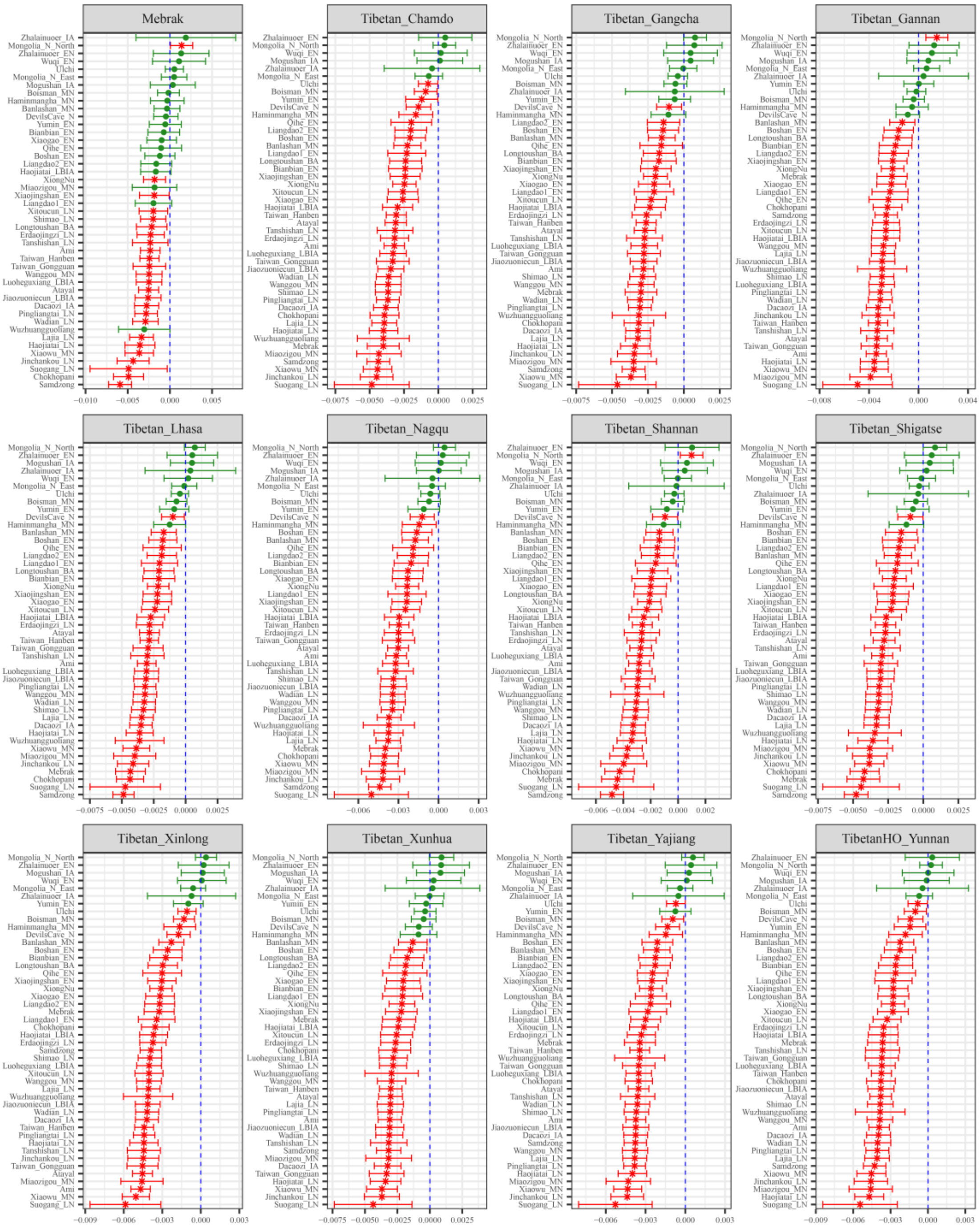

**Figure S98. Results of affinity-$f_4$ statistics for the form $f_4$*(Russia_UstIda_EBA, Modern Tibetan; Neolithic to Historic East Asians, Mbuti)* showed the additional shared derived alleles from source populations except for ancestral populations from Mongolia Plateau or Baikal Lake Region.**
Here, overlapping SNP loci included in the Affymetrix Human Origins platform among four analyzed populations were used. We used the genetic variation of Mbuti as the outgroup. Red asterisk point meant the significant value (Absolute value of Z-scores larger than three or equal to three) observed in the symmetry-$f_4$ statistics and green circle point denoted the non-significant $f_4$-statistic values (Absolute value of Z-scores less than three). All ancient East Asians were listed along the Y-axis and $f_4$ values were labeled along the X-axis. All results were faceted or grouped via Tibetan populations. Significant negative $f_4$ values indicated that the third population shared more alleles with the second population, and also means Tibetan obtained additional gene flow from the third source population (or related populations). And the significant positive $f_4$ value indicated that the third population shared more derived alleles with the first population. The value of $f_4$-statistics equal to zero was marked as the blue dash line.

The bar indicated three standard errors.

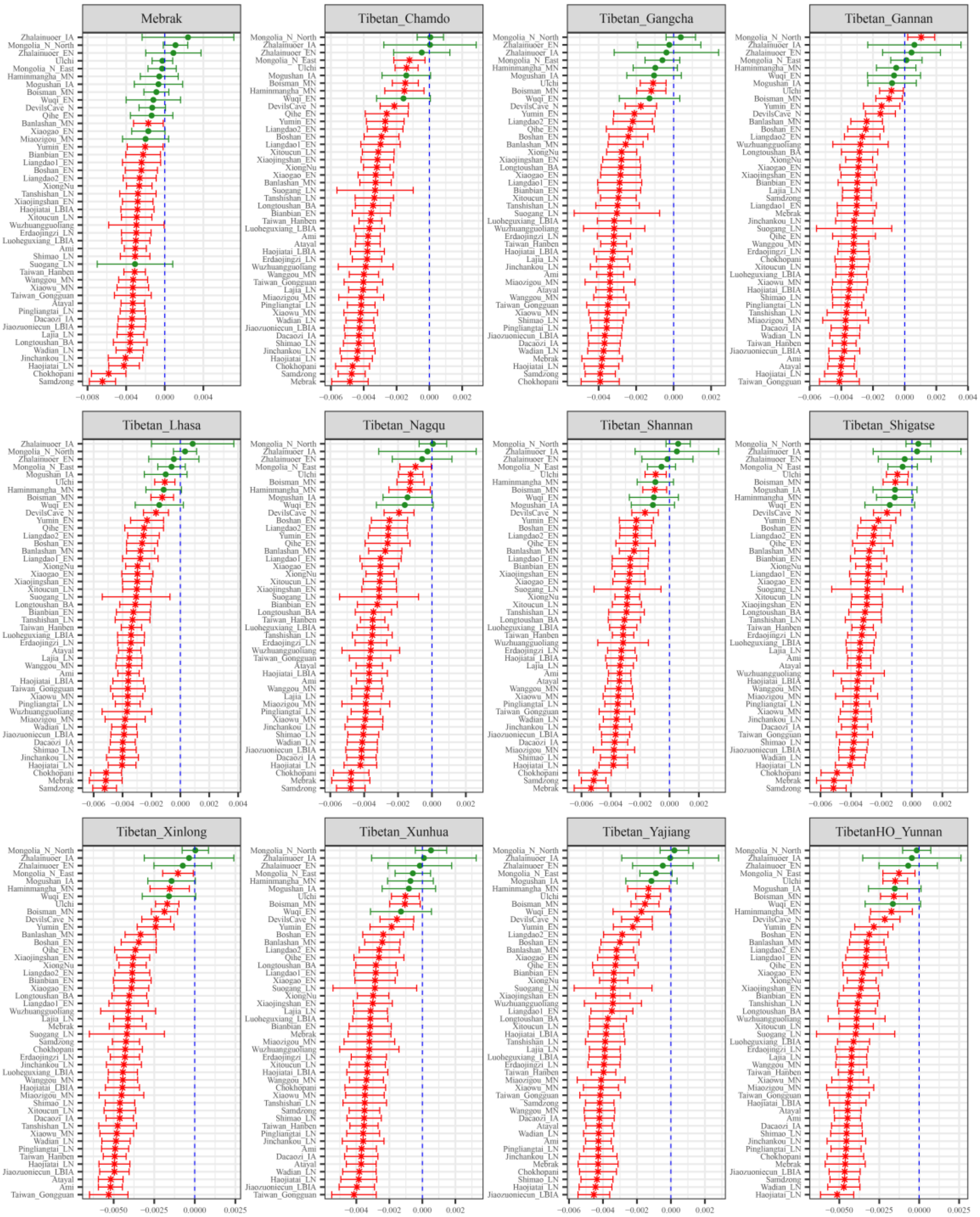

**Figure S99. Results of affinity-$f_4$ statistics for the form $f_4$*(Russia_UstIda_LN, Modern Tibetan; Neolithic to Historic East Asians, Mbuti)* showed the additional shared derived alleles from source populations except for ancestral populations from Mongolia Plateau or Baikal Lake Region.** Here, overlapping SNP loci included in the Affymetrix Human Origins platform among four analyzed populations were used. We used the genetic variation of Mbuti as the outgroup. Red asterisk point meant the significant value (Absolute value of Z-scores larger than three or equal to three) observed in the symmetry-$f_4$ statistics and green circle point denoted the non-significant $f_4$-statistic values (Absolute value of Z-scores less than three). All ancient East Asians were listed along the Y-axis and $f_4$ values were labeled along the X-axis. All results were faceted or grouped via Tibetan populations. Significant negative $f_4$ values indicated that the third population shared more alleles with the second population, and also means Tibetan obtained additional gene flow from the third source population (or related populations). And the significant positive $f_4$ value indicated that the third population shared more derived alleles with the first population. The value of $f_4$-statistics equal to zero was marked as the blue dash line.

The bar indicated three standard errors.

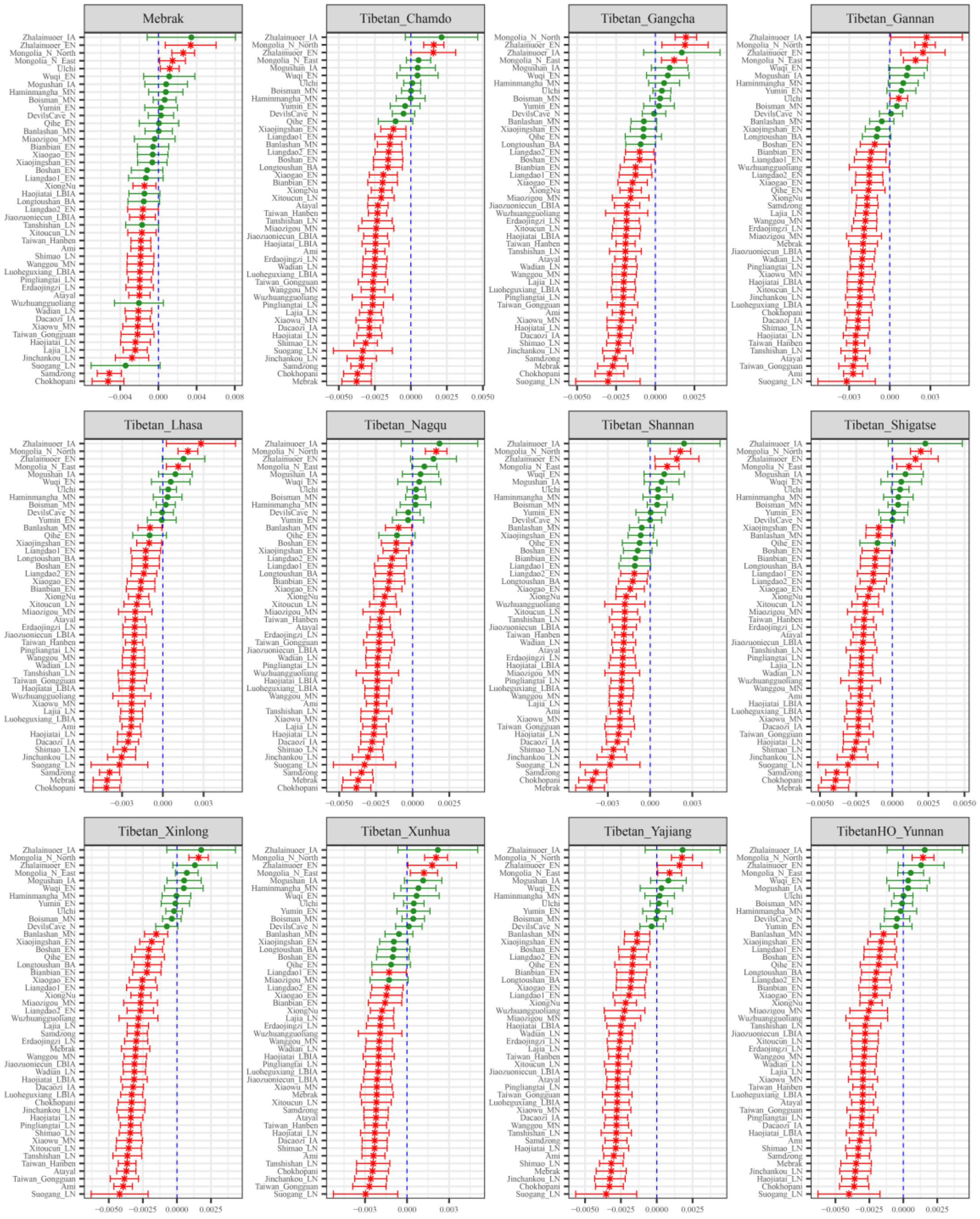

**Figure S100. Results of affinity-$f_4$ statistics for the form $f_4$*(UstBelaya_EBA, Modern Tibetan; Neolithic to Historic East Asians, Mbuti)* showed the additional shared derived alleles from source populations except for ancestral populations from Mongolia Plateau or Baikal Lake Region.**
Here, overlapping SNP loci included in the Affymetrix Human Origins platform among four analyzed populations were used. We used the genetic variation of Mbuti as the outgroup. Red asterisk point meant the significant value (Absolute value of Z-scores larger than three or equal to three) observed in the symmetry-$f_4$ statistics and green circle point denoted the non-significant $f_4$-statistic values (Absolute value of Z-scores less than three). All ancient East Asians were listed along the Y-axis and $f_4$ values were labeled along the X-axis. All results were faceted or grouped via Tibetan populations. Significant negative $f_4$ values indicated that the third population shared more alleles with the second population, and also means Tibetan obtained additional gene flow from the third source population (or related populations). And the significant positive $f_4$ value indicated that the third population shared more derived alleles with the first population. The value of $f_4$-statistics equal to zero was marked as the blue dash line.

The bar indicated three standard errors.

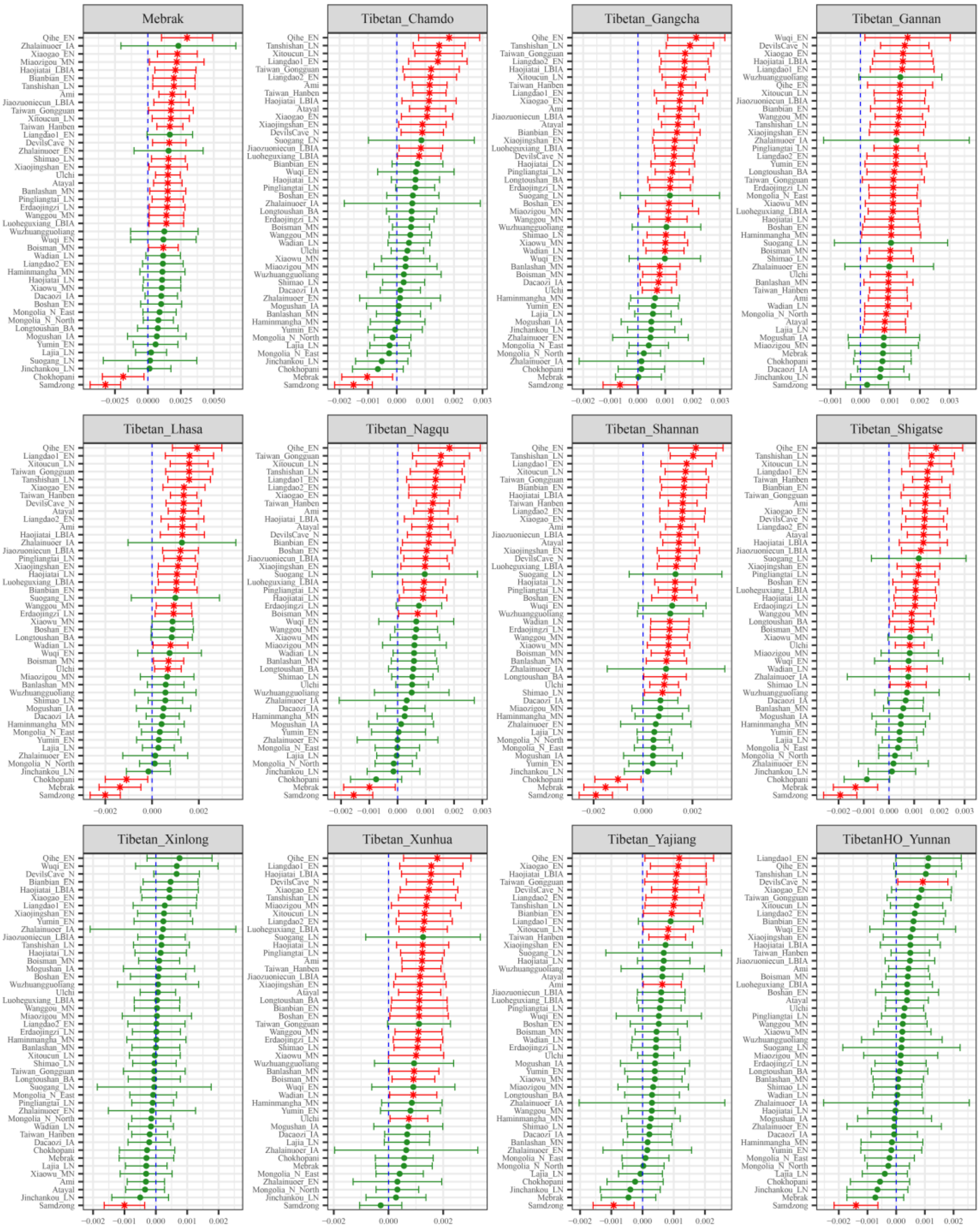

**Figure S101. Results of affinity-$f_4$ statistics for the form $f_4$*(XiongNu, Modern Tibetan; Neolithic to Historic East Asians, Mbuti)* showed the additional shared derived alleles from source populations except for ancestral populations from Mongolia Plateau or Baikal Lake Region.**
Here, overlapping SNP loci included in the Affymetrix Human Origins platform among four analyzed populations were used. We used the genetic variation of Mbuti as the outgroup. Red asterisk point meant the significant value (Absolute value of Z-scores larger than three or equal to three) observed in the symmetry-$f_4$ statistics and green circle point denoted the non-significant $f_4$-statistic values (Absolute value of Z-scores less than three). All ancient East Asians were listed along the Y-axis and $f_4$ values were labeled along the X-axis. All results were faceted or grouped via Tibetan populations. Significant negative $f_4$ values indicated that the third population shared more alleles with the second population, and also means Tibetan obtained additional gene flow from the third source population (or related populations). And the significant positive $f_4$ value indicated that the third population shared more derived alleles with the first population. The value of $f_4$-statistics equal to zero was marked as the blue dash line.

The bar indicated three standard errors.

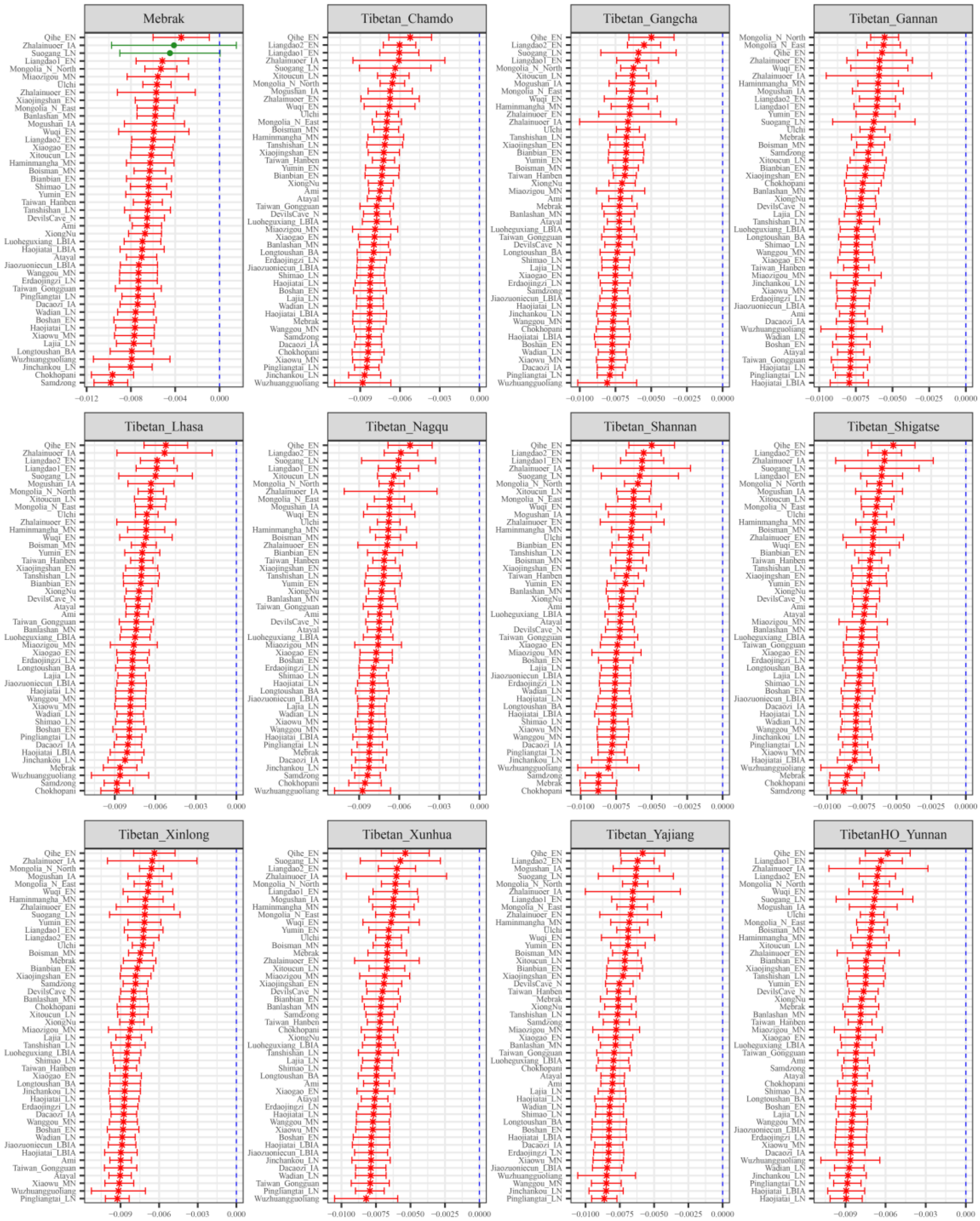

**Figure S102. Results of affinity-$f_4$ statistics for the form *$f_4$(Shirenzigou_IA, Modern Tibetan; Neolithic to Historic East Asians, Mbuti)* showed the additional shared derived alleles from source populations except for ancestral populations from Xinjiang or their related ancestral populations.** Here, overlapping SNP loci included in the Affymetrix Human Origins platform among four analyzed populations were used. We used the genetic variation of Mbuti as the outgroup. Red asterisk point meant the significant value (Absolute value of Z-scores larger than three or equal to three) observed in the symmetry-$f_4$ statistics and green circle point denoted the non-significant $f_4$-statistic values (Absolute value of Z-scores less than three). All ancient East Asians were listed along the Y-axis and $f_4$ values were labeled along the X-axis. All results were faceted or grouped via Tibetan populations. Significant negative $f_4$ values indicated that the third population shared more alleles with the second population, and also means Tibetan obtained additional gene flow from the third source population (or related populations). And the significant positive $f_4$ value indicated that the third population shared more derived alleles with the first population. The value of $f_4$-statistics equal to zero was marked as the blue dash line.

The bar indicated three standard errors.

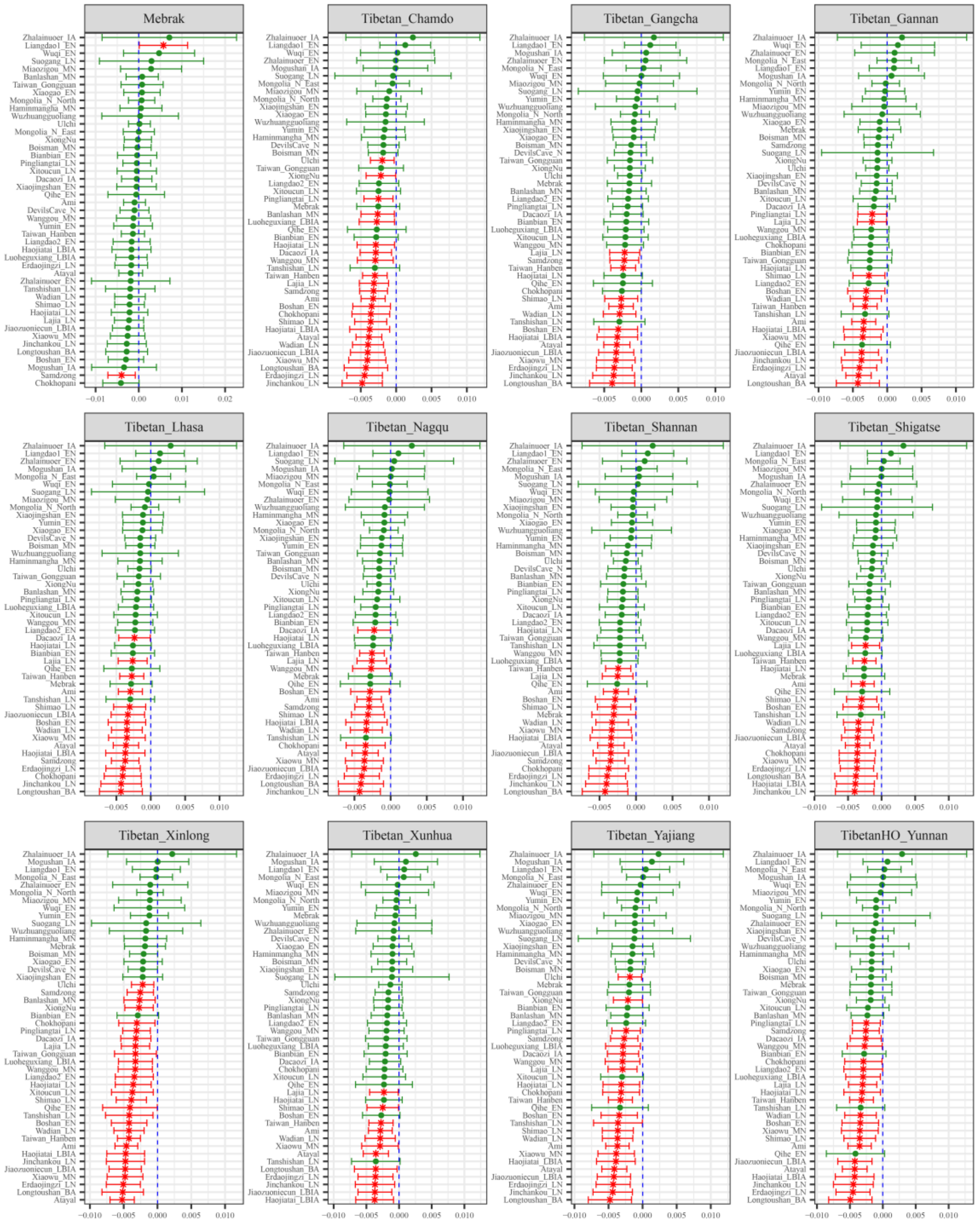

**Figure S103. Results of affinity-$f_4$ statistics for the form $f_4$*(Shirenzigou_IA_E, Modern Tibetan; Neolithic to Historic East Asians, Mbuti)* showed the additional shared derived alleles from source populations except for ancestral populations from Xinjiang or their related ancestral populations.** Here, overlapping SNP loci included in the Affymetrix Human Origins platform among four analyzed populations were used. We used the genetic variation of Mbuti as the outgroup. Red asterisk point meant the significant value (Absolute value of Z-scores larger than three or equal to three) observed in the symmetry-$f_4$ statistics and green circle point denoted the non-significant $f_4$-statistic values (Absolute value of Z-scores less than three). All ancient East Asians were listed along the Y-axis and $f_4$ values were labeled along the X-axis. All results were faceted or grouped via Tibetan populations. Significant negative $f_4$ values indicated that the third population shared more alleles with the second population, and also means Tibetan obtained additional gene flow from the third source population (or related populations). And the significant positive $f_4$ value indicated that the third population shared more derived alleles with the first population. The value of $f_4$-statistics equal to zero was marked as the blue dash line. The bar indicated three standard errors.

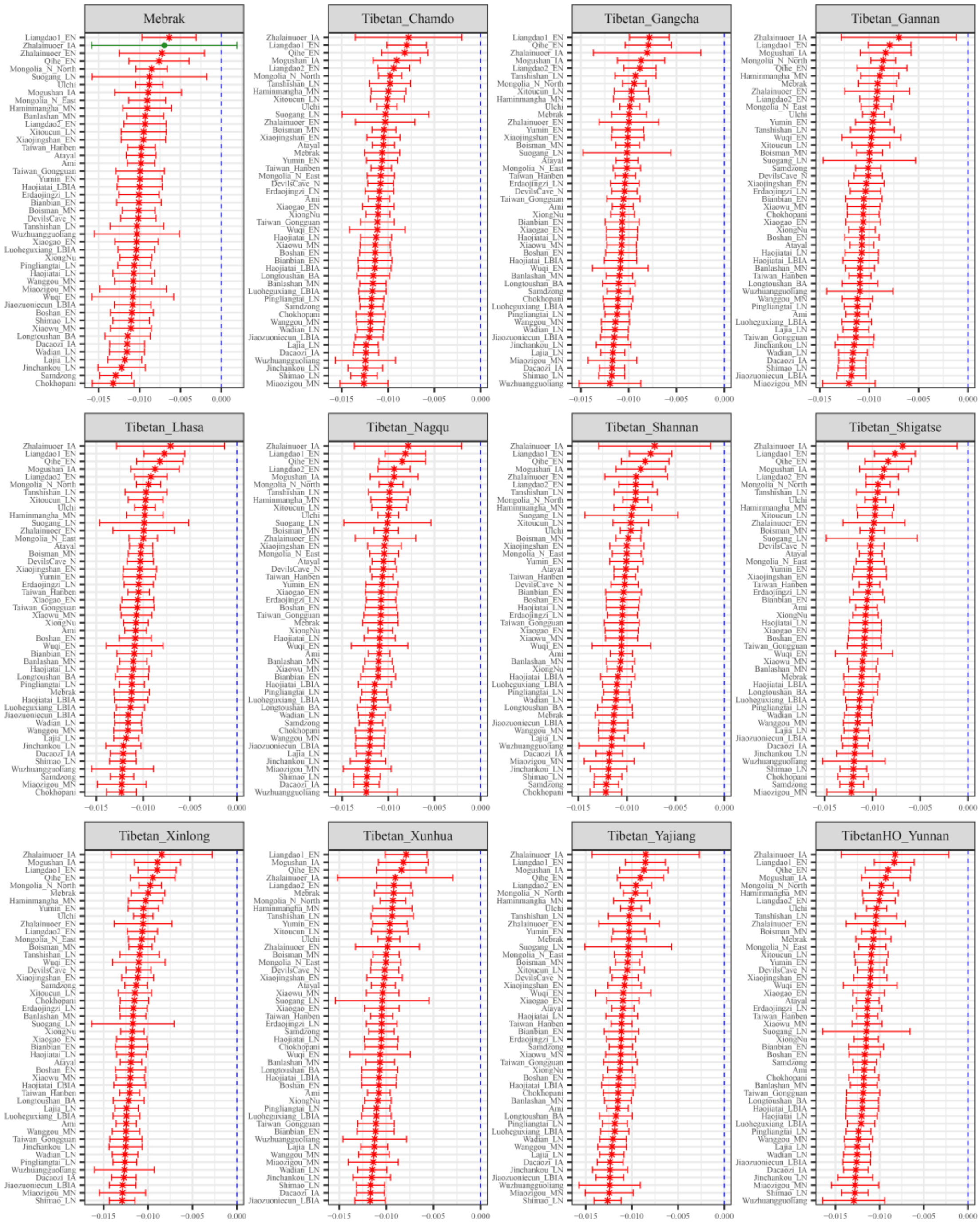

**Figure S104. Results of affinity-$f_4$ statistics for the form $f_4$*(Shirenzigou_IA_W, Modern Tibetan; Neolithic to Historic East Asians, Mbuti)* showed the additional shared derived alleles from source populations except for ancestral populations from Xinjiang or their related ancestral populations.** Here, overlapping SNP loci included in the Affymetrix Human Origins platform among four analyzed populations were used. We used the genetic variation of Mbuti as the outgroup. Red asterisk point meant the significant value (Absolute value of Z-scores larger than three or equal to three) observed in the symmetry-$f_4$ statistics and green circle point denoted the non-significant $f_4$-statistic values (Absolute value of Z-scores less than three). All ancient East Asians were listed along the Y-axis and $f_4$ values were labeled along the X-axis. All results were faceted or grouped via Tibetan populations. Significant negative $f_4$ values indicated that the third population shared more alleles with the second population, and also means Tibetan obtained additional gene flow from the third source population (or related populations). And the significant positive $f_4$ value indicated that the third population shared more derived alleles with the first population. The value of $f_4$-statistics equal to zero was marked as the blue dash line. The bar indicated three standard errors.

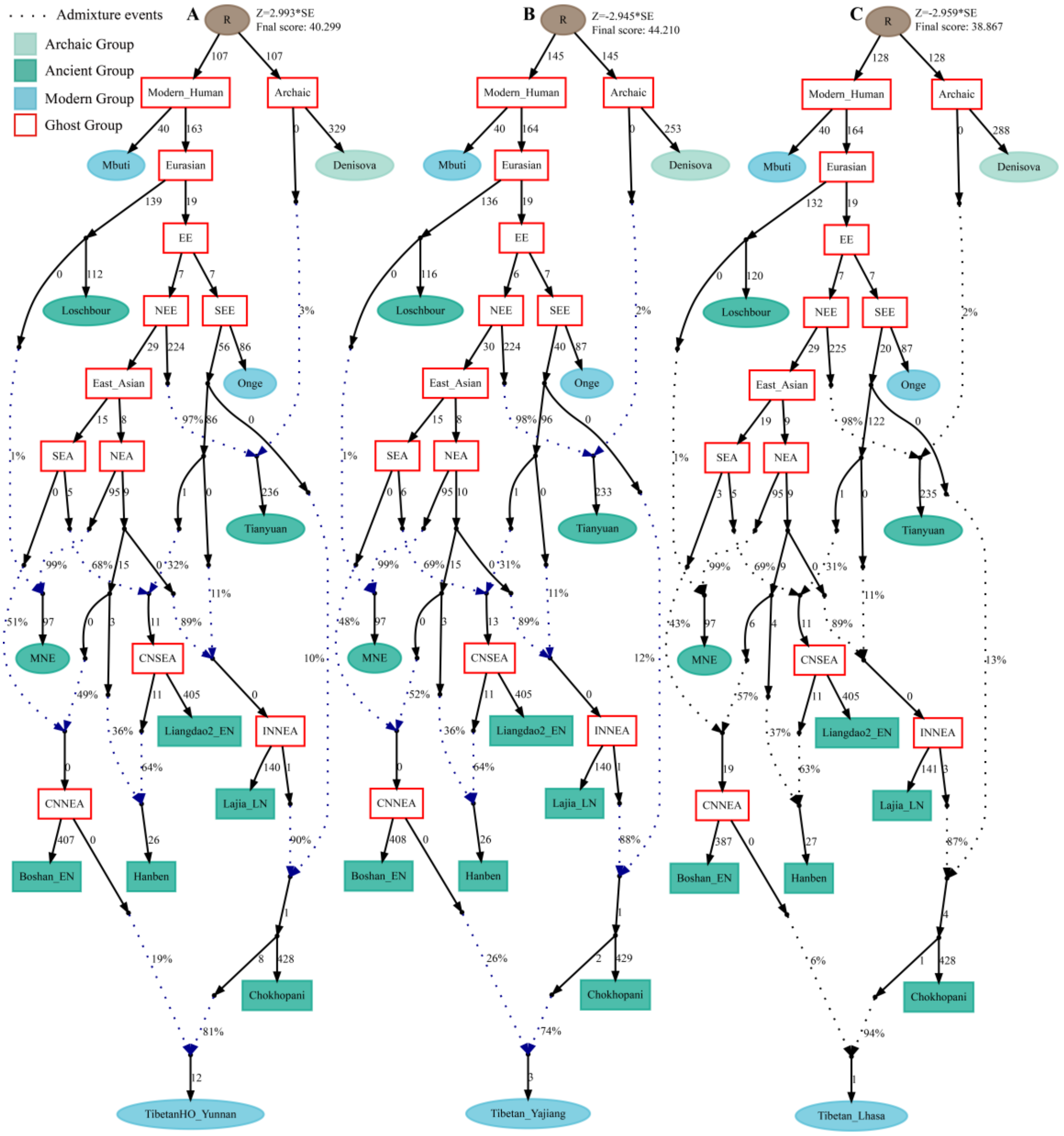

**Figure S105. Admixture graph model of modern highland and lowland Tibetans based on the Human Origin dataset using early Neolithic Boshan people as the source of the second migration into Tibet Plateau.**

Admixture history of lowland Tibetan from Yunnan (**A**), Tibetan from Yajiang (**B**) and highland Tibetan from Lhasa (**C**). Western Eurasian was represented by Loschbour. Deep southern Eurasian (SEE) and northern Eurasian (NEE) were represented by South Asian Hunter-Gatherer of Onge and 40,000-year-old Tianyuan people. East Asian was subsequently diverged as northern East Asian (NEA) and southern East Asian (SEA). Coastal Neolithic southern East Asian (CNSEA), coastal Neolithic Northern East Asian (CNNEA) and inland Neolithic northern East Asian (INNEA) were represented by Liangdao2_EN, Boshan_EN and Lajia_LN, respectively. All $f_4$-statistics of included populations are predicted to within 2.959 standard errors of their observed values. Branch lengths are given in units of 1000 times the $f_2$ drift distance (rounded to the nearest integer). Blue dotted lines denoted admixture events with admixture proportions as shown.

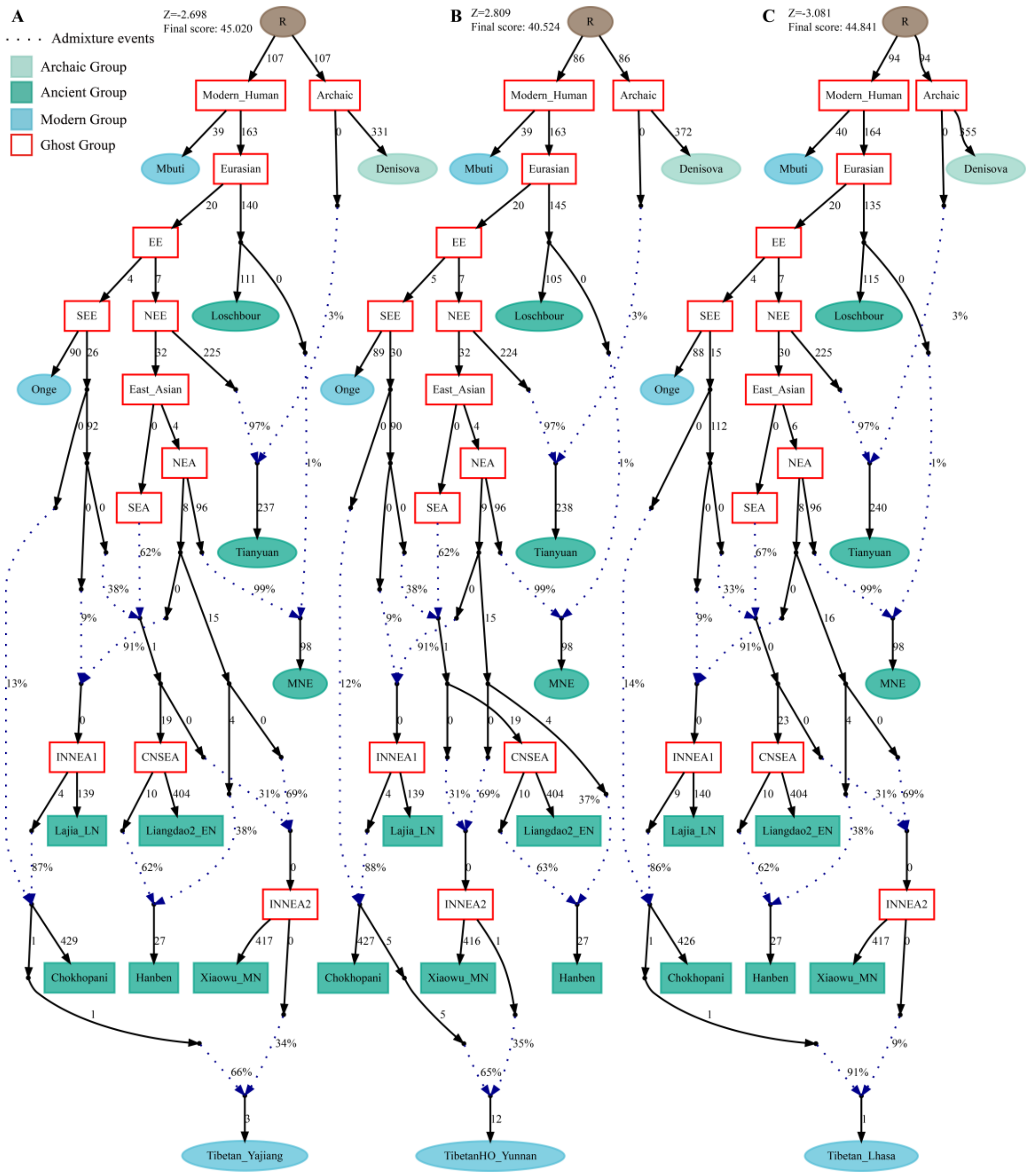

**Figure S106. Admixture graph model of modern highland and lowland Tibetans based on the Human Origin dataset using middle Neolithic Xiaowu people as the source of the second migration into Tibet Plateau.**

Admixture history of lowland Tibetan from Yunnan (**A**), Tibetan from Yajiang (**B**) and highland Tibetan from Lhasa (**C**). Western Eurasian was represented by Loschbour. Deep southern Eurasian (SEE) and northern Eurasian (NEE) were represented by South Asian Hunter-Gatherer of Onge and 40,000-year-old Tianyuan people. East Asian was subsequently diverged as northern East Asian (NEA) and southern East Asian (SEA). Coastal Neolithic southern East Asian (CNSEA), Inland Neolithic northern East Asian2 (INNEA2) and inland Neolithic northern East Asian1 (INNEA1) were represented by Liangdao2_EN, Xiaowu_MN and Lajia_LN, respectively. All $f_4$-statistics of included populations are predicted to within 3.018 standard errors of their observed values. Branch lengths are given in units of 1000 times the $f_2$ drift distance (rounded to the nearest integer). Blue dotted lines denoted admixture events with admixture proportions as shown.

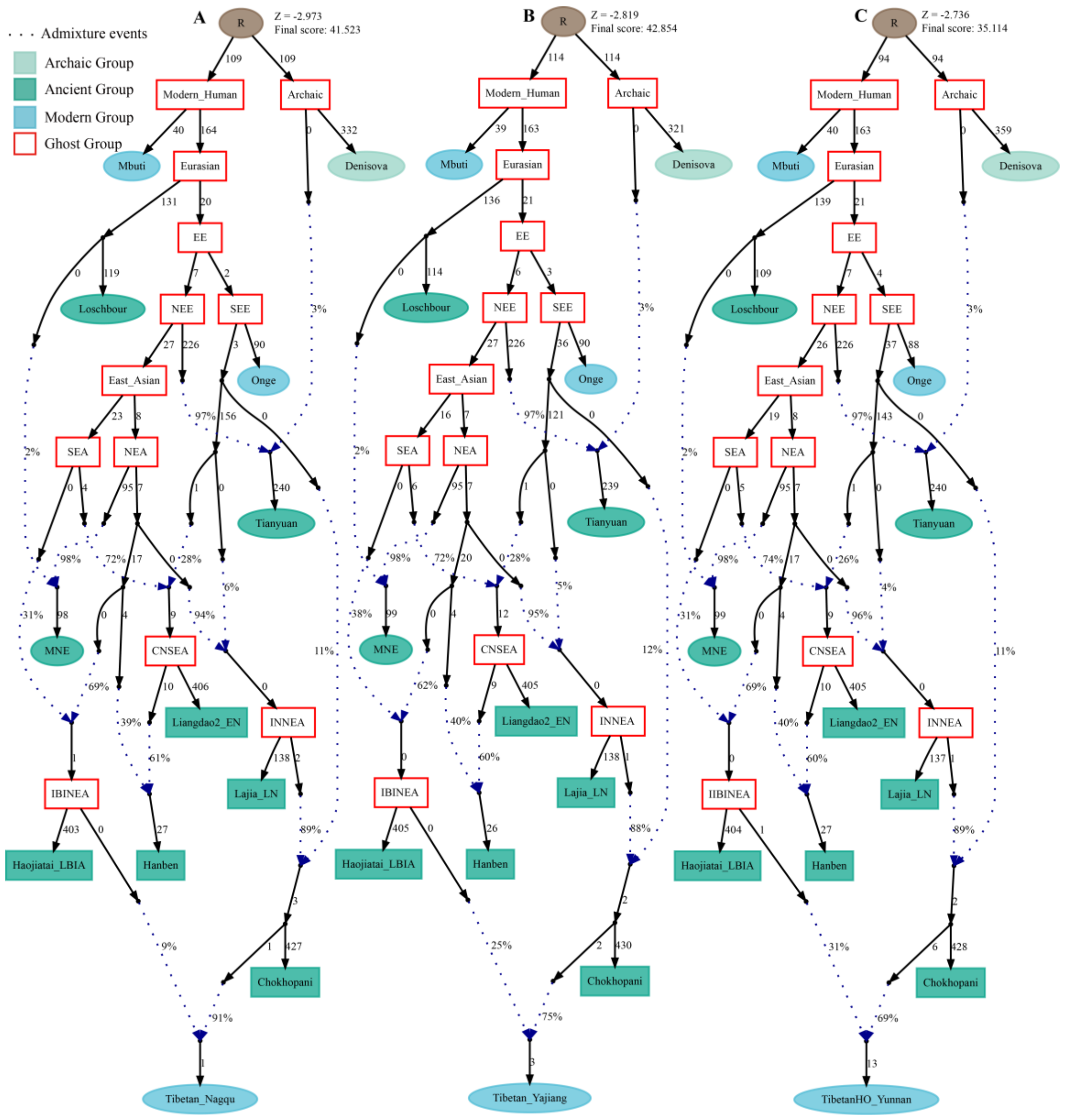

**Figure S107. Admixture graph model of modern highland and lowland Tibetans based on the Human Origin dataset using late Bronze Age to Iron Age Haojiatai people as the source of the second migration into Tibet Plateau.**

Admixture history of lowland Tibetan from Yunnan (**A**), Tibetan from Yajiang (**B**) and highland Tibetan from Lhasa (**C**). Western Eurasian was represented by Loschbour. Deep southern Eurasian (SEE) and northern Eurasian (NEE) were represented by South Asian Hunter-Gatherer of Onge and 40,000-year-old Tianyuan people. East Asian was subsequently diverged as northern East Asian (NEA) and southern East Asian (SEA). Coastal Neolithic southern East Asian (CNSEA), Inland Neolithic northern East Asian2 (INNEA2) and inland Neolithic northern East Asian1 (INNEA1) were represented by Liangdao2_EN, Haojiatai_LBIA and Lajia_LN, respectively. All $f_4$-statistics of included populations are predicted to within 2.959 standard errors of their observed values. Branch lengths are given in units of 1000 times the $f_2$ drift distance (rounded to the nearest integer). Blue dotted lines denoted admixture events with admixture proportions as shown.
